# Dynamic s-acylation of schizophrenia-linked antioxidants explicate the redox flexibility of human microglia during inflammation

**DOI:** 10.64898/2026.04.28.721236

**Authors:** Soumyadeep Mukherjee, Arushi Sharma, Prince Upadhyay, Evanka Madan, Soumya Pati, Shailja Singh

## Abstract

Cysteine proteome essentially regulates cellular adaptability during stress. Proteomic cysteines are reversibly modified to avoid loss through oxidation. Reversible cysteine-modifications beneficially preserve cellular stress-response mechanisms by regulating the transient activation of substrate proteins. Cysteine acylation (or s-acylation) is one such extensively documented reversible protein modification. Inflammation-responsive changes in s-acylation facilitate the plasticity of immune-partitioned brains by dynamically determining the ‘utilization’ of stress-buffering (substrate) cysteine-proteins in microglia, the primary immune-reactive brain cells.

However, the inflammation-responsiveness of cysteine-acylation in microglia is not elaborately measured at a large-scale. Therefore, we observed the differential (quantitative) s-acylation of proteins in cultured human (HMC3) microglia by implementing comparative MLCC-based chemoproteomic workflows. Parallelly, we characterized the molecular consequences of HMC3 cell treatment with 2-bromopalmitate, a general s-acylation inhibitor, by performing immunocytochemistry and proteomic experiments. We also identified the s-acylation-dependent microglial inflammatory responses through live spatiotemporal tracking and confocal microscopy, following treatment of microglia with PAMPs (LPS or IFNy). Finally, from our proteomic studies of postmortem human brain tissues, we confirmed major protein-causals of schizophrenia, a redox-related brain disorder, to undergo inflammation-responsive s-acylation.

This report provides a first comprehensive library of cysteine-proteins differentially processed through inflammation-responsive acylation in microglia. It also outlines the inflammation-responsive s-acylation of key NRF2-antioxidants, including peroxiredoxins (especially PRDX2) and glutathione synthetase (GSS), as a biochemical signature of early-stage inflammation and oxidative-stress in microglia. More importantly, the innovative proteo-informatic workflows designed for this research work can be reliably repurposed to identify bifunctional theragnostic protein markers of many complex disorders and diseases.

## Introduction

Environmental and internal immunogenic stimuli trigger inflammation through activation of innate immune cells (Cronkite & Strutt, 2018; Warrick et al., 2025). Immune responder cells such as neutrophils and macrophages, relay the inflammatory signals to immune-partitioned organs like the brain through the blood stream. Across the blood-brain barrier (BBB), these innate immune cells and the (humoral or chemical) inflammatory signals direct the polarization of glial cells. Inflammatory reactivity of glial cells (or gliosis) alters the homeodynamic functions of other neuronal system components including synaptic organizations and cells (Colonna & Butovsky, 2017; Hristovska & Pascual, 2016; Norden et al., 2016; Schramm & Waisman, 2022; Woodburn et al., 2021).

Particularly, events of persistent gliosis and inflammation supplement the oxidative stress states within the brain, thereby leading to bystander effects on neuronal systems like cellular excitotoxicity and neurodegeneration. Cellular homeostasis in such cases is maintained through the x_c_^-^ transporter-mediated export of glutamate in exchange of cystine (Barger et al., 2007; Ermakov et al., 2021; Wu et al., 2025). Cells can also readily absorb other chemically modified forms of cysteine such as *N*-acetylcysteine (NAC) from the periphery (McCarty & DiNicolantonio, 2015; Simpson & Oliver, 2020). Once inside the cell, these chemically modified molecules are converted to amino acid cysteine (Barger et al., 2007; McCarty & DiNicolantonio, 2015; Wu et al., 2025). Disruption of the cystine/cysteine-glutamate balance in brain elevates excitotoxicity, and accentuate symptoms like axonal shearing, brain cell necrosis, gliosis and hypoxia (Barger et al., 2007; Ermakov et al., 2021; Kigerl et al., 2012; McBean et al., 2015; Wu et al., 2025).

Cysteine is among the most infrequently found amino acids in the human proteome. Yet, it forms the conserved catalytic, regulatory and binding motifs of key oxidative stress-response proteins like antioxidants and other oxidative stress-response proteins (Castillo-Villanueva et al., 2023; Ermakov et al., 2021; Go et al., 2015; Jayaram & Krishnamurthy, 2021). These conserved cysteine residues can be reversibly inactivated by low (to mild) amounts of oxidants to form cysteine-sulfenic acid intermediates (Cys-SOH) of essential and stress-responsive proteins. The sulfenic acid protein intermediates can be further converted to inactive sulfinic acid forms (Cys-SO_2_H) or irreversibly inactivated sulfonic acid (Cys-SO_3_H) oxidative derivatives (Pace et al., 2025; Qiu et al., 2025; Rojo et al., 2014). Among these oxidative protein products, cysteine-sulfenic acid derivatives are reduced back to their functional states through disulfide bond formation with vicinal thiols, often spontaneously in microglial cells (Pace et al., 2025; Qiu et al., 2025; Rojo et al., 2014). Thus, microglia provide as excellent cell sources for the recovery and maintenance of endogenous antioxidant pools during oxidative stress. Additionally, inflammation-responsive microglial cell populations are the principal regulators of the cystine/glutamate balance (V. Singh et al., 2016). These cells are also the primary producers of both RONS free radicals (Ngo & Duennwald, 2022; Rojo et al., 2014) and antioxidants in the brain (Knudsen et al., 2026; Vilhardt et al., 2017). These factors underscore the paramount role of microglial cells and its master-regulatory signalling pathways, such as NRF2 responses, in restoration of brain health after neuroinflammatory and oxidative insults (Jayaram & Krishnamurthy, 2021; S. Singh et al., 2021; H. Zhang et al., 2024).

Within inflammatory microglia, protection of cysteines and cysteine-rich proteins from redox modifications is therefore expected to restore its homeostatic activation states and the cystine/glutamate balance. Simply, protection of cysteines within microglia would lower the cellular demands for cystine import and glutamate export, also allowing cysteine-rich stress-response proteins to function more efficiently during stress and inflammation (Barger et al., 2007). Protection of key protein thiol groups is carried out through reversible modifications like nitrosylation, cysteine(or s)-acylation and glutathiolation, which render the conserved cysteines impervious to ‘damaging’ oxidation (Qiu et al., 2025; Rojo et al., 2014). Almost all cysteine modifications can dynamically preserve redox homeostasis in microglia, and their aberrations are inherently pathognomonic for the onset and progression of microglial reactivity, inflammation and oxidative stress (Qiu et al., 2025; Rojo et al., 2014; Xin et al., 2018; Yang et al., 2016; X. Zhou et al., 2019).

S-acylation is a predominant cysteine modification, distinguished as the only, truly reversible lipid-based protein modification (Qiu et al., 2025). Apart from preventing the oxidation of functionally conserved protein thiols, S-acylation prevents a premature ‘utilization’ of stress-response proteins by steering them toward non-canonical localization and signalling fates (W. Li et al., 2025; Qiu et al., 2025). A transient deacylation of protein thiols again render them susceptible to oxidation, thereby restoring protein responses like reduction of free radical oxidants (Mesquita et al., 2024; Y. Wang et al., 2025). Hence, transient s-acylation in microglia must reflect its overall stress-responsiveness in different physiological states (Ngo & Duennwald, 2022; Rojo et al., 2014; Vilhardt et al., 2017).

Although many microglial proteins are discretely documented to undergo s-acylation in microglia, the inflammation-responsive ‘reversibility’ of this PTM had not been previously quantified. To advance the understanding of inflammation-responsive protein s-acylation in microglia, we undertook a data-intensive proteomic study of inflammation-responsive s-acylation in cultured human HMC3 microglia. We pulled down the s-acyl proteomes from resting-state and LPS/IFNy activated HMC3 microglia using the standard metabolic labelling and click chemistry techniques coupled with mass spectrometric analyses (MLCC-MS). A comparative study of these proteomes revealed 908 key microglia proteins to undergo variable s-acylation in inflammatory microglia. Subsequently, through studies of postmortem human brain autopsies, we could characterize differentially s-acylable microglia antioxidant proteins, like PRDX2 and P4HB, as conditional ‘hub’ causals of a psychotic dysfunction, schizophrenia, which is often pathologically correlated with microglial reactivity and oxidative stress (Trubalski et al., 2025). Additionally, we detected proteome level dysregulation of dynamically s-acylable antioxidants, PARK7 and PRDX5, verifying the role of redox disturbances in early onset of schizophrenia (De Simoni et al., 2013; Pap et al., 2022; Perkins et al., 2020; Tsoporis et al., 2022). Since s-acylation considerably maintains redox homeostasis through recycling and delocalization of antioxidants (Fisher, 2018; He et al., 2023; Lynes et al., 2012; Qiu et al., 2025), low levels of these proteins must suggest their utilization, plausibly through excessive deacylation in schizophrenia.

In this study, we described the cumulative impact of inflammation on the microglial cysteine-or s-acylation landscape. A comprehensive library, comprising of 908 microglial proteins undergoing differential s-acylation during inflammation, was drafted as a conclusion of this study. All the workflows and methodologies followed here, and all innovative software tools developed for this project, provide a fantastic framework for the characterization of ‘flexibly’ s-acylable proteins in primary microglia cells isolated from immediately recovered human brain autopsies. Insights gained from this investigation embellish our global research efforts to characterize the role of ‘reversible’ s-acylation in human immune responses to injuries, pathogens and diseases (Anam et al., 2022; Kumari et al., 2022).

## Methods

### 1. Preparation of an *in-silico* list of s-acylated proteins

To establish a **curated** list of high-confidence s-acylated **proteins** in humans, we recorded information from curated databases - UniprotKB and SwissPalm (filtered for proteins reported in at least 2 independent palmitoyl proteome studies) (Blanc et al., 2015; The UniProt Consortium, 2025), which was finalized on 1 October 2025 (Mattick & Amaral, 2022; The UniProt Consortium, 2025).

A catalogue of microglia-enriched proteins was separately constructed by using existing literature (Ayana et al., 2018; Patir et al., 2019) and information obtained from curated databases (Jiang et al., 2023; Meng et al., 2023; The UniProt Consortium, 2025; X. Zhang et al., 2019) to capture the s-acylable human proteins highly enriched in microglia (Supplementary Table 1A). Additionally, limma and edgeR packages were used in R programming to calculate the transcriptional prevalence of the genes encoding s-acylable proteins in cells. For this, the processed human cell type–specific gene expression (mean log2 of fragments per kilobase of transcript per million mapped reads values or FPKM) datasets were acquired from the Brain-RNAseq database (Y. Zhang et al., 2016).

Furthermore, to assess the signalling processes regulated by these s-acylated proteins in human cells, we performed extensive gene ontology (GO) analyses using the DAVID tool and the Reactome knowledgebase (release v94) (Milacic et al., 2024; Sherman et al., 2022).

### 2. HMC3 cell expansion

Labelled cultures of human microglia clone 3 (HMC3) cells were expanded in complete Eagle’s Minimal Essential Media (cEMEM) with 10% fetal bovine serum (FBS), 1× non-essential amino acids, 1 mM sodium pyruvate and 0.1% penicillin-streptomycin. HMC3 cells between passages 4 and 20 were used for all *in vitro* experiments. Careful matching of HMC3 cell passage numbers was performed for individual experiments (Mondal et al., 2024).

### 3. Cell viability tests – Trypan blue assay

To assess the viability of cells before each experiment, HMC3 microglia cells from each treatment group were counted and tested for living and dead cells using the 0.4% Trypan Blue Solution on hemocytometer chamber slides. Each of the viability tests were repeated at least 3 times to get multiple measurements. The percentage of live and dead cells were calculated within each grid relative to the total number of cells according to our previously published protocols (Mondal et al., 2024).

### 4. *In vitro* preparation of inflammatory PAMPs – A**β** fibrils and SARS-CoV2 ssRNAs

Pathogen or damage-associated molecular patterns (PAMPs or DAMPs) like Aβ fibril preparations and SARS-CoV-2 receptor-binding-domain (RBD) coding recombinant single stranded ribonucleic acids (ssRNAs) were used to trigger inflammation in HMC3 cells.

To prepare the 1 mg/ml stock solution of amyloid beta (Aβ) fibrils, HiLyte Fluor™ 488-labeled Aβ peptides (Anaspec, #AS-60479-01) were dissolved in phosphate-buffered saline (PBS) with 50 µl of 1% NH_4_OH solution (in dark). These stock solutions were diluted to 1uM working solutions in cEMEM. The Aβ peptide solutions were briefly sonicated (twice for 10 seconds with a break of 15 seconds at 30% amplitude) on ice to minimise aggregate formation. HMC3 cells from all treatment groups were incubated with 1uM Aβ peptide solutions. Aβ-treated cells (2.5*10^4^ cells) were either sub-cultured in 35mm dishes (cellvis, # D35-20-1.5P) for confocal-microscopic live cell imaging, or in a six-well plate (with 1*10^5^ cells) for RNA isolation (using techniques described in Section 5).

The recombinant SARS-CoV-2 RBD ssRNAs were transcribed *in vitro* from the PSFV3-SARS-CoV2 plasmid constructs with a gene expressing SARS-CoV-2 RBD proteins, using previously standardized protocols (Mondal et al., 2024). The MEGAscript SP6 transcription kit (Invitrogen, #AM1330) was used to perform *in vitro* transcription experiments with SpeI-HF (NEB, #R3133) linearized products of the PSFV3-SARS-CoV2-RBD vector. The PSFV3-SARS-CoV2-RBD plasmid was a gift from Dr. Milan Surjit’s laboratory in THSTI, India. The stability of the transcribed ssRNAs was increased by capping at the 5’-end using the ScriptCap m7G Capping System (CellScript, #C-SCCE0610). The capped ssRNAs were finally purified using TRIzol™ reagent (Thermo-Fisher Scientific, #15596026).

To perform the transfection of the recombinant SARS-CoV-2 RBD ssRNAs, 24-well cell culture plates pre-seeded with 5*10^4^ HMC3 cells (counted by the trypan blue assay) for 24 hours, were incubated with purified ssRNAs in solution with lipofectamine 3000 reagent. The ssRNA transfected cells were allowed to expand in cEMEM for 48 hours inside a 37°C incubator. After expansion, the successfully transfected HMC3 cells were selected based on the expression of SARS-CoV-2 RBD ssRNA protein products detected using the SARS-CoV-2 (COVID-19) specific Spike RBD antibodies [HL257]. The successfully transfected HMC3 cell populations were either sub-cultured in 6-well plates for RNA isolation or in chambered coverslips (with 4 wells) for immunofluorescence assays. The used consumables were carefully discarded in sealed autoclave bags after proper treatment with sodium hypochlorite.

### 5. Gene expression studies

To evaluate the effects of PAMPs treatment on the gene expression dynamics of HMC3 microglia, standard real time quantitative PCR (RT-qPCR) methodologies were followed.

Firstly, the total RNA was isolated from treatment and control groups of HMC3 cells. For this, six-well plates preseeded with 1*10^5^ HMC3 cells. After 24 hours of expansion, untreated HMC3 and HMC3 cells treated with either 1μM Aβ peptides, 1μg SARS-CoV-2 ssRNAs or 500ng/mLPS+20ng/mLIFNy, were incubated for another 12 hours in a 37°C. Total RNA content was isolated from these cell populations using the TRIzol™ reagent as per manufacturer’s instructions.

Finally, RT-PCR experiments were performed with the isolated RNA samples. Complementary deoxyribonucleic acids (cDNAs) were synthesised using the high-capacity cDNA reverse transcription kit (Applied Biosystems, #4368814). Freshly synthesised cDNA templates were resuspended in the Powerup SYBR green master mix (Applied Biosystems, #A25742) with gene-specific primer pairs. RT-qPCR experiments were performed with this cocktail using the Applied Biosystems™ StepOnePlus™ Real-Time PCR System (Applied Biosystems, Thermo-Fisher Scientific). At least 3 technical replicate reactions were set up for each gene of interest. Fold changes (2^−ΔΔCt^) were calculated after normalizing with the readouts of housekeeping gene 18S in Microsoft Excel. The standard error of mean (SEM) and the bar plots were plotted in R Studio using R programming. All the gene specific primer pairs have been described in a tabular format (Materials Table).

### 6. Cytotoxicity assay

The cytotoxicity of 2-bromopalmitate (2-BMP or 2-BP or 2BP, used interchangeably) was evaluated before proceeding for the treatment of HMC3 cells. For these cytotoxicity assays, 100mM stock solutions of 2-BMP were prepared in dimethyl sulphoxide (DMSO; SRL, #24075). Varying working concentrations of 2-BMP (0 to 100 μM and 150 μM) were prepared from the stock solution in cEMEM. These dilutions were systematically added to 96-well plates pre-seeded (for 24 hours) with approximately 4*10^3^ HMC3 cells/well. Additionally, a 0.15% DMSO solution in cEMEM (same volume in 150 μM 2-BMP) was added to the control wells to nullify the solvent toxicity. These cell culture plates were allowed to stand in a CO_2_ incubator at 37°C for approximately 18 hours. After 18 hours, the treatment media were replaced with 0.5 mg/mL MTT (3-(4,5-dimethylthiazol-2-yl)-2,5-diphenyl-2H-tetrazolium bromide) solution in 1x phosphate buffer saline (PBS). The HMC3 cell cultures were incubated for 3 hours with the MTT solution, after which the excess solutions were removed from each well. The formazan crystals, visible after the removal of excess (liquid) MTT, were resuspended in DMSO. To ensure complete dissolution, formazan crystals were incubated with 200μL DMSO in a 37°C incubator for 30 minutes more. The absorbance values were recorded in a plate reader (Biorad) at 570 nm wavelength. Replicate absorbance values were used for plotting errors (SEM), cytotoxic concentrations (CC50), and graphs in Graphpad Prism (version 9).

Although the cytotoxicity assays were performed for 2-BMP dissolved in DMSO, DMSO was substituted with ethanol as a solvent for mass spectrometric studies to reduce impedance and toxicity.

### 7. Visualization of dynamic s-acylation using click chemistry

To visualize dynamic s-acylation within HMC3 microglia, we optimized our previously published click-chemistry-based visualization methods used with *Plasmodium falciparum* cells (Anam et al., 2022). The unintended background signals (noise or artefacts) were removed after comparing the s-acylation patterns of untreated and treated HMC3 cells with the s-acylation patterns of HMC3 cells treated with 2-BMP, a general inhibitor of the eukaryotic s-acylation machinery.

Initially, 3*10^4^ HMC3 cells were seeded in chambered coverslips (Ibidi, #80426) and allowed to stand for 18 hours in standard culture conditions. The HMC3 cells were distributed into four experimental groups; 1. untreated HMC3 cells, 2. HMC3 treated with 50µM 2-BMP (nearly twice the CC50 concentration values), 3. HMC3 treated with 500ng/ml LPS and 20ng/ml IFNγ, and 4. HMC3 pretreated with 50µM 2-BMP prior to treatment with LPS and IFNγ.

The treated and untreated HMC3 were metabolically labelled with 17-octadecynoic acid (17-ODYA; Cayman chemicals #90270), a ‘biorthogonal’ alkyne-conjugated long-chain fatty acid analogue, which is incorporated innately into proteins through s-acylation processes. The cells were initially incubated for 6 hours with 10μM 17-ODYA. The ‘clickable’ alkyne side chains on 17-ODYA provided as baits for the Cu(I)-catalyzed cycloaddition-based (‘click’) tagging of s-acylated proteins in HMC3 cells.

While the metabolic labelling of s-acylated proteins was performed in live cells, their click chemistry-based tagging was performed in fixed, permeabilized and 17-ODYA incorporated HMC3 cells, which were also prestained with golgi-ER tagging BODIPY™ TR Ceramide dye solution (Invitrogen, #D7540). For click chemistry-based visualization of dynamic s-acylation processes, HMC3 cells were incubated with a reaction cocktail containing 30μM azide-conjugated Oregon Green 488 dye or OG488 (Invitrogen, #O10180), 1 mM solution of Tris (2-carboxyethyl) phosphine or TCEP (Sigma-Aldrich #75259) in 1x PBS, and 1 mM copper sulfate (CuSO_4_) for 1 hour at 37°C. The cell nuclei were subsequently stained with 4′,6-diamidino-2-phenylindole (DAPI) in the Vectashield antifade mounting media (Vector Lab, H-1200).

Our MLCC-visualization methods were assisted with confocal microscopy assisted imaging. The spectral profiles of BODIPY™ TR Ceramide dye (Excitation: 586nm/Emission: 622nm), nuclear stain DAPI (Excitation: 360nm/Emission: 460nm) and Oregon Green 488 dye (Excitation: 498nm/Emission: 526nm) were considered for capturing images with the Nikon Ti2 confocal microscope. The OG488 signals captured from all experimental groups were compared with those captured from untreated (control) groups of HMC3 cells. All analyses and image parsing tasks were performed in multiple replicates using the Fiji ImageJ software.

### 8. Immunocytochemistry to monitor the nuclear activity of NRF2

For the immunochemistry experiments, 2.5*10^4^ HMC3 cells were seeded in separate chambered coverslip with wells (Ibidi, #80426). After 18 hours of incubation under standard culture conditions (at nearly 40-50% confluency), the HMC3 cells were distributed into four experimental groups; 1. untreated HMC3, 2. HMC3 treated with 100µM 2-BMP (more than four times the CC50 value estimated with HMC3 cells), 3. HMC3 treated with 500ng/ml LPS and 20ng/ml IFNγ, 4. HMC3 pretreated with 100µM 2-BMP prior to treatment with LPS and IFNγ. A higher dosage of 2-BMP was provided to generate more prominent and stronger oxidative stress responses in HMC3 cells for morphological characterization.

All four groups of cells were fixed with 4% paraformaldehyde (PFA) for 10 minutes and washed using 1x PBS. The membranes of fixed cells were then permeabilized with 0.1% Triton X-100, after which blocking was performed by 5% BSA for 1 hour. The treated cell groups were allowed to be incubated with primary anti-human antibodies (1:100 dilution) from rabbit against the human nuclear factor erythroid 2-related factor 2 (NFE2L2 or NRF2; Affinity, #AF7904) protein for 3 hours at room temperature in a humid chamber. Three successive wash steps were performed with 1x PBS to remove excess antibodies. FITC conjugated secondary anti-rabbit (Bio-RAD #STAR34B) antibodies (1:200 dilution) were then added to the respective wells. The coverslip slides were allowed to stand for 90 minutes in the same humid chamber at room temperature. Then, the wells were washed thrice using 1x PBS and then mounted using the Vectashield antifade mounting media with DAPI (Vector Lab, H-1200). Images were captured using the Nikon Ti2 Confocal microscope and analysed by Fiji ImageJ software.

### 9. Preparation of cell lysates for proteomics studies

To evaluate the proteomics-level changes induced by the treatment of HMC3 cells, total proteins were isolated from RIPA buffer lysates of control HMC3, 2-BMP-treated HMC3, LPS and IFNy–treated HMC3, and HMC3 cells treated with 2-BMP, LPS and IFNy.

After BCA estimation, 1mg of proteins were considered for the *in solution* trypsinization process. The lysate samples with 1mg of proteins were purified using chloroform-methanol precipitation methods. The purified pellets were resuspended in a 6M urea solution in 50mMTris-HCl (pH8.0) for initial denaturation. The samples were then reduced with 10mM 1,4-dithio-DL-threitol (DTT) solution in 50mMTris-HCl (pH8.0) for 1 hour at room temperature in dark. Alkylation was performed using 30mM iodoacetamide, again for 1 hour at room temperature in dark. Excess (unreacted) iodoacetamide were removed by incubating the samples with 30mM DTT for another 1 hour at room temperature in dark. Finally, the samples were diluted (10x) in a 1mM CaCl_2_ solution (pH=7.6) in 50mMTris-HCl to reduce the urea concentration. MS-grade porcine trypsin (G-biosciences, #786-245) was added in a 1:50 ratio (w/w trypsin:protein) to the diluted samples. All the samples were allowed to stand in a 37°C water bath for approximately 20 hours. The samples were acidified with formic acid until a pH of 3-4 was achieved on a pH strip.

The final peptide mixtures were normalized by BCA assay performed using the bicinchoninic acid (BCA) protein assay as per the manufacturer’s protocols and instructions (Sigma-Aldrich, #QPBCA-1KT) before proceeding for MS. Statistical significance was substantiated through proteomic analysis of 3 independent biological replicate samples from each of the 4 HMC3 treatment groups. Analytical precision was ensured by considering 3 independent technical repeats of each biological replicate sample.

### 10. Preparation of tissue lysates for proteomics studies

To study the proteomic changes related to schizophrenia, schizophrenia-affected and matched control human brain tissues were collected from the Human Brain Tissue Repository for Neurobiological Studies (HBTR) at National Institute of Mental Health and Neurosciences, Bengaluru, through standardized collection procedures following approved ethical guidelines (Materials Table).

The tissues of interest were cut into small pieces using sterile scalpel and normalized by weight. Approximately 100mg of tissues were weighed in sterile conditions, taken inside a laminar flow cabinet, and cut into even smaller pieces before adding to 50 ml centrifuge tubes with 2ml RIPA buffer. The macerated tissues were further broken-down using a handheld tissue homogenizer on ice. After the debris were removed by centrifugation, all samples were purified using chloroform-methanol precipitation methods. Rest of the protocol was the same as followed for preparing and enriching cellular peptides for MS analyses. Like cellular peptides obtained from different treatment groups, tissue specific peptides were also normalized using BCA assay before proceeding for MS experiments. Statistical significance was validated through proteomic analysis of white matter or hippocampal tissue autopsies obtained from 3 control and 3 schizophrenia-affected individuals (biological replicates=3). 2 technical repeat runs were performed for each tissue lysate to minimize analytical errors.

### 11. MS settings and instrumentation

Sample preparation methods were optimized in-house for the proteomics-based studies. All peptide mixtures were desalted using C18 spin columns (G-biosciences, #786-930) before proceeding to MS analyses. Data-independent acquisition (DIA, SWATH-MS) was performed by the hybrid Quadrupole Time-of-Flight TripleTOF® 6600 mass spectrometer (Sciex) coupled to Eksigent NanoLC™ 425 (Sciex), according to previously published protocols (Bian et al., 2021).

Injection of peptides were performed with a 10µm SilicaTip electrospray emitter (New Objective, #FS360-20-10-N-20-C12). Tryptic peptide samples were loaded in equal amounts onto a micro trap column (5µm, 120L, 0.3*10mm; Chrom XP, #C18-CL-120A) using loading buffer (0.1% Formic acid and 2 % Acetonitrile in water) by isocratically running the system at a flow rate of 5LµL/min for 60Lminutes and on to the analytical column (3µm, 120L, 0.3*150mm; ChromXP, #3C18-CL-120).

A gradient flow was setup in a conserved flow mode using the following compositions:

1. Mobile phase A (0.1% Formic acid and 97% Acetonitrile in water) and B (0.1% Formic acid and 3% Acetonitrile in water) for initial 2 minutes.
2. Mobile phase A (0.1% Formic acid and 65% Acetonitrile in water) and B (0.1% Formic acid and 35% Acetonitrile in water) until the 40^th^ minute.
3. Mobile phase A (0.1% Formic acid and 50% Acetonitrile in water) and B (0.1% Formic acid and 50% Acetonitrile in water) for the next 5 minutes.
4. Mobile phase A (0.1% Formic acid and 20% Acetonitrile in water) and B (0.1% Formic acid and 80% Acetonitrile in water) for another 5 minutes until the 50^th^ minute.
5. Mobile phase A (0.1% Formic acid and 3% Acetonitrile in water) and B (0.1% Formic acid and 97% Acetonitrile in water) for flushing till the 60^th^ minute.

A TOF-MS scan range of 350 to 1200 m/z was applied with an accumulation time of 50 ms, followed by an MS/MS scan at 100 to 1500 m/z in high sensitivity mode with an accumulation time of 30 ms. 1161 cycles were set in a looped mode over the mass range of 100 to 1500 m/z for 1 hour with an accumulation time of 30 ms, resulting in a total cycle time of nearly 3 seconds. The MS detection data and metadata were recorded in raw read files (with ‘.wiff.scan’ and ‘.wiff’ file extension).

### 12. Generation of spectral libraries and peptide quantification

Prior to the quantification of peptides found in MS detection data, spectral ion libraries were generated in the DIA-NN (v2.2.0) tool (Demichev et al., 2020; Messner et al., 2021) using the canonical proteins listed in UniprotKB (The UniProt Consortium, 2025). The contaminants mapped with camprotR were removed. FASTA digestion settings in DIA-NN tool were carefully set to match the MS settings used for the detection of peptides in MLCC-products, cell samples and tissue samples. Quantification was subsequently performed with the raw read files obtained from the MS analyses. All replicate files from individual experiments were loaded and quantified at one go for most effective normalization. The program logs of FASTA digestion and quantification tasks performed with DIA-NN are provided as Supplementary Files 1-4.

Additionally, the Skyline (64-bit, version 25.1.0.237) tool (MacLean et al., 2010; Pino et al., 2017) was used for the detection and quantification of peptides of all known human PATs and deacylases (reviewed and unreviewed sequences downloaded from UniprotKB), which were initially not detected by DIA-NN (possibly due to low detection levels). A FASTA file including all reviewed and unreviewed proteins listed in UniprotKB were used as background proteome for this analysis. In Skyline, the peptide settings were set according to the DIA-NN settings, and the transition settings were set according to the default settings of the TripleTOF® 6600 mass spectrometer (Sciex) instrument.

### 13. Differential Expression analyses

Reliable tools, such as MS-Stats (R programming) and MetaboAnalyst (web), were used for quantifying the relative expression of peptides and proteins detected by DIA-NN and Skyline (Demichev et al., 2020; Kohler et al., 2023; MacLean et al., 2010; Pang et al., 2024, 2024; Pino et al., 2017). Additionally, the built-in MS-Stats algorithm in Skyline were used for the quantification of significantly detected peptides (absolute mass error≤5 and detection q value≤0.05) in Skyline. Gene ontology tasks were performed using the DAVID tool (Sherman et al., 2022) and the analysis tools provided by the Reactome database (Milacic et al., 2024).

### 14. MLCC-MS based enrichment studies of s-acylated proteins

Large-scale enrichment of s-acyl proteins were performed with HMC3 cells using conventional methods of metabolic labelling followed by click chemistry (MLCC) coupled with mass spectrometric analyses (Martin & Cravatt, 2009). MLCC methods used for the MS-based identification of s-acylated proteins, were optimized from the MLCC-visualization methods used (in Methods, Section 7) to detect dynamic s-acylation under a confocal microscope.

1*10^6^ HMC3 cells were firstly seeded in T-75 flasks. After approximately 20 hours of expansion in cell culture conditions, cells were either left untreated or pretreated with 50µM of 2-BMP (dissolved in ethanol) for 12 hours. 500ng/ml LPS and 20ng/ml IFNγ were subsequently added to half of the flasks from each pretreatment group.

A 20mM stock solution of 17-ODYA was diluted to 20μM in cEMEM and briefly sonicated (twice for 10 seconds with a break of 15 seconds at 30% amplitude) on ice before adding to all T-75 flasks. The T-75 flasks were incubated further in standard cell culture conditions for 12 hours. Treated and untreated cells labelled with 17-ODYA were washed, trypsinized and resuspended in 1x PBS supplemented with 2.5µM phenylmethylsulphonyl fluoride or PMSF (Sigma-Aldrich, #78830), 0.5% Triton X-100 and 1× Protease inhibitor cocktail ProteaseArrest™ (G-biosciences, #786-108) without EDTA. Cell suspensions were briefly sonicated (twice for 20 seconds with a break of 30 seconds at 30% amplitude) to obtain cell lysates for click chemistry reactions. The cell lysates were passed through the QIAshredder columns (Qiagen, #79654) before proceeding for purification of proteins with 1.5:4:4 solution of chloroform:methanol:PBS. All purified samples were then obtained as pellets by centrifugation at 4°C, which were solubilized in the initial buffer composition for lysate preparation. The protein concentrations in the purified samples were measured using the standard BCA assay (Sigma-Aldrich, #QPBCA-1KT) as described earlier. Samples carrying approximately 1.5mg of purified proteins were incubated in dark with 50μM biotin-azide (MedChemExpress, #HY-129832; solubilized in DMSO), 1mM sodium ascorbate (Sigma-Aldrich, #11140), and 1mM CuSO_4_ in 50mM HEPES solution (Sigma-Aldrich, #H3375; pH set to 7.4) on a benchtop rotator (6 rpm) for 1 hour. This was followed by one more round of purification with chloroform:methanol:PBS. The purified contents were then solubilised in 1.2% SDS in 50mM HEPES (pH=7.4). The solution was then diluted (6x) further in 50mM HEPES (pH=7.4) and were rested in a-80°C refrigerator for around 24 hours. While the samples were thawed on ice, streptavidin-agarose beads (Thermo Scientific, #20347) were washed once with 0.2% SDS in 50mM HEPES (pH=7.4), thrice with 1x PBS and thrice more with 50mM HEPES (pH=7.4) in a refrigerated centrifuge. The streptavidin-agarose beads were added to the thawed samples and rotated for 90 minutes at room temperature. Same washing procedures were repeated for the bound beads. A final wash was performed with 50mM Tris (pH=8).

The beads recovered from all the samples were resuspended in 6M urea solution in 50mM Tris (pH=8). The beads were then incubated with 10mM TCEP (pH=7, neutralized by KOH) for 30 minutes at room temperature. The samples were further incubated for 1 hour in dark with 30mM iodoacetamide in 50mM Tris (pH=8). The solution was diluted (10x) in a 1mM calcium chloride (CaCl_2_) solution in 50mM Tris (pH=8) before proceeding for trypsin digestion. Mass-spectrometry (MS) grade porcine trypsin (G-biosciences, #786-245) was added in a 1:50 ratio (w/w trypsin:protein) to the reduced solutions containing the beads. The samples were allowed to stand in a 37°C water bath for approximately 20 hours. The supernatants were collected on the next day and concentrated using a vacuum-coupled Concentrator Plus instrument (Eppendorf). The minimal protein pellets were resuspended in 50mM Tris (pH=8.0; volume adjusted according to the pellet size) and acidified with 5% formic acid. The final peptide mixtures were normalized by BCA assay before proceeding for MS. The peptides enriched from three different pulldown tasks were pooled and the entire pooling experiment was repeated four times (n=4).

The enrichment calculations were performed using the differential expression analyses methods (described in Results Section 13) in two steps: Firstly, the relative abundances of proteins were calculated by normalizing MLCC-MS outputs from each treatment group with their corresponding biotin-azide untreated controls; secondly, the normalized MLCC-MS protein abundance values from untreated and ‘activated’ HMC3 cells were compared with MLCC-MS protein abundance values from ‘inhibited’ HMC3 cells to receive the final enrichment results. Individual proteins detected in MLCC-MS enrichment results were subsequently assigned a confidence grade (from 1 to 4, with a value of 1 assigned to MLCC-MS-detected proteins which could not be documented as s-acylated after database mapping and prediction methods) based on the source of curation.

### 15. Differential Network analyses

To identify the schizophrenia-specific regulation of protein associations in human brains, differential network analyses were performed.

For the differential network analyses, bootstrapped consensus protein-protein co-expression models were predicted by integration of multiple inference methods: Differential co-expression (DiffCoExp), GENIE3, Graphical lasso (GLASSO) or Pearson correlation (PCC) in R-programming. These prediction tools independently identified different aspects of conditional relationships between the MS-quantified expression of proteins. DiffCoExp identified the differential co-expression relationships between conditions, genie3 used tree-based machine-learning methods to infer directed regulatory networks, glasso modelled the partial correlation networks to predict the direct regulatory interactions, and pcc estimated the pairwise linear correlations between normalised MS-expression values of interacting proteins.

A network chart was prepared from every weighted adjacency matrix generated by these tools during each resampling round across 1000 bootstrap iterations. This was followed by the summarisation (identification and ranking) of differentially connected protein hubs in schizophrenia. The ‘hub’ summarisation was performed in multiple steps.

Initially, the degree differences of connecting edges were computed for all protein nodes between disease and control networks within each method. Degree differences were then standardized for each method by transformation of absolute outputs into Z scores to remove scale bias and to enable direct comparison across methods. For individual proteins, the degree differences calculated by each method (within that bootstrap round) were simply compared with the mean degree differences and the standard deviation for the calculation of Z scores.

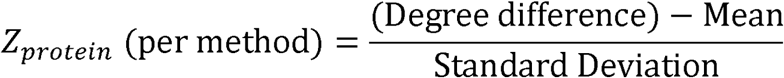

For each bootstrap, a model consensus hub score (MCS1) was calculated for each protein as the mean of standardized degree differences across all methods, capturing method-independent network connectivity changes.

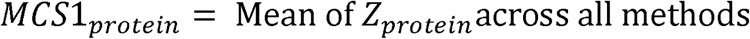

A high positive MCS1 would indicate a gain of connections predicted by an agreement of most methods. Similarly, a high negative MCS1 would indicate a loss of connections in schizophrenia. An MCS1 closer to zero would indicate more inconsistent signals.

MCS1 values were then aggregated across bootstrap iterations to estimate mean consensus hub scores (MCS2), mean effect size and variability.

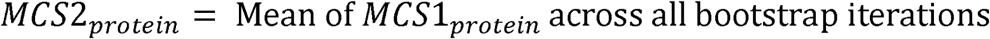

Finally, the statistical significance was assessed using Z tests, with two-sided P-values and 95% confidence intervals computed under a normal approximation.

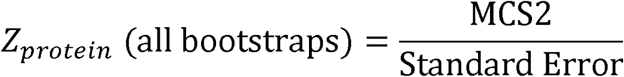

Proteins were then ranked based on statistical significance and magnitude of consensus hub scores, yielding a robust set of reproducible network hubs in schizophrenia.

The results of the differential network analyses were mapped on physical protein interaction charts obtained from curated databases to calculate the conditional metabolic network charts in control and schizophrenia brain tissues (del Toro et al., 2021; Oughtred et al., 2021; Szklarczyk et al., 2022). Specifically, the larger network charts predicted by different network inference methods were filtered for significant physical or direct PPIs obtained from curated databases. Physically or directly interacting proteins and the strength of their association were obtained from curated databases – StringDB version 12 (cutoff score = 500), Biogrid version 5 (cutoff score = 100) and IntAct (cutoff score = 0.5) (del Toro et al., 2021; Oughtred et al., 2021; Szklarczyk et al., 2022). The large tabular data from each were pre-processed using R programming. The pre-processed tables featuring PPIs with strengths above cutoff scores were stored in MS Excel. The interactions were scored either zero, one, two or three based on the number of databases recognizing it as a physical or direct interaction.

The curated protein associations of differentially s-acylated proteins (identified using a workflow described in Methods, Section 14) were thereby mapped in the protein network maps generated for schizophrenia and control autopsy samples. These ‘conditional’ protein networks were later filtered to reveal the differentially s-acylated antioxidant ‘hub’ proteins.

### 16. Statistical relevance

All data points were collected to calculate the standard error mean (SEM) using R programming and the Prism tool (Graphpad). Values justifying statistical significance were calculated based on experimental parameters using unpaired t-tests, one-way and two-way ANOVA tests. All legends, methods, statistical tests and calculations (like p-, q- and FDR values) were mentioned in the respective figures, tables, graphs and network charts. Performance of all normalization methods applied to datasets were monitored using principal component-based dimensionality reduction analyses (PCA) in R programming. The best normalization methods were selected based on the number of significant proteins returned by differential expression analyses. All results were collected with well-documented logs from at least 3 independent biological replicates to avoid statistical bias.

## Results

### 1. S-acylation is a critical component of microglia biology

#### A. *In silico* catalogue of genes coding for s-acylation substrates

To identify the metabolically ‘reversible’ inflammatory pathways, ‘s-acylable’ human proteins were recovered only from the curated databases. Since these proteins cannot be ‘s-acylated’ at all biological states, we used the term’s-acylable’ to describe these proteins herein.

We found a total of 4562 ‘s-acylable’ human proteins including 25 PATs and 7 deacylases. These ‘s-acylable’ proteins mapped to less than 19% (3584 non-redundant primary gene symbols) of all human protein-coding genes (19433 protein-coding genes recorded in Gencode version 49) (Mudge et al., 2024; The UniProt Consortium, 2025). Pathway analyses revealed a significant (Entities FDR≤0.05) association of these proteins with approximately 15% of Reactome (knowledgebase version 95) curated human reactions (organized into ∼8% of human biological pathways; Supplementary Table 1A) (Milacic et al., 2024).

The subsequent GO-based classification of these genes highlighted 78 significantly enriched biological clusters (Benjamini–Hochberg procedure derived False Discovery Rate or FDR ≤ 0.05), including the most significant cluster (cluster 1) of 1946 ‘transmembrane proteins’ (FDR ≤ 0.05). This cluster of ‘transmembrane proteins’ included 114 microglia-enriched protein markers, which mapped to Reactome database-documented cell processes such as innate immunity and stress signalling.

The Brain-RNAseq datasets indicated LYPLA1, ZDHHC2 and the amyloid beta (Aβ) phagocytic factor ZDHHC6 (Ouyang et al., 2024) as the most highly transcribed (FPKM) s-acylation enzyme genes in microglia. However, no ZDHHC-PATs or deacylases were found as significantly enriched in microglia (log2 FC ≥ 1.5, adjusted p ≤ 0.05) relative to other cell types of the brain (Table 1; Supplementary Table 1A).

**Table 1.** Relative transcription rates of s-acylation enzyme genes in human microglia with respect to other cell types of the human brain. The relative abundances were calculated using gene expression data (FPKM) provided in the Brain RNA-seq database. The log2 FC, FDR and p values were calculated using the edgeR and limma packages in R programming. ^#^

| Accession | Type | Gene | Log2 FC | P value | FDR | Mean Log2 Abundance in Microglia |
| --- | --- | --- | --- | --- | --- | --- |
| O75608 | APT | LYPLA1 | 1.39120866 | 0.008527852 | 0.052784111 | 6.03652842 |
| Q9H6R6 | PAT | ZDHHC6 | 0.786933428 | 0.031150492 | 0.143309527 | 6.93377648 |
| Q9C0B5 | PAT | ZDHHC5 | 0.778288737 | 0.010107481 | 0.060349566 | 5.27427195 |
| Q9NXF8 | PAT | ZDHHC7 | 0.713727014 | 0.050341516 | 0.203378676 | 4.81307792 |
| Q9NYG2 | PAT | ZDHHC3 | 0.473730164 | 0.109308278 | 0.246573103 | 4.44277498 |
| Q8IZN3 | PAT | ZDHHC14 | 0.452085552 | 0.300105802 | 0.430798534 | 4.19956153 |
| Q5W0Z9 | PAT | ZDHHC20 | 0.441568665 | 0.415324488 | 0.549713653 | 4.84498776 |
| O15498 | PAT | YKT6 | 0.395003259 | 0.383483762 | 0.517952672 | 4.41844767 |
| O43617 | PAT | TRAPPC3 | 0.286437985 | 0.622874658 | 0.738012479 | 5.92328348 |
| O95870 | APT | ABHD16A | 0.14729067 | 0.40639732 | 0.541195582 | 3.14599055 |
| Q9UIJ5 | PAT | ZDHHC2 | 0.113429051 | 0.857412987 | 0.910577685 | 6.35810312 |

#### **B.** Global expression dynamics of s-acylation genes during inflammation

RT-qPCR studies were performed to test the dynamic expression of microglia-enriched s-acylation enzyme genes – ZDHHC6, ZDHHC2 and ZDHHC5, and the most highly expressed deacylase LYPLA1, under the influence of PAMPs (and DAMPs) including CoV2 ssRNA and Aβ.

Although relatively steady transcription rates of s-acylation enzyme genes are reported after treatment with inflammatory agents like LPS, TNFα and IL-1β (De Kleijn et al., 2022), we could observe a significant variance of gene expression for ZDHHC6 in microglia after treatment with LPS and IFNγ (pairwise t-test p value<0.001 relative to control; Supplementary Table 1B). Simultaneously, the global gene expression of microglia-enriched deacylase PPT1 (Supplementary Table 1A) were found to be significantly altered (anova p value) in HMC3 cells treated with different PAMPs (Figure 1A, Supplementary Table 1B).

**Figure 1.**
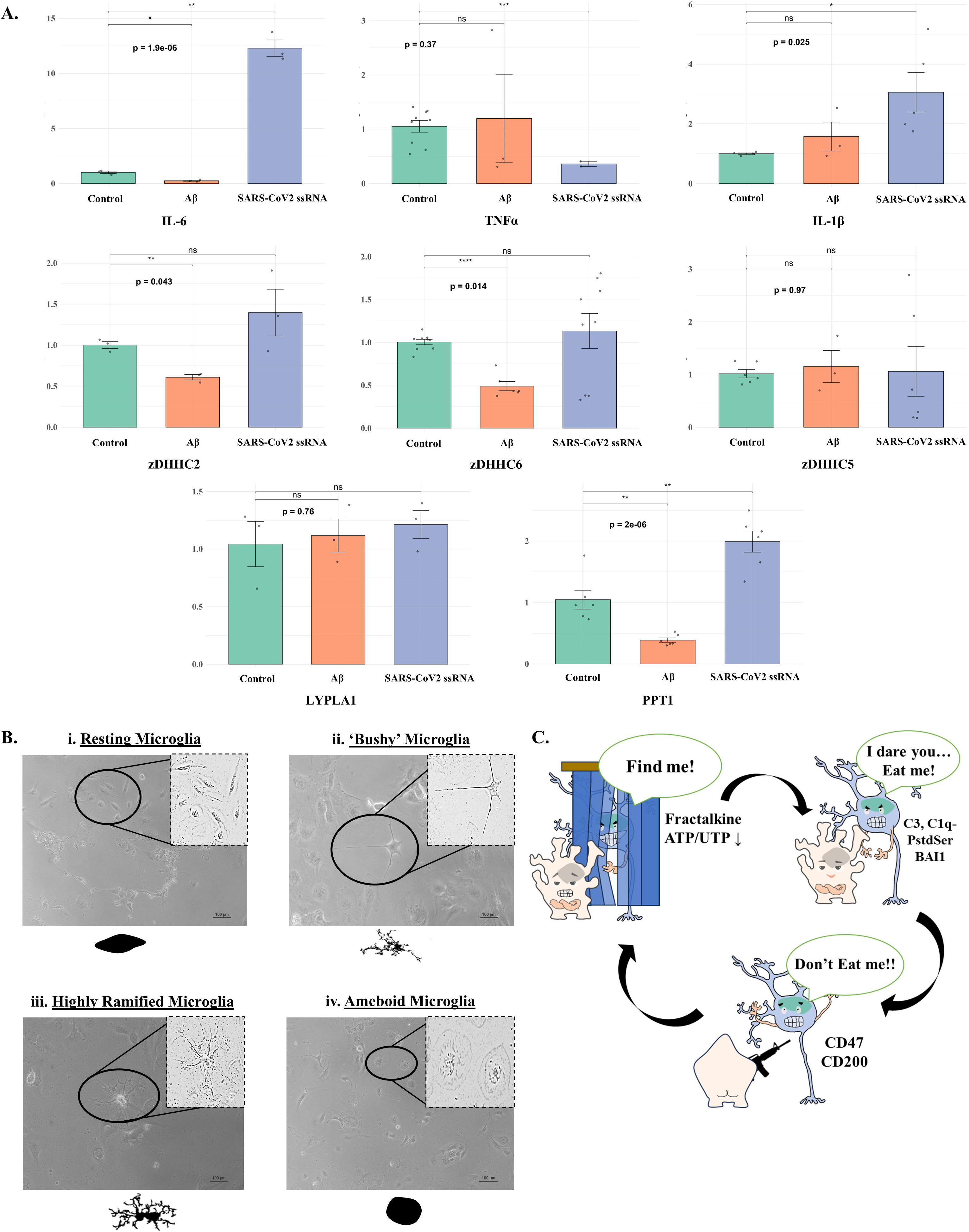
Condition-dependent transcription landscape of s-acylation enzyme genes in resting and activated microglia. **A.** Faceted bar plot panels of major s-acylation enzyme coding genes to represent their relative gene expression (X axes represent Average gene expression (Fold change or 2^-ΔΔCt^) values ± SEM) in HMC3 cells treated with Aβ (orange bars) or SARS-CoV2 ssRNA (blue bars) as compared with untreated HMC3 cells (green bars). Statistical significance between conditions is indicated as t-test p values (*p≤0.05, **p≤0.01, ***p≤0.001, ****p≤0.0001) and the global dyregulation of each gene across the treatments is indicated as a global anova p value. **B.** Representative images showcasing distinct morphological forms of activated HMC3 microglia as captured in regular cultures using a DIC microscope. **C.** A cartoon representation of neurological cues which activate microglia in physiological systems.

### 2. Immune activation of microglia involves an organellar displacement of intracellular s-acylation machineries

In the human brain, morphological dynamics of human microglia cells are dependent on the health of neuronal systems and effectively suggest inflammatory activation in stress states (Figure 1B-C). However, the extent and strength of stress-responses in the brain can be better characterized through an evaluation of molecular processes underlying these morphological shifts.

Since s-acylation processes are key upstream regulators of membrane dynamics and signalling activities of immune and inflammatory proteins in microglia, we presumed that the visualization of this process can be used to characterize the activation states of microglia. To detect the transient and spatial alterations of microglial s-acylation processes under the influence of PAMPs (LPS and IFNy), MLCC-visualization methods (previously validated in Plasmodium and human blood cells) were applied for untreated and LPS/IFNy-treated HMC3 cells (Anam et al., 2022; Kumari et al., 2022) (Figure 2A).

**Figure 2.**
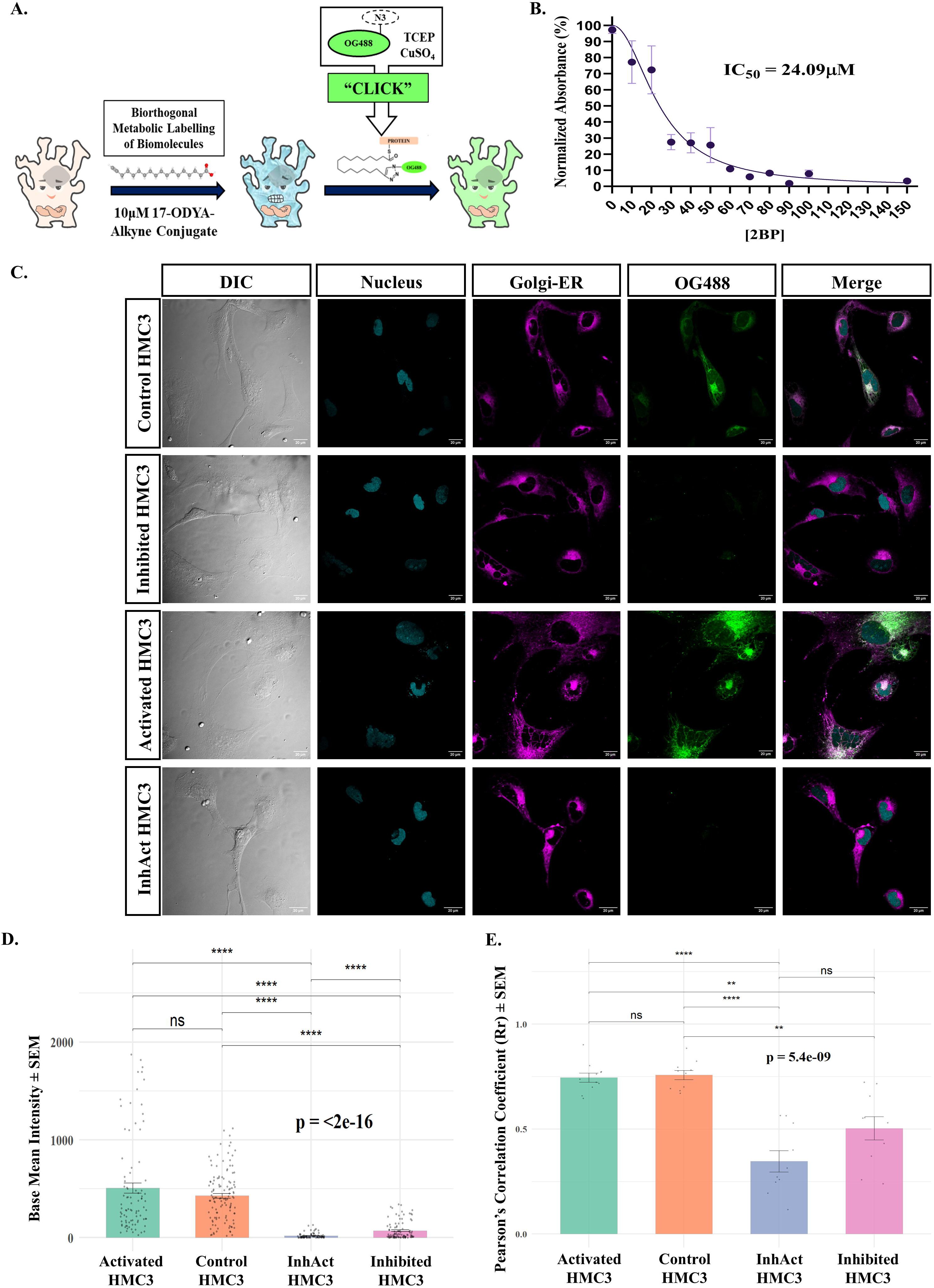
Immune activation of microglia involves a complex interplay of morphological changes and s-acylation assisted signalling. **A.** A cartoon representation of the procedures followed for click-chemistry based labelling of s-acylated biomolecules in HMC3 microglia cells. **B.** MTT assay-based calculations of CC50 values corresponding to the treatment of HMC3 cells with s-acylation inhibitor 2-BMP. **C.** Images of s-acylation in different treatment groups of HMC3 cells, as observed under a confocal microscope. 2-BMP treated HMC3 cells effectively lost maximum (nearly 83%) signals from the OG488 derivatives of 17ODYA-labelled biomolecules. **D.** Bar graphs of mean intensities recorded from control (orange), ‘activated’ (green), ‘inhibited’ (pink) and ‘InhAct’ (blue) groups of HMC3 cells. The error bars represent the Standard Error of the Mean (SEM) error calculated for each group. The mean intensity values were calculated using the OG488 signals recorded from individual cell bodies to reduce bias by avoiding the inclusion of signals from cell peripheries. **E.** Bar graphs summarising the Pearson correlation coefficient (PCC)-based colocalization of OG488-tagged s-acylable biomolecules with the BODIPY-TR-ceramide labelled Golgi-ER network in each of the ascribed HMC3 treatment groups. Through these experiments, s-acylated biomolecules (and biomolecular complexes) including proteins were observed to be enriched in the ER-Golgi network of the control HMC3 cells (PCC=0.76). Although these proteins were found to be significantly associated with the Golgi-ER network in activated HMC3 cells as well (PCC=0.75), but both types of signals were found to be dispersed or delocalized within these cells highlighting the redistribution of s-acylated biomolecules (or biomolecular complexes) during inflammation. The error bars represent the Standard Error of the Mean (SEM) error calculated for each group.

For all MLCC-visualization experiments, specificity of OG488 signals as markers of s-acylated biomolecules (or complexes) was measured in all treatment populations of HMC3 cells relative to signals recorded from HMC3 cells treated with the s-acylation inhibitor 2-BMP. The CC50 value of the s-acylation inhibitor 2-BMP was calculated to be 24.09μM for cultured HMC3 populations (Figure 2B). A treatment with 20μM 2-BMP (below CC50) could reduce OG488 signals in (‘inhibited’) HMC3 cells by 83% after 6 hours, and by more than 91% after 12 hours of treatment. This confirmed a high (roughly 86%) specificity of 2-BMP as an inhibitor of s-acylation in HMC3 cells (Figure 2C; Supplementary Table 2A).

In resting HMC3 cells, the OG488 signals showed strong colocalization with the signals recorded from BODIPY TR-Ceramide-stained Golgi-ER networks (PCC = 0.76). LPS and IFNγ stimulation of HMC3 did not significantly disturb the intensity of OG488 signals and spatial association of these signals with Golgi-ER signals (PCC = 0.74) (Figure 2C-E, Supplementary Table 2B). However, signals recorded from both s-acylated biomolecules and Golgi-ER networks were found to be dispersed within these (‘activated’) HMC3 cells (Figure 2C). These results suggested that the delocalization of s-acylated biomolecules (or complexes) and Golgi-ER network are mutually associated events in inflammatory microglia, Both intensity and the Golgi-ER colocalization of OG488 signals were most significantly reduced in (‘InhAct’) HMC3 cells stimulated with LPS and IFNγ after pretreatment with 2-BMP (PCC = 0.35) (Figure 2C-E, Supplementary Table 2B). While the ‘inhibited’ group of HMC3 cells displayed HMC3-like morphological properties, ‘InhAct’ group of cells were found to be visibly injured (Figure 2C). Therefore, we hypothesised that s-acylation must modulate fundamental molecular functions altering (inducing or buffering) stress-responsive ‘plasticity’ of microglia, and cells with dysregulated s-acylation should consequently ‘break down’ into injury states.

From these results, we concluded the delocalization (dispersion) of s-acylated biomolecules (or complexes) as a more prominent consequence of inflammation than its global dysregulation.

### 3. LPS/IFN**γ** and 2-BMP treatments induce NRF2-dependent expression of antioxidants in microglia

#### A. A comparative proteomic profiling of HMC3 cells treated with LPS, IFNγ or 2-BMP

To resolve the molecular signalling cascades and processes driving stress-responsive spatial and morphological changes in microglia, we performed comprehensive MS-based proteomic studies of control, ‘inhibited’, ‘activated’ and ‘InhAct’ HMC3 cell populations (Figure 3, Supplementary Table 3A).

**Figure 3.**
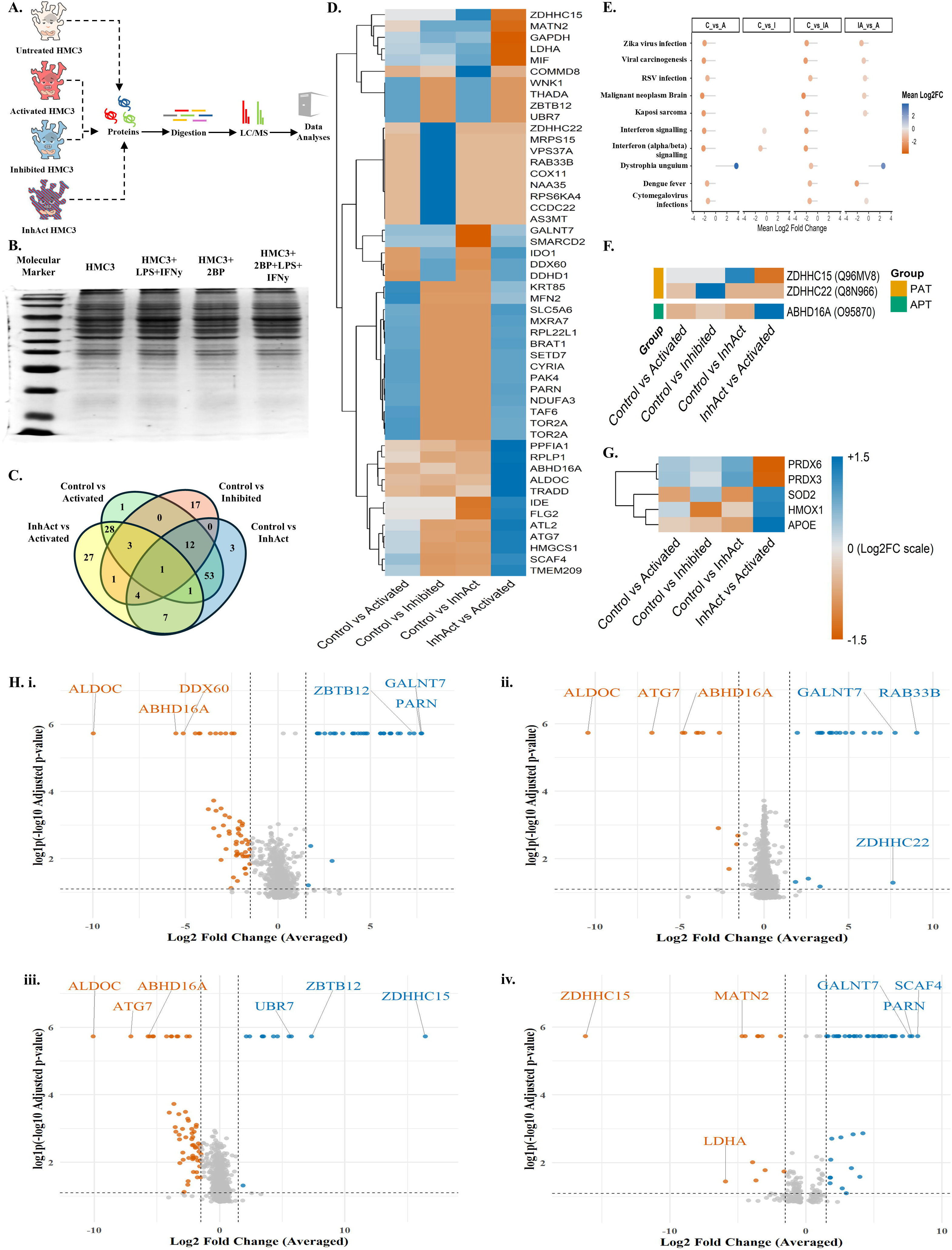
Proteomic-level characterization of microglia responses to treatment with 2-BMP or LPS and IFNy. **A.** A cartoon overview of the experimental workflow followed for the proteomic characterization of cell responses induced in control, activated, inhibited and InhAct HMC3 microglia cells. **B.** A representative SDS-PAGE gel of cell lysates purified from the control and treatment groups of HMC3 cells as a reference for equal loading. **C.** Venn diagram to suggest the number of unique proteins found in each of the comparison groups studied (Control HMC3 vs Activated HMC3, Control HMC3 vs Inhibited HMC3, Control HMC3 vs InhAct HMC3 and InhAct HMC3 vs Activated HMC3). **D.** A heatmap representation of most highly dysregulated proteins found across the ascribed comparison groups of HMC3 cells. **E.** A combined lollipop plot to highlight the dysregulated biological functions identified in the ascribed comparison groups of HMC3 cells. **F.** A heatmap plot to exhibit the most prominently dysregulated s-acylation enzyme proteins (PATs and Deacylases/APTs) across the comparison groups of HMC3 cells studied. **G.** A heatmap plot to exhibit the most prominently dysregulated antioxidant proteins across the comparison groups of HMC3 cells studied. **H.** Volcano plots representing differentially expressing proteins in various treatment groups of HMC3 cells^#^; (i). Relative protein expression patterns in control as compared with LPS and IFNy stimulated (activated) HMC3 cells. (ii). Relative protein expression patterns in control as compared with 2-BMP treated HMC3 cells. (iii). Relative protein expression patterns in control as compared with InhAct (2-BMP pretreated and LPS and IFNy activated) HMC3 cells. (iv). Relative protein expression patterns in InhAct as compared with activated HMC3 cells to summarise the effects of 2-BMP treatment in resting and activated HMC3 cells. ^#^Plotted points are either labelled blue (upregulated in the initial group or condition) or red (upregulated in the second condition with which the relative expressions are calculated in the initial condition) based on a threshold of absolute log2 fold change (absLog2 FC) ≥ 1.5 (represented on X-axis as averaged values of outputs from multiple calculation methods) and adjusted p value ≤ 0.01 (represented on Y-axis as log1p{-log10 of adjusted p values}).

In our proteomic results we identified a significant dysregulation (absolute log2 FC ≥ 1.5, adjusted p ≤ 0.05) of 158 unique proteins (Figure 3C). Specifically, an induced expression of proinflammatory markers TLR2/4 effector RIPK2, JAK-STAT signalling factors like STAT1 and pattern recognition receptors (PRRs) such as MDA5 (or IFIH1) and RIG-I were detected in these cells (Supplementary Table 3A). Conversely, the expression of nonsense-mediated mRNA decay protein poly(A)-specific ribonuclease (PARN) was found to be significantly downregulated in these ‘activated’ HMC3 (Figure 3D), possibly to support increased protein synthesis demands during immune polarization (J. E. Lee et al., 2012) of microglia (Figure 3E, Supplementary Table 3A).

Quite remarkably, a treatment of HMC3 cells with 2-BMP (‘inhibited’ HMC3) was sufficient to reduce the expression of cell activation markers such as TRIM14, suggesting a strong disruption of IFN regulatory factor 3 (IRF3), NF-kappa-B (Z. Zhou et al., 2014) and autophagic signalling pathways in these cells (D. Liu et al., 2022). However, the expression of proinflammatory proteins found earlier in ‘activated’ HMC3 cells were also observed as upregulated in 2-BMP pretreated HMC3 treated with LPS and IFNy (‘InhAct’ HMC3), suggesting a synergistic contribution of LPS/IFNy and 2-BMP to inflammation in microglia.

Among the s-acylation (and deacylation) enzymes, ABHD16A was consistently found to be most dysregulated in all treatment groups relative to control HMC3 cells. PATs ZDHHC15 and ZDHHC22 were found to be dysregulated in ‘InhAct’ and ‘inhibited’ HMC3 cell groups respectively (Figure 3F, Data provided in Supplementary Table 3A). Additionally, the antioxidant SOD2 protein, was found as most significantly upregulated in ‘activated’ and ‘InhAct’ HMC3 cell populations, highlighting its centrality in regulating the oxidative stress in inflammation-responsive microglia. Other NRF2-pathway antioxidants like peroxiredoxins (PRDX3 and PRDX6) and protein disulfide-isomerase A6 (PDIA6) were also found as upregulated in ‘activated’ HMC3 cells as compared to ‘InhAct’ HMC3 cells (Figure 3G, Supplementary Table 3A), indicating that the signalling pathways inducing these antioxidant responses in microglia are sensitive to 2-BMP. Moreover, a slightly higher enrichment of PDIA6 proteins indicate a better resolution of unfolded protein aggregation and stress in ‘activated’ HMC3 than ‘InhAct’ HMC3.

#### **B.** Nuclear activation of NRF2 mediates microglial responses to inflammation

To understand the extent of oxidative stress triggered in each of the HMC3 treatment groups, we monitored the nuclear colocalization (activation) of ‘master’ oxidative-stress-response factor NRF2 using confocal microscopy (Figure 4A).

**Figure 4.**
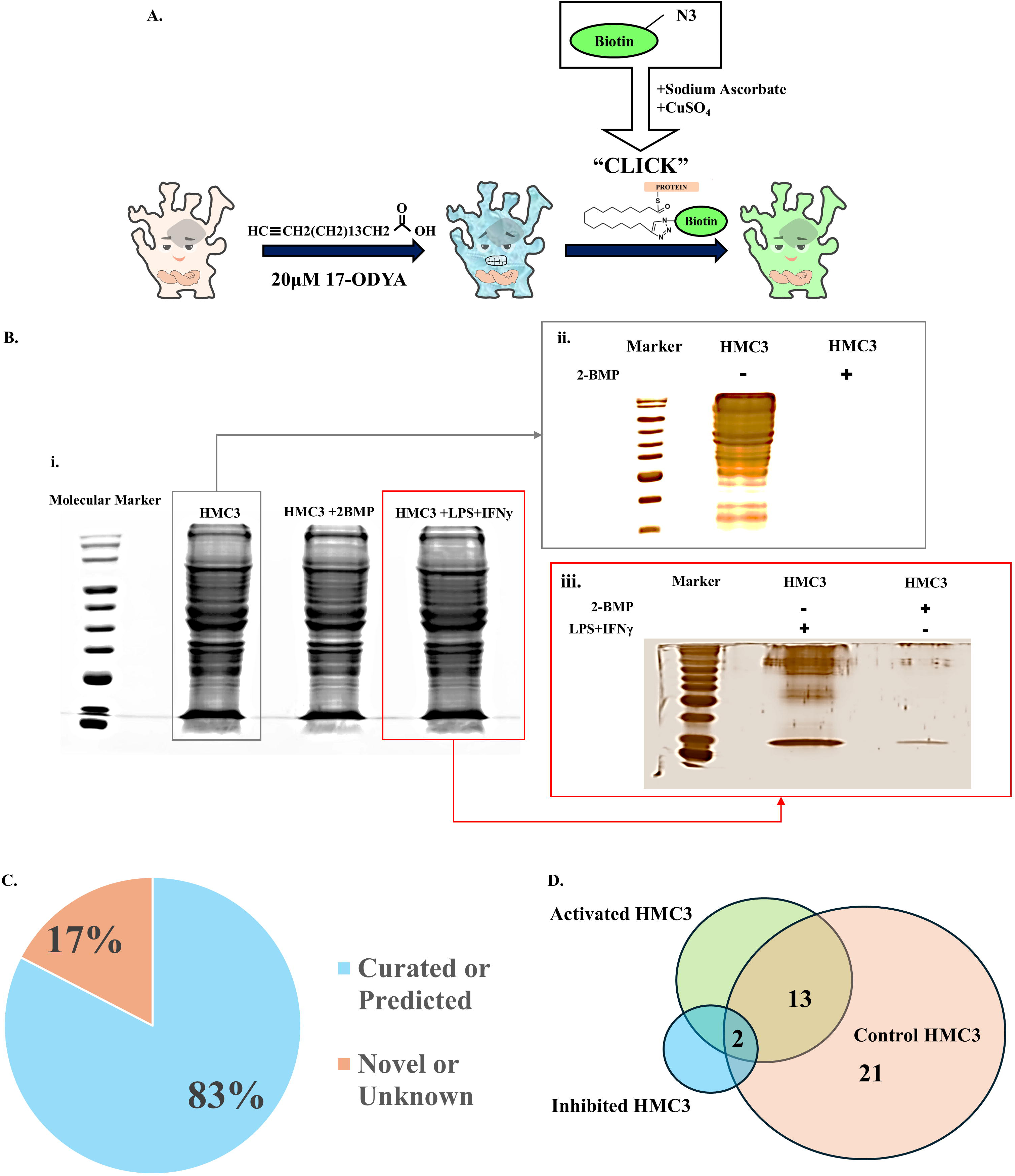
Characterization of the levels and routes of oxidative stress induced in microglia after treatment with 2-BMP, LPS and IFNy. **A.** A schematic view of the experimental workflow to visualize the nuclear activation of NRF2 in control, activated, inhibited and InhAct HMC3 cells. Some images were obtained from the BioArt repository (NIAID Visual & Medical Arts. (10/7/2024). Confocal Microscope. NIAID NIH BIOART Source. bioart.niaid.nih.gov/bioart/86 and NIAID Visual & Medical Arts. (10/7/2024). Antibody. NIAID NIH BIOART Source. bioart.niaid.nih.gov/bioart/17). **B.** Confocal microscope-captured images exhibit differential localization patterns of NRF2 proteins within HMC3 cells belonging to different treatment groups. **C.** Quantitative analysis of NRF2 nuclear localization in treated and untreated HMC3 cells. Bar graphs summarising the Pearson correlation coefficients (PCC values) for colocalization of fluorescence-tagged NRF2 proteins with the nuclear dye DAPI in control (green), 2-BMP treated (inhibited, orange), LPS and IFNy activated (blue) and 2-BMP pretreated and LPS and IFNy activated (InhAct, pink) HMC3 cell populations. **D.** A simple mechanistic view of oxidative stress responses and cell fate outcomes triggered as inferred in HMC3 microglia after treatment with LPS and IFNy or 2-BMP.

Expectedly, the nuclear localization of NRF2 was increased in LPS and IFNγ stimulated HMC3 cell populations (PCC=0.69) as compared to the resting state HMC3 cells (PCC=0.08), suggesting a significant increase in oxidative stress. Higher than CC50 concentrations of 2-BMP (100μM) were also found to increase the nuclear translocation of NRF2 (PCC=0.43). The 2-BMP (100μM) pretreated HMC3 cells stimulated with LPS and IFNγ displayed the maximal nuclear translocation of NRF2 (PCC=0.8), showcasing the synergistic effects of individual treatments on the oxidative stress state of microglia (Figure 4B, Supplementary Table 3B-C).

Based on our observations and current evidence, a molecular model describing the effects of 2-BMP and LPS/IFNy treatment on morphological and survival dynamics of HMC3 cells was predicted (Figure 4D).

### 4. Microglia antioxidants can be reversibly s-acylated during inflammation

To establish s-acylation as a direct causal of antioxidant responses in microglia, we performed the MS-coupled MLCC-based enrichment of ‘s-acylable’ proteins in untreated and ‘activated’ HMC3 cells (Figure 5A-B).

**Figure 5.**
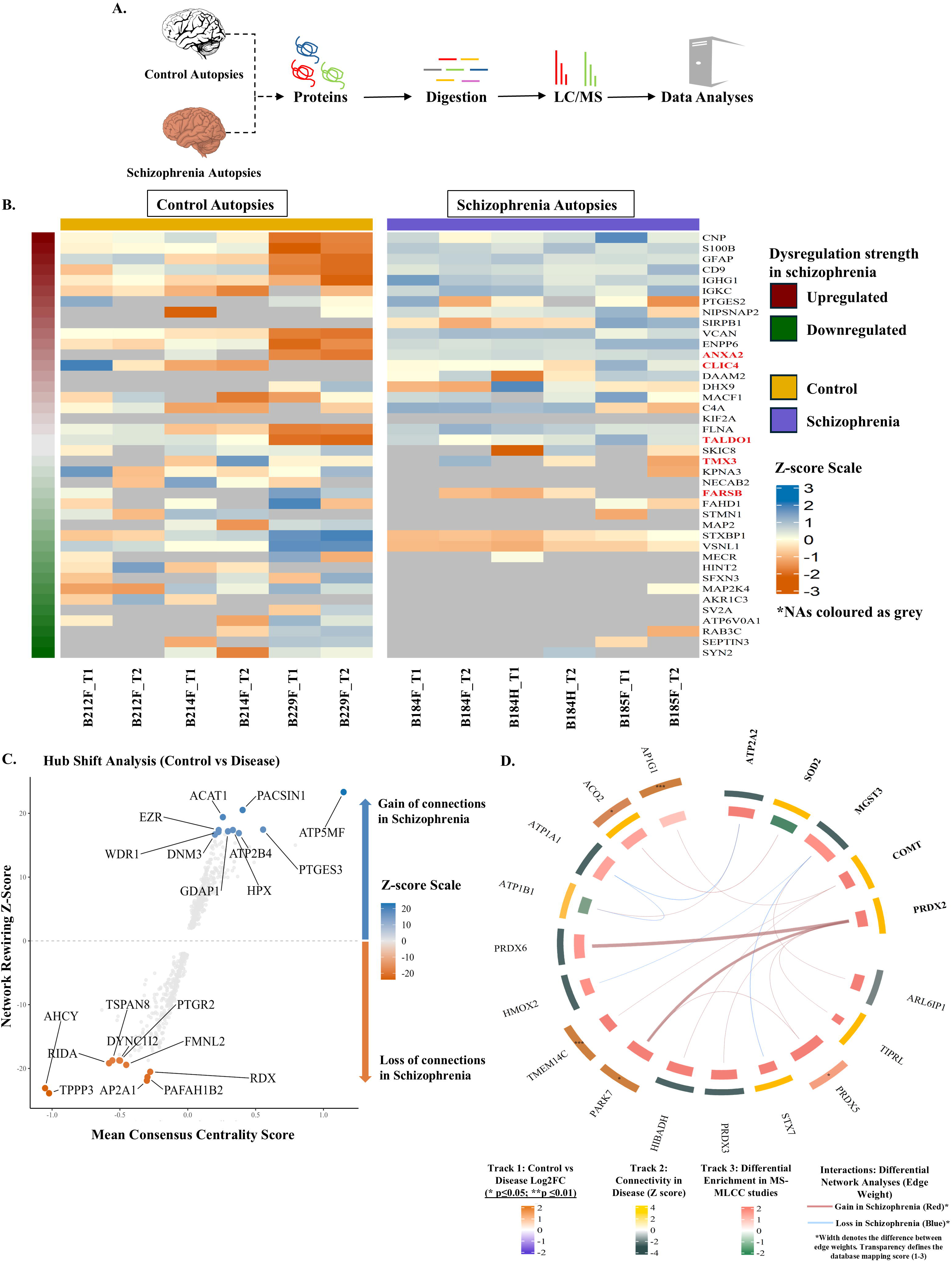
Overview of metabolic labelling and click chemistry (MLCC)-based proteomic experiments. **A.** A cartoon representation of the MLCC based capture of s-acylated biomolecules from treated and untreated HMC3 microglia. Small adjustments were made to the earlier MLCC-visualization workflows (described in Figure 2), such as azide-conjugated OG488 dye was replaced with azide-conjugated biotin to capture proteins (potentially with other biomolecules) metabolically labelled with the fatty acid analogue 17-octadecynoic acid (17-ODYA). **B.** Representative SDS-PAGE gel pictures suggesting the MLCC-based capture of s-acylated proteins. (i.) Whole proteins purified from the control, inhibited and activated groups of HMC3 cells as a reference for equal loading. (ii.) MLCC-captured proteins from control HMC3 cells. (iii.) MLCC-captured proteins from activated HMC3 cells. Total MLCC-captured proteins from both control and activated HMC3 cells, were individually normalized with total MLCC-captured proteins from inhibited HMC3 cells. **All protein pulldown products were (pre-) normalized using BCA method before Mass Spectrometric analyses. **C.** A pie chart summary of MLCC-MS-characterized proteins validated as s-acylated proteins after database mapping. Detailed confidence calculations are provided in Supplementary Table 4. **D.** Venn diagram to represent the number of PATs and deacylases (APTs) detected among the MLCC-MS characterized s-acylated proteins in untreated (control), LPS and IFNy stimulated (activated) and 2-BMP treated (inhibited) HMC3 cell populations.

Robust and reproducible libraries of high confidence s-acylated proteins were prepared from the final enrichment results. While 1568 proteins were found to be significantly enriched in MLCC pools obtained from the untreated HMC3 cells, only 883 proteins were detected as significantly enriched in MLCC pools of activated HMC3 cells (MLCC-MS Log2FC≥1.5, adjusted p≤0.05; Supplementary Table 4). Over 83% of these microglia-proteins detected in our MLCC-MS libraries could be found in public datasets or were predicted as high-confidence substrates of s-acylation using the GPS-Palm tool (with cutoff score ≥ 0.892; Figure 5C, Supplementary Table 4). Also, among all the HMC3 proteins detected in MLCC-MS studies, 908 proteins were characterized as differentially enriched (Difference of MLCC-MS Log2FC≥1.5; Data provided in Supplementary Table 4).

24 protein accessions correlating to ZDHHC-PATs and 10 protein accessions correlating to deacylases could be found across all MLCC-MS libraries from HMC3 microglia, corresponding to nearly 62% of all known s-acylation enzymes (Supplementary Table 4). While the s-acylated forms (or complexes) of ZDHHC17, IFIT1 and IFIT2 were detected as significantly increased in the MLCC-MS libraries of ‘activated’ HMC3 cells (difference of log2FC between control and activated HMC3≤-5.71), the s-acylated forms (or complexes) of ZDHHC7, RHOB and ZDHHC16 were identified as significantly reduced in the MLCC-MS libraries of ‘activated’ HMC3 cells (difference of log2FC between control and activated HMC3≥7.76).

Remarkably, the MLCC-MS experiments revealed a lower enrichment of around 1151 s-acylated proteins and groups (corresponding to 1157 isoforms) in inflammatory microglia relative to untreated microglia populations (difference of log2FC between control and activated HMC3>1, adjusted p≤0.05). Among these, we could designate 11 proteins as antioxidants (with s-acylation confidence grade of 2 or above) by performing gene ontology analysis. This included s-acylable antioxidant proteins like calgranulin B (S100A9 or MRP-14), peroxiredoxins (PRDXs 5, 3, 2 and 6), thioredoxin-signalling related thioredoxin reductase 1 (TXNRD1), glutathione-metabolism related glutathione s-transferase omega 1 (GSTO1) and glutamate-cysteine ligase catalytic subunit (GCLC), and NRF2-regulatory deglycase (PARK7 or DJ-1). In contrast, the s-acylated forms of mitochondrial superoxide dismutase 2 (often called manganese superoxide dismutase, MnSOD or SOD2 (featuring s-acylation confidence grade of 3), were found to be significantly enriched in inflammatory microglia relative to resting microglia populations (difference of log2 FC=-2.22, adjusted p ≤ 0.02). These quantification studies highlighted s-acylation of NRF2-antioxidants as a metabolically ‘flexible’ response to immune polarization of microglia (Supplementary Table 4).

### 5. ‘Flexibly s-acylable’ microglia antioxidant proteins drive brain dysfunction

The MLCC-MS-identified ‘flexibly’ s-acylated microglia antioxidants were finally characterized as protein ‘drivers’ of schizophrenia, after mapping to results from differential expression and differential network (conditional) analyses of MS-quantified proteins in control and schizophrenia-affected tissues.

Since the microglia cells from frontal white matter and hippocampal regions of the human brain share morphological attributes including soma size, cytoplasmic area, ramification complexity and total size (Tan et al., 2020), and microglia inflammation in these regions are associated with psychotic disorders such as schizophrenia (Arnold et al., 2015; Brisch et al., 2022; Ermakov et al., 2021; Kochunov et al., 2017; Lieberman et al., 2018; Nazeri et al., 2013; Yamada et al., 2022), we expected the overall proteomic-level changes within these brain regions to summarise microglia activity in schizophrenia progression.

Notably, the dysregulation of expression markers S100A1, CAP1, the C3-SIRPA axis and complementary cascades suggested a significant impairment of microglial activity and inflammation in schizophrenia (Supplementary Table 5). Hallmarks of glial activation (like GFAP) and oxidative stress were also observed in differential expression studies of schizophrenia tissues (Davalieva et al., 2016).

Interestingly, many differentially s-acylable proteins identified in microglia including ANXA2, CLIC4, TALDO1, TMX3 and FARSB were also found to be dysregulated in schizophrenia-affected brain autopsies (Figure 6B). Although, the general expression of ZDHHC-PATs and deacylases were noted as relatively low within the tissue-derived proteomes, several peptides correlating to the s-acylation enzyme ZDHHC13 were found to be consistently downregulated in schizophrenia-affected tissues, a loss of which is typically associated with elevated RONS production (Shen et al., 2017) (Supplementary Table 5).

**Figure 6.**
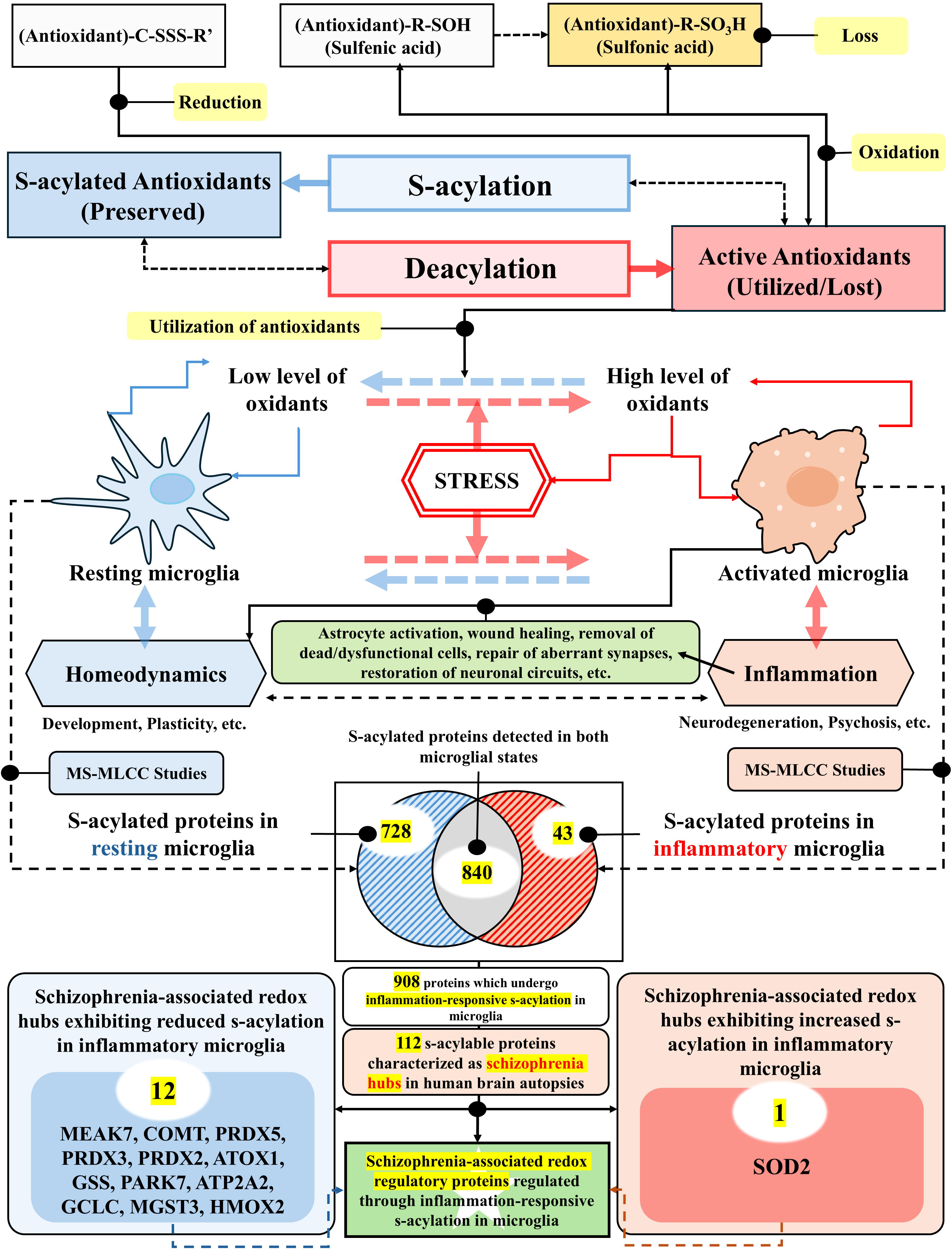
Characterization of differentially s-acylated proteins as conditional drivers of schizophrenia. **A.** A cartoon summary of experimental workflows followed to identify dysregulated proteins and causal proteins in schizophrenia. **B.** A heatmap representation of protein expression in control and schizophrenia affected autopsies for the highly dysregulated proteins. The proteins characterized earlier in MLCC-MS studies as differentially s-acylated in activated HMC3, are highlighted as ‘red’ fonts. **C.** A hub-shift analysis plot to suggest the most dysregulated protein network hubs in schizophrenia. **D.** An integrated Circos plot to visualize the systemic changes of interactions between differentially s-acylated proteins in schizophrenia relative to ‘homeodynamic’ control states. The outermost track features gene symbols of the five most dysregulated schizophrenia hubs also inferred as differentially s-acylable through MLCC-MS studies (in bold fonts) and their plausible interactors selected by absolute enrichment scores and differential connectivity. The annotation track 1 exhibits the relative (Log2FC) expression of each protein in schizophrenia-affected brain tissues relative to control. The colour scale of track 1 ranges from purple denoting the lowest log2FC value (downregulation) to orange denoting the highest log2FC values (upregulation). Statistical significance is denoted by asterisks (* = adjusted p-value<0.05, ** = adjusted p-value<0.01, *** = adjusted p-value<0.001). Annotation track 2 is a heatmap representation to indicate the differential ‘hubness’ of each protein. The colour scale of track 2 ranges from teal denoting the lowest Z-score values (overall loss of connections) to gold denoting the highest Z-score values (overall gain of connections). Annotation track 3 demarcates the differential enrichment of each protein in the MLCC-MS studies. The colour scale of track 3 ranges from green denoting the highest log2FC enrichment in s-acyl proteome of activated HMC3 to salmon denoting the lowest log2FC enrichment in s-acyl proteome of activated HMC3. The width of the bars in track 3 indicates the confidence weight of the protein to be part of the s-acyl proteome. The central interaction chords represent the interaction rewiring between important hubs identified in our study. While red chords indicate a gain of interaction strength in schizophrenia, blue chords indicate a loss of the interaction strength. The thickness of the chords represents the absolute difference of a protein interaction in schizophrenia brain autopsies studied, thereby highlighting the most significant disruptions of interactions between proteins which undergo inflammation-responsive s-acylation.

From these differential expression studies, the differentially ‘s-acylable’ NRF2-regulator and antioxidant PARK7 was detected as significantly downregulated in schizophrenia-affected tissues (differential expression log2FC=2.89, adjusted p ≤ 0.05; Supplementary Table 5). Evidently, 8 other differentially ‘s-acylable’ (absolute MLCC-MS enrichment in control vs activated HMC3 cells ≥ 1.5) antioxidant proteins including MGST3, ATOX1 and PRDX2, were identified as significant network ‘hubs’ through the ‘conditional’ network prediction models (consensus pvalue≤0.05), highlighting these proteins as high-confidence, schizophrenia-relevant and metabolically ‘flexible’ proteins dependent on dynamic s-acylation states (Table 2; DNA, Supplementary Table 5). Differentially ‘s-acylable’ proteins AHCY (Z score=-23.09) and ATP5MF (Z score=23.34), were found as the most dysregulated hubs associated with schizophrenia progression (Figure 6C). All the consensus weighted network graphs, generated with proteomic data (DIA-SWATH-MS) recorded from control and schizophrenia-affected brain tissue lysates, were stored as raw (.graphml) files (Supplementary Files 5-12).

**Table 2.** Differentially ‘s-acylable’ antioxidant hubs or ‘drivers’ of schizophrenia.

| **Protein** | **Gene** | **diffMLCC_Log2FC** | **Remarks** | **AcylationConfidence** | **MicrogliaEnriched?** | **DNA_Z (only significant)** | **DEA (only significant)** |
| --- | --- | --- | --- | --- | --- | --- | --- |
| P04179 | SOD2 | -2.22 | More s-acylated in actHMC3 | high | No | 12.15 | 0 |
| O14880 | MGST3 | 1.54 | More s-acylated in conHMC3 | very high | No | -8.37 | 0 |
| P32119 | PRDX2 | 2.91 | More s-acylated in conHMC3 | mild | MicrogliaEnriched | 7.35 | 0 |
| P30048 | PRDX3 | 3.37 | More s-acylate d in<br>conH<br>MC3 | high | No | -4.47 | 0 |
| P305<br>19 | HMO<br>X2 | 1.54 | More<br>s-acylate<br>d in<br>conH<br>MC3 | mild | No | -4.29 | 0 |
| P486<br>37 | GSS | 2.24 | More<br>s-acylate<br>d in<br>conH<br>MC3 | low | No | -3.23 | 0 |
| O00<br>244 | ATO<br>X1 | 2.28 | More<br>s-acylate<br>d in<br>conH<br>MC3 | low | No | -2.57 | 0 |
| P485<br>06 | GCL<br>C | 2.04 | More<br>s-acylate<br>d in<br>conH<br>MC3 | mild | No | -2.48 | 0 |
| P300<br>44 | PRD<br>X5 | 3.68 | More<br>s-acylate<br>d in<br>conH<br>MC3 | very high | No | 0 | 1.14 |
| Q99<br>497 | PAR<br>K7 | 2.23 | More<br>s-acylate<br>d in<br>conH<br>MC3 | high | No | 0 | 2.89 |

Dysregulated schizophrenia driving proteins which were also identified to be differentially s-acylated in ‘activated’ HMC3 cells, were found to coordinate various biological functions such as mechanotransduction, protein folding responses, mitophagy and apoptosis (Supplementary Table 5). Collectively, these findings highlighted the dynamic s-acylation-deacylation mechanism as a tractable regulator of schizophrenia pathogenesis in the brain (Figure 6, Supplementary Figure 1).

## Discussion

The results from our study strengthened s-acylation of proteins as a biochemical driver of inflammatory signalling within brain-resident microglia cells. Moving beyond previous omics studies, we could present a comprehensive map of reversibly s-acylable proteins in untreated and inflammatory microglia, whose modification status can be pharmacologically ‘hacked’ to diagnose or treat inflammatory diseases of the brain such as schizophrenia.

We initially prepared a reliable catalogue of all ‘s-acylable’ (substrate) human proteins using *in silico* methods. Within this catalogue, we were able to identify a subset of s-acylation (catalyst) enzymes which exhibit maximal transcriptomic enrichment in human microglia, as reported by the Brain-RNAseq database. RT-qPCR studies further revealed a context-dependent transcriptional prevalence of s-acylation enzymes in HMC3 microglia. This provided us the evidence of an endogenous s-acylation machinery in microglia and its responsiveness to inflammation.

Confocal microscopic analyses were subsequently performed with HMC3 cells to determine the impact of inflammation on intracellular arrangement (or localization) of the s-acylation machinery and the s-acylated biomolecules. The redistribution of s-acylated biomolecules strongly correlated with the spatial reorientation of ER-Golgi network fragments in inflammatory microglia, suggesting a functional coupling of both. Since s-acylation is understood to drive ER-Golgi trafficking by promoting the membrane affinity and lateral clustering of cargo proteins at the vesicle forming rims of Golgi cisternae (Ernst et al., 2019), we hypothesized that the activation of microglia under inflammation is reflected through an induced Golgi-ER based trafficking (or sorting) of s-acylated biomolecules.

To elaborate the global impacts of inflammation in microglia, we performed proteomic studies with untreated HMC3 cells and ‘activated’ HMC3 cells treated with proinflammatory PAMPs, LPS and IFNy. Similar workflows were applied to study proteomic changes in HMC3 cells treated with the s-acylation inhibitor 2-BMP alone (‘inhibited’ HMC3) or in combination with LPS and IFNy (‘InhAct’ HMC3). Through this study, we could detect a higher enrichment of ALDOC in all treatment groups relative to control HMC3 cells (Figure 3D, Supplementary Table 3A), indicating that all treatments can induce glycolytic shift and mitochondrial dysfunction in HMC3 microglia. Additionally, a remarkable upregulation of TALDO1, TRIM21, TLRs, JAK-STAT pathway factors and PRRs was observed in ‘activated’ and ‘InhAct’ HMC3 microglia (Supplementary Table 3A). This highlighted the activation of NF-κB-NLRP3 signalling axes in inflammatory HMC3 cells, leading to a release of proinflammatory cytokines such as IL1β (Hetz & Saxena, 2017; Rojo et al., 2014; Sikorski et al., 2012; P. Wang et al., 2017; Wei et al., 2024; Xiao et al., 2021). Simultaneously, the 3’-exoribonuclease PARN was detected in all treatment groups, but not in ‘activated’ HMC3 cells in the raw datasets. An inflammation-responsive loss of this gene elevates the stabilization of more gene transcripts, thereby increasing the production of proinflammatory proteins and ER stress in HMC3 microglia, which can be recovered through PDIA6-dependent activation of unfolded protein response (UPR) processes, particularly of the PERK/p-eIF2α/ATF4/CHOP, IRE1α/XBP1 or ATF6 signalling pathways. Strikingly, the expression of chaperone protein PDIA6 was augmented in ‘activated’ cells relative to ‘InhAct’ HMC3 cells, confirming more efficient regulation of proteostatic stress in these cells (Chen et al., 2023; Hetz & Saxena, 2017; Jaronen et al., 2014; Junjappa et al., 2018; Mo et al., 2021; Ransohoff, 2016). Moreover, HO-1 were found to be more enriched in ‘InhAct’ and ‘inhibited’ groups, but not in ‘activated’ HMC3 cells. Since HO-1 protein is induced by intracellular ER-stress (X. Liu et al., 2005), we presumed 2-BMP treatment to elicit more prominent ER-stress in HMC3 cells as compared to treatment with LPS and IFNy. These results suggested lower ER-stress in ‘activated’ cells, but increased ER-stress in ‘inhibited’ and ‘InhAct’ HMC3 microglia. Consequently, when confocal microscopy was performed with these cells, ‘inhibited’ and ‘InhAct’ HMC3 microglia displayed ‘injured’ morphologies. Together, increased ER-stress, ER-Golgi trafficking processes, glycolysis, injuries and mitochondrial dysfunction can stimulate CNS microglia to release excessive amounts of oxidants (Fujita et al., 2014; Kettenmann et al., 2011; Y. Li et al., 2022; Liemburg-Apers et al., 2015).

The tissue redox balance disturbed by the oxidants and excitotoxins is reciprocally stabilized through the NRF2-mediated transcription of cytoprotective and antioxidant genes (Rojo et al., 2014). Since the upregulation and nuclear engagement of intracellular NRF2 is a direct mechanistic marker of oxidative stress in mammalian cells (Ma, 2013), we used it as a measure of oxidative stress induced by each of the treatment conditions in HMC3 cells. An elevated nuclear activity of NRF2 was observed in HMC3 cell populations treated separately with 2-BMP or LPS and IFNy. However, the maximal enrichment and nuclear translocation of NRF2 was observed in ‘InhAct’ HMC3 microglia (measured colocalization of tagged NRF2 with DAPI; PCC=0.8). These results validate our proteomic studies where we found the stress marker DNAJA1 (log2FC=1.2, adjusted p << 0.01) and the NRF2-antioxidant HO-1 (log2FC=1.5, adjusted p << 0.01) to be higher enriched in ‘InhAct’ HMC3 cells relative to ‘activated’ HMC3 microglia (Figure 3G, Supplementary Table 3A).

In the DIA-SWATH proteomic studies, almost all ZDHHC-PATs were observed to remain relatively stable after co-stimulation of HMC3 microglia with LPS and IFNγ, except the deacylase ABDH16A (absolute log2FC ≥ 1.5, adjusted p ≤ 0.05). Similar trends were observed after treatment with the s-acylation inhibitor, 2-BMP. These results suggested the expression of s-acylation enzymes as constitutive than inflammation-responsive, thus underscoring other factors, like kinetic flexibility of these enzymes or dynamic availability of metabolic fatty acyl groups, as more important determinants of s-acylation ‘flexibility’ in inflammatory microglia.

Beyond protein enrichment, the differential prevalence of cellular s-acylation processes was observed as a biochemical marker to determine the redox-sensitive activation of microglial antioxidants during inflammation (Qiu et al., 2025), which modulate microglia responses to oxidative stress, ultimately shaping the inflammatory responsiveness of the cell type. For this, comparative (and quantitative) MLCC-MS studies were initially performed between resting, ‘inhibited’ and ‘activated’ HMC3 microglia. A comparison of MLCC-MS enrichment results from resting or ‘activated’ HMC3 with MLCC-MS enrichment results from ‘inhibited’ HMC3 cells subsequently revealed a cohort of proteins which could be s-acylated in HMC3 cells.

In our MLCC-MS studies, the peptides and proteins were enriched in resting and activated HMC3 cells relative to 2-BMP treated cells, in presence or absence of biotin-azide molecules (used for the capture of 17-ODYA-alkyne labelled proteins), but hydroxylamine-based enrichment strategies were not applied to minimize the exclusion of false-negative signals (reduced recovery or loss of true signals, especially phosphopetides) (Antorini et al., 1997; Demyanenko et al., 2025; Kwon et al., 2018). However, hydroxylamine treatment steps can be reintroduced in targeted studies for confirming s-acylation status of proteins or as part of more advanced workflows such as those coupled with iodoTMT0 labelling for the identification of s-acylated sites within these proteins (Cai et al., 2024). To address the significant limitations of the project and to restore the authenticity of our microglia-specific s-acylation datasets, we employed rigorous data mining and database searches for assigning confidence values to all identified proteins. In addition, we validated the specificity of 2-BMP for isolating s-acylated proteins, by correlating the newly established s-acylation libraries with global proteomic changes observed after the treatment of HMC3 with 2-BMP.

The likelihood of each protein to be s-acylated were next measured by confidence weights, derived from curated databases SwissPalm and UniprotKB, and the prediction tool GPS-Palm. Specifically, 1395 and 801 high confidence (with confidence weights or scores≥2) s-acylated proteins were detected in MLCC-MS fractions obtained from resting and ‘activated’ HMC3 microglia respectively (absolute log2FC≥1.5, adjusted p ≤ 0.05). Among these, 780 proteins were detected as differentially ‘s-acylable’ proteins in ‘activated’ HMC3 cells relative to the untreated control cells (Difference_Log2FC≥1.5). In the light of these findings, we observed a significant number of membrane proteins and molecular processes such as oxidative-stress response signalling to undergo differential s-acylation in inflammation-responsive microglia (Guns et al., 2022; Ouyang et al., 2024).

12 major microglia-inducible antioxidants, including PRDXs and TRXRs, were found to be consistently reduced in HMC3 cells treated with PAMPs – LPS and IFNy. Only one antioxidant, SOD2, was found to ‘atypically’ exhibit elevated s-acylation in inflammatory or (LPS and IFNγ)-triggered HMC3 cells than from untreated HMC3 cells, maybe to keep the dramatically-induced SOD2 proteins from getting activated all at once. However, no targeted records could be found to associate dynamic s-acylation with SOD2 function. Thus, no technical assumptions were made for the differential s-acylation of SOD2. Since, similar redox-sensitive s-acylation mechanisms have been previously described for the activation of microglia PRDXs (Qiu et al., 2025), we presumed the inflammation-responsive downregulation of s-acylated antioxidants as a crucial means to sensitize related oxidative-stress responses in microglia. Collectively, a reduced s-acylation of antioxidant proteins in ‘activated’ HMC3 microglia, correlate with morphological plasticity and anti-inflammatory polarization of microglia (specifically the disease-associated microglia or DAMs) to facilitate tissue repairs in schizophrenia or other brain dysfunctions caused by oxidative stress and inflammation (Kim et al., 2022; Vilhardt et al., 2017).

The results of our comparative MLCC-MS experiments corroborated published studies, highlighting two most plausible correlations of differentially s-acylated antioxidant proteins with context-dependent plasticity of microglia activities. First, the s-acylation of TXNRD1 is expected to reduce the antioxidant’s activity by promoting its membrane association, potentially redirecting it towards non-canonical pathways such as the formation of membrane protrusions (Blanc et al., 2015; Cebula et al., 2013). High levels of s-acylated TXNRD1 detected in only untreated microglia, overlaps well with high degree of ramifications in these cells. Dynamic s-acylation of TXNRD1 may trigger morphological rearrangements in microglia, or conversely, reconfiguration of ramification may induce deacylation to restore canonical antioxidant function of proteins like TXNRD1. Second, the s-acylation of PARK7 protein (or DJ-1) is expected to keep it associated with the membrane lipid rafts, and its deacylation should promote its delocalization towards the cytoplasm, where it can engage the NRF2 signalling pathway in stress conditions. This correlates with relatively lower enrichment of s-acylated DJ-1 observed in inflammatory microglia than resting microglia (J. Liu et al., 2025; Zgorzynska et al., 2021; F. Zhang et al., 2021). While these mechanistic models may be reliable, these are not proven in our study and require proper experimental validation through future studies.

Comparative proteomic-expression studies of control and schizophrenia-affected postmortem brain tissues were performed to measure the role of microglial inflammation in progression of brain dysfunctions. Through these comparative proteomic studies, we could detect a higher enrichment of astrocytic activation marker GFAP and a lower expression of microglia-enriched DYNLT1 proteins in schizophrenia-affected tissues, highlighting glial activation as a significant clinical factor of schizophrenia progression in patient autopsies studied (Davalieva et al., 2016). In addition, an altered expression of key oxidative stress–responsive proteins like S100B, RHOA and MAP2K4, highlighted the inflammatory progression in these tissues. Specifically, a stimulated expression of RhoA/ROCK signalling pathway components indicated a probable blood-brain barrier dysfunction and oxidative stress states in schizophrenia tissues (Hwang et al., 2025). A significant dysregulation of other crucial factors regulating glial-cell specific innate immunity, extracellular matrix reorganization, redox sensor NRF2 activity and developmental pathways provided as prominent hallmarks of neuroinflammation, oxidative stress and structural changes in the schizophrenia tissue samples studied (Bitanihirwe & Woo, 2014; Murray et al., 2021; Traina et al., 2022).

Consensus (conditionally) weighted network graphs were constructed from the quantitative proteomic data (DIA-SWATH-MS). These networks revealed metabolic protein hubs and dysregulated protein interactions driving the progression of schizophrenia in the brain tissue. Finally, the results from earlier comparative MLCC-MS studies were mapped on these conditional graphs to investigate the clinical relevance of proteins undergoing inflammation-responsive s-acylation in microglia. This delineated potential therapeutic targets and schizophrenia-sensitive signalling pathways, which can be ‘flexibly’ modulated by redox-sensitive s-acylation machineries in microglia (Supplementary Figure 1).

Interactions of the flexibly s-acylable antioxidant PRDX2 with other antioxidant proteins PRDX6 and PARK7 were found to be significantly enhanced in schizophrenia brains relative to control. Since PRDX2 has been previously characterized as a potential biomarker of early psychosis (H. Lee et al., 2024), we presumed that the inflammation-responsive s-acylation of PRDX2 and its interactions with other s-acylable antioxidants must be important factors determining the onset and progression of schizophrenia (Figure 6D, Supplementary Figure 1). Importantly, a gain of interactions for the ‘hub’ antioxidant PRDX2 indicated increased microglial/macrophage-driven astrocyte-inflammation, mitochondrial dysfunction and RONS production in schizophrenia-affected tissues studied herein (Voigt et al., 2017). Moreover, a reduction of interactions observed between the transmembrane P-type ATPase ion-pumps such as the sodium/potassium-transporting ATPases ATP1A1 and ATP1B1 and the SERCA2 (Sarco/Endoplasmic Reticulum Calcium-ATPase 2) pump ATP2A2 (Figure 6D) provided as a clear biological hallmark of cellular ER stress, disrupted ion ‘homeodynamics’ (mainly of sodium, potassium and calcium ions), impaired dopamine signalling and neuroinflammation in the tissue samples studied (Kong et al., 2025; Lin et al., 2021; Nakajima et al., 2021; Selvakumar et al., 2014; Wen et al., 2018). Results from our MLCC-MS studies further highlighted the s-acylation of these proteins (or their biomolecular complexes) as metabolically ‘switchable’ during inflammation, this emphasizes the role of this posttranslational modification in maintaining brain homeodynamics and preventing schizophrenia progression.

Some more mechanistic hypotheses could also be derived after mapping of MLCC-MS results to the results of differential network analyses. For instance, in ‘homeodynamic’ microglia, a synchronous s-acylation of antioxidant PRDX6 and mitochondrial protein VDAC1, should induce their co-localization and association on similar cellular microdomain structures within the mitochondria-associated ER membranes (MAMs) and the outer mitochondria membrane (OMM). This facilitates repair processes and oxidative-stress (sensitization) signalling through the Ca²L-independent phospholipase AL (iPLAL) activity of PRDX6. Similarly, s-acylation triggers VDAC1 to interact with MAMs-enriched proteins such as GRP75 and PRDX6. While a dynamic deacylation of PRDX6 can restore its antioxidant and peroxidase activity (leading to its utilisation), the deacylation of VDAC1 decouples it from cellular organizations like MAMs and signalling processes including glycolysis. A weakening of these predicted causal interactions in schizophrenia tissues support a model in which redox-sensitive s-acylation drives the functional retooling of membrane-associated signalling complexes in inflammatory microglia, specifically during schizophrenia. Such metabolic reversal events allow immune cells like microglia to transition from sensitive to responsive phenotypes during oxidative stress in the CNS, thereby reducing membrane microdomain stability, while inducing cellular apoptosis (Fisher, 2018; He et al., 2023; Lynes et al., 2012). Although many such schizophrenia-sensitive s-acylation dependent protein interaction (causal) models could be predicted from the results of this study, these findings would require thorough experimental validation.

By suggesting the dynamic molecular, cellular, and clinical consequences of differential s-acylation (and s-acylated proteins) in schizophrenia, this report demonstrated s-acylation to actively govern the microglial adaptation processes under inflammation-induced oxidative stress in the human brain. Additionally, it also introduces a framework for integrating the results of differential expression and differential network analyses for the derivation of dysregulated proteins in disease, injuries or treatment. At its core, this report provides the first proteomic catalogue of microglia antioxidants undergoing inflammation-responsive s-acylation and their conditional protein interactors in the brain, thereby laying a conceptual foundation for future investigations on the role of dynamic s-acylation in inflammatory brain dysfunctions.

However, an extensive (and thorough) experimental validation of the reported ‘metabolically flexible’ proteins and protein-interactions within schizophrenia-associated primary microglia remains a goal for future work.

## Author Approvals

All authors have seen and approved the manuscript, and it hasn’t been published elsewhere.

## Declaration

The authors hold full accountability for the originality and accuracy of the submitted manuscript. It has been drafted in a scientific (IMRaD) reports format for easy comprehension.

## Data availability statement

The data that support the findings of this study are available from the corresponding author upon reasonable request.

## Ethics Approval and Safety Statement

The research work was approved and certified by the Shiv Nadar IoE Ethics Committee [SNIoE/2024/0026].

All experimental work were conducted in Biosafety level 2 (BSL-2) cell culture facilities. All postmortem tissues were obtained and studied after getting ethical clearances from the concerned internal ethics committee.

## Patient Consent Statement

Patient consent statements were recorded by the Human Brain Bank at the Human Brain Tissue Repository for Neurobiological Studies (HBTR), National Institute of Mental Health and Neurosciences (NIMHANS), Bengaluru, before the dissection of postmortem autopsies. The biological human tissues were provided free of cost by the Human Brain Bank, HBTR, NIMHANS, Bengaluru as an academic activity. All tissues were provided with proper identifiers and details (like postmortem delay, anatomical area, age and gender) for research work, without disclosing the patient identities.

## Permission to Reproduce Material from Other Sources

Data reproduced from databases are attributed with a Creative Commons Attribution 4.0 International (CC BY 4.0) License for academic usage and has been duly cited in text. All other sources have been carefully credited in text as citations and bibliography references.

## Graphical Summary. Dynamic protein s-acylation regulates the inflammatory plasticity of microglia

Immune-polarized microglia is a primary inducer of stress, mainly oxidative stress, in the brain. During stressful states, the differential activities and localization of redox-responsive proteins are attributed to dynamic s-acylation. We performed comprehensive proteomic studies of human microglia and human brain postmortem autopsies to characterize inflammation-responsive protein s-acylation libraries associated with microglia-induced inflammation, stress and a redox-related brain disorder, schizophrenia. 908 microglial proteins, including 13 redox-regulatory proteins, were annotated to be differentially processed through inflammation-responsive s-acylation.

---

**Supplementary Figure 1.**
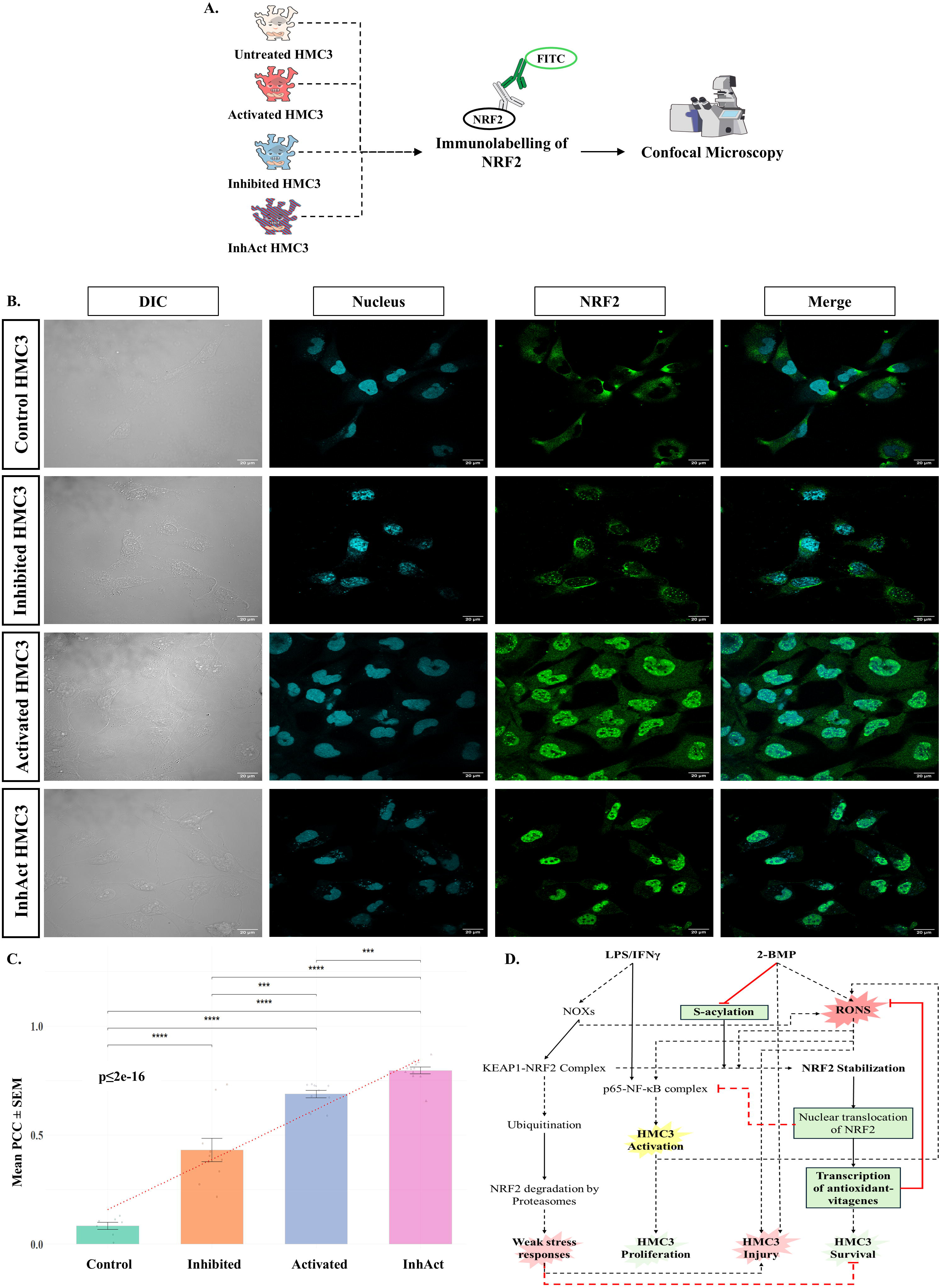
Conditional interactions of variably ‘s-acylable’ antioxidants with other proteins which can undergo inflammation-responsive s-acylation in microglia (identified in MLCC-MS studies of HMC3 microglia). Only high-confidence interactions (featuring consensus edge weights equal to or more than 100) are shown in the protein network graphs to reduce visual clutter. **A.** Protein network graphs generated by DifCoExp for proteomic analysis results obtained from (i) control and, (ii) schizophrenia postmortem autopsies. **B.** Protein network graphs generated by GLASSO for proteomic analysis results obtained from (i) control and, (ii) schizophrenia postmortem autopsies. **C.** Protein network graphs generated by PCC for proteomic analysis results obtained from (i) control and, (ii) schizophrenia postmortem autopsies. Red nodes indicate the MLCC-MS identified ‘s-acylable’ proteins (or complexes) uniquely enriched in LPS/IFNy activated HMC3; Green nodes indicate the MLCC-MS identified ‘s-acylable’ proteins (or protein complexes) uniquely enriched in untreated HMC3; Blue nodes indicate the MLCC-MS identified ‘s-acylable’ proteins (or protein complexes) detected in both untreated and LPS/IFNy activated HMC3. Each of the edges are labelled with respective weights averaged from differential network predictions made by respective network predictors. A higher weight is shaded darker and suggests a stronger correlation of expression between the nodes. The differentially s-acylable NRF2-antioxidants feature Red borders.

## References

1. Anam, Z., Kumari, G., Mukherjee, S., Rex, D. A. B., Biswas, S., Maurya, P., Ravikumar, S., Gupta, N., Kushawaha, A. K., Sah, R. K., Chaurasiya, A., Singhal, J., Singh, N., Kaushik, S., Prasad, T. S. K., Pati, S., Ranganathan, A., & Singh, S. (2022). Complementary crosstalk between palmitoylation and phosphorylation events in MTIP regulates its role during Plasmodium falciparum invasion. Frontiers in Cellular and Infection Microbiology, 12. 10.3389/fcimb.2022.924424

2. Antorini, M., Breme, U., Caccia, P., Grassi, C., Lebrun, S., Orsini, G., Taylor, G., Valsasina, B., Marengo, E., Todeschini, R., Andersson, C., Gellerfors, P., & Gustafsson, J.-G. (1997). Hydroxylamine-Induced Cleavage of the Asparaginyl–Glycine Motif in the Production of Recombinant Proteins: The Case of Insulin-like Growth Factor I. Protein Expression and Purification, 11(1), 135–147. 10.1006/prep.1997.0771

3. Arnold, S. J. M., Ivleva, E. I., Gopal, T. A., Reddy, A. P., Jeon-Slaughter, H., Sacco, C. B., Francis, A. N., Tandon, N., Bidesi, A. S., Witte, B., Poudyal, G., Pearlson, G. D., Sweeney, J. A., Clementz, B. A., Keshavan, M. S., & Tamminga, C. A. (2015). Hippocampal Volume Is Reduced in Schizophrenia and Schizoaffective Disorder But Not in Psychotic Bipolar I Disorder Demonstrated by Both Manual Tracing and Automated Parcellation (FreeSurfer). Schizophrenia Bulletin, 41(1), 233–249. 10.1093/schbul/sbu009

4. Ayana, R., Singh, S., & Pati, S. (2018). Deconvolution of Human Brain Cell Type Transcriptomes Unraveled Microglia-Specific Potential Biomarkers. Frontiers in Neurology, 9. 10.3389/fneur.2018.00266

5. Barger, S. W., Goodwin, M. E., Porter, M. M., & Beggs, M. L. (2007). Glutamate release from activated microglia requires the oxidative burst and lipid peroxidation. Journal of Neurochemistry, 101(5), 1205–1213. 10.1111/j.1471-4159.2007.04487.x

6. Bian, J., Sze, Y.-H., Tse, D. Y.-Y., To, C.-H., McFadden, S. A., Lam, C. S.-Y., Li, K.-K., & Lam, T. C. (2021). SWATH Based Quantitative Proteomics Reveals Significant Lipid Metabolism in Early Myopic Guinea Pig Retina. International Journal of Molecular Sciences, 22(9), 4721. 10.3390/ijms22094721

7. Bitanihirwe, B. K. Y., & Woo, T.-U. W. (2014). Perineuronal Nets and Schizophrenia: The Importance of Neuronal Coatings. Neuroscience and Biobehavioral Reviews, 45, 85–99. 10.1016/j.neubiorev.2014.03.018

8. Blanc, M., David, F., Abrami, L., Migliozzi, D., Armand, F., Bürgi, J., & van der Goot, F. G. (2015). SwissPalm: Protein Palmitoylation database. F1000Research, 4, 261. 10.12688/f1000research.6464.1

9. Brisch, R., Wojtylak, S., Saniotis, A., Steiner, J., Gos, T., Kumaratilake, J., Henneberg, M., & Wolf, R. (2022). The role of microglia in neuropsychiatric disorders and suicide. European Archives of Psychiatry and Clinical Neuroscience, 272(6), 929–945. 10.1007/s00406-021-01334-z

10. Cai, J., Song, M., Li, M., Merchant, M., Benz, F., McClain, C., & Klein, J. (2024). Site-Specific Identification of Protein S-Acylation by IodoTMT0 Labeling and Immobilized Anti-TMT Antibody Resin Enrichment. Journal of Proteome Research, 23(2), 673–683. 10.1021/acs.jproteome.3c00525

11. Castillo-Villanueva, A., Reyes-Vivas, H., & Oria-Hernández, J. (2023). Comparison of cysteine content in whole proteomes across the three domains of life. PLOS ONE, 18(11), e0294268. 10.1371/journal.pone.0294268

12. Cebula, M., Moolla, N., Capovilla, A., & Arnér, E. S. J. (2013). The Rare TXNRD1_v3 (“v3”) Splice Variant of Human Thioredoxin Reductase 1 Protein Is Targeted to Membrane Rafts by N-Acylation and Induces Filopodia Independently of Its Redox Active Site Integrity. The Journal of Biological Chemistry, 288(14), 10002–10011. 10.1074/jbc.M112.445932

13. Chen, X., Shi, C., He, M., Xiong, S., & Xia, X. (2023). Endoplasmic reticulum stress: Molecular mechanism and therapeutic targets. Signal Transduction and Targeted Therapy, 8(1), 352. 10.1038/s41392-023-01570-w

14. Colonna, M., & Butovsky, O. (2017). Microglia Function in the Central Nervous System During Health and Neurodegeneration. Annual Review of Immunology, 35, 441–468. 10.1146/annurev-immunol-051116-052358

15. Cronkite, D. A., & Strutt, T. M. (2018). The Regulation of Inflammation by Innate and Adaptive Lymphocytes. Journal of Immunology Research, 2018, 1467538. 10.1155/2018/1467538

16. Davalieva, K., Maleva Kostovska, I., & Dwork, A. J. (2016). Proteomics Research in Schizophrenia. Frontiers in Cellular Neuroscience, 10. 10.3389/fncel.2016.00018

17. De Kleijn, K. M. A., Straasheijm, K. R., Zuure, W. A., & Martens, G. J. M. (2022). Molecular Signature of Neuroinflammation Induced in Cytokine-Stimulated Human Cortical Spheroids. Biomedicines, 10(5), 1025. 10.3390/biomedicines10051025

18. De Simoni, S., Linard, D., Hermans, E., Knoops, B., & Goemaere, J. (2013). Mitochondrial peroxiredoxin-5 as potential modulator of mitochondria-ER crosstalk in MPP+-induced cell death. Journal of Neurochemistry, 125(3), 473–485. 10.1111/jnc.12117

19. del Toro, N., Shrivastava, A., Ragueneau, E., Meldal, B., Combe, C., Barrera, E., Perfetto, L., How, K., Ratan, P., Shirodkar, G., Lu, O., Mészáros, B., Watkins, X., Pundir, S., Licata, L., Iannuccelli, M., Pellegrini, M., Martin, M. J., Panni, S.,…Hermjakob, H. (2021). The IntAct database: Efficient access to fine-grained molecular interaction data. Nucleic Acids Research, 50(D1), D648–D653. 10.1093/nar/gkab1006

20. Demichev, V., Messner, C. B., Vernardis, S. I., Lilley, K. S., & Ralser, M. (2020). DIA-NN: Neural networks and interference correction enable deep proteome coverage in high throughput. Nature Methods, 17(1), 41–44. 10.1038/s41592-019-0638-x

21. Demyanenko, Y., Sui, X., Giltrap, A. M., Davis, B. G., Kuster, B., & Mohammed, S. (2025). Addressing NHS Chemistry: Efficient Quenching of Excess TMT Reagent and Reversing TMT Overlabeling in Proteomic Samples by Methylamine. Molecular & Cellular ProteomicslJ: MCP, 24(4), 100948. 10.1016/j.mcpro.2025.100948

22. Ermakov, E. A., Dmitrieva, E. M., Parshukova, D. A., Kazantseva, D. V., Vasilieva, A. R., & Smirnova, L. P. (2021). Oxidative Stress-Related Mechanisms in Schizophrenia Pathogenesis and New Treatment Perspectives. Oxidative Medicine and Cellular Longevity, 2021, 8881770. 10.1155/2021/8881770

23. Ernst, A. M., Toomre, D., & Bogan, J. S. (2019). Acylation – A New Means to Control Traffic Through the Golgi. Frontiers in Cell and Developmental Biology, 7. 10.3389/fcell.2019.00109

24. Fisher, A. B. (2018). The phospholipase A2 activity of peroxiredoxin 6. Journal of Lipid Research, 59(7), 1132–1147. 10.1194/jlr.R082578

25. Fujita, H., Aoki, H., Ajioka, I., Yamazaki, M., Abe, M., Oh-Nishi, A., Sakimura, K., & Sugihara, I. (2014). Detailed Expression Pattern of Aldolase C (Aldoc) in the Cerebellum, Retina and Other Areas of the CNS Studied in Aldoc-Venus Knock-In Mice. PLOS ONE, 9(1), e86679. 10.1371/journal.pone.0086679

26. Go, Y.-M., Chandler, J. D., & Jones, D. P. (2015). The Cysteine Proteome. Free Radical Biology & Medicine, 84, 227–245. 10.1016/j.freeradbiomed.2015.03.022

27. Guns, J., Vanherle, S., Hendriks, J. J. A., & Bogie, J. F. J. (2022). Protein Lipidation by Palmitate Controls Macrophage Function. Cells, 11(3), 565. 10.3390/cells11030565

28. He, Q., Qu, M., Shen, T., Su, J., Xu, Y., Xu, C., Barkat, M. Q., Cai, J., Zhu, H., Zeng, L.-H., & Wu, X. (2023). Control of mitochondria-associated endoplasmic reticulum membranes by protein S-palmitoylation: Novel therapeutic targets for neurodegenerative diseases. Ageing Research Reviews, 87, 101920. 10.1016/j.arr.2023.101920

29. Hetz, C., & Saxena, S. (2017). ER stress and the unfolded protein response in neurodegeneration. Nature Reviews. Neurology, 13(8), 477–491. 10.1038/nrneurol.2017.99

30. Hristovska, I., & Pascual, O. (2016). Deciphering Resting Microglial Morphology and Process Motility from a Synaptic Prospect. Frontiers in Integrative Neuroscience, 9, 73. 10.3389/fnint.2015.00073

31. Hwang, J. S., Vo, T. T. L., Kim, M., Cha, E. H., Mun, K. C., Ha, E., & Seo, J. H. (2025). Involvement of RhoA/ROCK Signaling Pathway in Methamphetamine-Induced Blood-Brain Barrier Disruption. Biomolecules, 15(3), 340. 10.3390/biom15030340

32. Jaronen, M., Goldsteins, G., & Koistinaho, J. (2014). ER stress and unfolded protein response in amyotrophic lateral sclerosis—A controversial role of protein disulphide isomerase. Frontiers in Cellular Neuroscience, 8. 10.3389/fncel.2014.00402

33. Jayaram, S., & Krishnamurthy, P. T. (2021). Role of microgliosis, oxidative stress and associated neuroinflammation in the pathogenesis of Parkinson’s disease: The therapeutic role of Nrf2 activators. Neurochemistry International, 145, 105014. 10.1016/j.neuint.2021.105014

34. Jiang, S., Qian, Q., Zhu, T., Zong, W., Shang, Y., Jin, T., Zhang, Y., Chen, M., Wu, Z., Chu, Y., Zhang, R., Luo, S., Jing, W., Zou, D., Bao, Y., Xiao, J., & Zhang, Z. (2023). Cell Taxonomy: A curated repository of cell types with multifaceted characterization. Nucleic Acids Research, 51(D1), D853–D860. 10.1093/nar/gkac816

35. Junjappa, R. P., Patil, P., Bhattarai, K. R., Kim, H.-R., & Chae, H.-J. (2018). IRE1α Implications in Endoplasmic Reticulum Stress-Mediated Development and Pathogenesis of Autoimmune Diseases. Frontiers in Immunology, 9. 10.3389/fimmu.2018.01289

36. Kettenmann, H., Hanisch, U.-K., Noda, M., & Verkhratsky, A. (2011). Physiology of Microglia. Physiological Reviews, 91(2), 461–553. 10.1152/physrev.00011.2010

37. Kigerl, K. A., Ankeny, D. P., Garg, S. K., Wei, P., Guan, Z., Lai, W., McTigue, D. M., Banerjee, R., & Popovich, P. G. (2012). System xc− regulates microglia and macrophage glutamate excitotoxicity *in vivo*. *Experimental Neurology*, Special Issue: Stress and Neurological Disease, 233(1), 333–341. 10.1016/j.expneurol.2011.10.025

38. Kim, S., Lee, W., Jo, H., Sonn, S.-K., Jeong, S.-J., Seo, S., Suh, J., Jin, J., Kweon, H. Y., Kim, T. K., Moon, S. H., Jeon, S., Kim, J. W., Kim, Y. R., Lee, E.-W., Shin, H. K., Park, S. H., & Oh, G. T. (2022). The antioxidant enzyme Peroxiredoxin-1 controls stroke-associated microglia against acute ischemic stroke. Redox Biology, 54, 102347. 10.1016/j.redox.2022.102347

39. Knudsen, E., Tadje, J., Coggins, C., & Venketaraman, V. (2026). Glutathione and neurodegenerative diseases: Immunopharmacological implications. Frontiers in Pharmacology, 16. 10.3389/fphar.2025.1737199

40. Kochunov, P., Coyle, T. R., Rowland, L. M., Jahanshad, N., Thompson, P. M., Kelly, S., Du, X., Sampath, H., Bruce, H., Chiappelli, J., Ryan, M., Fisseha, F., Savransky, A., Adhikari, B., Chen, S., Paciga, S. A., Whelan, C. D., Xie, Z., Hyde, C. L.,…Hong, L. E. (2017). Association of White Matter With Core Cognitive Deficits in Patients With Schizophrenia. JAMA Psychiatry, 74(9), 958–966. 10.1001/jamapsychiatry.2017.2228

41. Kohler, D., Staniak, M., Tsai, T.-H., Huang, T., Shulman, N., Bernhardt, O. M., MacLean, B. X., Nesvizhskii, A. I., Reiter, L., Sabido, E., Choi, M., & Vitek, O. (2023). MSstats Version 4.0: Statistical Analyses of Quantitative Mass Spectrometry-Based Proteomic Experiments with Chromatography-Based Quantification at Scale. Journal of Proteome Research, 22(5), 1466–1482. 10.1021/acs.jproteome.2c00834

42. Kong, W., Jiang, P., Miao, X., Sang, B., Hu, S., & Feng, L. (2025). The role of Atp2a2-mediated calcium imbalance and endoplasmic reticulum stress in hydrocortisone-induced neurotoxicity. Cell Stress & Chaperones, 30(6), 100112. 10.1016/j.cstres.2025.100112

43. Kumari, G., Rex, D. A. B., Goswami, S., Mukherjee, S., Biswas, S., Maurya, P., Jain, R., Garg, S., Prasad, T. S. K., Pati, S., Ramalingam, S., Mohandas, N., & Singh, S. (2022). Dynamic Palmitoylation of Red Cell Membrane Proteins Governs Susceptibility to Invasion by the Malaria Parasite, Plasmodium falciparum. ACS Infectious Diseases, 8(10), 2106–2118. 10.1021/acsinfecdis.2c00199

44. Kwon, Y., Ju, S., Kaushal, P., Lee, J.-W., & Lee, C. (2018). Neutralizing the Detrimental Effect of an N-Hydroxysuccinimide Quenching Reagent on Phosphopeptide in Quantitative Proteomics. Analytical Chemistry, 90(5), 3019–3023. 10.1021/acs.analchem.7b04678

45. Lee, H., Kim, M., Kim, S. H., Lee, J., Lee, T. Y., Rhee, S. J., Roh, S., Baik, M., Jung, H. Y., Kim, H., Han, D. H., Ha, K., Ahn, Y. M., & Kwon, J. S. (2024). Proteomic profiling in the progression of psychosis: Analysis of clinical high-risk, first episode psychosis, and healthy controls. Journal of Psychiatric Research, 169, 264–271. 10.1016/j.jpsychires.2023.11.031

46. Lee, J. E., Lee, J. Y., Trembly, J., Wilusz, J., Tian, B., & Wilusz, C. J. (2012). The PARN Deadenylase Targets a Discrete Set of mRNAs for Decay and Regulates Cell Motility in Mouse Myoblasts. PLoS Genetics, 8(8), e1002901. 10.1371/journal.pgen.1002901

47. Li, W., Shen, J., Zhuang, A., Wang, R., Li, Q., Rabata, A., Zhang, Y., & Cao, D. (2025). Palmitoylation: An emerging therapeutic target bridging physiology and disease. Cellular & Molecular Biology Letters, 30(1), 98. 10.1186/s11658-025-00776-w

48. Li, Y., Xia, X., Wang, Y., & Zheng, J. C. (2022). Mitochondrial dysfunction in microglia: A novel perspective for pathogenesis of Alzheimer’s disease. Journal of Neuroinflammation, 19, 248. 10.1186/s12974-022-02613-9

49. Lieberman, J. A., Girgis, R. R., Brucato, G., Moore, H., Provenzano, F., Kegeles, L., Javitt, D., Kantrowitz, J., Wall, M. M., Corcoran, C. M., Schobel, S. A., & Small, S. A. (2018). Hippocampal dysfunction in the pathophysiology of schizophrenia: A selective review and hypothesis for early detection and intervention. Molecular Psychiatry, 23(8), 1764–1772. 10.1038/mp.2017.249

50. Liemburg-Apers, D. C., Willems, P. H. G. M., Koopman, W. J. H., & Grefte, S. (2015). Interactions between mitochondrial reactive oxygen species and cellular glucose metabolism. Archives of Toxicology, 89(8), 1209–1226. 10.1007/s00204-015-1520-y

51. Lin, Z., Li, J., Ji, T., Wu, Y., Gao, K., & Jiang, Y. (2021). ATP1A1 de novo Mutation-Related Disorders: Clinical and Genetic Features. Frontiers in Pediatrics, 9, 657256. 10.3389/fped.2021.657256

52. Liu, D., Zhao, Z., She, Y., Zhang, L., Chen, X., Ma, L., & Cui, J. (2022). TRIM14 inhibits OPTN-mediated autophagic degradation of KDM4D to epigenetically regulate inflammation. Proceedings of the National Academy of Sciences of the United States of America, 119(7), e2113454119. 10.1073/pnas.2113454119

53. Liu, J., Wang, S., Fan, L., Zhou, X., Zhang, S., Wang, Q., Dong, P., & Yu, B. (2025). Protein regulatory network mediated by palmitoylation modifications in the pathological progression of Parkinson’s disease: A narrative review. Frontiers in Immunology, 16, 1615001. 10.3389/fimmu.2025.1615001

54. Liu, X., Peyton, K. J., Ensenat, D., Wang, H., Schafer, A. I., Alam, J., & Durante, W. (2005). Endoplasmic Reticulum Stress Stimulates Heme Oxygenase-1 Gene Expression in Vascular Smooth Muscle: ROLE IN CELL SURVIVAL*. Journal of Biological Chemistry, 280(2), 872–877. 10.1074/jbc.M410413200

55. Lynes, E. M., Bui, M., Yap, M. C., Benson, M. D., Schneider, B., Ellgaard, L., Berthiaume, L. G., & Simmen, T. (2012). Palmitoylated TMX and calnexin target to the mitochondriaLassociated membrane. The EMBO Journal, 31(2), 457–470. 10.1038/emboj.2011.384

56. Ma, Q. (2013). Role of Nrf2 in Oxidative Stress and Toxicity. Annual Review of Pharmacology and Toxicology, 53, 401–426. 10.1146/annurev-pharmtox-011112-140320

57. MacLean, B., Tomazela, D. M., Shulman, N., Chambers, M., Finney, G. L., Frewen, B., Kern, R., Tabb, D. L., Liebler, D. C., & MacCoss, M. J. (2010). Skyline: An open source document editor for creating and analyzing targeted proteomics experiments. Bioinformatics, 26(7), 966–968. 10.1093/bioinformatics/btq054

58. Martin, B. R., & Cravatt, B. F. (2009). Large-Scale Profiling of Protein Palmitoylation in Mammalian Cells. Nature Methods, 6(2), 135–138. 10.1038/nmeth.1293

59. Mattick, J., & Amaral, P. (2022). The Human Genome. In RNA, the Epicenter of Genetic Information: A new understanding of molecular biology. CRC Press. 10.1201/9781003109242-11

60. McBean, G. J., Aslan, M., Griffiths, H. R., & Torrão, R. C. (2015). Thiol redox homeostasis in neurodegenerative disease. Redox Biology, 5, 186–194. 10.1016/j.redox.2015.04.004

61. McCarty, M. F., & DiNicolantonio, J. J. (2015). An increased need for dietary cysteine in support of glutathione synthesis may underlie the increased risk for mortality associated with low protein intake in the elderly. Age, 37(5), 96. 10.1007/s11357-015-9823-8

62. Meng, F.-L., Huang, X.-L., Qin, W.-Y., Liu, K.-B., Wang, Y., Li, M., Ren, Y.-H., Li, Y.-Z., & Sun, Y.-M. (2023). singleCellBase: A high-quality manually curated database of cell markers for single cell annotation across multiple species. Biomarker Research, 11(1), 83. 10.1186/s40364-023-00523-3

63. Mesquita, F. S., Abrami, L., Linder, M. E., Bamji, S. X., Dickinson, B. C., & van der Goot, F. G. (2024). Mechanisms and functions of protein S-acylation. Nature Reviews. Molecular Cell Biology, 25(6), 488–509. 10.1038/s41580-024-00700-8

64. Messner, C. B., Demichev, V., Bloomfield, N., Yu, J. S. L., White, M., Kreidl, M., Egger, A.-S., Freiwald, A., Ivosev, G., Wasim, F., Zelezniak, A., Jürgens, L., Suttorp, N., Sander, L. E., Kurth, F., Lilley, K. S., Mülleder, M., Tate, S., & Ralser, M. (2021). Ultra-fast proteomics with Scanning SWATH. Nature Biotechnology, 39(7), 846–854. 10.1038/s41587-021-00860-4

65. Milacic, M., Beavers, D., Conley, P., Gong, C., Gillespie, M., Griss, J., Haw, R., Jassal, B., Matthews, L., May, B., Petryszak, R., Ragueneau, E., Rothfels, K., Sevilla, C., Shamovsky, V., Stephan, R., Tiwari, K., Varusai, T., Weiser, J.,…D’Eustachio, P. (2024). The Reactome Pathway Knowledgebase 2024. Nucleic Acids Research, 52(D1), D672–D678. 10.1093/nar/gkad1025

66. Mo, Z.-T., Zheng, J., & Liao, Y. (2021). Icariin inhibits the expression of IL-1β, IL-6 and TNF-α induced by OGD/R through the IRE1/XBP1s pathway in microglia. Pharmaceutical Biology, 59(1), 1471–1477. 10.1080/13880209.2021.1991959

67. Mondal, A., Munan, S., Saxena, I., Mukherjee, S., Upadhyay, P., Gupta, N., Dar, W., Samanta, A., Singh, S., & Pati, S. (2024). G6PD deficiency mediated impairment of iNOS and lysosomal acidification affecting phagocytotic clearance in microglia in response to SARS-CoV-2. Biochimica et Biophysica Acta (BBA) - Molecular Basis of Disease, 1870(7), 167444. 10.1016/j.bbadis.2024.167444

68. Mudge, J. M., Carbonell-Sala, S., Diekhans, M., Martinez, J. G., Hunt, T., Jungreis, I., Loveland, J. E., Arnan, C., Barnes, I., Bennett, R., Berry, A., Bignell, A., Cerdán-Vélez, D., Cochran, K., Cortés, L. T., Davidson, C., Donaldson, S., Dursun, C., Fatima, R.,…Frankish, A. (2024). GENCODE 2025: Reference gene annotation for human and mouse. Nucleic Acids Research, 53(D1), D966–D975. 10.1093/nar/gkae1078

69. Murray, A. J., Rogers, J. C., Katshu, M. Z. U. H., Liddle, P. F., & Upthegrove, R. (2021). Oxidative Stress and the Pathophysiology and Symptom Profile of Schizophrenia Spectrum Disorders. Frontiers in Psychiatry, 12, 703452. 10.3389/fpsyt.2021.703452

70. Nakajima, K., Ishiwata, M., Weitemier, A. Z., Shoji, H., Monai, H., Miyamoto, H., Yamakawa, K., Miyakawa, T., McHugh, T. J., & Kato, T. (2021). Brain-specific heterozygous loss-of-function of ATP2A2, endoplasmic reticulum Ca2+ pump responsible for Darier’s disease, causes behavioral abnormalities and a hyper-dopaminergic state. Human Molecular Genetics, 30(18), 1762–1772. 10.1093/hmg/ddab137

71. Nazeri, A., Chakravarty, M. M., Felsky, D., Lobaugh, N. J., Rajji, T. K., Mulsant, B. H., & Voineskos, A. N. (2013). Alterations of Superficial White Matter in Schizophrenia and Relationship to Cognitive Performance. Neuropsychopharmacology, 38(10), 1954–1962. 10.1038/npp.2013.93

72. Ngo, V., & Duennwald, M. L. (2022). Nrf2 and Oxidative Stress: A General Overview of Mechanisms and Implications in Human Disease. Antioxidants, 11(12), 2345. 10.3390/antiox11122345

73. Norden, D. M., Trojanowski, P. J., Villanueva, E., Navarro, E., & Godbout, J. P. (2016). Sequential Activation of Microglia and Astrocyte Cytokine Expression Precedes Increased Iba-1 or GFAP Immunoreactivity following Systemic Immune Challenge. Glia, 64(2), 300–316. 10.1002/glia.22930

74. Oughtred, R., Rust, J., Chang, C., Breitkreutz, B., Stark, C., Willems, A., Boucher, L., Leung, G., Kolas, N., Zhang, F., Dolma, S., CoulombeLHuntington, J., ChatrLaryamontri, A., Dolinski, K., & Tyers, M. (2021). The BioGRID database: A comprehensive biomedical resource of curated protein, genetic, and chemical interactions. Protein SciencelJ: A Publication of the Protein Society, 30(1), 187–200. 10.1002/pro.3978

75. Ouyang, P., Cai, Z., Peng, J., Lin, S., Chen, X., Chen, C., Feng, Z., Wang, L., Song, G., & Zhang, Z. (2024). SELENOK-dependent CD36 palmitoylation regulates microglial functions and Aβ phagocytosis. Redox Biology, 70, 103064. 10.1016/j.redox.2024.103064

76. Pace, M., Giorgi, C., Lombardozzi, G., Cimini, A., Castelli, V., & d’Angelo, M. (2025). Exploring the Antioxidant Roles of Cysteine and Selenocysteine in Cellular Aging and Redox Regulation. Biomolecules, 15(8), 1115. 10.3390/biom15081115

77. Pang, Z., Xu, L., Viau, C., Lu, Y., Salavati, R., Basu, N., & Xia, J. (2024). MetaboAnalystR 4.0: A unified LC-MS workflow for global metabolomics. Nature Communications, 15(1), 3675. 10.1038/s41467-024-48009-6

78. Pap, D., Veres-Székely, A., Szebeni, B., & Vannay, Á. (2022). PARK7/DJ-1 as a Therapeutic Target in Gut-Brain Axis Diseases. International Journal of Molecular Sciences, 23(12), 6626. 10.3390/ijms23126626

79. Patir, A., Shih, B., McColl, B. W., & Freeman, T. C. (2019). A core transcriptional signature of human microglia: Derivation and utility in describing region-dependent alterations associated with Alzheimer’s disease. Glia, 67(7), 1240–1253. 10.1002/glia.23572

80. Perkins, D. O., Jeffries, C. D., & Do, K. Q. (2020). Potential Roles of Redox Dysregulation in the Development of Schizophrenia. Biological Psychiatry, 88(4), 326–336. 10.1016/j.biopsych.2020.03.016

81. Pino, L. K., Searle, B. C., Bollinger, J. G., Nunn, B., MacLean, B., & MacCoss, M. J. (2017). The Skyline Ecosystem: Informatics for Quantitative Mass Spectrometry Proteomics. Mass Spectrometry Reviews, 10.1002/mas.21540.

82. Qiu, T., Azizi, S.-A., Pani, S., & Dickinson, B. C. (2025). Dynamic PRDX S-acylation modulates ROS stress and signaling. Cell Chemical Biology, 32(3), 511–519.e5. 10.1016/j.chembiol.2025.01.009

83. Ransohoff, R. M. (2016). How neuroinflammation contributes to neurodegeneration. *Science (New York*, N.Y.), 353(6301), 777–783. 10.1126/science.aag2590

84. Rojo, A. I., McBean, G., Cindric, M., Egea, J., López, M. G., Rada, P., Zarkovic, N., & Cuadrado, A. (2014). Redox Control of Microglial Function: Molecular Mechanisms and Functional Significance. Antioxidants & Redox Signaling, 21(12), 1766–1801. 10.1089/ars.2013.5745

85. Schramm, E., & Waisman, A. (2022). Microglia as Central Protagonists in the Chronic Stress Response. Neurology® Neuroimmunology & Neuroinflammation, 9(6), e200023. 10.1212/NXI.0000000000200023

86. Selvakumar, P., Owens, T. A., David, J. M., Petrelli, N. J., Christensen, B. C., Lakshmikuttyamma, A., & Rajasekaran, A. K. (2014). Epigenetic silencing of Na,K-ATPase β1 subunit gene ATP1B1 by methylation in clear cell renal cell carcinoma. Epigenetics, 9(4), 579–586. 10.4161/epi.27795

87. Shen, L.-F., Chen, Y.-J., Liu, K.-M., Haddad, A. N. S., Song, I.-W., Roan, H.-Y., Chen, L.-Y., Yen, J. J. Y., Chen, Y.-J., Wu, J.-Y., & Chen, Y.-T. (2017). Role of S-Palmitoylation by ZDHHC13 in Mitochondrial function and Metabolism in Liver. Scientific Reports, 7, 2182. 10.1038/s41598-017-02159-4

88. Sherman, B. T., Hao, M., Qiu, J., Jiao, X., Baseler, M. W., Lane, H. C., Imamichi, T., & Chang, W. (2022). DAVID: A web server for functional enrichment analysis and functional annotation of gene lists (2021 update). Nucleic Acids Research, 50(W1), W216–W221. 10.1093/nar/gkac194

89. Sikorski, K., Chmielewski, S., Olejnik, A., Wesoly, J. Z., Heemann, U., Baumann, M., & Bluyssen, H. (2012). STAT1 as a central mediator of IFNγ and TLR4 signal integration in vascular dysfunction. JAK-STAT, 1(4), 241–249. 10.4161/jkst.22469

90. Simpson, D. S. A., & Oliver, P. L. (2020). ROS Generation in Microglia: Understanding Oxidative Stress and Inflammation in Neurodegenerative Disease. Antioxidants, 9(8), 743. 10.3390/antiox9080743

91. Singh, S., Nagalakshmi, D., Sharma, K. K., & Ravichandiran, V. (2021). Natural antioxidants for neuroinflammatory disorders and possible involvement of Nrf2 pathway: A review. Heliyon, 7(2), e06216. 10.1016/j.heliyon.2021.e06216

92. Singh, V., Gera, R., Kushwaha, R., Sharma, A. K., Patnaik, S., & Ghosh, D. (2016). Hijacking microglial glutathione by inorganic arsenic impels bystander death of immature neurons through extracellular cystine/glutamate imbalance. Scientific Reports, 6, 30601. 10.1038/srep30601

93. Szklarczyk, D., Kirsch, R., Koutrouli, M., Nastou, K., Mehryary, F., Hachilif, R., Gable, A. L., Fang, T., Doncheva, N. T., Pyysalo, S., Bork, P., Jensen, L. J., & von Mering, C. (2022). The STRING database in 2023: Protein–protein association networks and functional enrichment analyses for any sequenced genome of interest. Nucleic Acids Research, 51(D1), D638–D646. 10.1093/nar/gkac1000

94. Tan, Y.-L., Yuan, Y., & Tian, L. (2020). Microglial regional heterogeneity and its role in the brain. Molecular Psychiatry, 25(2), 351–367. 10.1038/s41380-019-0609-8

95. The UniProt Consortium. (2025). UniProt: The Universal Protein Knowledgebase in 2025. Nucleic Acids Research, 53(D1), D609–D617. 10.1093/nar/gkae1010

96. Traina, G., Tuszynski, J. A., & Cocchi, M. (2022). Molecular Insights in Psychiatry. International Journal of Molecular Sciences, 23(9), 4878. 10.3390/ijms23094878

97. Trubalski, M., Markiewicz-Gospodarek, A., Żerebiec, M., Poleszak, J., Szczotka, M., Markiewicz, R., Łoza, B., & Szymańczyk, S. (2025). Oxidative Stress-Mediated Neuroinflammation in the Pathophysiology of Schizophrenia. International Journal of Molecular Sciences, 26(22), 11139. 10.3390/ijms262211139

98. Tsoporis, J. N., Ektesabi, A. M., Gupta, S., Izhar, S., Salpeas, V., Rizos, I. K., Kympouropoulos, S. P., dos Santos, C. C., Parker, T. G., & Rizos, E. (2022). A longitudinal study of alterations of circulating DJ-1 and miR203a-3p in association to olanzapine medication in a sample of first episode patients with schizophrenia. Journal of Psychiatric Research, 146, 109–117. 10.1016/j.jpsychires.2021.12.049

99. Vilhardt, F., HaslundLVinding, J., Jaquet, V., & McBean, G. (2017). Microglia antioxidant systems and redox signalling. British Journal of Pharmacology, 174(12), 1719–1732. 10.1111/bph.13426

100. Voigt, D., Scheidt, U., Derfuss, T., Brück, W., & Junker, A. (2017). Expression of the Antioxidative Enzyme Peroxiredoxin 2 in Multiple Sclerosis Lesions in Relation to Inflammation. International Journal of Molecular Sciences, 18(4), 760. 10.3390/ijms18040760

101. Wang, P., Han, X., Mo, B., Huang, G., & Wang, C. (2017). LPS enhances TLR4 expression and IFN-γ production via the TLR4/IRAK/NF-κB signaling pathway in rat pulmonary arterial smooth muscle cells. Molecular Medicine Reports, 16(3), 3111–3116. 10.3892/mmr.2017.6983

102. Wang, Y., Zhu, W., Wang, W., Zhang, J., Hu, D., Shao, H., zhou, Y., Wang, S., & Zhao, L. (2025). The role of protein S-acylation in vascular injury associated with metabolic disorders. Frontiers in Endocrinology, 16, 1643008. 10.3389/fendo.2025.1643008

103. Warrick, K. A., Vallez, C. N., Meibers, H. E., & Pasare, C. (2025). Bidirectional Communication Between the Innate and Adaptive Immune Systems. Annual Review of Immunology, 43(Volume 43, 2025), 489–514. 10.1146/annurev-immunol-083122-040624

104. Wei, Y., Gu, Y., Zhou, Z., Wu, C., Liu, Y., & Sun, H. (2024). TRIM21 Promotes Oxidative Stress and Ferroptosis through the SQSTM1-NRF2-KEAP1 Axis to Increase the Titers of H5N1 Highly Pathogenic Avian Influenza Virus. International Journal of Molecular Sciences, 25(6), 3315. 10.3390/ijms25063315

105. Wen, H., Liu, L., Zhan, L., Liang, D., Li, L., Liu, D., Sun, W., & Xu, E. (2018). Neuroglobin mediates neuroprotection of hypoxic postconditioning against transient global cerebral ischemia in rats through preserving the activity of Na+/K+ ATPases. Cell Death & Disease, 9(6), 635. 10.1038/s41419-018-0656-0

106. Woodburn, S. C., Bollinger, J. L., & Wohleb, E. S. (2021). The semantics of microglia activation: Neuroinflammation, homeostasis, and stress. Journal of Neuroinflammation, 18(1), 258. 10.1186/s12974-021-02309-6

107. Wu, W., Gong, X., Qin, Z., & Wang, Y. (2025). Molecular mechanisms of excitotoxicity and their relevance to the pathogenesis of neurodegenerative diseases—An update. Acta Pharmacologica Sinica, 46(12), 3129–3142. 10.1038/s41401-025-01576-w

108. Xiao, T., Wan, J., Qu, H., & Li, Y. (2021). Tripartite-motif protein 21 knockdown extenuates LPS-triggered neurotoxicity by inhibiting microglial M1 polarization via suppressing NF-κB-mediated NLRP3 inflammasome activation. Archives of Biochemistry and Biophysics, 706, 108918. 10.1016/j.abb.2021.108918

109. Xin, D., Chu, X., Bai, X., Ma, W., Yuan, H., Qiu, J., Liu, C., Li, T., Zhou, X., Chen, W., Liu, D., & Wang, Z. (2018). L-Cysteine suppresses hypoxia-ischemia injury in neonatal mice by reducing glial activation, promoting autophagic flux and mediating synaptic modification via H2S formation. Brain, Behavior, and Immunity, 73, 222–234. 10.1016/j.bbi.2018.05.007

110. Yamada, S., Takahashi, S., Malchow, B., Papazova, I., Stöcklein, S., Ertl-Wagner, B., Papazov, B., Kumpf, U., Wobrock, T., Keller-Varady, K., Hasan, A., Falkai, P., Wagner, E., Raabe, F. J., & Keeser, D. (2022). Cognitive and functional deficits are associated with white matter abnormalities in two independent cohorts of patients with schizophrenia. European Archives of Psychiatry and Clinical Neuroscience, 272(6), 957–969. 10.1007/s00406-021-01363-8

111. Yang, J., Carroll, K. S., & Liebler, D. C. (2016). The Expanding Landscape of the Thiol Redox Proteome*. Molecular & Cellular Proteomics, 15(1), 1–11. 10.1074/mcp.O115.056051

112. Zgorzynska, E., Dziedzic, B., & Walczewska, A. (2021). An Overview of the Nrf2/ARE Pathway and Its Role in Neurodegenerative Diseases. International Journal of Molecular Sciences, 22(17), 9592. 10.3390/ijms22179592

113. Zhang, F., Yan, Y., Peng, W., Wang, L., Wang, T., Xie, Z., Luo, H., Zhang, J., & Dong, W. (2021). PARK7 promotes repair in early steroid-induced osteonecrosis of the femoral head by enhancing resistance to stress-induced apoptosis in bone marrow mesenchymal stem cells via regulation of the Nrf2 signaling pathway. Cell Death & Disease, 12(10), 940. 10.1038/s41419-021-04226-1

114. Zhang, H., Yang, S., Lu, Y.-L., Zhou, L.-Q., Dong, M.-H., Chu, Y.-H., Pang, X.-W., Chen, L., Xu, L.-L., Zhang, L.-Y., Zhu, L.-F., Xu, T., Wang, W., Shang, K., Tian, D.-S., & Qin, C. (2024). Microglial Nrf2-mediated lipid and iron metabolism reprogramming promotes remyelination during white matter ischemia. Redox Biology, 79, 103473. 10.1016/j.redox.2024.103473

115. Zhang, X., Lan, Y., Xu, J., Quan, F., Zhao, E., Deng, C., Luo, T., Xu, L., Liao, G., Yan, M., Ping, Y., Li, F., Shi, A., Bai, J., Zhao, T., Li, X., & Xiao, Y. (2019). CellMarker: A manually curated resource of cell markers in human and mouse. Nucleic Acids Research, 47(D1), D721–D728. 10.1093/nar/gky900

116. Zhang, Y., Sloan, S. A., Clarke, L. E., Caneda, C., Plaza, C. A., Blumenthal, P. D., Vogel, H., Steinberg, G. K., Edwards, M. S. B., Li, G., Duncan, J. A., Cheshier, S. H., Shuer, L. M., Chang, E. F., Grant, G. A., Hayden Gephart, M. G., & Barres, B. A. (2016). Purification and characterization of progenitor and mature human astrocytes reveals transcriptional and functional differences with mouse. Neuron, 89(1), 37–53. 10.1016/j.neuron.2015.11.013

117. Zhou, X., Chu, X., Xin, D., Li, T., Bai, X., Qiu, J., Yuan, H., Liu, D., Wang, D., & Wang, Z. (2019). L-Cysteine-Derived H2S Promotes Microglia M2 Polarization via Activation of the AMPK Pathway in Hypoxia-Ischemic Neonatal Mice. Frontiers in Molecular Neuroscience, 12, 58. 10.3389/fnmol.2019.00058

118. Zhou, Z., Jia, X., Xue, Q., Dou, Z., Ma, Y., Zhao, Z., Jiang, Z., He, B., Jin, Q., & Wang, J. (2014). TRIM14 is a mitochondrial adaptor that facilitates retinoic acid-inducible gene-I–like receptor-mediated innate immune response. Proceedings of the National Academy of Sciences of the United States of America, 111(2), E245–E254. 10.1073/pnas.1316941111

