## Supplementary figures and images for "Dynamic s-acylation of schizophrenia-linked antioxidants explicate the redox flexibility of human microglia during inflammation"

### Supplementary Figure 1

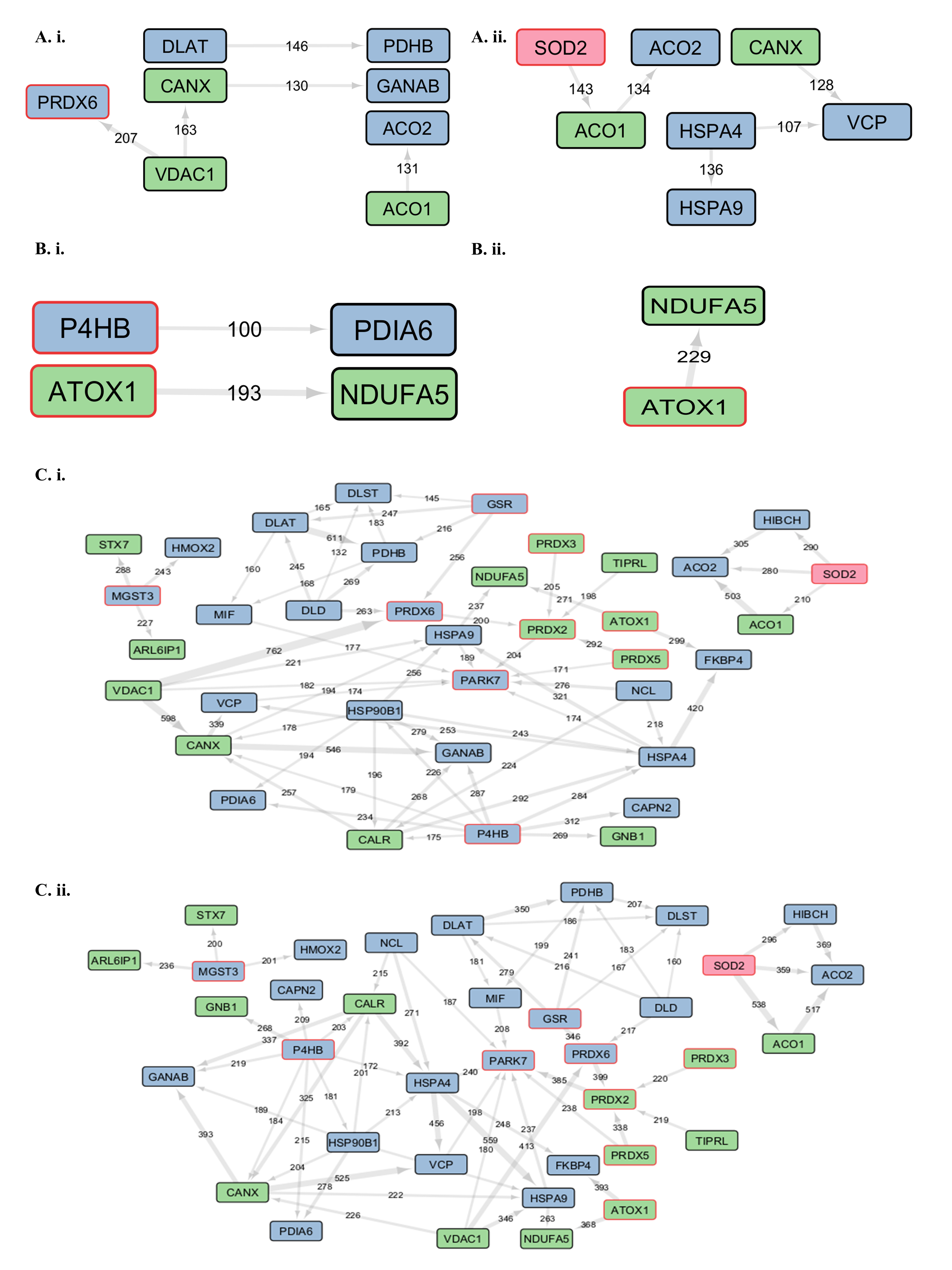
