## Supplementary File 1 for "Dynamic s-acylation of schizophrenia-linked antioxidants explicate the redox flexibility of human microglia during inflammation"

DIA-NN 2.2.0 Academia  (Data-Independent Acquisition by Neural Networks)
Compiled on May 29 2025 21:29:29
Current date and time: Sat Jul 12 00:41:36 2025
CPU: GenuineIntel Intel(R) Core(TM) i5-6300U CPU @ 2.40GHz
SIMD instructions: AVX AVX2 FMA SSE4.1 SSE4.2 
Logical CPU cores: 4
diann.exe --lib  --threads 3 --verbose 1 --out E:\mass_spec\final analyses_diann\click_chem\out\sphumprots_analysis_out.parquet --qvalue 0.05 --matrices --temp E:\mass_spec\final analyses_diann\click_chem\click_palm_withoutcontam\temp --out-lib E:\mass_spec\final analyses_diann\click_chem\click_palm_withoutcontam\lib\uniprotcan_wp_wocontam.parquet --gen-spec-lib --predictor --fasta camprotR_240512_cRAP_20190401_full_tags.fasta --cont-quant-exclude cRAP- --fasta E:\mass_spec\final analyses_diann\click_chem\click_palm_withoutcontam\clean_humwholeprot_reviewed_can.fasta --fasta-search --min-fr-mz 50 --max-fr-mz 1300 --met-excision --min-pep-len 7 --max-pep-len 30 --min-pr-mz 300 --max-pr-mz 1300 --min-pr-charge 1 --max-pr-charge 4 --cut K*,R*,!*P --missed-cleavages 2 --unimod4 --var-mods 3 --var-mod UniMod:35,15.994915,M --reanalyse --rt-profiling --duplicate-proteins 

Thread number set to 3
Output will be filtered at 0.05 FDR
Precursor/protein x samples expression level matrices will be saved along with the main report
A spectral library will be generated
Deep learning will be used to generate a new in silico spectral library from peptides provided
Peptides corresponding to protein sequence IDs tagged with cRAP- will be excluded from normalisation as well as quantification of protein groups that do not include proteins bearing the tag
DIA-NN will carry out FASTA digest for in silico lib generation
Min fragment m/z set to 50
Max fragment m/z set to 1300
N-terminal methionine excision enabled
Min peptide length set to 7
Max peptide length set to 30
Min precursor m/z set to 300
Max precursor m/z set to 1300
Min precursor charge set to 1
Max precursor charge set to 4
In silico digest will involve cuts at K*,R*
But excluding cuts at *P
Maximum number of missed cleavages set to 2
Cysteine carbamidomethylation enabled as a fixed modification
Maximum number of variable modifications set to 3
Modification UniMod:35 with mass delta 15.9949 at M will be considered as variable
MBR enabled; .quant files will only be saved to disk during the first pass
The spectral library (if generated) will retain the original spectra but will include empirically-aligned RTs
Duplicate proteins in FASTA files will not be skipped
WARNING: peptidoform scoring enabled because variable modifications have been declared; to disable, use --no-peptidoforms
The following variable modifications will be localised: UniMod:35 
WARNING: protein inference will only be performed for precursors identified with global q-value <= 0.05 or q-value <= 0.05.
WARNING: MBR turned off, at least two files are required

0 files will be processed
[0:00] Loading FASTA camprotR_240512_cRAP_20190401_full_tags.fasta
[0:00] Loading FASTA E:\mass_spec\final analyses_diann\click_chem\click_palm_withoutcontam\clean_humwholeprot_reviewed_can.fasta
[0:17] Processing FASTA
[0:25] Assembling elution groups
[0:40] 7707491 precursors generated
[0:40] Gene names missing for some isoforms
[0:40] Library contains 20493 proteins, and 20269 genes
[0:50] [1:14] [104:42] [115:49] [117:10] [117:12] Saving the library to E:\mass_spec\final analyses_diann\click_chem\click_palm_withoutcontam\lib\uniprotcan_wp_wocontam.predicted.speclib
[117:30] Initialising library
