## Supplementary File 2 for "Dynamic s-acylation of schizophrenia-linked antioxidants explicate the redox flexibility of human microglia during inflammation"

DIA-NN 2.2.0 Academia  (Data-Independent Acquisition by Neural Networks)
Compiled on May 29 2025 21:29:29
Current date and time: Wed Jul  9 17:00:56 2025
CPU: GenuineIntel 12th Gen Intel(R) Core(TM) i7-12700
SIMD instructions: AVX AVX2 FMA SSE4.1 SSE4.2 
Logical CPU cores: 20
diann.exe --f D:\Soumyadeep\Cell Click\Data\Cell Run 1 MS Data\Positive Click\HMC3\HA_EL3_T1.wiff  --f D:\Soumyadeep\Cell Click\Data\Cell Run 1 MS Data\Positive Click\HMC3\HA_EL3_T2.wiff  --f D:\Soumyadeep\Cell Click\Data\Cell Run 1 MS Data\Positive Click\HMC3\HA_EL3_T3.wiff  --f D:\Soumyadeep\Cell Click\Data\Cell Run 1 MS Data\Positive Click\HMC3\HA_EL3_T4.wiff  --f D:\Soumyadeep\Cell Click\Data\Cell Run 1 MS Data\Positive Click\Inhibited\HiA_EL3_T1.wiff  --f D:\Soumyadeep\Cell Click\Data\Cell Run 1 MS Data\Positive Click\Inhibited\HiA_EL3_T2.wiff  --f D:\Soumyadeep\Cell Click\Data\Cell Run 1 MS Data\Positive Click\Inhibited\HiA_EL3_T3.wiff  --f D:\Soumyadeep\Cell Click\Data\Cell Run 1 MS Data\Positive Click\Inhibited\HiA_EL3_T4.wiff  --f D:\Soumyadeep\Cell Click\Data\Cell Run 1 MS Data\Positive Click\Activated\HaA_EL3_T1.wiff  --f D:\Soumyadeep\Cell Click\Data\Cell Run 1 MS Data\Positive Click\Activated\HaA_EL3_T2.wiff  --f D:\Soumyadeep\Cell Click\Data\Cell Run 1 MS Data\Positive Click\Activated\HaA_EL3_T3.wiff  --f D:\Soumyadeep\Cell Click\Data\Cell Run 1 MS Data\Positive Click\Activated\HaA_EL3_T4.wiff  --f D:\Soumyadeep\Cell Click\Data\Cell Run 1 MS Data\No Azide Control\HMC3\HO_EL3_T1.wiff  --f D:\Soumyadeep\Cell Click\Data\Cell Run 1 MS Data\No Azide Control\HMC3\HO_EL3_T2.wiff  --f D:\Soumyadeep\Cell Click\Data\Cell Run 1 MS Data\No Azide Control\HMC3\HO_EL3_T3.wiff  --f D:\Soumyadeep\Cell Click\Data\Cell Run 1 MS Data\No Azide Control\HMC3\HO_EL3_T4.wiff  --f D:\Soumyadeep\Cell Click\Data\Cell Run 1 MS Data\No Azide Control\Inhibited\HiO_EL3_T1.wiff  --f D:\Soumyadeep\Cell Click\Data\Cell Run 1 MS Data\No Azide Control\Inhibited\HiO_EL3_T2.wiff  --f D:\Soumyadeep\Cell Click\Data\Cell Run 1 MS Data\No Azide Control\Inhibited\HiO_EL3_T3.wiff  --f D:\Soumyadeep\Cell Click\Data\Cell Run 1 MS Data\No Azide Control\Inhibited\HiO_EL3_T4.wiff  --f D:\Soumyadeep\Cell Click\Data\Cell Run 1 MS Data\No Azide Control\Activated\HaO_EL3_T1.wiff  --f D:\Soumyadeep\Cell Click\Data\Cell Run 1 MS Data\No Azide Control\Activated\HaO_EL3_T2.wiff  --f D:\Soumyadeep\Cell Click\Data\Cell Run 1 MS Data\No Azide Control\Activated\HaO_EL3_T3.wiff  --f D:\Soumyadeep\Cell Click\Data\Cell Run 1 MS Data\No Azide Control\Activated\HaO_EL3_T4.wiff  --lib D:\Soumyadeep\Cell Click\lib\click_canfinal_out.predicted.speclib --threads 18 --verbose 1 --out D:\Soumyadeep\Cell Click\out\click_canfinal_out.parquet --qvalue 0.05 --matrices --temp D:\Soumyadeep\Cell Click\temp --xic --fasta camprotR_240512_cRAP_20190401_full_tags.fasta --cont-quant-exclude cRAP- --fasta D:\Soumyadeep\Cell Click\clean_humwholeprot_reviewed_can.fasta --met-excision --cut K*,R*,!*P --missed-cleavages 2 --unimod4 --var-mods 3 --var-mod UniMod:35,15.994915,M --reanalyse --rt-profiling 

Thread number set to 18
Output will be filtered at 0.05 FDR
Precursor/protein x samples expression level matrices will be saved along with the main report
XICs within 10 seconds from the apex will be extracted for each precursor and saved in .parquet format, a folder will be created next to the main report for the XICs storage
Peptides corresponding to protein sequence IDs tagged with cRAP- will be excluded from normalisation as well as quantification of protein groups that do not include proteins bearing the tag
N-terminal methionine excision enabled
In silico digest will involve cuts at K*,R*
But excluding cuts at *P
Maximum number of missed cleavages set to 2
Cysteine carbamidomethylation enabled as a fixed modification
Maximum number of variable modifications set to 3
Modification UniMod:35 with mass delta 15.9949 at M will be considered as variable
MBR enabled; .quant files will only be saved to disk during the first pass
The spectral library (if generated) will retain the original spectra but will include empirically-aligned RTs
DIA-NN will automatically optimise the mass accuracy for the first run of the experiment, use this mode for preliminary analyses only
WARNING: peptidoform scoring enabled because variable modifications have been declared; to disable, use --no-peptidoforms
The following variable modifications will be localised: UniMod:35 
WARNING: protein inference will only be performed for precursors identified with global q-value <= 0.05 or q-value <= 0.05.

24 files will be processed
[0:00] Loading spectral library D:\Soumyadeep\Cell Click\lib\click_canfinal_out.predicted.speclib
[0:05] Library annotated with sequence database(s): D:\Soumyadeep\Cell Click\clean_humwholeprot_reviewed_can.fasta
[0:06] Spectral library loaded: 20368 protein isoforms, 29405 protein groups and 7700783 precursors in 2804689 elution groups.
[0:06] Loading protein annotations from FASTA camprotR_240512_cRAP_20190401_full_tags.fasta
[0:06] Loading protein annotations from FASTA D:\Soumyadeep\Cell Click\clean_humwholeprot_reviewed_can.fasta
[0:07] Annotating library proteins with information from the FASTA database
[0:07] Gene names missing for some isoforms
[0:07] Library contains 20368 proteins, and 20174 genes
[0:11] Initialising library

First pass: generating a spectral library from DIA data

[0:26] File #1/24
[0:26] Loading run D:\Soumyadeep\Cell Click\Data\Cell Run 1 MS Data\Positive Click\HMC3\HA_EL3_T1.wiff
[3:09] Pre-processing...
[3:11] 1117 MS1 and 111700 MS2 scans in 1117 (inferred) and 1117 (encoded) cycles, 6606085 precursors in range
[3:12] Calibrating with mass accuracies 25 (MS1), 25 (MS2)
[7:04] RT window set to 2.17164
[7:04] Peak width: 2.832
[7:04] Scan window radius set to 6
[7:04] Recommended MS1 mass accuracy setting: 20 ppm
[14:06] Optimised mass accuracy: 17 ppm
[15:54] Searching decoys
[17:54] Main search
[21:47] Removing low confidence identifications
[21:51] Removing interfering precursors
[21:54] Training neural networks on 15508 target and 7949 decoy PSMs
[22:06] Training neural networks on 15508 target and 7420 decoy PSMs
[22:15] Number of IDs at 0.01 FDR: 8721
[22:15] Precursors at 1% peptidoform FDR: 7871
[22:15] Calculating protein q-values
[22:16] Number of genes identified at 1% FDR: 1836 (precursor-level), 1633 (protein-level) (inference performed using proteotypic peptides only)
[22:16] Quantification
[22:16] Precursors with scored PTMs at 1% FDR: 144 out of 179 considered
[22:16] Precursors with all scored PTM sites unoccupied at 1% FDR: 7727
[22:16] Precursors with PTMs localised (when required) with > 90% confidence: 142 out of 144
[22:17] Quantification information saved to D:\Soumyadeep\Cell Click\temp/D__Soumyadeep_Cell Click_Data_Cell Run 1 MS Data_Positive Click_HMC3_HA_EL3_T1_wiff.quant

[22:17] File #2/24
[22:17] Loading run D:\Soumyadeep\Cell Click\Data\Cell Run 1 MS Data\Positive Click\HMC3\HA_EL3_T2.wiff
[25:27] Pre-processing...
[25:29] 1117 MS1 and 111700 MS2 scans in 1117 (inferred) and 1117 (encoded) cycles, 6606085 precursors in range
[25:30] Calibrating with mass accuracies 25 (MS1), 30 (MS2)
[29:37] RT window set to 2.32906
[29:37] Recommended MS1 mass accuracy setting: 21 ppm
[32:02] Searching decoys
[34:43] Main search
[39:59] Removing low confidence identifications
[40:04] Removing interfering precursors
[40:07] Training neural networks on 16403 target and 8597 decoy PSMs
[40:19] Training neural networks on 16403 target and 7880 decoy PSMs
[40:28] Number of IDs at 0.01 FDR: 9423
[40:28] Precursors at 1% peptidoform FDR: 8575
[40:29] Calculating protein q-values
[40:29] Number of genes identified at 1% FDR: 1908 (precursor-level), 1716 (protein-level) (inference performed using proteotypic peptides only)
[40:29] Quantification
[40:29] Precursors with scored PTMs at 1% FDR: 185 out of 233 considered
[40:29] Precursors with all scored PTM sites unoccupied at 1% FDR: 8390
[40:29] Precursors with PTMs localised (when required) with > 90% confidence: 181 out of 185
[40:30] Quantification information saved to D:\Soumyadeep\Cell Click\temp/D__Soumyadeep_Cell Click_Data_Cell Run 1 MS Data_Positive Click_HMC3_HA_EL3_T2_wiff.quant

[40:30] File #3/24
[40:30] Loading run D:\Soumyadeep\Cell Click\Data\Cell Run 1 MS Data\Positive Click\HMC3\HA_EL3_T3.wiff
[43:11] Pre-processing...
[43:12] 1117 MS1 and 111700 MS2 scans in 1117 (inferred) and 1117 (encoded) cycles, 6606085 precursors in range
[43:13] Calibrating with mass accuracies 25 (MS1), 30 (MS2)
[47:05] RT window set to 2.18571
[47:05] Recommended MS1 mass accuracy setting: 20 ppm
[48:58] Searching decoys
[50:50] Main search
[54:29] Removing low confidence identifications
[54:33] Removing interfering precursors
[54:36] Training neural networks on 14603 target and 7180 decoy PSMs
[54:47] Training neural networks on 14603 target and 6518 decoy PSMs
[54:56] Number of IDs at 0.01 FDR: 8705
[54:56] Precursors at 1% peptidoform FDR: 8017
[54:57] Calculating protein q-values
[54:57] Number of genes identified at 1% FDR: 1910 (precursor-level), 1627 (protein-level) (inference performed using proteotypic peptides only)
[54:57] Quantification
[54:57] Precursors with scored PTMs at 1% FDR: 137 out of 172 considered
[54:57] Precursors with all scored PTM sites unoccupied at 1% FDR: 7880
[54:57] Precursors with PTMs localised (when required) with > 90% confidence: 134 out of 137
[54:58] Quantification information saved to D:\Soumyadeep\Cell Click\temp/D__Soumyadeep_Cell Click_Data_Cell Run 1 MS Data_Positive Click_HMC3_HA_EL3_T3_wiff.quant

[54:58] File #4/24
[54:58] Loading run D:\Soumyadeep\Cell Click\Data\Cell Run 1 MS Data\Positive Click\HMC3\HA_EL3_T4.wiff
[56:18] Pre-processing...
[56:19] 1117 MS1 and 111700 MS2 scans in 1117 (inferred) and 1117 (encoded) cycles, 6606085 precursors in range
[56:20] Calibrating with mass accuracies 25 (MS1), 25 (MS2)
[88:52] Cannot perform mass calibration, too few confidently identified precursors
[88:52] Cannot perform MS1 mass calibration, too few confidently identified precursors
[88:53] Searching decoys
[94:47] Main search
[106:21] Removing low confidence identifications
[106:22] Cannot perform MS1 mass calibration, too few confidently identified precursors
[106:24] Removing interfering precursors
[106:27] Training neural networks on 3091 target and 3088 decoy PSMs
[106:34] Training neural networks on 3091 target and 3007 decoy PSMs
[106:39] Number of IDs at 0.01 FDR: 4
[106:39] Disabling neural networks
[106:40] Number of IDs at 0.01 FDR: 0
[106:40] Precursors at 1% peptidoform FDR: 0
[106:40] Calculating protein q-values
[106:40] Number of genes identified at 1% FDR: 0 (precursor-level), 0 (protein-level) (inference performed using proteotypic peptides only)
[106:41] Quantification
[106:41] Precursors with scored PTMs at 1% FDR: 0 out of 0 considered
[106:41] Precursors with all scored PTM sites unoccupied at 1% FDR: 0
[106:41] Precursors with PTMs localised (when required) with > 90% confidence: 0 out of 0
[106:41] Quantification information saved to D:\Soumyadeep\Cell Click\temp/D__Soumyadeep_Cell Click_Data_Cell Run 1 MS Data_Positive Click_HMC3_HA_EL3_T4_wiff.quant

[106:41] File #5/24
[106:41] Loading run D:\Soumyadeep\Cell Click\Data\Cell Run 1 MS Data\Positive Click\Inhibited\HiA_EL3_T1.wiff
[108:04] Pre-processing...
[108:06] 1117 MS1 and 111700 MS2 scans in 1117 (inferred) and 1117 (encoded) cycles, 6606085 precursors in range
[108:06] Calibrating with mass accuracies 25 (MS1), 25 (MS2)
[117:57] RT window set to 3.68297
[117:57] Recommended MS1 mass accuracy setting: 15 ppm
[121:51] Searching decoys
[122:46] Main search
[124:34] Removing low confidence identifications
[124:38] Removing interfering precursors
[124:40] Training neural networks on 1502 target and 1078 decoy PSMs
[124:49] Training neural networks on 1502 target and 981 decoy PSMs
[124:55] Number of IDs at 0.01 FDR: 384
[124:55] Precursors at 1% peptidoform FDR: 0
[124:56] Calculating protein q-values
[124:56] Number of genes identified at 1% FDR: 122 (precursor-level), 120 (protein-level) (inference performed using proteotypic peptides only)
[124:56] Quantification
[124:56] Precursors with scored PTMs at 1% FDR: 0 out of 5 considered
[124:56] Precursors with all scored PTM sites unoccupied at 1% FDR: 0
[124:56] Precursors with PTMs localised (when required) with > 90% confidence: 0 out of 0
[124:57] Quantification information saved to D:\Soumyadeep\Cell Click\temp/D__Soumyadeep_Cell Click_Data_Cell Run 1 MS Data_Positive Click_Inhibited_HiA_EL3_T1_wiff.quant

[124:57] File #6/24
[124:57] Loading run D:\Soumyadeep\Cell Click\Data\Cell Run 1 MS Data\Positive Click\Inhibited\HiA_EL3_T2.wiff
[126:15] Pre-processing...
[126:17] 1117 MS1 and 111700 MS2 scans in 1117 (inferred) and 1117 (encoded) cycles, 6606085 precursors in range
[126:17] Calibrating with mass accuracies 25 (MS1), 26 (MS2)
[136:14] RT window set to 3.07922
[136:14] Recommended MS1 mass accuracy setting: 16 ppm
[140:06] Searching decoys
[140:56] Main search
[142:34] Removing low confidence identifications
[142:37] Removing interfering precursors
[142:40] Training neural networks on 1567 target and 1004 decoy PSMs
[142:48] Training neural networks on 1567 target and 895 decoy PSMs
[142:53] Number of IDs at 0.01 FDR: 572
[142:54] Precursors at 1% peptidoform FDR: 436
[142:54] Calculating protein q-values
[142:54] Number of genes identified at 1% FDR: 178 (precursor-level), 157 (protein-level) (inference performed using proteotypic peptides only)
[142:54] Quantification
[142:55] Precursors with scored PTMs at 1% FDR: 3 out of 5 considered
[142:55] Precursors with all scored PTM sites unoccupied at 1% FDR: 433
[142:55] Precursors with PTMs localised (when required) with > 90% confidence: 3 out of 3
[142:55] Quantification information saved to D:\Soumyadeep\Cell Click\temp/D__Soumyadeep_Cell Click_Data_Cell Run 1 MS Data_Positive Click_Inhibited_HiA_EL3_T2_wiff.quant

[142:55] File #7/24
[142:55] Loading run D:\Soumyadeep\Cell Click\Data\Cell Run 1 MS Data\Positive Click\Inhibited\HiA_EL3_T3.wiff
[144:23] Pre-processing...
[144:25] 1117 MS1 and 111700 MS2 scans in 1117 (inferred) and 1117 (encoded) cycles, 6606085 precursors in range
[144:25] Calibrating with mass accuracies 25 (MS1), 40 (MS2)
[150:48] Recommended MS1 mass accuracy setting: 18 ppm
[163:24] RT window set to 5.22887
[163:25] Recommended MS1 mass accuracy setting: 18 ppm
[169:17] Searching decoys
[170:43] Main search
[173:32] Removing low confidence identifications
[173:35] Removing interfering precursors
[173:38] Training neural networks on 1302 target and 883 decoy PSMs
[173:45] Training neural networks on 1302 target and 776 decoy PSMs
[173:49] Number of IDs at 0.01 FDR: 460
[173:50] Precursors at 1% peptidoform FDR: 206
[173:50] Calculating protein q-values
[173:50] Number of genes identified at 1% FDR: 153 (precursor-level), 0 (protein-level) (inference performed using proteotypic peptides only)
[173:51] Quantification
[173:51] Precursors with scored PTMs at 1% FDR: 0 out of 5 considered
[173:51] Precursors with all scored PTM sites unoccupied at 1% FDR: 206
[173:51] Precursors with PTMs localised (when required) with > 90% confidence: 0 out of 0
[173:51] Quantification information saved to D:\Soumyadeep\Cell Click\temp/D__Soumyadeep_Cell Click_Data_Cell Run 1 MS Data_Positive Click_Inhibited_HiA_EL3_T3_wiff.quant

[173:51] File #8/24
[173:51] Loading run D:\Soumyadeep\Cell Click\Data\Cell Run 1 MS Data\Positive Click\Inhibited\HiA_EL3_T4.wiff
[175:19] Pre-processing...
[175:21] 1117 MS1 and 111700 MS2 scans in 1117 (inferred) and 1117 (encoded) cycles, 6606085 precursors in range
[175:21] Calibrating with mass accuracies 25 (MS1), 25 (MS2)
[185:30] RT window set to 4.27784
[185:31] Recommended MS1 mass accuracy setting: 14 ppm
[189:47] Searching decoys
[190:54] Main search
[193:06] Removing low confidence identifications
[193:10] Removing interfering precursors
[193:13] Training neural networks on 1132 target and 594 decoy PSMs
[193:19] Training neural networks on 1132 target and 538 decoy PSMs
[193:23] Number of IDs at 0.01 FDR: 473
[193:23] Precursors at 1% peptidoform FDR: 138
[193:23] Calculating protein q-values
[193:24] Number of genes identified at 1% FDR: 161 (precursor-level), 109 (protein-level) (inference performed using proteotypic peptides only)
[193:24] Quantification
[193:24] Precursors with scored PTMs at 1% FDR: 0 out of 3 considered
[193:24] Precursors with all scored PTM sites unoccupied at 1% FDR: 138
[193:24] Precursors with PTMs localised (when required) with > 90% confidence: 0 out of 0
[193:24] Quantification information saved to D:\Soumyadeep\Cell Click\temp/D__Soumyadeep_Cell Click_Data_Cell Run 1 MS Data_Positive Click_Inhibited_HiA_EL3_T4_wiff.quant

[193:24] File #9/24
[193:24] Loading run D:\Soumyadeep\Cell Click\Data\Cell Run 1 MS Data\Positive Click\Activated\HaA_EL3_T1.wiff
[195:12] Pre-processing...
[195:13] 1117 MS1 and 111700 MS2 scans in 1117 (inferred) and 1117 (encoded) cycles, 6606085 precursors in range
[195:14] Calibrating with mass accuracies 25 (MS1), 30 (MS2)
[202:12] RT window set to 4.82624
[202:13] Recommended MS1 mass accuracy setting: 16 ppm
[205:39] Searching decoys
[207:38] Main search
[211:27] Removing low confidence identifications
[211:31] Removing interfering precursors
[211:34] Training neural networks on 4474 target and 2419 decoy PSMs
[211:41] Training neural networks on 4474 target and 2170 decoy PSMs
[211:47] Number of IDs at 0.01 FDR: 2624
[211:47] Precursors at 1% peptidoform FDR: 2345
[211:48] Calculating protein q-values
[211:48] Number of genes identified at 1% FDR: 783 (precursor-level), 741 (protein-level) (inference performed using proteotypic peptides only)
[211:48] Quantification
[211:48] Precursors with scored PTMs at 1% FDR: 29 out of 43 considered
[211:48] Precursors with all scored PTM sites unoccupied at 1% FDR: 2316
[211:48] Precursors with PTMs localised (when required) with > 90% confidence: 29 out of 29
[211:49] Quantification information saved to D:\Soumyadeep\Cell Click\temp/D__Soumyadeep_Cell Click_Data_Cell Run 1 MS Data_Positive Click_Activated_HaA_EL3_T1_wiff.quant

[211:49] File #10/24
[211:49] Loading run D:\Soumyadeep\Cell Click\Data\Cell Run 1 MS Data\Positive Click\Activated\HaA_EL3_T2.wiff
[213:33] Pre-processing...
[213:34] 1117 MS1 and 111700 MS2 scans in 1117 (inferred) and 1117 (encoded) cycles, 6606085 precursors in range
[213:35] Calibrating with mass accuracies 25 (MS1), 29 (MS2)
[220:30] RT window set to 3.45057
[220:31] Recommended MS1 mass accuracy setting: 19 ppm
[225:06] Searching decoys
[227:33] Main search
[232:19] Removing low confidence identifications
[232:23] Removing interfering precursors
[232:26] Training neural networks on 5173 target and 2592 decoy PSMs
[232:34] Training neural networks on 5173 target and 2341 decoy PSMs
[232:39] Number of IDs at 0.01 FDR: 3085
[232:39] Precursors at 1% peptidoform FDR: 2807
[232:40] Calculating protein q-values
[232:40] Number of genes identified at 1% FDR: 894 (precursor-level), 760 (protein-level) (inference performed using proteotypic peptides only)
[232:40] Quantification
[232:40] Precursors with scored PTMs at 1% FDR: 35 out of 46 considered
[232:40] Precursors with all scored PTM sites unoccupied at 1% FDR: 2772
[232:40] Precursors with PTMs localised (when required) with > 90% confidence: 32 out of 35
[232:41] Quantification information saved to D:\Soumyadeep\Cell Click\temp/D__Soumyadeep_Cell Click_Data_Cell Run 1 MS Data_Positive Click_Activated_HaA_EL3_T2_wiff.quant

[232:41] File #11/24
[232:41] Loading run D:\Soumyadeep\Cell Click\Data\Cell Run 1 MS Data\Positive Click\Activated\HaA_EL3_T3.wiff
[234:26] Pre-processing...
[234:27] 1117 MS1 and 111700 MS2 scans in 1117 (inferred) and 1117 (encoded) cycles, 6606085 precursors in range
[234:28] Calibrating with mass accuracies 25 (MS1), 30 (MS2)
[241:20] RT window set to 3.49733
[241:21] Recommended MS1 mass accuracy setting: 16 ppm
[245:06] Searching decoys
[247:33] Main search
[252:19] Removing low confidence identifications
[252:23] Removing interfering precursors
[252:26] Training neural networks on 5033 target and 2285 decoy PSMs
[252:34] Training neural networks on 5033 target and 2033 decoy PSMs
[252:39] Number of IDs at 0.01 FDR: 3243
[252:39] Precursors at 1% peptidoform FDR: 2979
[252:39] Calculating protein q-values
[252:40] Number of genes identified at 1% FDR: 926 (precursor-level), 872 (protein-level) (inference performed using proteotypic peptides only)
[252:40] Quantification
[252:40] Precursors with scored PTMs at 1% FDR: 34 out of 47 considered
[252:40] Precursors with all scored PTM sites unoccupied at 1% FDR: 2945
[252:40] Precursors with PTMs localised (when required) with > 90% confidence: 33 out of 34
[252:41] Quantification information saved to D:\Soumyadeep\Cell Click\temp/D__Soumyadeep_Cell Click_Data_Cell Run 1 MS Data_Positive Click_Activated_HaA_EL3_T3_wiff.quant

[252:41] File #12/24
[252:41] Loading run D:\Soumyadeep\Cell Click\Data\Cell Run 1 MS Data\Positive Click\Activated\HaA_EL3_T4.wiff
[254:38] Pre-processing...
[254:40] 1117 MS1 and 111700 MS2 scans in 1117 (inferred) and 1117 (encoded) cycles, 6606085 precursors in range
[254:40] Calibrating with mass accuracies 25 (MS1), 25 (MS2)
[259:32] RT window set to 3.19512
[259:33] Recommended MS1 mass accuracy setting: 21 ppm
[263:24] Searching decoys
[265:41] Main search
[270:06] Removing low confidence identifications
[270:10] Removing interfering precursors
[270:13] Training neural networks on 5777 target and 2781 decoy PSMs
[270:21] Training neural networks on 5777 target and 2543 decoy PSMs
[270:26] Number of IDs at 0.01 FDR: 3570
[270:27] Precursors at 1% peptidoform FDR: 3129
[270:27] Calculating protein q-values
[270:27] Number of genes identified at 1% FDR: 1006 (precursor-level), 930 (protein-level) (inference performed using proteotypic peptides only)
[270:27] Quantification
[270:28] Precursors with scored PTMs at 1% FDR: 32 out of 51 considered
[270:28] Precursors with all scored PTM sites unoccupied at 1% FDR: 3097
[270:28] Precursors with PTMs localised (when required) with > 90% confidence: 31 out of 32
[270:28] Quantification information saved to D:\Soumyadeep\Cell Click\temp/D__Soumyadeep_Cell Click_Data_Cell Run 1 MS Data_Positive Click_Activated_HaA_EL3_T4_wiff.quant

[270:28] File #13/24
[270:28] Loading run D:\Soumyadeep\Cell Click\Data\Cell Run 1 MS Data\No Azide Control\HMC3\HO_EL3_T1.wiff
[276:07] Pre-processing...
[276:09] 1117 MS1 and 111700 MS2 scans in 1117 (inferred) and 1117 (encoded) cycles, 6606085 precursors in range
[276:09] Calibrating with mass accuracies 25 (MS1), 40 (MS2)
[287:16] Recommended MS1 mass accuracy setting: 11 ppm
[299:01] RT window set to 6.4152
[299:02] Recommended MS1 mass accuracy setting: 11 ppm
[323:39] Searching decoys
[331:01] Main search
[345:32] Removing low confidence identifications
[345:35] Removing interfering precursors
[345:38] Training neural networks on 625 target and 578 decoy PSMs
[345:44] Training neural networks on 625 target and 524 decoy PSMs
[345:46] Number of IDs at 0.01 FDR: 27
[345:46] Disabling neural networks
[345:47] Number of IDs at 0.01 FDR: 0
[345:47] Precursors at 1% peptidoform FDR: 0
[345:47] Calculating protein q-values
[345:48] Number of genes identified at 1% FDR: 0 (precursor-level), 0 (protein-level) (inference performed using proteotypic peptides only)
[345:48] Quantification
[345:48] Precursors with scored PTMs at 1% FDR: 0 out of 0 considered
[345:48] Precursors with all scored PTM sites unoccupied at 1% FDR: 0
[345:48] Precursors with PTMs localised (when required) with > 90% confidence: 0 out of 0
[345:48] Quantification information saved to D:\Soumyadeep\Cell Click\temp/D__Soumyadeep_Cell Click_Data_Cell Run 1 MS Data_No Azide Control_HMC3_HO_EL3_T1_wiff.quant

[345:48] File #14/24
[345:48] Loading run D:\Soumyadeep\Cell Click\Data\Cell Run 1 MS Data\No Azide Control\HMC3\HO_EL3_T2.wiff
[351:41] Pre-processing...
[351:43] 1117 MS1 and 111700 MS2 scans in 1117 (inferred) and 1117 (encoded) cycles, 6606085 precursors in range
[351:43] Calibrating with mass accuracies 25 (MS1), 40 (MS2)
[362:37] Recommended MS1 mass accuracy setting: 8 ppm
[375:44] RT window set to 5.71474
[375:45] Recommended MS1 mass accuracy setting: 7 ppm
[402:49] Searching decoys
[410:31] Main search
[425:32] Removing low confidence identifications
[425:35] Removing interfering precursors
[425:38] Training neural networks on 641 target and 589 decoy PSMs
[425:44] Training neural networks on 641 target and 581 decoy PSMs
[425:47] Number of IDs at 0.01 FDR: 16
[425:47] Disabling neural networks
[425:47] Number of IDs at 0.01 FDR: 0
[425:48] Precursors at 1% peptidoform FDR: 0
[425:48] Calculating protein q-values
[425:48] Number of genes identified at 1% FDR: 0 (precursor-level), 0 (protein-level) (inference performed using proteotypic peptides only)
[425:48] Quantification
[425:49] Precursors with scored PTMs at 1% FDR: 0 out of 0 considered
[425:49] Precursors with all scored PTM sites unoccupied at 1% FDR: 0
[425:49] Precursors with PTMs localised (when required) with > 90% confidence: 0 out of 0
[425:49] Quantification information saved to D:\Soumyadeep\Cell Click\temp/D__Soumyadeep_Cell Click_Data_Cell Run 1 MS Data_No Azide Control_HMC3_HO_EL3_T2_wiff.quant

[425:49] File #15/24
[425:49] Loading run D:\Soumyadeep\Cell Click\Data\Cell Run 1 MS Data\No Azide Control\HMC3\HO_EL3_T3.wiff
[431:42] Pre-processing...
[431:43] 1117 MS1 and 111700 MS2 scans in 1117 (inferred) and 1117 (encoded) cycles, 6606085 precursors in range
[431:44] Calibrating with mass accuracies 25 (MS1), 40 (MS2)
[445:00] Recommended MS1 mass accuracy setting: 9 ppm
[460:05] RT window set to 6.47459
[460:07] Recommended MS1 mass accuracy setting: 8 ppm
[491:08] Searching decoys
[499:49] Main search
[516:48] Removing low confidence identifications
[516:54] Removing interfering precursors
[516:57] Training neural networks on 547 target and 511 decoy PSMs
[517:04] Too few confident identifications, neural networks will not be used
[517:04] Number of IDs at 0.01 FDR: 0
[517:05] Precursors at 1% peptidoform FDR: 0
[517:05] Calculating protein q-values
[517:06] Number of genes identified at 1% FDR: 0 (precursor-level), 0 (protein-level) (inference performed using proteotypic peptides only)
[517:06] Quantification
[517:06] Precursors with scored PTMs at 1% FDR: 0 out of 0 considered
[517:06] Precursors with all scored PTM sites unoccupied at 1% FDR: 0
[517:06] Precursors with PTMs localised (when required) with > 90% confidence: 0 out of 0
[517:07] Quantification information saved to D:\Soumyadeep\Cell Click\temp/D__Soumyadeep_Cell Click_Data_Cell Run 1 MS Data_No Azide Control_HMC3_HO_EL3_T3_wiff.quant

[517:07] File #16/24
[517:07] Loading run D:\Soumyadeep\Cell Click\Data\Cell Run 1 MS Data\No Azide Control\HMC3\HO_EL3_T4.wiff
[522:36] Pre-processing...
[522:38] 1117 MS1 and 111700 MS2 scans in 1117 (inferred) and 1117 (encoded) cycles, 6606085 precursors in range
[522:38] Calibrating with mass accuracies 25 (MS1), 40 (MS2)
[567:48] Searching decoys
[573:28] Main search
[584:36] Removing low confidence identifications
[584:40] Removing interfering precursors
[584:45] Training neural networks on 8543 target and 8633 decoy PSMs
[584:55] Training neural networks on 8543 target and 8428 decoy PSMs
[585:03] Number of IDs at 0.01 FDR: 0
[585:03] Disabling neural networks
[585:03] Number of IDs at 0.01 FDR: 0
[585:04] Precursors at 1% peptidoform FDR: 0
[585:04] Calculating protein q-values
[585:04] Number of genes identified at 1% FDR: 0 (precursor-level), 0 (protein-level) (inference performed using proteotypic peptides only)
[585:04] Quantification
[585:05] Precursors with scored PTMs at 1% FDR: 0 out of 0 considered
[585:05] Precursors with all scored PTM sites unoccupied at 1% FDR: 0
[585:05] Precursors with PTMs localised (when required) with > 90% confidence: 0 out of 0
[585:05] Quantification information saved to D:\Soumyadeep\Cell Click\temp/D__Soumyadeep_Cell Click_Data_Cell Run 1 MS Data_No Azide Control_HMC3_HO_EL3_T4_wiff.quant

[585:05] File #17/24
[585:05] Loading run D:\Soumyadeep\Cell Click\Data\Cell Run 1 MS Data\No Azide Control\Inhibited\HiO_EL3_T1.wiff
[590:31] Pre-processing...
[590:33] 1117 MS1 and 111700 MS2 scans in 1117 (inferred) and 1117 (encoded) cycles, 6606085 precursors in range
[590:33] Calibrating with mass accuracies 25 (MS1), 36 (MS2)
[601:06] Recommended MS1 mass accuracy setting: 17 ppm
[613:53] RT window set to 5.72866
[613:54] Recommended MS1 mass accuracy setting: 15 ppm
[641:07] Searching decoys
[647:40] Main search
[660:32] Removing low confidence identifications
[660:35] Removing interfering precursors
[660:38] Training neural networks on 801 target and 793 decoy PSMs
[660:44] Training neural networks on 801 target and 752 decoy PSMs
[660:48] Number of IDs at 0.01 FDR: 12
[660:48] Disabling neural networks
[660:48] Number of IDs at 0.01 FDR: 0
[660:49] Precursors at 1% peptidoform FDR: 0
[660:49] Calculating protein q-values
[660:49] Number of genes identified at 1% FDR: 0 (precursor-level), 0 (protein-level) (inference performed using proteotypic peptides only)
[660:49] Quantification
[660:49] Precursors with scored PTMs at 1% FDR: 0 out of 0 considered
[660:49] Precursors with all scored PTM sites unoccupied at 1% FDR: 0
[660:49] Precursors with PTMs localised (when required) with > 90% confidence: 0 out of 0
[660:50] Quantification information saved to D:\Soumyadeep\Cell Click\temp/D__Soumyadeep_Cell Click_Data_Cell Run 1 MS Data_No Azide Control_Inhibited_HiO_EL3_T1_wiff.quant

[660:50] File #18/24
[660:50] Loading run D:\Soumyadeep\Cell Click\Data\Cell Run 1 MS Data\No Azide Control\Inhibited\HiO_EL3_T2.wiff
[666:31] Pre-processing...
[666:32] 1117 MS1 and 111700 MS2 scans in 1117 (inferred) and 1117 (encoded) cycles, 6606085 precursors in range
[666:33] Calibrating with mass accuracies 25 (MS1), 25 (MS2)
[676:57] RT window set to 4.79103
[676:58] Recommended MS1 mass accuracy setting: 11 ppm
[704:02] Searching decoys
[710:50] Main search
[724:14] Removing low confidence identifications
[724:18] Removing interfering precursors
[724:21] Too few confident identifications, neural networks will not be used
[724:24] Too few confident identifications, neural networks will not be used
[724:24] Number of IDs at 0.01 FDR: 0
[724:24] Precursors at 1% peptidoform FDR: 0
[724:25] Calculating protein q-values
[724:25] Number of genes identified at 1% FDR: 0 (precursor-level), 0 (protein-level) (inference performed using proteotypic peptides only)
[724:25] Quantification
[724:25] Precursors with scored PTMs at 1% FDR: 0 out of 0 considered
[724:25] Precursors with all scored PTM sites unoccupied at 1% FDR: 0
[724:25] Precursors with PTMs localised (when required) with > 90% confidence: 0 out of 0
[724:26] Quantification information saved to D:\Soumyadeep\Cell Click\temp/D__Soumyadeep_Cell Click_Data_Cell Run 1 MS Data_No Azide Control_Inhibited_HiO_EL3_T2_wiff.quant

[724:26] File #19/24
[724:26] Loading run D:\Soumyadeep\Cell Click\Data\Cell Run 1 MS Data\No Azide Control\Inhibited\HiO_EL3_T3.wiff
[730:03] Pre-processing...
[730:04] 1117 MS1 and 111700 MS2 scans in 1117 (inferred) and 1117 (encoded) cycles, 6606085 precursors in range
[730:05] Calibrating with mass accuracies 25 (MS1), 25 (MS2)
[740:07] RT window set to 6.67305
[740:07] Recommended MS1 mass accuracy setting: 10 ppm
[767:02] Searching decoys
[774:16] Main search
[788:30] Removing low confidence identifications
[788:34] Removing interfering precursors
[788:36] Training neural networks on 586 target and 554 decoy PSMs
[788:42] Training neural networks on 586 target and 541 decoy PSMs
[788:44] Number of IDs at 0.01 FDR: 23
[788:44] Disabling neural networks
[788:45] Number of IDs at 0.01 FDR: 0
[788:45] Precursors at 1% peptidoform FDR: 0
[788:45] Calculating protein q-values
[788:46] Number of genes identified at 1% FDR: 0 (precursor-level), 0 (protein-level) (inference performed using proteotypic peptides only)
[788:46] Quantification
[788:46] Precursors with scored PTMs at 1% FDR: 0 out of 0 considered
[788:46] Precursors with all scored PTM sites unoccupied at 1% FDR: 0
[788:46] Precursors with PTMs localised (when required) with > 90% confidence: 0 out of 0
[788:46] Quantification information saved to D:\Soumyadeep\Cell Click\temp/D__Soumyadeep_Cell Click_Data_Cell Run 1 MS Data_No Azide Control_Inhibited_HiO_EL3_T3_wiff.quant

[788:47] File #20/24
[788:47] Loading run D:\Soumyadeep\Cell Click\Data\Cell Run 1 MS Data\No Azide Control\Inhibited\HiO_EL3_T4.wiff
[793:54] Pre-processing...
[793:56] 1117 MS1 and 111700 MS2 scans in 1117 (inferred) and 1117 (encoded) cycles, 6606085 precursors in range
[793:57] Calibrating with mass accuracies 25 (MS1), 39 (MS2)
[838:30] Searching decoys
[844:17] Main search
[855:37] Removing low confidence identifications
[855:42] Removing interfering precursors
[855:45] Training neural networks on 36360 target and 36148 decoy PSMs
[856:15] Training neural networks on 36360 target and 35510 decoy PSMs
[856:43] Number of IDs at 0.01 FDR: 0
[856:43] Disabling neural networks
[856:43] Number of IDs at 0.01 FDR: 0
[856:44] Precursors at 1% peptidoform FDR: 0
[856:44] Calculating protein q-values
[856:44] Number of genes identified at 1% FDR: 0 (precursor-level), 0 (protein-level) (inference performed using proteotypic peptides only)
[856:44] Quantification
[856:45] Precursors with scored PTMs at 1% FDR: 0 out of 0 considered
[856:45] Precursors with all scored PTM sites unoccupied at 1% FDR: 0
[856:45] Precursors with PTMs localised (when required) with > 90% confidence: 0 out of 0
[856:45] Quantification information saved to D:\Soumyadeep\Cell Click\temp/D__Soumyadeep_Cell Click_Data_Cell Run 1 MS Data_No Azide Control_Inhibited_HiO_EL3_T4_wiff.quant

[856:45] File #21/24
[856:45] Loading run D:\Soumyadeep\Cell Click\Data\Cell Run 1 MS Data\No Azide Control\Activated\HaO_EL3_T1.wiff
[862:21] Pre-processing...
[862:22] 1117 MS1 and 111700 MS2 scans in 1117 (inferred) and 1117 (encoded) cycles, 6606085 precursors in range
[862:23] Calibrating with mass accuracies 25 (MS1), 62 (MS2)
[873:47] Recommended MS1 mass accuracy setting: 7 ppm
[886:07] RT window set to 5.32379
[886:08] Recommended MS1 mass accuracy setting: 7 ppm
[911:45] Searching decoys
[918:08] Main search
[930:45] Removing low confidence identifications
[930:49] Removing interfering precursors
[930:51] Too few confident identifications, neural networks will not be used
[930:54] Too few confident identifications, neural networks will not be used
[930:55] Number of IDs at 0.01 FDR: 0
[930:55] Precursors at 1% peptidoform FDR: 0
[930:55] Calculating protein q-values
[930:56] Number of genes identified at 1% FDR: 0 (precursor-level), 0 (protein-level) (inference performed using proteotypic peptides only)
[930:56] Quantification
[930:56] Precursors with scored PTMs at 1% FDR: 0 out of 0 considered
[930:56] Precursors with all scored PTM sites unoccupied at 1% FDR: 0
[930:56] Precursors with PTMs localised (when required) with > 90% confidence: 0 out of 0
[930:56] Quantification information saved to D:\Soumyadeep\Cell Click\temp/D__Soumyadeep_Cell Click_Data_Cell Run 1 MS Data_No Azide Control_Activated_HaO_EL3_T1_wiff.quant

[930:57] File #22/24
[930:57] Loading run D:\Soumyadeep\Cell Click\Data\Cell Run 1 MS Data\No Azide Control\Activated\HaO_EL3_T2.wiff
[936:28] Pre-processing...
[936:30] 1117 MS1 and 111700 MS2 scans in 1117 (inferred) and 1117 (encoded) cycles, 6606085 precursors in range
[936:31] Calibrating with mass accuracies 25 (MS1), 33 (MS2)
[947:10] Recommended MS1 mass accuracy setting: 10 ppm
[959:43] RT window set to 7.9172
[959:44] Recommended MS1 mass accuracy setting: 12 ppm
[983:31] Searching decoys
[990:40] Main search
[1004:45] Removing low confidence identifications
[1004:49] Removing interfering precursors
[1004:51] Training neural networks on 1641 target and 1591 decoy PSMs
[1004:58] Training neural networks on 1641 target and 1473 decoy PSMs
[1005:02] Number of IDs at 0.01 FDR: 16
[1005:02] Disabling neural networks
[1005:03] Number of IDs at 0.01 FDR: 0
[1005:03] Precursors at 1% peptidoform FDR: 0
[1005:03] Calculating protein q-values
[1005:04] Number of genes identified at 1% FDR: 0 (precursor-level), 0 (protein-level) (inference performed using proteotypic peptides only)
[1005:04] Quantification
[1005:04] Precursors with scored PTMs at 1% FDR: 0 out of 0 considered
[1005:04] Precursors with all scored PTM sites unoccupied at 1% FDR: 0
[1005:04] Precursors with PTMs localised (when required) with > 90% confidence: 0 out of 0
[1005:04] Quantification information saved to D:\Soumyadeep\Cell Click\temp/D__Soumyadeep_Cell Click_Data_Cell Run 1 MS Data_No Azide Control_Activated_HaO_EL3_T2_wiff.quant

[1005:04] File #23/24
[1005:04] Loading run D:\Soumyadeep\Cell Click\Data\Cell Run 1 MS Data\No Azide Control\Activated\HaO_EL3_T3.wiff
[1010:43] Pre-processing...
[1010:45] 1117 MS1 and 111700 MS2 scans in 1117 (inferred) and 1117 (encoded) cycles, 6606085 precursors in range
[1010:45] Calibrating with mass accuracies 25 (MS1), 30 (MS2)
[1021:23] RT window set to 3.71034
[1021:23] Recommended MS1 mass accuracy setting: 10 ppm
[1048:44] Searching decoys
[1056:01] Main search
[1070:24] Removing low confidence identifications
[1070:28] Removing interfering precursors
[1070:31] Training neural networks on 540 target and 511 decoy PSMs
[1070:36] Too few confident identifications, neural networks will not be used
[1070:36] Number of IDs at 0.01 FDR: 0
[1070:36] Precursors at 1% peptidoform FDR: 0
[1070:37] Calculating protein q-values
[1070:37] Number of genes identified at 1% FDR: 0 (precursor-level), 0 (protein-level) (inference performed using proteotypic peptides only)
[1070:37] Quantification
[1070:37] Precursors with scored PTMs at 1% FDR: 0 out of 0 considered
[1070:37] Precursors with all scored PTM sites unoccupied at 1% FDR: 0
[1070:37] Precursors with PTMs localised (when required) with > 90% confidence: 0 out of 0
[1070:38] Quantification information saved to D:\Soumyadeep\Cell Click\temp/D__Soumyadeep_Cell Click_Data_Cell Run 1 MS Data_No Azide Control_Activated_HaO_EL3_T3_wiff.quant

[1070:38] File #24/24
[1070:38] Loading run D:\Soumyadeep\Cell Click\Data\Cell Run 1 MS Data\No Azide Control\Activated\HaO_EL3_T4.wiff
[1076:05] Pre-processing...
[1076:07] 1117 MS1 and 111700 MS2 scans in 1117 (inferred) and 1117 (encoded) cycles, 6606085 precursors in range
[1076:08] Calibrating with mass accuracies 25 (MS1), 34 (MS2)
[1086:36] Recommended MS1 mass accuracy setting: 12 ppm
[1099:42] RT window set to 6.91185
[1099:43] Recommended MS1 mass accuracy setting: 5 ppm
[1127:05] Searching decoys
[1133:36] Main search
[1146:24] Removing low confidence identifications
[1146:28] Removing interfering precursors
[1146:31] Training neural networks on 772 target and 760 decoy PSMs
[1146:37] Training neural networks on 772 target and 704 decoy PSMs
[1146:40] Number of IDs at 0.01 FDR: 18
[1146:40] Disabling neural networks
[1146:40] Number of IDs at 0.01 FDR: 0
[1146:41] Precursors at 1% peptidoform FDR: 0
[1146:41] Calculating protein q-values
[1146:41] Number of genes identified at 1% FDR: 0 (precursor-level), 0 (protein-level) (inference performed using proteotypic peptides only)
[1146:41] Quantification
[1146:42] Precursors with scored PTMs at 1% FDR: 0 out of 0 considered
[1146:42] Precursors with all scored PTM sites unoccupied at 1% FDR: 0
[1146:42] Precursors with PTMs localised (when required) with > 90% confidence: 0 out of 0
[1146:42] Quantification information saved to D:\Soumyadeep\Cell Click\temp/D__Soumyadeep_Cell Click_Data_Cell Run 1 MS Data_No Azide Control_Activated_HaO_EL3_T4_wiff.quant

[1146:42] Cross-run analysis
[1146:42] Reading quantification information: 24 files
[1146:57] Quantifying peptides
[1147:03] Assembling protein groups
[1147:04] Quantifying proteins
[1147:04] Calculating q-values for protein and gene groups
[1147:11] Calculating global q-values for protein and gene groups
[1147:11] Protein groups with global q-value <= 0.01: 2072
[1147:11] Compressed report saved to D:\Soumyadeep\Cell Click\out\click_canfinal_out-first-pass.parquet. Use R 'arrow' or Python 'PyArrow' package to process
[1147:11] Site report saved to D:\Soumyadeep\Cell Click\out\click_canfinal_out-first-pass.site_report.parquet
[1147:11] Saving precursor levels matrix
[1147:11] Precursor levels matrix (1% precursor and protein group FDR) saved to D:\Soumyadeep\Cell Click\out\click_canfinal_out-first-pass.pr_matrix.tsv.
[1147:11] Manifest saved to D:\Soumyadeep\Cell Click\out\click_canfinal_out-first-pass.manifest.txt
[1147:11] Stats report saved to D:\Soumyadeep\Cell Click\out\click_canfinal_out-first-pass.stats.tsv
[1147:11] Generating spectral library:
[1147:11] 13265 target and 662 decoy precursors saved
[1147:12] Spectral library saved to D:\Soumyadeep\Cell Click\out\click_canfinal_out-lib.parquet

[1147:13] Loading spectral library D:\Soumyadeep\Cell Click\out\click_canfinal_out-lib.parquet
[1147:13] Spectral library loaded: 3625 protein isoforms, 3487 protein groups and 13927 precursors in 11906 elution groups.
[1147:13] Loading protein annotations from FASTA camprotR_240512_cRAP_20190401_full_tags.fasta
[1147:13] Loading protein annotations from FASTA D:\Soumyadeep\Cell Click\clean_humwholeprot_reviewed_can.fasta
[1147:13] Annotating library proteins with information from the FASTA database
[1147:13] Gene names missing for some isoforms
[1147:13] Library contains 3625 proteins, and 3617 genes
[1147:13] Initialising library
[1147:14] Saving the library to D:\Soumyadeep\Cell Click\out\click_canfinal_out-lib.parquet.skyline.speclib


Second pass: using the newly created spectral library to reanalyse the data

[1147:14] File #1/24
[1147:14] Loading run D:\Soumyadeep\Cell Click\Data\Cell Run 1 MS Data\Positive Click\HMC3\HA_EL3_T1.wiff
[1153:59] Pre-processing...
[1153:59] 1117 MS1 and 111700 MS2 scans in 1117 (inferred) and 1117 (encoded) cycles, 13265 precursors in range
[1153:59] Calibrating with mass accuracies 25 (MS1), 25 (MS2)
[1154:00] RT window set to 1.62703
[1154:00] Recommended MS1 mass accuracy setting: 28 ppm
[1154:00] Searching decoys
[1154:01] Main search
[1154:01] Removing low confidence identifications
[1154:02] Removing interfering precursors
[1154:02] Training neural networks on 11244 target and 6094 decoy PSMs
[1154:09] Training neural networks on 11235 target and 5948 decoy PSMs
[1154:15] Number of IDs at 0.01 FDR: 9621
[1154:15] Precursors at 1% peptidoform FDR: 6425
[1154:15] Calculating protein q-values
[1154:15] Number of genes identified at 1% FDR: 2025 (precursor-level), 1781 (protein-level) (inference performed using proteotypic peptides only)
[1154:15] Quantification
[1154:16] Precursors with scored PTMs at 1% FDR: 147 out of 203 considered
[1154:16] Precursors with all scored PTM sites unoccupied at 1% FDR: 6278
[1154:16] Precursors with PTMs localised (when required) with > 90% confidence: 143 out of 147
[1154:16] XICs saved to D:\Soumyadeep\Cell Click\out/click_canfinal_out_xic/HA_EL3_T1.xic.parquet

[1154:16] File #2/24
[1154:16] Loading run D:\Soumyadeep\Cell Click\Data\Cell Run 1 MS Data\Positive Click\HMC3\HA_EL3_T2.wiff
[1162:00] Pre-processing...
[1162:00] 1117 MS1 and 111700 MS2 scans in 1117 (inferred) and 1117 (encoded) cycles, 13265 precursors in range
[1162:00] Calibrating with mass accuracies 25 (MS1), 30 (MS2)
[1162:01] RT window set to 1.68753
[1162:01] Recommended MS1 mass accuracy setting: 26 ppm
[1162:01] Searching decoys
[1162:02] Main search
[1162:03] Removing low confidence identifications
[1162:03] Removing interfering precursors
[1162:04] Training neural networks on 11276 target and 6152 decoy PSMs
[1162:10] Training neural networks on 11270 target and 5982 decoy PSMs
[1162:17] Number of IDs at 0.01 FDR: 9760
[1162:17] Precursors at 1% peptidoform FDR: 6748
[1162:17] Calculating protein q-values
[1162:17] Number of genes identified at 1% FDR: 2062 (precursor-level), 1832 (protein-level) (inference performed using proteotypic peptides only)
[1162:17] Quantification
[1162:17] Precursors with scored PTMs at 1% FDR: 158 out of 205 considered
[1162:17] Precursors with all scored PTM sites unoccupied at 1% FDR: 6590
[1162:17] Precursors with PTMs localised (when required) with > 90% confidence: 154 out of 158
[1162:17] XICs saved to D:\Soumyadeep\Cell Click\out/click_canfinal_out_xic/HA_EL3_T2.xic.parquet

[1162:18] File #3/24
[1162:18] Loading run D:\Soumyadeep\Cell Click\Data\Cell Run 1 MS Data\Positive Click\HMC3\HA_EL3_T3.wiff
[1169:37] Pre-processing...
[1169:38] 1117 MS1 and 111700 MS2 scans in 1117 (inferred) and 1117 (encoded) cycles, 13265 precursors in range
[1169:38] Calibrating with mass accuracies 25 (MS1), 29 (MS2)
[1169:38] RT window set to 1.73964
[1169:38] Recommended MS1 mass accuracy setting: 26 ppm
[1169:39] Searching decoys
[1169:39] Main search
[1169:40] Removing low confidence identifications
[1169:41] Removing interfering precursors
[1169:41] Training neural networks on 11239 target and 6083 decoy PSMs
[1169:47] Training neural networks on 11229 target and 5962 decoy PSMs
[1169:54] Number of IDs at 0.01 FDR: 9738
[1169:54] Precursors at 1% peptidoform FDR: 5955
[1169:54] Calculating protein q-values
[1169:54] Number of genes identified at 1% FDR: 2043 (precursor-level), 1758 (protein-level) (inference performed using proteotypic peptides only)
[1169:54] Quantification
[1169:54] Precursors with scored PTMs at 1% FDR: 137 out of 201 considered
[1169:54] Precursors with all scored PTM sites unoccupied at 1% FDR: 5818
[1169:54] Precursors with PTMs localised (when required) with > 90% confidence: 134 out of 137
[1169:55] XICs saved to D:\Soumyadeep\Cell Click\out/click_canfinal_out_xic/HA_EL3_T3.xic.parquet

[1169:55] File #4/24
[1169:55] Loading run D:\Soumyadeep\Cell Click\Data\Cell Run 1 MS Data\Positive Click\HMC3\HA_EL3_T4.wiff
[1175:40] Pre-processing...
[1175:40] 1117 MS1 and 111700 MS2 scans in 1117 (inferred) and 1117 (encoded) cycles, 13265 precursors in range
[1175:40] Calibrating with mass accuracies 25 (MS1), 25 (MS2)
[1175:41] RT window set to 4.06779
[1175:41] Cannot perform MS1 mass calibration, too few confidently identified precursors
[1175:44] Searching decoys
[1175:45] Main search
[1175:46] Removing low confidence identifications
[1175:47] Removing interfering precursors
[1175:47] Training neural networks on 4456 target and 3991 decoy PSMs
[1175:51] Training neural networks on 4448 target and 3933 decoy PSMs
[1175:56] Number of IDs at 0.01 FDR: 44
[1175:56] Precursors at 1% peptidoform FDR: 0
[1175:56] Calculating protein q-values
[1175:56] Number of genes identified at 1% FDR: 17 (precursor-level), 0 (protein-level) (inference performed using proteotypic peptides only)
[1175:56] Quantification
[1175:56] Precursors with scored PTMs at 1% FDR: 0 out of 0 considered
[1175:56] Precursors with all scored PTM sites unoccupied at 1% FDR: 0
[1175:56] Precursors with PTMs localised (when required) with > 90% confidence: 0 out of 0
[1175:56] XICs saved to D:\Soumyadeep\Cell Click\out/click_canfinal_out_xic/HA_EL3_T4.xic.parquet

[1175:57] File #5/24
[1175:57] Loading run D:\Soumyadeep\Cell Click\Data\Cell Run 1 MS Data\Positive Click\Inhibited\HiA_EL3_T1.wiff
[1181:26] Pre-processing...
[1181:26] 1117 MS1 and 111700 MS2 scans in 1117 (inferred) and 1117 (encoded) cycles, 13265 precursors in range
[1181:26] Calibrating with mass accuracies 25 (MS1), 25 (MS2)
[1181:27] RT window set to 1.35502
[1181:27] Recommended MS1 mass accuracy setting: 22 ppm
[1181:27] Searching decoys
[1181:28] Main search
[1181:28] Removing low confidence identifications
[1181:28] Removing interfering precursors
[1181:28] Training neural networks on 5066 target and 3178 decoy PSMs
[1181:33] Training neural networks on 5065 target and 3176 decoy PSMs
[1181:37] Number of IDs at 0.01 FDR: 1079
[1181:37] Precursors at 1% peptidoform FDR: 145
[1181:37] Calculating protein q-values
[1181:37] Number of genes identified at 1% FDR: 401 (precursor-level), 328 (protein-level) (inference performed using proteotypic peptides only)
[1181:37] Quantification
[1181:38] Precursors with scored PTMs at 1% FDR: 0 out of 15 considered
[1181:38] Precursors with all scored PTM sites unoccupied at 1% FDR: 145
[1181:38] Precursors with PTMs localised (when required) with > 90% confidence: 0 out of 0
[1181:38] XICs saved to D:\Soumyadeep\Cell Click\out/click_canfinal_out_xic/HiA_EL3_T1.xic.parquet

[1181:38] File #6/24
[1181:38] Loading run D:\Soumyadeep\Cell Click\Data\Cell Run 1 MS Data\Positive Click\Inhibited\HiA_EL3_T2.wiff
[1186:39] Pre-processing...
[1186:39] 1117 MS1 and 111700 MS2 scans in 1117 (inferred) and 1117 (encoded) cycles, 13265 precursors in range
[1186:39] Calibrating with mass accuracies 25 (MS1), 25 (MS2)
[1186:40] RT window set to 1.56491
[1186:40] Recommended MS1 mass accuracy setting: 24 ppm
[1186:41] Searching decoys
[1186:41] Main search
[1186:41] Removing low confidence identifications
[1186:41] Removing interfering precursors
[1186:41] Training neural networks on 5829 target and 4021 decoy PSMs
[1186:46] Training neural networks on 5822 target and 3891 decoy PSMs
[1186:51] Number of IDs at 0.01 FDR: 1204
[1186:51] Precursors at 1% peptidoform FDR: 757
[1186:51] Calculating protein q-values
[1186:51] Number of genes identified at 1% FDR: 433 (precursor-level), 318 (protein-level) (inference performed using proteotypic peptides only)
[1186:51] Quantification
[1186:51] Precursors with scored PTMs at 1% FDR: 11 out of 19 considered
[1186:51] Precursors with all scored PTM sites unoccupied at 1% FDR: 746
[1186:51] Precursors with PTMs localised (when required) with > 90% confidence: 11 out of 11
[1186:51] XICs saved to D:\Soumyadeep\Cell Click\out/click_canfinal_out_xic/HiA_EL3_T2.xic.parquet

[1186:51] File #7/24
[1186:51] Loading run D:\Soumyadeep\Cell Click\Data\Cell Run 1 MS Data\Positive Click\Inhibited\HiA_EL3_T3.wiff
[1192:18] Pre-processing...
[1192:19] 1117 MS1 and 111700 MS2 scans in 1117 (inferred) and 1117 (encoded) cycles, 13265 precursors in range
[1192:19] Calibrating with mass accuracies 25 (MS1), 25 (MS2)
[1192:20] RT window set to 1.25053
[1192:20] Recommended MS1 mass accuracy setting: 24 ppm
[1192:20] Searching decoys
[1192:20] Main search
[1192:20] Removing low confidence identifications
[1192:20] Removing interfering precursors
[1192:20] Training neural networks on 5065 target and 3334 decoy PSMs
[1192:25] Training neural networks on 5067 target and 3151 decoy PSMs
[1192:30] Number of IDs at 0.01 FDR: 1022
[1192:30] Precursors at 1% peptidoform FDR: 368
[1192:30] Calculating protein q-values
[1192:30] Number of genes identified at 1% FDR: 364 (precursor-level), 246 (protein-level) (inference performed using proteotypic peptides only)
[1192:30] Quantification
[1192:30] Precursors with scored PTMs at 1% FDR: 3 out of 17 considered
[1192:30] Precursors with all scored PTM sites unoccupied at 1% FDR: 365
[1192:30] Precursors with PTMs localised (when required) with > 90% confidence: 3 out of 3
[1192:30] XICs saved to D:\Soumyadeep\Cell Click\out/click_canfinal_out_xic/HiA_EL3_T3.xic.parquet

[1192:30] File #8/24
[1192:30] Loading run D:\Soumyadeep\Cell Click\Data\Cell Run 1 MS Data\Positive Click\Inhibited\HiA_EL3_T4.wiff
[1197:58] Pre-processing...
[1197:58] 1117 MS1 and 111700 MS2 scans in 1117 (inferred) and 1117 (encoded) cycles, 13265 precursors in range
[1197:58] Calibrating with mass accuracies 25 (MS1), 25 (MS2)
[1197:59] RT window set to 2.76095
[1197:59] Recommended MS1 mass accuracy setting: 21 ppm
[1197:59] Searching decoys
[1197:59] Main search
[1198:00] Removing low confidence identifications
[1198:00] Removing interfering precursors
[1198:00] Training neural networks on 5972 target and 3879 decoy PSMs
[1198:05] Training neural networks on 5973 target and 3728 decoy PSMs
[1198:10] Number of IDs at 0.01 FDR: 1205
[1198:10] Precursors at 1% peptidoform FDR: 317
[1198:10] Calculating protein q-values
[1198:10] Number of genes identified at 1% FDR: 429 (precursor-level), 332 (protein-level) (inference performed using proteotypic peptides only)
[1198:10] Quantification
[1198:10] Precursors with scored PTMs at 1% FDR: 0 out of 13 considered
[1198:10] Precursors with all scored PTM sites unoccupied at 1% FDR: 317
[1198:10] Precursors with PTMs localised (when required) with > 90% confidence: 0 out of 0
[1198:10] XICs saved to D:\Soumyadeep\Cell Click\out/click_canfinal_out_xic/HiA_EL3_T4.xic.parquet

[1198:10] File #9/24
[1198:10] Loading run D:\Soumyadeep\Cell Click\Data\Cell Run 1 MS Data\Positive Click\Activated\HaA_EL3_T1.wiff
[1204:02] Pre-processing...
[1204:02] 1117 MS1 and 111700 MS2 scans in 1117 (inferred) and 1117 (encoded) cycles, 13265 precursors in range
[1204:02] Calibrating with mass accuracies 25 (MS1), 31 (MS2)
[1204:03] Recommended MS1 mass accuracy setting: 21 ppm
[1204:04] RT window set to 2.24017
[1204:04] Recommended MS1 mass accuracy setting: 22 ppm
[1204:05] Searching decoys
[1204:05] Main search
[1204:05] Removing low confidence identifications
[1204:06] Removing interfering precursors
[1204:06] Training neural networks on 6207 target and 4058 decoy PSMs
[1204:11] Training neural networks on 6197 target and 3998 decoy PSMs
[1204:16] Number of IDs at 0.01 FDR: 3680
[1204:16] Precursors at 1% peptidoform FDR: 2867
[1204:16] Calculating protein q-values
[1204:16] Number of genes identified at 1% FDR: 1098 (precursor-level), 880 (protein-level) (inference performed using proteotypic peptides only)
[1204:16] Quantification
[1204:16] Precursors with scored PTMs at 1% FDR: 41 out of 55 considered
[1204:16] Precursors with all scored PTM sites unoccupied at 1% FDR: 2826
[1204:16] Precursors with PTMs localised (when required) with > 90% confidence: 40 out of 41
[1204:16] XICs saved to D:\Soumyadeep\Cell Click\out/click_canfinal_out_xic/HaA_EL3_T1.xic.parquet

[1204:17] File #10/24
[1204:17] Loading run D:\Soumyadeep\Cell Click\Data\Cell Run 1 MS Data\Positive Click\Activated\HaA_EL3_T2.wiff
[1209:59] Pre-processing...
[1209:59] 1117 MS1 and 111700 MS2 scans in 1117 (inferred) and 1117 (encoded) cycles, 13265 precursors in range
[1209:59] Calibrating with mass accuracies 25 (MS1), 30 (MS2)
[1210:00] RT window set to 1.87039
[1210:00] Recommended MS1 mass accuracy setting: 25 ppm
[1210:01] Searching decoys
[1210:01] Main search
[1210:02] Removing low confidence identifications
[1210:02] Removing interfering precursors
[1210:02] Training neural networks on 7194 target and 4245 decoy PSMs
[1210:07] Training neural networks on 7187 target and 4116 decoy PSMs
[1210:13] Number of IDs at 0.01 FDR: 4009
[1210:13] Precursors at 1% peptidoform FDR: 2931
[1210:13] Calculating protein q-values
[1210:13] Number of genes identified at 1% FDR: 1135 (precursor-level), 843 (protein-level) (inference performed using proteotypic peptides only)
[1210:13] Quantification
[1210:13] Precursors with scored PTMs at 1% FDR: 45 out of 67 considered
[1210:13] Precursors with all scored PTM sites unoccupied at 1% FDR: 2886
[1210:13] Precursors with PTMs localised (when required) with > 90% confidence: 42 out of 45
[1210:13] XICs saved to D:\Soumyadeep\Cell Click\out/click_canfinal_out_xic/HaA_EL3_T2.xic.parquet

[1210:13] File #11/24
[1210:13] Loading run D:\Soumyadeep\Cell Click\Data\Cell Run 1 MS Data\Positive Click\Activated\HaA_EL3_T3.wiff
[1216:12] Pre-processing...
[1216:12] 1117 MS1 and 111700 MS2 scans in 1117 (inferred) and 1117 (encoded) cycles, 13265 precursors in range
[1216:12] Calibrating with mass accuracies 25 (MS1), 30 (MS2)
[1216:14] RT window set to 2.06548
[1216:14] Recommended MS1 mass accuracy setting: 26 ppm
[1216:15] Searching decoys
[1216:15] Main search
[1216:15] Removing low confidence identifications
[1216:16] Removing interfering precursors
[1216:16] Training neural networks on 7544 target and 4266 decoy PSMs
[1216:21] Training neural networks on 7531 target and 4163 decoy PSMs
[1216:26] Number of IDs at 0.01 FDR: 4172
[1216:26] Precursors at 1% peptidoform FDR: 2695
[1216:26] Calculating protein q-values
[1216:26] Number of genes identified at 1% FDR: 1150 (precursor-level), 944 (protein-level) (inference performed using proteotypic peptides only)
[1216:26] Quantification
[1216:27] Precursors with scored PTMs at 1% FDR: 41 out of 60 considered
[1216:27] Precursors with all scored PTM sites unoccupied at 1% FDR: 2654
[1216:27] Precursors with PTMs localised (when required) with > 90% confidence: 41 out of 41
[1216:27] XICs saved to D:\Soumyadeep\Cell Click\out/click_canfinal_out_xic/HaA_EL3_T3.xic.parquet

[1216:27] File #12/24
[1216:27] Loading run D:\Soumyadeep\Cell Click\Data\Cell Run 1 MS Data\Positive Click\Activated\HaA_EL3_T4.wiff
[1222:35] Pre-processing...
[1222:35] 1117 MS1 and 111700 MS2 scans in 1117 (inferred) and 1117 (encoded) cycles, 13265 precursors in range
[1222:35] Calibrating with mass accuracies 25 (MS1), 25 (MS2)
[1222:37] RT window set to 2.90668
[1222:37] Recommended MS1 mass accuracy setting: 28 ppm
[1222:38] Searching decoys
[1222:38] Main search
[1222:39] Removing low confidence identifications
[1222:40] Removing interfering precursors
[1222:40] Training neural networks on 8048 target and 4451 decoy PSMs
[1222:45] Training neural networks on 8043 target and 4332 decoy PSMs
[1222:50] Number of IDs at 0.01 FDR: 4712
[1222:50] Precursors at 1% peptidoform FDR: 3778
[1222:50] Calculating protein q-values
[1222:50] Number of genes identified at 1% FDR: 1222 (precursor-level), 1037 (protein-level) (inference performed using proteotypic peptides only)
[1222:50] Quantification
[1222:51] Precursors with scored PTMs at 1% FDR: 51 out of 59 considered
[1222:51] Precursors with all scored PTM sites unoccupied at 1% FDR: 3727
[1222:51] Precursors with PTMs localised (when required) with > 90% confidence: 50 out of 51
[1222:51] XICs saved to D:\Soumyadeep\Cell Click\out/click_canfinal_out_xic/HaA_EL3_T4.xic.parquet

[1222:51] File #13/24
[1222:51] Loading run D:\Soumyadeep\Cell Click\Data\Cell Run 1 MS Data\No Azide Control\HMC3\HO_EL3_T1.wiff
[1228:28] Pre-processing...
[1228:28] 1117 MS1 and 111700 MS2 scans in 1117 (inferred) and 1117 (encoded) cycles, 13265 precursors in range
[1228:28] Calibrating with mass accuracies 25 (MS1), 31 (MS2)
[1228:30] Recommended MS1 mass accuracy setting: 12 ppm
[1228:31] RT window set to 7.09285
[1228:31] Recommended MS1 mass accuracy setting: 12 ppm
[1228:34] Searching decoys
[1228:35] Main search
[1228:37] Removing low confidence identifications
[1228:37] Removing interfering precursors
[1228:37] Training neural networks on 4215 target and 3768 decoy PSMs
[1228:42] Training neural networks on 4207 target and 3779 decoy PSMs
[1228:47] Number of IDs at 0.01 FDR: 2
[1228:47] Disabling neural networks
[1228:47] Number of IDs at 0.01 FDR: 0
[1228:47] Precursors at 1% peptidoform FDR: 0
[1228:47] Calculating protein q-values
[1228:47] Number of genes identified at 1% FDR: 0 (precursor-level), 0 (protein-level) (inference performed using proteotypic peptides only)
[1228:47] Quantification
[1228:48] Precursors with scored PTMs at 1% FDR: 0 out of 0 considered
[1228:48] Precursors with all scored PTM sites unoccupied at 1% FDR: 0
[1228:48] Precursors with PTMs localised (when required) with > 90% confidence: 0 out of 0
[1228:48] XICs saved to D:\Soumyadeep\Cell Click\out/click_canfinal_out_xic/HO_EL3_T1.xic.parquet

[1228:48] File #14/24
[1228:48] Loading run D:\Soumyadeep\Cell Click\Data\Cell Run 1 MS Data\No Azide Control\HMC3\HO_EL3_T2.wiff
[1234:39] Pre-processing...
[1234:39] 1117 MS1 and 111700 MS2 scans in 1117 (inferred) and 1117 (encoded) cycles, 13265 precursors in range
[1234:39] Calibrating with mass accuracies 25 (MS1), 26 (MS2)
[1234:40] RT window set to 5.65269
[1234:40] Recommended MS1 mass accuracy setting: 13 ppm
[1234:43] Searching decoys
[1234:44] Main search
[1234:46] Removing low confidence identifications
[1234:47] Removing interfering precursors
[1234:47] Training neural networks on 10012 target and 9409 decoy PSMs
[1234:54] Training neural networks on 9993 target and 9366 decoy PSMs
[1235:01] Number of IDs at 0.01 FDR: 0
[1235:01] Disabling neural networks
[1235:01] Number of IDs at 0.01 FDR: 0
[1235:01] Precursors at 1% peptidoform FDR: 0
[1235:01] Calculating protein q-values
[1235:01] Number of genes identified at 1% FDR: 0 (precursor-level), 0 (protein-level) (inference performed using proteotypic peptides only)
[1235:01] Quantification
[1235:01] Precursors with scored PTMs at 1% FDR: 0 out of 0 considered
[1235:01] Precursors with all scored PTM sites unoccupied at 1% FDR: 0
[1235:01] Precursors with PTMs localised (when required) with > 90% confidence: 0 out of 0
[1235:01] XICs saved to D:\Soumyadeep\Cell Click\out/click_canfinal_out_xic/HO_EL3_T2.xic.parquet

[1235:01] File #15/24
[1235:01] Loading run D:\Soumyadeep\Cell Click\Data\Cell Run 1 MS Data\No Azide Control\HMC3\HO_EL3_T3.wiff
[1236:32] Pre-processing...
[1236:32] 1117 MS1 and 111700 MS2 scans in 1117 (inferred) and 1117 (encoded) cycles, 13265 precursors in range
[1236:32] Calibrating with mass accuracies 25 (MS1), 27 (MS2)
[1236:33] RT window set to 6.15308
[1236:33] Recommended MS1 mass accuracy setting: 9 ppm
[1236:36] Searching decoys
[1236:37] Main search
[1236:39] Removing low confidence identifications
[1236:39] Removing interfering precursors
[1236:39] Training neural networks on 4108 target and 3690 decoy PSMs
[1236:44] Training neural networks on 4093 target and 3607 decoy PSMs
[1236:50] Number of IDs at 0.01 FDR: 10
[1236:50] Disabling neural networks
[1236:50] Number of IDs at 0.01 FDR: 0
[1236:50] Precursors at 1% peptidoform FDR: 0
[1236:50] Calculating protein q-values
[1236:50] Number of genes identified at 1% FDR: 0 (precursor-level), 0 (protein-level) (inference performed using proteotypic peptides only)
[1236:50] Quantification
[1236:50] Precursors with scored PTMs at 1% FDR: 0 out of 0 considered
[1236:50] Precursors with all scored PTM sites unoccupied at 1% FDR: 0
[1236:50] Precursors with PTMs localised (when required) with > 90% confidence: 0 out of 0
[1236:51] XICs saved to D:\Soumyadeep\Cell Click\out/click_canfinal_out_xic/HO_EL3_T3.xic.parquet

[1236:51] File #16/24
[1236:51] Loading run D:\Soumyadeep\Cell Click\Data\Cell Run 1 MS Data\No Azide Control\HMC3\HO_EL3_T4.wiff
[1238:11] Pre-processing...
[1238:11] 1117 MS1 and 111700 MS2 scans in 1117 (inferred) and 1117 (encoded) cycles, 13265 precursors in range
[1238:11] Calibrating with mass accuracies 25 (MS1), 27 (MS2)
[1238:15] Searching decoys
[1238:16] Main search
[1238:18] Removing low confidence identifications
[1238:18] Removing interfering precursors
[1238:19] Training neural networks on 9832 target and 9275 decoy PSMs
[1238:27] Training neural networks on 9842 target and 9258 decoy PSMs
[1238:35] Number of IDs at 0.01 FDR: 0
[1238:35] Disabling neural networks
[1238:35] Number of IDs at 0.01 FDR: 0
[1238:35] Precursors at 1% peptidoform FDR: 0
[1238:35] Calculating protein q-values
[1238:35] Number of genes identified at 1% FDR: 0 (precursor-level), 0 (protein-level) (inference performed using proteotypic peptides only)
[1238:35] Quantification
[1238:35] Precursors with scored PTMs at 1% FDR: 0 out of 0 considered
[1238:35] Precursors with all scored PTM sites unoccupied at 1% FDR: 0
[1238:35] Precursors with PTMs localised (when required) with > 90% confidence: 0 out of 0
[1238:35] XICs saved to D:\Soumyadeep\Cell Click\out/click_canfinal_out_xic/HO_EL3_T4.xic.parquet

[1238:35] File #17/24
[1238:35] Loading run D:\Soumyadeep\Cell Click\Data\Cell Run 1 MS Data\No Azide Control\Inhibited\HiO_EL3_T1.wiff
[1239:56] Pre-processing...
[1239:56] 1117 MS1 and 111700 MS2 scans in 1117 (inferred) and 1117 (encoded) cycles, 13265 precursors in range
[1239:56] Calibrating with mass accuracies 25 (MS1), 29 (MS2)
[1239:58] RT window set to 6.20103
[1239:58] Recommended MS1 mass accuracy setting: 14 ppm
[1240:01] Searching decoys
[1240:02] Main search
[1240:04] Removing low confidence identifications
[1240:05] Removing interfering precursors
[1240:05] Training neural networks on 4289 target and 3913 decoy PSMs
[1240:10] Training neural networks on 4292 target and 3808 decoy PSMs
[1240:16] Number of IDs at 0.01 FDR: 0
[1240:16] Disabling neural networks
[1240:16] Number of IDs at 0.01 FDR: 0
[1240:16] Precursors at 1% peptidoform FDR: 0
[1240:16] Calculating protein q-values
[1240:16] Number of genes identified at 1% FDR: 0 (precursor-level), 0 (protein-level) (inference performed using proteotypic peptides only)
[1240:16] Quantification
[1240:17] Precursors with scored PTMs at 1% FDR: 0 out of 0 considered
[1240:17] Precursors with all scored PTM sites unoccupied at 1% FDR: 0
[1240:17] Precursors with PTMs localised (when required) with > 90% confidence: 0 out of 0
[1240:17] XICs saved to D:\Soumyadeep\Cell Click\out/click_canfinal_out_xic/HiO_EL3_T1.xic.parquet

[1240:17] File #18/24
[1240:17] Loading run D:\Soumyadeep\Cell Click\Data\Cell Run 1 MS Data\No Azide Control\Inhibited\HiO_EL3_T2.wiff
[1241:45] Pre-processing...
[1241:45] 1117 MS1 and 111700 MS2 scans in 1117 (inferred) and 1117 (encoded) cycles, 13265 precursors in range
[1241:45] Calibrating with mass accuracies 25 (MS1), 36 (MS2)
[1241:46] Recommended MS1 mass accuracy setting: 12 ppm
[1241:48] RT window set to 5.13613
[1241:48] Recommended MS1 mass accuracy setting: 13 ppm
[1241:51] Searching decoys
[1241:52] Main search
[1241:54] Removing low confidence identifications
[1241:54] Removing interfering precursors
[1241:54] Training neural networks on 4526 target and 4033 decoy PSMs
[1242:00] Training neural networks on 4526 target and 4042 decoy PSMs
[1242:06] Number of IDs at 0.01 FDR: 18
[1242:06] Disabling neural networks
[1242:06] Number of IDs at 0.01 FDR: 0
[1242:06] Precursors at 1% peptidoform FDR: 0
[1242:06] Calculating protein q-values
[1242:06] Number of genes identified at 1% FDR: 0 (precursor-level), 0 (protein-level) (inference performed using proteotypic peptides only)
[1242:06] Quantification
[1242:06] Precursors with scored PTMs at 1% FDR: 0 out of 0 considered
[1242:06] Precursors with all scored PTM sites unoccupied at 1% FDR: 0
[1242:06] Precursors with PTMs localised (when required) with > 90% confidence: 0 out of 0
[1242:06] XICs saved to D:\Soumyadeep\Cell Click\out/click_canfinal_out_xic/HiO_EL3_T2.xic.parquet

[1242:06] File #19/24
[1242:06] Loading run D:\Soumyadeep\Cell Click\Data\Cell Run 1 MS Data\No Azide Control\Inhibited\HiO_EL3_T3.wiff
[1243:29] Pre-processing...
[1243:29] 1117 MS1 and 111700 MS2 scans in 1117 (inferred) and 1117 (encoded) cycles, 13265 precursors in range
[1243:29] Calibrating with mass accuracies 25 (MS1), 39 (MS2)
[1243:31] Recommended MS1 mass accuracy setting: 12 ppm
[1243:32] RT window set to 4.63458
[1243:32] Recommended MS1 mass accuracy setting: 11 ppm
[1243:36] Searching decoys
[1243:37] Main search
[1243:39] Removing low confidence identifications
[1243:39] Removing interfering precursors
[1243:39] Training neural networks on 4436 target and 4121 decoy PSMs
[1243:45] Training neural networks on 4430 target and 4004 decoy PSMs
[1243:51] Number of IDs at 0.01 FDR: 9
[1243:51] Disabling neural networks
[1243:51] Number of IDs at 0.01 FDR: 0
[1243:51] Precursors at 1% peptidoform FDR: 0
[1243:51] Calculating protein q-values
[1243:51] Number of genes identified at 1% FDR: 0 (precursor-level), 0 (protein-level) (inference performed using proteotypic peptides only)
[1243:51] Quantification
[1243:51] Precursors with scored PTMs at 1% FDR: 0 out of 0 considered
[1243:51] Precursors with all scored PTM sites unoccupied at 1% FDR: 0
[1243:51] Precursors with PTMs localised (when required) with > 90% confidence: 0 out of 0
[1243:51] XICs saved to D:\Soumyadeep\Cell Click\out/click_canfinal_out_xic/HiO_EL3_T3.xic.parquet

[1243:51] File #20/24
[1243:51] Loading run D:\Soumyadeep\Cell Click\Data\Cell Run 1 MS Data\No Azide Control\Inhibited\HiO_EL3_T4.wiff
[1245:03] Pre-processing...
[1245:03] 1117 MS1 and 111700 MS2 scans in 1117 (inferred) and 1117 (encoded) cycles, 13265 precursors in range
[1245:03] Calibrating with mass accuracies 25 (MS1), 39 (MS2)
[1245:09] Searching decoys
[1245:09] Main search
[1245:11] Removing low confidence identifications
[1245:12] Removing interfering precursors
[1245:12] Training neural networks on 10020 target and 9458 decoy PSMs
[1245:20] Training neural networks on 10020 target and 9430 decoy PSMs
[1245:29] Number of IDs at 0.01 FDR: 0
[1245:29] Disabling neural networks
[1245:29] Number of IDs at 0.01 FDR: 0
[1245:29] Precursors at 1% peptidoform FDR: 0
[1245:29] Calculating protein q-values
[1245:29] Number of genes identified at 1% FDR: 0 (precursor-level), 0 (protein-level) (inference performed using proteotypic peptides only)
[1245:29] Quantification
[1245:29] Precursors with scored PTMs at 1% FDR: 0 out of 0 considered
[1245:29] Precursors with all scored PTM sites unoccupied at 1% FDR: 0
[1245:29] Precursors with PTMs localised (when required) with > 90% confidence: 0 out of 0
[1245:29] XICs saved to D:\Soumyadeep\Cell Click\out/click_canfinal_out_xic/HiO_EL3_T4.xic.parquet

[1245:29] File #21/24
[1245:29] Loading run D:\Soumyadeep\Cell Click\Data\Cell Run 1 MS Data\No Azide Control\Activated\HaO_EL3_T1.wiff
[1246:52] Pre-processing...
[1246:53] 1117 MS1 and 111700 MS2 scans in 1117 (inferred) and 1117 (encoded) cycles, 13265 precursors in range
[1246:53] Calibrating with mass accuracies 25 (MS1), 37 (MS2)
[1246:54] Recommended MS1 mass accuracy setting: 11 ppm
[1246:55] RT window set to 5.61095
[1246:55] Recommended MS1 mass accuracy setting: 8 ppm
[1246:59] Searching decoys
[1247:00] Main search
[1247:02] Removing low confidence identifications
[1247:02] Removing interfering precursors
[1247:02] Training neural networks on 4484 target and 4055 decoy PSMs
[1247:08] Training neural networks on 4476 target and 3942 decoy PSMs
[1247:14] Number of IDs at 0.01 FDR: 38
[1247:14] Disabling neural networks
[1247:14] Number of IDs at 0.01 FDR: 0
[1247:14] Precursors at 1% peptidoform FDR: 0
[1247:14] Calculating protein q-values
[1247:14] Number of genes identified at 1% FDR: 0 (precursor-level), 0 (protein-level) (inference performed using proteotypic peptides only)
[1247:14] Quantification
[1247:14] Precursors with scored PTMs at 1% FDR: 0 out of 0 considered
[1247:14] Precursors with all scored PTM sites unoccupied at 1% FDR: 0
[1247:14] Precursors with PTMs localised (when required) with > 90% confidence: 0 out of 0
[1247:14] XICs saved to D:\Soumyadeep\Cell Click\out/click_canfinal_out_xic/HaO_EL3_T1.xic.parquet

[1247:14] File #22/24
[1247:14] Loading run D:\Soumyadeep\Cell Click\Data\Cell Run 1 MS Data\No Azide Control\Activated\HaO_EL3_T2.wiff
[1248:38] Pre-processing...
[1248:38] 1117 MS1 and 111700 MS2 scans in 1117 (inferred) and 1117 (encoded) cycles, 13265 precursors in range
[1248:38] Calibrating with mass accuracies 25 (MS1), 27 (MS2)
[1248:39] RT window set to 5.57139
[1248:39] Recommended MS1 mass accuracy setting: 17 ppm
[1248:43] Searching decoys
[1248:44] Main search
[1248:46] Removing low confidence identifications
[1248:46] Removing interfering precursors
[1248:46] Training neural networks on 4561 target and 4173 decoy PSMs
[1248:52] Training neural networks on 4547 target and 4192 decoy PSMs
[1248:58] Number of IDs at 0.01 FDR: 29
[1248:58] Disabling neural networks
[1248:58] Number of IDs at 0.01 FDR: 0
[1248:58] Precursors at 1% peptidoform FDR: 0
[1248:58] Calculating protein q-values
[1248:58] Number of genes identified at 1% FDR: 0 (precursor-level), 0 (protein-level) (inference performed using proteotypic peptides only)
[1248:58] Quantification
[1248:58] Precursors with scored PTMs at 1% FDR: 0 out of 0 considered
[1248:58] Precursors with all scored PTM sites unoccupied at 1% FDR: 0
[1248:58] Precursors with PTMs localised (when required) with > 90% confidence: 0 out of 0
[1248:58] XICs saved to D:\Soumyadeep\Cell Click\out/click_canfinal_out_xic/HaO_EL3_T2.xic.parquet

[1248:58] File #23/24
[1248:58] Loading run D:\Soumyadeep\Cell Click\Data\Cell Run 1 MS Data\No Azide Control\Activated\HaO_EL3_T3.wiff
[1250:23] Pre-processing...
[1250:23] 1117 MS1 and 111700 MS2 scans in 1117 (inferred) and 1117 (encoded) cycles, 13265 precursors in range
[1250:23] Calibrating with mass accuracies 25 (MS1), 28 (MS2)
[1250:24] RT window set to 4.99385
[1250:24] Recommended MS1 mass accuracy setting: 9 ppm
[1250:27] Searching decoys
[1250:28] Main search
[1250:30] Removing low confidence identifications
[1250:31] Removing interfering precursors
[1250:31] Training neural networks on 4436 target and 3943 decoy PSMs
[1250:36] Training neural networks on 4433 target and 3976 decoy PSMs
[1250:42] Number of IDs at 0.01 FDR: 10
[1250:42] Disabling neural networks
[1250:42] Number of IDs at 0.01 FDR: 0
[1250:42] Precursors at 1% peptidoform FDR: 0
[1250:42] Calculating protein q-values
[1250:42] Number of genes identified at 1% FDR: 0 (precursor-level), 0 (protein-level) (inference performed using proteotypic peptides only)
[1250:42] Quantification
[1250:42] Precursors with scored PTMs at 1% FDR: 0 out of 0 considered
[1250:42] Precursors with all scored PTM sites unoccupied at 1% FDR: 0
[1250:42] Precursors with PTMs localised (when required) with > 90% confidence: 0 out of 0
[1250:42] XICs saved to D:\Soumyadeep\Cell Click\out/click_canfinal_out_xic/HaO_EL3_T3.xic.parquet

[1250:43] File #24/24
[1250:43] Loading run D:\Soumyadeep\Cell Click\Data\Cell Run 1 MS Data\No Azide Control\Activated\HaO_EL3_T4.wiff
[1252:08] Pre-processing...
[1252:08] 1117 MS1 and 111700 MS2 scans in 1117 (inferred) and 1117 (encoded) cycles, 13265 precursors in range
[1252:08] Calibrating with mass accuracies 25 (MS1), 27 (MS2)
[1252:09] RT window set to 4.79637
[1252:09] Recommended MS1 mass accuracy setting: 9 ppm
[1252:13] Searching decoys
[1252:14] Main search
[1252:16] Removing low confidence identifications
[1252:17] Removing interfering precursors
[1252:17] Training neural networks on 4319 target and 3832 decoy PSMs
[1252:22] Training neural networks on 4319 target and 3807 decoy PSMs
[1252:28] Number of IDs at 0.01 FDR: 36
[1252:28] Disabling neural networks
[1252:28] Number of IDs at 0.01 FDR: 0
[1252:28] Precursors at 1% peptidoform FDR: 0
[1252:28] Calculating protein q-values
[1252:28] Number of genes identified at 1% FDR: 0 (precursor-level), 0 (protein-level) (inference performed using proteotypic peptides only)
[1252:28] Quantification
[1252:28] Precursors with scored PTMs at 1% FDR: 0 out of 0 considered
[1252:28] Precursors with all scored PTM sites unoccupied at 1% FDR: 0
[1252:28] Precursors with PTMs localised (when required) with > 90% confidence: 0 out of 0
[1252:28] XICs saved to D:\Soumyadeep\Cell Click\out/click_canfinal_out_xic/HaO_EL3_T4.xic.parquet

[1252:29] Cross-run analysis
[1252:29] Reading quantification information: 24 files
[1252:29] Quantifying peptides
[1252:38] Quantification parameters: 0.385436, 0.0730652, 1.35519, 0.165904, 0.28189, 0.259526, 0.354243, 0.033201, 0.0850453, 0.141483, 0.117781, 0.128412, 0.547002, 0.301688, 0.378496, 0.0104543
[1252:41] Quantifying proteins
[1252:41] Calculating q-values for protein and gene groups
[1252:42] Calculating global q-values for protein and gene groups
[1252:42] Protein groups with global q-value <= 0.01: 1964
[1252:42] Compressed report saved to D:\Soumyadeep\Cell Click\out\click_canfinal_out.parquet. Use R 'arrow' or Python 'PyArrow' package to process
[1252:42] Site report saved to D:\Soumyadeep\Cell Click\out\click_canfinal_out.site_report.parquet
[1252:42] Saving precursor levels matrix
[1252:42] Precursor levels matrix (1% precursor and protein group FDR) saved to D:\Soumyadeep\Cell Click\out\click_canfinal_out.pr_matrix.tsv.
[1252:42] Saving protein group levels matrix
[1252:42] Protein groups matrix saved to D:\Soumyadeep\Cell Click\out\click_canfinal_out.pg_matrix.tsv.
[1252:42] Saving gene group levels matrix
[1252:42] Gene groups matrix saved to D:\Soumyadeep\Cell Click\out\click_canfinal_out.gg_matrix.tsv.
[1252:42] Saving unique genes levels matrix
[1252:42] Unique genes matrix saved to D:\Soumyadeep\Cell Click\out\click_canfinal_out.unique_genes_matrix.tsv.
[1252:42] Manifest saved to D:\Soumyadeep\Cell Click\out\click_canfinal_out.manifest.txt
[1252:42] Stats report saved to D:\Soumyadeep\Cell Click\out\click_canfinal_out.stats.tsv
