## Supplementary File 3 for "Dynamic s-acylation of schizophrenia-linked antioxidants explicate the redox flexibility of human microglia during inflammation"

DIA-NN 2.2.0 Academia  (Data-Independent Acquisition by Neural Networks)
Compiled on May 29 2025 21:29:29
Current date and time: Fri Jul 11 11:32:04 2025
CPU: GenuineIntel Intel(R) Xeon(R) Gold 6134 CPU @ 3.20GHz
SIMD instructions: AVX AVX2 AVX512CD AVX512F FMA SSE4.1 SSE4.2 
Logical CPU cores: 32
diann.exe --f D:\USERS\Soumyadeep\Cell_Lysates\270625_cell\H_T1.wiff  --f D:\USERS\Soumyadeep\Cell_Lysates\270625_cell\H_T2.wiff  --f D:\USERS\Soumyadeep\Cell_Lysates\270625_cell\H_T3.wiff  --f D:\USERS\Soumyadeep\Cell_Lysates\270625_cell\HA_T1.wiff  --f D:\USERS\Soumyadeep\Cell_Lysates\270625_cell\HA_T2.wiff  --f D:\USERS\Soumyadeep\Cell_Lysates\270625_cell\HA_T3.wiff  --f D:\USERS\Soumyadeep\Cell_Lysates\270625_cell\HI_T1.wiff  --f D:\USERS\Soumyadeep\Cell_Lysates\270625_cell\HI_T2.wiff  --f D:\USERS\Soumyadeep\Cell_Lysates\270625_cell\HI_T3.wiff  --f D:\USERS\Soumyadeep\Cell_Lysates\270625_cell\HIA_T1.wiff  --f D:\USERS\Soumyadeep\Cell_Lysates\270625_cell\HIA_T2.wiff  --f D:\USERS\Soumyadeep\Cell_Lysates\270625_cell\HIA_T3.wiff  --f D:\USERS\Soumyadeep\020725\E2\H_E2_T1.wiff  --f D:\USERS\Soumyadeep\020725\E2\H_E2_T2.wiff  --f D:\USERS\Soumyadeep\020725\E2\H_E2_T3.wiff  --f D:\USERS\Soumyadeep\020725\E2\HA_E2_T1.wiff  --f D:\USERS\Soumyadeep\020725\E2\HA_E2_T2.wiff  --f D:\USERS\Soumyadeep\020725\E2\HA_E2_T3.wiff  --f D:\USERS\Soumyadeep\020725\E2\HI_E2_T1.wiff  --f D:\USERS\Soumyadeep\020725\E2\HI_E2_T2.wiff  --f D:\USERS\Soumyadeep\020725\E2\HI_E2_T3.wiff  --f D:\USERS\Soumyadeep\020725\E2\HIA_E2_T1.wiff  --f D:\USERS\Soumyadeep\020725\E2\HIA_E2_T2.wiff  --f D:\USERS\Soumyadeep\020725\E2\HIA_E2_T3.wiff  --f D:\USERS\Soumyadeep\020725\E3\H_E3_T1.wiff  --f D:\USERS\Soumyadeep\020725\E3\H_E3_T2.wiff  --f D:\USERS\Soumyadeep\020725\E3\H_E3_T3.wiff  --f D:\USERS\Soumyadeep\020725\E3\HA_E3_T1.wiff  --f D:\USERS\Soumyadeep\020725\E3\HA_E3_T2.wiff  --f D:\USERS\Soumyadeep\020725\E3\HA_E3_T3.wiff  --f D:\USERS\Soumyadeep\020725\E3\HI_E3_T1.wiff  --f D:\USERS\Soumyadeep\020725\E3\HI_E3_T2.wiff  --f D:\USERS\Soumyadeep\020725\E3\HI_E3_T3.wiff  --f D:\USERS\Soumyadeep\020725\E3\HIA_E3_T1.wiff  --f D:\USERS\Soumyadeep\020725\E3\HIA_E3_T2.wiff  --f D:\USERS\Soumyadeep\020725\E3\HIA_E3_T3.wiff  --lib D:\USERS\Soumyadeep\020725\lib.predicted.speclib --threads 20 --verbose 1 --out D:\USERS\Soumyadeep\CellLysate_DIANN2\HvHIvHA_out\all_celllysates_comparison.parquet --qvalue 0.05 --matrices --xic --fasta camprotR_240512_cRAP_20190401_full_tags.fasta --cont-quant-exclude cRAP- --fasta D:\USERS\Soumyadeep\Fasta 070725\clean_humwholeprot_reviewed_can.fasta --met-excision --cut K*,R*,!*P --missed-cleavages 2 --unimod4 --var-mods 3 --var-mod UniMod:35,15.994915,M --reanalyse --rt-profiling --report-lib-info 

Thread number set to 20
Output will be filtered at 0.05 FDR
Precursor/protein x samples expression level matrices will be saved along with the main report
XICs within 10 seconds from the apex will be extracted for each precursor and saved in .parquet format, a folder will be created next to the main report for the XICs storage
Peptides corresponding to protein sequence IDs tagged with cRAP- will be excluded from normalisation as well as quantification of protein groups that do not include proteins bearing the tag
N-terminal methionine excision enabled
In silico digest will involve cuts at K*,R*
But excluding cuts at *P
Maximum number of missed cleavages set to 2
Cysteine carbamidomethylation enabled as a fixed modification
Maximum number of variable modifications set to 3
Modification UniMod:35 with mass delta 15.9949 at M will be considered as variable
MBR enabled; .quant files will only be saved to disk during the first pass
The spectral library (if generated) will retain the original spectra but will include empirically-aligned RTs
DIA-NN will automatically optimise the mass accuracy for the first run of the experiment, use this mode for preliminary analyses only
WARNING: peptidoform scoring enabled because variable modifications have been declared; to disable, use --no-peptidoforms
The following variable modifications will be localised: UniMod:35 
WARNING: protein inference will only be performed for precursors identified with global q-value <= 0.05 or q-value <= 0.05.

36 files will be processed
[0:00] Loading spectral library D:\USERS\Soumyadeep\020725\lib.predicted.speclib
[0:14] Library annotated with sequence database(s): D:\USERS\Soumyadeep\Fasta 070725\clean_humwholeprot_reviewed_can.fasta
[0:17] Spectral library loaded: 20368 protein isoforms, 29406 protein groups and 7700783 precursors in 2804689 elution groups.
[0:17] Loading protein annotations from FASTA camprotR_240512_cRAP_20190401_full_tags.fasta
[0:17] Loading protein annotations from FASTA D:\USERS\Soumyadeep\Fasta 070725\clean_humwholeprot_reviewed_can.fasta
[0:18] Annotating library proteins with information from the FASTA database
[0:18] Gene names missing for some isoforms
[0:18] Library contains 20368 proteins, and 20174 genes
[0:29] Initialising library

First pass: generating a spectral library from DIA data

[1:05] File #1/36
[1:05] Loading run D:\USERS\Soumyadeep\Cell_Lysates\270625_cell\H_T1.wiff
[10:13] Pre-processing...
[10:17] 1117 MS1 and 111700 MS2 scans in 1117 (inferred) and 1117 (encoded) cycles, 6606089 precursors in range
[10:18] Calibrating with mass accuracies 25 (MS1), 25 (MS2)
[14:44] RT window set to 3.11041
[14:44] Peak width: 2.98
[14:44] Scan window radius set to 6
[14:44] Recommended MS1 mass accuracy setting: 28 ppm
[25:44] Optimised mass accuracy: 20 ppm
[28:34] Searching decoys
[32:05] Main search
[38:56] Removing low confidence identifications
[39:10] Removing interfering precursors
[39:18] Training neural networks on 24875 target and 13431 decoy PSMs
[39:53] Training neural networks on 24875 target and 12146 decoy PSMs
[40:19] Number of IDs at 0.01 FDR: 14319
[40:20] Precursors at 1% peptidoform FDR: 13420
[40:22] Calculating protein q-values
[40:23] Number of genes identified at 1% FDR: 2750 (precursor-level), 2500 (protein-level) (inference performed using proteotypic peptides only)
[40:23] Quantification
[40:24] Precursors with scored PTMs at 1% FDR: 21 out of 68 considered
[40:24] Precursors with all scored PTM sites unoccupied at 1% FDR: 13399
[40:24] Precursors with PTMs localised (when required) with > 90% confidence: 18 out of 21
[40:25] Quantification information saved to D:\USERS\Soumyadeep\Cell_Lysates\270625_cell\H_T1.wiff.quant

[40:26] File #2/36
[40:26] Loading run D:\USERS\Soumyadeep\Cell_Lysates\270625_cell\H_T2.wiff
[49:42] Pre-processing...
[49:46] 1117 MS1 and 111700 MS2 scans in 1117 (inferred) and 1117 (encoded) cycles, 6606089 precursors in range
[49:47] Calibrating with mass accuracies 28 (MS1), 25 (MS2)
[54:15] RT window set to 2.95146
[54:15] Recommended MS1 mass accuracy setting: 29 ppm
[57:36] Searching decoys
[60:44] Main search
[66:49] Removing low confidence identifications
[67:02] Removing interfering precursors
[67:10] Training neural networks on 22896 target and 11980 decoy PSMs
[67:43] Training neural networks on 22896 target and 10920 decoy PSMs
[68:07] Number of IDs at 0.01 FDR: 14143
[68:07] Precursors at 1% peptidoform FDR: 13311
[68:09] Calculating protein q-values
[68:10] Number of genes identified at 1% FDR: 2689 (precursor-level), 2363 (protein-level) (inference performed using proteotypic peptides only)
[68:10] Quantification
[68:11] Precursors with scored PTMs at 1% FDR: 28 out of 68 considered
[68:11] Precursors with all scored PTM sites unoccupied at 1% FDR: 13283
[68:11] Precursors with PTMs localised (when required) with > 90% confidence: 26 out of 28
[68:13] Quantification information saved to D:\USERS\Soumyadeep\Cell_Lysates\270625_cell\H_T2.wiff.quant

[68:14] File #3/36
[68:14] Loading run D:\USERS\Soumyadeep\Cell_Lysates\270625_cell\H_T3.wiff
[78:04] Pre-processing...
[78:08] 1117 MS1 and 111700 MS2 scans in 1117 (inferred) and 1117 (encoded) cycles, 6606089 precursors in range
[78:09] Calibrating with mass accuracies 28 (MS1), 25 (MS2)
[82:44] RT window set to 2.93168
[82:44] Recommended MS1 mass accuracy setting: 27 ppm
[86:15] Searching decoys
[89:56] Main search
[97:13] Removing low confidence identifications
[97:26] Removing interfering precursors
[97:35] Training neural networks on 23435 target and 12087 decoy PSMs
[98:08] Training neural networks on 23435 target and 10927 decoy PSMs
[98:32] Number of IDs at 0.01 FDR: 14345
[98:33] Precursors at 1% peptidoform FDR: 13543
[98:35] Calculating protein q-values
[98:35] Number of genes identified at 1% FDR: 2743 (precursor-level), 2455 (protein-level) (inference performed using proteotypic peptides only)
[98:36] Quantification
[98:36] Precursors with scored PTMs at 1% FDR: 33 out of 72 considered
[98:36] Precursors with all scored PTM sites unoccupied at 1% FDR: 13510
[98:36] Precursors with PTMs localised (when required) with > 90% confidence: 28 out of 33
[98:38] Quantification information saved to D:\USERS\Soumyadeep\Cell_Lysates\270625_cell\H_T3.wiff.quant

[98:39] File #4/36
[98:39] Loading run D:\USERS\Soumyadeep\Cell_Lysates\270625_cell\HA_T1.wiff
[107:02] Pre-processing...
[107:05] 1117 MS1 and 111700 MS2 scans in 1117 (inferred) and 1117 (encoded) cycles, 6606089 precursors in range
[107:07] Calibrating with mass accuracies 28 (MS1), 25 (MS2)
[111:18] RT window set to 2.59995
[111:19] Recommended MS1 mass accuracy setting: 26 ppm
[113:53] Searching decoys
[116:09] Main search
[120:39] Removing low confidence identifications
[120:51] Removing interfering precursors
[120:59] Training neural networks on 22298 target and 10341 decoy PSMs
[121:30] Training neural networks on 22298 target and 9659 decoy PSMs
[121:52] Number of IDs at 0.01 FDR: 14109
[121:52] Precursors at 1% peptidoform FDR: 13208
[121:54] Calculating protein q-values
[121:55] Number of genes identified at 1% FDR: 2421 (precursor-level), 2086 (protein-level) (inference performed using proteotypic peptides only)
[121:55] Quantification
[121:56] Precursors with scored PTMs at 1% FDR: 21 out of 44 considered
[121:56] Precursors with all scored PTM sites unoccupied at 1% FDR: 13187
[121:56] Precursors with PTMs localised (when required) with > 90% confidence: 20 out of 21
[121:57] Quantification information saved to D:\USERS\Soumyadeep\Cell_Lysates\270625_cell\HA_T1.wiff.quant

[121:58] File #5/36
[121:58] Loading run D:\USERS\Soumyadeep\Cell_Lysates\270625_cell\HA_T2.wiff
[130:24] Pre-processing...
[130:27] 1117 MS1 and 111700 MS2 scans in 1117 (inferred) and 1117 (encoded) cycles, 6606089 precursors in range
[130:29] Calibrating with mass accuracies 28 (MS1), 25 (MS2)
[134:38] RT window set to 2.5081
[134:39] Recommended MS1 mass accuracy setting: 26 ppm
[137:09] Searching decoys
[139:16] Main search
[143:32] Removing low confidence identifications
[143:45] Removing interfering precursors
[143:53] Training neural networks on 22201 target and 10527 decoy PSMs
[144:23] Training neural networks on 22201 target and 9817 decoy PSMs
[144:45] Number of IDs at 0.01 FDR: 14033
[144:46] Precursors at 1% peptidoform FDR: 13241
[144:48] Calculating protein q-values
[144:49] Number of genes identified at 1% FDR: 2442 (precursor-level), 2139 (protein-level) (inference performed using proteotypic peptides only)
[144:49] Quantification
[144:49] Precursors with scored PTMs at 1% FDR: 25 out of 47 considered
[144:49] Precursors with all scored PTM sites unoccupied at 1% FDR: 13216
[144:49] Precursors with PTMs localised (when required) with > 90% confidence: 23 out of 25
[144:51] Quantification information saved to D:\USERS\Soumyadeep\Cell_Lysates\270625_cell\HA_T2.wiff.quant

[144:52] File #6/36
[144:52] Loading run D:\USERS\Soumyadeep\Cell_Lysates\270625_cell\HA_T3.wiff
[153:47] Pre-processing...
[153:50] 1117 MS1 and 111700 MS2 scans in 1117 (inferred) and 1117 (encoded) cycles, 6606089 precursors in range
[153:52] Calibrating with mass accuracies 28 (MS1), 25 (MS2)
[158:10] RT window set to 2.42646
[158:11] Recommended MS1 mass accuracy setting: 27 ppm
[160:50] Searching decoys
[163:06] Main search
[167:41] Removing low confidence identifications
[167:54] Removing interfering precursors
[168:02] Training neural networks on 22594 target and 10511 decoy PSMs
[168:33] Training neural networks on 22594 target and 9787 decoy PSMs
[168:55] Number of IDs at 0.01 FDR: 14571
[168:55] Precursors at 1% peptidoform FDR: 13664
[168:57] Calculating protein q-values
[168:58] Number of genes identified at 1% FDR: 2530 (precursor-level), 2246 (protein-level) (inference performed using proteotypic peptides only)
[168:58] Quantification
[168:59] Precursors with scored PTMs at 1% FDR: 21 out of 45 considered
[168:59] Precursors with all scored PTM sites unoccupied at 1% FDR: 13643
[168:59] Precursors with PTMs localised (when required) with > 90% confidence: 21 out of 21
[169:00] Quantification information saved to D:\USERS\Soumyadeep\Cell_Lysates\270625_cell\HA_T3.wiff.quant

[169:01] File #7/36
[169:01] Loading run D:\USERS\Soumyadeep\Cell_Lysates\270625_cell\HI_T1.wiff
[178:24] Pre-processing...
[178:27] 1117 MS1 and 111700 MS2 scans in 1117 (inferred) and 1117 (encoded) cycles, 6606089 precursors in range
[178:29] Calibrating with mass accuracies 28 (MS1), 25 (MS2)
[182:46] RT window set to 2.76918
[182:47] Recommended MS1 mass accuracy setting: 27 ppm
[185:35] Searching decoys
[188:08] Main search
[193:08] Removing low confidence identifications
[193:21] Removing interfering precursors
[193:30] Training neural networks on 25022 target and 13020 decoy PSMs
[194:04] Training neural networks on 25022 target and 12109 decoy PSMs
[194:29] Number of IDs at 0.01 FDR: 14780
[194:30] Precursors at 1% peptidoform FDR: 13845
[194:32] Calculating protein q-values
[194:33] Number of genes identified at 1% FDR: 2737 (precursor-level), 2375 (protein-level) (inference performed using proteotypic peptides only)
[194:33] Quantification
[194:34] Precursors with scored PTMs at 1% FDR: 25 out of 46 considered
[194:34] Precursors with all scored PTM sites unoccupied at 1% FDR: 13820
[194:34] Precursors with PTMs localised (when required) with > 90% confidence: 23 out of 25
[194:35] Quantification information saved to D:\USERS\Soumyadeep\Cell_Lysates\270625_cell\HI_T1.wiff.quant

[194:37] File #8/36
[194:37] Loading run D:\USERS\Soumyadeep\Cell_Lysates\270625_cell\HI_T2.wiff
[204:00] Pre-processing...
[204:03] 1117 MS1 and 111700 MS2 scans in 1117 (inferred) and 1117 (encoded) cycles, 6606089 precursors in range
[204:05] Calibrating with mass accuracies 28 (MS1), 25 (MS2)
[208:30] RT window set to 3.04044
[208:30] Recommended MS1 mass accuracy setting: 26 ppm
[211:45] Searching decoys
[214:38] Main search
[220:20] Removing low confidence identifications
[220:32] Removing interfering precursors
[220:40] Training neural networks on 22816 target and 11083 decoy PSMs
[221:12] Training neural networks on 22816 target and 10364 decoy PSMs
[221:35] Number of IDs at 0.01 FDR: 14426
[221:36] Precursors at 1% peptidoform FDR: 13563
[221:38] Calculating protein q-values
[221:38] Number of genes identified at 1% FDR: 2687 (precursor-level), 2384 (protein-level) (inference performed using proteotypic peptides only)
[221:39] Quantification
[221:39] Precursors with scored PTMs at 1% FDR: 25 out of 54 considered
[221:39] Precursors with all scored PTM sites unoccupied at 1% FDR: 13538
[221:39] Precursors with PTMs localised (when required) with > 90% confidence: 24 out of 25
[221:41] Quantification information saved to D:\USERS\Soumyadeep\Cell_Lysates\270625_cell\HI_T2.wiff.quant

[221:42] File #9/36
[221:42] Loading run D:\USERS\Soumyadeep\Cell_Lysates\270625_cell\HI_T3.wiff
[231:06] Pre-processing...
[231:09] 1117 MS1 and 111700 MS2 scans in 1117 (inferred) and 1117 (encoded) cycles, 6606089 precursors in range
[231:10] Calibrating with mass accuracies 28 (MS1), 25 (MS2)
[235:57] RT window set to 2.55059
[235:58] Recommended MS1 mass accuracy setting: 27 ppm
[238:44] Searching decoys
[241:18] Main search
[246:24] Removing low confidence identifications
[246:36] Removing interfering precursors
[246:44] Training neural networks on 21172 target and 10007 decoy PSMs
[247:15] Training neural networks on 21172 target and 9270 decoy PSMs
[247:37] Number of IDs at 0.01 FDR: 13548
[247:37] Precursors at 1% peptidoform FDR: 12826
[247:39] Calculating protein q-values
[247:40] Number of genes identified at 1% FDR: 2510 (precursor-level), 2206 (protein-level) (inference performed using proteotypic peptides only)
[247:40] Quantification
[247:41] Precursors with scored PTMs at 1% FDR: 16 out of 36 considered
[247:41] Precursors with all scored PTM sites unoccupied at 1% FDR: 12810
[247:41] Precursors with PTMs localised (when required) with > 90% confidence: 13 out of 16
[247:43] Quantification information saved to D:\USERS\Soumyadeep\Cell_Lysates\270625_cell\HI_T3.wiff.quant

[247:44] File #10/36
[247:44] Loading run D:\USERS\Soumyadeep\Cell_Lysates\270625_cell\HIA_T1.wiff
[257:37] Pre-processing...
[257:40] 1117 MS1 and 111700 MS2 scans in 1117 (inferred) and 1117 (encoded) cycles, 6606089 precursors in range
[257:42] Calibrating with mass accuracies 28 (MS1), 25 (MS2)
[262:42] RT window set to 2.95285
[262:43] Recommended MS1 mass accuracy setting: 27 ppm
[266:06] Searching decoys
[269:04] Main search
[274:37] Removing low confidence identifications
[274:50] Removing interfering precursors
[274:58] Training neural networks on 21029 target and 10430 decoy PSMs
[275:28] Training neural networks on 21029 target and 9371 decoy PSMs
[275:49] Number of IDs at 0.01 FDR: 13379
[275:50] Precursors at 1% peptidoform FDR: 12645
[275:52] Calculating protein q-values
[275:52] Number of genes identified at 1% FDR: 2445 (precursor-level), 2224 (protein-level) (inference performed using proteotypic peptides only)
[275:53] Quantification
[275:53] Precursors with scored PTMs at 1% FDR: 21 out of 55 considered
[275:53] Precursors with all scored PTM sites unoccupied at 1% FDR: 12624
[275:53] Precursors with PTMs localised (when required) with > 90% confidence: 21 out of 21
[275:55] Quantification information saved to D:\USERS\Soumyadeep\Cell_Lysates\270625_cell\HIA_T1.wiff.quant

[275:56] File #11/36
[275:56] Loading run D:\USERS\Soumyadeep\Cell_Lysates\270625_cell\HIA_T2.wiff
[285:53] Pre-processing...
[285:56] 1117 MS1 and 111700 MS2 scans in 1117 (inferred) and 1117 (encoded) cycles, 6606089 precursors in range
[285:57] Calibrating with mass accuracies 28 (MS1), 25 (MS2)
[290:53] RT window set to 2.37428
[290:54] Recommended MS1 mass accuracy setting: 24 ppm
[293:42] Searching decoys
[296:26] Main search
[301:57] Removing low confidence identifications
[302:10] Removing interfering precursors
[302:18] Training neural networks on 23137 target and 11970 decoy PSMs
[302:51] Training neural networks on 23137 target and 10946 decoy PSMs
[303:16] Number of IDs at 0.01 FDR: 13758
[303:16] Precursors at 1% peptidoform FDR: 12702
[303:18] Calculating protein q-values
[303:19] Number of genes identified at 1% FDR: 2548 (precursor-level), 2306 (protein-level) (inference performed using proteotypic peptides only)
[303:19] Quantification
[303:20] Precursors with scored PTMs at 1% FDR: 18 out of 60 considered
[303:20] Precursors with all scored PTM sites unoccupied at 1% FDR: 12684
[303:20] Precursors with PTMs localised (when required) with > 90% confidence: 17 out of 18
[303:22] Quantification information saved to D:\USERS\Soumyadeep\Cell_Lysates\270625_cell\HIA_T2.wiff.quant

[303:23] File #12/36
[303:23] Loading run D:\USERS\Soumyadeep\Cell_Lysates\270625_cell\HIA_T3.wiff
[313:31] Pre-processing...
[313:34] 1117 MS1 and 111700 MS2 scans in 1117 (inferred) and 1117 (encoded) cycles, 6606089 precursors in range
[313:36] Calibrating with mass accuracies 28 (MS1), 25 (MS2)
[318:40] RT window set to 2.7618
[318:41] Recommended MS1 mass accuracy setting: 29 ppm
[321:40] Searching decoys
[324:35] Main search
[330:47] Removing low confidence identifications
[331:01] Removing interfering precursors
[331:11] Training neural networks on 24048 target and 12743 decoy PSMs
[331:47] Training neural networks on 24048 target and 11622 decoy PSMs
[332:10] Number of IDs at 0.01 FDR: 14308
[332:11] Precursors at 1% peptidoform FDR: 13201
[332:13] Calculating protein q-values
[332:14] Number of genes identified at 1% FDR: 2632 (precursor-level), 2342 (protein-level) (inference performed using proteotypic peptides only)
[332:14] Quantification
[332:15] Precursors with scored PTMs at 1% FDR: 26 out of 61 considered
[332:15] Precursors with all scored PTM sites unoccupied at 1% FDR: 13175
[332:15] Precursors with PTMs localised (when required) with > 90% confidence: 23 out of 26
[332:17] Quantification information saved to D:\USERS\Soumyadeep\Cell_Lysates\270625_cell\HIA_T3.wiff.quant

[332:18] File #13/36
[332:18] Loading run D:\USERS\Soumyadeep\020725\E2\H_E2_T1.wiff
[345:22] Pre-processing...
[345:26] 1117 MS1 and 111700 MS2 scans in 1117 (inferred) and 1117 (encoded) cycles, 6606089 precursors in range
[345:27] Calibrating with mass accuracies 28 (MS1), 25 (MS2)
[349:22] RT window set to 2.6949
[349:22] Recommended MS1 mass accuracy setting: 29 ppm
[352:01] Searching decoys
[355:41] Main search
[362:46] Removing low confidence identifications
[363:00] Removing interfering precursors
[363:08] Training neural networks on 31550 target and 16477 decoy PSMs
[363:47] Training neural networks on 31550 target and 15349 decoy PSMs
[364:18] Number of IDs at 0.01 FDR: 18945
[364:19] Precursors at 1% peptidoform FDR: 17856
[364:21] Calculating protein q-values
[364:22] Number of genes identified at 1% FDR: 3059 (precursor-level), 2661 (protein-level) (inference performed using proteotypic peptides only)
[364:22] Quantification
[364:23] Precursors with scored PTMs at 1% FDR: 37 out of 75 considered
[364:23] Precursors with all scored PTM sites unoccupied at 1% FDR: 17819
[364:23] Precursors with PTMs localised (when required) with > 90% confidence: 34 out of 37
[364:25] Quantification information saved to D:\USERS\Soumyadeep\020725\E2\H_E2_T1.wiff.quant

[364:26] File #14/36
[364:26] Loading run D:\USERS\Soumyadeep\020725\E2\H_E2_T2.wiff
[374:20] Pre-processing...
[374:23] 1117 MS1 and 111700 MS2 scans in 1117 (inferred) and 1117 (encoded) cycles, 6606089 precursors in range
[374:25] Calibrating with mass accuracies 28 (MS1), 25 (MS2)
[378:47] RT window set to 2.54833
[378:47] Recommended MS1 mass accuracy setting: 28 ppm
[381:31] Searching decoys
[385:08] Main search
[392:10] Removing low confidence identifications
[392:24] Removing interfering precursors
[392:33] Training neural networks on 32393 target and 17039 decoy PSMs
[393:12] Training neural networks on 32393 target and 15855 decoy PSMs
[393:44] Number of IDs at 0.01 FDR: 19030
[393:45] Precursors at 1% peptidoform FDR: 18036
[393:47] Calculating protein q-values
[393:48] Number of genes identified at 1% FDR: 2995 (precursor-level), 2701 (protein-level) (inference performed using proteotypic peptides only)
[393:48] Quantification
[393:49] Precursors with scored PTMs at 1% FDR: 39 out of 80 considered
[393:49] Precursors with all scored PTM sites unoccupied at 1% FDR: 17997
[393:49] Precursors with PTMs localised (when required) with > 90% confidence: 38 out of 39
[393:50] Quantification information saved to D:\USERS\Soumyadeep\020725\E2\H_E2_T2.wiff.quant

[393:52] File #15/36
[393:52] Loading run D:\USERS\Soumyadeep\020725\E2\H_E2_T3.wiff
[403:33] Pre-processing...
[403:37] 1117 MS1 and 111700 MS2 scans in 1117 (inferred) and 1117 (encoded) cycles, 6606089 precursors in range
[403:38] Calibrating with mass accuracies 28 (MS1), 25 (MS2)
[408:00] RT window set to 2.5512
[408:01] Recommended MS1 mass accuracy setting: 28 ppm
[410:45] Searching decoys
[414:20] Main search
[421:25] Removing low confidence identifications
[421:40] Removing interfering precursors
[421:48] Training neural networks on 35227 target and 18779 decoy PSMs
[422:29] Training neural networks on 35227 target and 17544 decoy PSMs
[423:03] Number of IDs at 0.01 FDR: 20119
[423:04] Precursors at 1% peptidoform FDR: 18999
[423:06] Calculating protein q-values
[423:07] Number of genes identified at 1% FDR: 3130 (precursor-level), 2710 (protein-level) (inference performed using proteotypic peptides only)
[423:07] Quantification
[423:07] Precursors with scored PTMs at 1% FDR: 19 out of 56 considered
[423:07] Precursors with all scored PTM sites unoccupied at 1% FDR: 18980
[423:07] Precursors with PTMs localised (when required) with > 90% confidence: 19 out of 19
[423:09] Quantification information saved to D:\USERS\Soumyadeep\020725\E2\H_E2_T3.wiff.quant

[423:11] File #16/36
[423:11] Loading run D:\USERS\Soumyadeep\020725\E2\HA_E2_T1.wiff
[431:01] Pre-processing...
[431:04] 1117 MS1 and 111700 MS2 scans in 1117 (inferred) and 1117 (encoded) cycles, 6606089 precursors in range
[431:06] Calibrating with mass accuracies 28 (MS1), 25 (MS2)
[435:24] RT window set to 2.46977
[435:24] Recommended MS1 mass accuracy setting: 24 ppm
[437:37] Searching decoys
[439:51] Main search
[444:16] Removing low confidence identifications
[444:28] Removing interfering precursors
[444:36] Training neural networks on 25423 target and 12054 decoy PSMs
[445:09] Training neural networks on 25423 target and 11564 decoy PSMs
[445:33] Number of IDs at 0.01 FDR: 15272
[445:33] Precursors at 1% peptidoform FDR: 14230
[445:35] Calculating protein q-values
[445:36] Number of genes identified at 1% FDR: 2518 (precursor-level), 2151 (protein-level) (inference performed using proteotypic peptides only)
[445:36] Quantification
[445:37] Precursors with scored PTMs at 1% FDR: 22 out of 36 considered
[445:37] Precursors with all scored PTM sites unoccupied at 1% FDR: 14208
[445:37] Precursors with PTMs localised (when required) with > 90% confidence: 21 out of 22
[445:38] Quantification information saved to D:\USERS\Soumyadeep\020725\E2\HA_E2_T1.wiff.quant

[445:39] File #17/36
[445:39] Loading run D:\USERS\Soumyadeep\020725\E2\HA_E2_T2.wiff
[454:11] Pre-processing...
[454:14] 1117 MS1 and 111700 MS2 scans in 1117 (inferred) and 1117 (encoded) cycles, 6606089 precursors in range
[454:16] Calibrating with mass accuracies 28 (MS1), 25 (MS2)
[459:00] RT window set to 2.79755
[459:00] Recommended MS1 mass accuracy setting: 28 ppm
[461:21] Searching decoys
[463:56] Main search
[469:05] Removing low confidence identifications
[469:17] Removing interfering precursors
[469:26] Training neural networks on 23868 target and 11049 decoy PSMs
[469:56] Training neural networks on 23868 target and 10143 decoy PSMs
[470:19] Number of IDs at 0.01 FDR: 15343
[470:19] Precursors at 1% peptidoform FDR: 14600
[470:22] Calculating protein q-values
[470:22] Number of genes identified at 1% FDR: 2519 (precursor-level), 2164 (protein-level) (inference performed using proteotypic peptides only)
[470:22] Quantification
[470:23] Precursors with scored PTMs at 1% FDR: 31 out of 45 considered
[470:23] Precursors with all scored PTM sites unoccupied at 1% FDR: 14569
[470:23] Precursors with PTMs localised (when required) with > 90% confidence: 26 out of 31
[470:25] Quantification information saved to D:\USERS\Soumyadeep\020725\E2\HA_E2_T2.wiff.quant

[470:25] File #18/36
[470:25] Loading run D:\USERS\Soumyadeep\020725\E2\HA_E2_T3.wiff
[479:01] Pre-processing...
[479:04] 1117 MS1 and 111700 MS2 scans in 1117 (inferred) and 1117 (encoded) cycles, 6606089 precursors in range
[479:06] Calibrating with mass accuracies 28 (MS1), 25 (MS2)
[483:56] RT window set to 2.75578
[483:56] Recommended MS1 mass accuracy setting: 27 ppm
[486:19] Searching decoys
[488:50] Main search
[493:53] Removing low confidence identifications
[494:06] Removing interfering precursors
[494:14] Training neural networks on 24296 target and 11299 decoy PSMs
[494:46] Training neural networks on 24296 target and 10753 decoy PSMs
[495:10] Number of IDs at 0.01 FDR: 15242
[495:10] Precursors at 1% peptidoform FDR: 14481
[495:12] Calculating protein q-values
[495:13] Number of genes identified at 1% FDR: 2491 (precursor-level), 2157 (protein-level) (inference performed using proteotypic peptides only)
[495:13] Quantification
[495:14] Precursors with scored PTMs at 1% FDR: 26 out of 40 considered
[495:14] Precursors with all scored PTM sites unoccupied at 1% FDR: 14455
[495:14] Precursors with PTMs localised (when required) with > 90% confidence: 24 out of 26
[495:15] Quantification information saved to D:\USERS\Soumyadeep\020725\E2\HA_E2_T3.wiff.quant

[495:16] File #19/36
[495:16] Loading run D:\USERS\Soumyadeep\020725\E2\HI_E2_T1.wiff
[503:00] Pre-processing...
[503:03] 1117 MS1 and 111700 MS2 scans in 1117 (inferred) and 1117 (encoded) cycles, 6606089 precursors in range
[503:05] Calibrating with mass accuracies 28 (MS1), 25 (MS2)
[507:36] RT window set to 2.55076
[507:36] Recommended MS1 mass accuracy setting: 25 ppm
[510:09] Searching decoys
[512:19] Main search
[516:38] Removing low confidence identifications
[516:51] Removing interfering precursors
[516:59] Training neural networks on 24154 target and 11013 decoy PSMs
[517:30] Training neural networks on 24154 target and 10293 decoy PSMs
[517:53] Number of IDs at 0.01 FDR: 15033
[517:54] Precursors at 1% peptidoform FDR: 13983
[517:56] Calculating protein q-values
[517:57] Number of genes identified at 1% FDR: 2598 (precursor-level), 2189 (protein-level) (inference performed using proteotypic peptides only)
[517:57] Quantification
[517:57] Precursors with scored PTMs at 1% FDR: 19 out of 40 considered
[517:57] Precursors with all scored PTM sites unoccupied at 1% FDR: 13964
[517:57] Precursors with PTMs localised (when required) with > 90% confidence: 18 out of 19
[517:59] Quantification information saved to D:\USERS\Soumyadeep\020725\E2\HI_E2_T1.wiff.quant

[518:00] File #20/36
[518:00] Loading run D:\USERS\Soumyadeep\020725\E2\HI_E2_T2.wiff
[526:32] Pre-processing...
[526:35] 1117 MS1 and 111700 MS2 scans in 1117 (inferred) and 1117 (encoded) cycles, 6606089 precursors in range
[526:37] Calibrating with mass accuracies 28 (MS1), 25 (MS2)
[531:18] RT window set to 2.53447
[531:19] Recommended MS1 mass accuracy setting: 26 ppm
[533:45] Searching decoys
[535:50] Main search
[539:54] Removing low confidence identifications
[540:06] Removing interfering precursors
[540:14] Training neural networks on 21740 target and 9794 decoy PSMs
[540:43] Training neural networks on 21740 target and 9065 decoy PSMs
[541:04] Number of IDs at 0.01 FDR: 14074
[541:05] Precursors at 1% peptidoform FDR: 13384
[541:08] Calculating protein q-values
[541:09] Number of genes identified at 1% FDR: 2408 (precursor-level), 2105 (protein-level) (inference performed using proteotypic peptides only)
[541:10] Quantification
[541:10] Precursors with scored PTMs at 1% FDR: 19 out of 37 considered
[541:10] Precursors with all scored PTM sites unoccupied at 1% FDR: 13365
[541:10] Precursors with PTMs localised (when required) with > 90% confidence: 19 out of 19
[541:12] Quantification information saved to D:\USERS\Soumyadeep\020725\E2\HI_E2_T2.wiff.quant

[541:13] File #21/36
[541:13] Loading run D:\USERS\Soumyadeep\020725\E2\HI_E2_T3.wiff
[549:50] Pre-processing...
[549:53] 1117 MS1 and 111700 MS2 scans in 1117 (inferred) and 1117 (encoded) cycles, 6606089 precursors in range
[549:55] Calibrating with mass accuracies 28 (MS1), 25 (MS2)
[554:41] RT window set to 2.82766
[554:42] Recommended MS1 mass accuracy setting: 27 ppm
[557:28] Searching decoys
[559:52] Main search
[564:32] Removing low confidence identifications
[564:44] Removing interfering precursors
[564:52] Training neural networks on 22777 target and 10656 decoy PSMs
[565:22] Training neural networks on 22777 target and 9826 decoy PSMs
[565:43] Number of IDs at 0.01 FDR: 14623
[565:44] Precursors at 1% peptidoform FDR: 13750
[565:46] Calculating protein q-values
[565:47] Number of genes identified at 1% FDR: 2498 (precursor-level), 2162 (protein-level) (inference performed using proteotypic peptides only)
[565:47] Quantification
[565:47] Precursors with scored PTMs at 1% FDR: 21 out of 44 considered
[565:47] Precursors with all scored PTM sites unoccupied at 1% FDR: 13729
[565:47] Precursors with PTMs localised (when required) with > 90% confidence: 21 out of 21
[565:49] Quantification information saved to D:\USERS\Soumyadeep\020725\E2\HI_E2_T3.wiff.quant

[565:50] File #22/36
[565:50] Loading run D:\USERS\Soumyadeep\020725\E2\HIA_E2_T1.wiff
[574:57] Pre-processing...
[575:05] 1117 MS1 and 111700 MS2 scans in 1117 (inferred) and 1117 (encoded) cycles, 6606089 precursors in range
[575:06] Calibrating with mass accuracies 28 (MS1), 25 (MS2)
[580:01] RT window set to 2.60816
[580:02] Recommended MS1 mass accuracy setting: 26 ppm
[582:17] Searching decoys
[584:39] Main search
[589:18] Removing low confidence identifications
[589:31] Removing interfering precursors
[589:39] Training neural networks on 24484 target and 11334 decoy PSMs
[590:11] Training neural networks on 24484 target and 10679 decoy PSMs
[590:34] Number of IDs at 0.01 FDR: 15318
[590:35] Precursors at 1% peptidoform FDR: 14264
[590:37] Calculating protein q-values
[590:38] Number of genes identified at 1% FDR: 2572 (precursor-level), 2166 (protein-level) (inference performed using proteotypic peptides only)
[590:38] Quantification
[590:39] Precursors with scored PTMs at 1% FDR: 28 out of 53 considered
[590:39] Precursors with all scored PTM sites unoccupied at 1% FDR: 14236
[590:39] Precursors with PTMs localised (when required) with > 90% confidence: 27 out of 28
[590:40] Quantification information saved to D:\USERS\Soumyadeep\020725\E2\HIA_E2_T1.wiff.quant

[590:41] File #23/36
[590:41] Loading run D:\USERS\Soumyadeep\020725\E2\HIA_E2_T2.wiff
[600:01] Pre-processing...
[600:04] 1117 MS1 and 111700 MS2 scans in 1117 (inferred) and 1117 (encoded) cycles, 6606089 precursors in range
[600:06] Calibrating with mass accuracies 28 (MS1), 25 (MS2)
[605:10] RT window set to 2.76785
[605:10] Recommended MS1 mass accuracy setting: 27 ppm
[608:04] Searching decoys
[610:34] Main search
[615:26] Removing low confidence identifications
[615:39] Removing interfering precursors
[615:47] Training neural networks on 26950 target and 13113 decoy PSMs
[616:20] Training neural networks on 26950 target and 12436 decoy PSMs
[616:45] Number of IDs at 0.01 FDR: 15564
[616:46] Precursors at 1% peptidoform FDR: 14686
[616:48] Calculating protein q-values
[616:49] Number of genes identified at 1% FDR: 2606 (precursor-level), 2287 (protein-level) (inference performed using proteotypic peptides only)
[616:49] Quantification
[616:50] Precursors with scored PTMs at 1% FDR: 31 out of 51 considered
[616:50] Precursors with all scored PTM sites unoccupied at 1% FDR: 14655
[616:50] Precursors with PTMs localised (when required) with > 90% confidence: 28 out of 31
[616:51] Quantification information saved to D:\USERS\Soumyadeep\020725\E2\HIA_E2_T2.wiff.quant

[616:52] File #24/36
[616:52] Loading run D:\USERS\Soumyadeep\020725\E2\HIA_E2_T3.wiff
[626:19] Pre-processing...
[626:22] 1117 MS1 and 111700 MS2 scans in 1117 (inferred) and 1117 (encoded) cycles, 6606089 precursors in range
[626:23] Calibrating with mass accuracies 28 (MS1), 25 (MS2)
[631:26] RT window set to 2.55737
[631:27] Recommended MS1 mass accuracy setting: 26 ppm
[633:42] Searching decoys
[636:04] Main search
[640:45] Removing low confidence identifications
[640:57] Removing interfering precursors
[641:05] Training neural networks on 22519 target and 10103 decoy PSMs
[641:35] Training neural networks on 22519 target and 9472 decoy PSMs
[641:57] Number of IDs at 0.01 FDR: 14323
[641:58] Precursors at 1% peptidoform FDR: 13488
[642:00] Calculating protein q-values
[642:00] Number of genes identified at 1% FDR: 2407 (precursor-level), 2112 (protein-level) (inference performed using proteotypic peptides only)
[642:01] Quantification
[642:01] Precursors with scored PTMs at 1% FDR: 27 out of 46 considered
[642:01] Precursors with all scored PTM sites unoccupied at 1% FDR: 13461
[642:01] Precursors with PTMs localised (when required) with > 90% confidence: 24 out of 27
[642:03] Quantification information saved to D:\USERS\Soumyadeep\020725\E2\HIA_E2_T3.wiff.quant

[642:04] File #25/36
[642:04] Loading run D:\USERS\Soumyadeep\020725\E3\H_E3_T1.wiff
[652:54] Pre-processing...
[652:57] 1117 MS1 and 111700 MS2 scans in 1117 (inferred) and 1117 (encoded) cycles, 6606089 precursors in range
[652:59] Calibrating with mass accuracies 28 (MS1), 25 (MS2)
[656:16] RT window set to 2.1887
[656:16] Recommended MS1 mass accuracy setting: 31 ppm
[658:04] Searching decoys
[661:41] Main search
[668:53] Removing low confidence identifications
[669:10] Removing interfering precursors
[669:19] Training neural networks on 55159 target and 29562 decoy PSMs
[670:19] Training neural networks on 55159 target and 28146 decoy PSMs
[671:11] Number of IDs at 0.01 FDR: 28908
[671:12] Precursors at 1% peptidoform FDR: 27165
[671:14] Calculating protein q-values
[671:15] Number of genes identified at 1% FDR: 3675 (precursor-level), 3220 (protein-level) (inference performed using proteotypic peptides only)
[671:15] Quantification
[671:16] Precursors with scored PTMs at 1% FDR: 49 out of 72 considered
[671:16] Precursors with all scored PTM sites unoccupied at 1% FDR: 27116
[671:16] Precursors with PTMs localised (when required) with > 90% confidence: 48 out of 49
[671:18] Quantification information saved to D:\USERS\Soumyadeep\020725\E3\H_E3_T1.wiff.quant

[671:20] File #26/36
[671:20] Loading run D:\USERS\Soumyadeep\020725\E3\H_E3_T2.wiff
[682:47] Pre-processing...
[682:50] 1117 MS1 and 111700 MS2 scans in 1117 (inferred) and 1117 (encoded) cycles, 6606089 precursors in range
[682:52] Calibrating with mass accuracies 28 (MS1), 25 (MS2)
[686:26] RT window set to 2.67034
[686:27] Recommended MS1 mass accuracy setting: 32 ppm
[688:34] Searching decoys
[693:09] Main search
[702:09] Removing low confidence identifications
[702:26] Removing interfering precursors
[702:35] Training neural networks on 45596 target and 21885 decoy PSMs
[703:26] Training neural networks on 45596 target and 20785 decoy PSMs
[704:09] Number of IDs at 0.01 FDR: 27467
[704:10] Precursors at 1% peptidoform FDR: 26038
[704:12] Calculating protein q-values
[704:13] Number of genes identified at 1% FDR: 3566 (precursor-level), 2975 (protein-level) (inference performed using proteotypic peptides only)
[704:13] Quantification
[704:14] Precursors with scored PTMs at 1% FDR: 46 out of 74 considered
[704:14] Precursors with all scored PTM sites unoccupied at 1% FDR: 25992
[704:14] Precursors with PTMs localised (when required) with > 90% confidence: 44 out of 46
[704:16] Quantification information saved to D:\USERS\Soumyadeep\020725\E3\H_E3_T2.wiff.quant

[704:18] File #27/36
[704:18] Loading run D:\USERS\Soumyadeep\020725\E3\H_E3_T3.wiff
[715:20] Pre-processing...
[715:23] 1117 MS1 and 111700 MS2 scans in 1117 (inferred) and 1117 (encoded) cycles, 6606089 precursors in range
[715:25] Calibrating with mass accuracies 28 (MS1), 25 (MS2)
[718:45] RT window set to 2.04882
[718:46] Recommended MS1 mass accuracy setting: 30 ppm
[720:14] Searching decoys
[723:54] Main search
[731:13] Removing low confidence identifications
[731:29] Removing interfering precursors
[731:38] Training neural networks on 50081 target and 25195 decoy PSMs
[732:34] Training neural networks on 50081 target and 23936 decoy PSMs
[733:21] Number of IDs at 0.01 FDR: 28438
[733:22] Precursors at 1% peptidoform FDR: 26862
[733:24] Calculating protein q-values
[733:25] Number of genes identified at 1% FDR: 3618 (precursor-level), 3154 (protein-level) (inference performed using proteotypic peptides only)
[733:25] Quantification
[733:26] Precursors with scored PTMs at 1% FDR: 34 out of 59 considered
[733:26] Precursors with all scored PTM sites unoccupied at 1% FDR: 26828
[733:26] Precursors with PTMs localised (when required) with > 90% confidence: 30 out of 34
[733:28] Quantification information saved to D:\USERS\Soumyadeep\020725\E3\H_E3_T3.wiff.quant

[733:30] File #28/36
[733:30] Loading run D:\USERS\Soumyadeep\020725\E3\HA_E3_T1.wiff
[744:09] Pre-processing...
[744:13] 1117 MS1 and 111700 MS2 scans in 1117 (inferred) and 1117 (encoded) cycles, 6606089 precursors in range
[744:14] Calibrating with mass accuracies 28 (MS1), 25 (MS2)
[748:21] RT window set to 2.20413
[748:22] Recommended MS1 mass accuracy setting: 29 ppm
[749:59] Searching decoys
[753:38] Main search
[761:34] Removing low confidence identifications
[761:51] Removing interfering precursors
[762:04] Training neural networks on 48974 target and 25928 decoy PSMs
[762:58] Training neural networks on 48974 target and 24771 decoy PSMs
[763:44] Number of IDs at 0.01 FDR: 26018
[763:44] Precursors at 1% peptidoform FDR: 24687
[763:47] Calculating protein q-values
[763:47] Number of genes identified at 1% FDR: 3456 (precursor-level), 3024 (protein-level) (inference performed using proteotypic peptides only)
[763:47] Quantification
[763:48] Precursors with scored PTMs at 1% FDR: 34 out of 51 considered
[763:48] Precursors with all scored PTM sites unoccupied at 1% FDR: 24653
[763:48] Precursors with PTMs localised (when required) with > 90% confidence: 34 out of 34
[763:50] Quantification information saved to D:\USERS\Soumyadeep\020725\E3\HA_E3_T1.wiff.quant

[763:54] File #29/36
[763:54] Loading run D:\USERS\Soumyadeep\020725\E3\HA_E3_T2.wiff
[774:59] Pre-processing...
[775:05] 1117 MS1 and 111700 MS2 scans in 1117 (inferred) and 1117 (encoded) cycles, 6606089 precursors in range
[775:07] Calibrating with mass accuracies 28 (MS1), 25 (MS2)
[778:27] RT window set to 2.03886
[778:28] Recommended MS1 mass accuracy setting: 28 ppm
[780:06] Searching decoys
[783:13] Main search
[789:14] Removing low confidence identifications
[789:29] Removing interfering precursors
[789:38] Training neural networks on 42249 target and 20649 decoy PSMs
[790:24] Training neural networks on 42249 target and 19476 decoy PSMs
[791:03] Number of IDs at 0.01 FDR: 24768
[791:04] Precursors at 1% peptidoform FDR: 23478
[791:06] Calculating protein q-values
[791:07] Number of genes identified at 1% FDR: 3362 (precursor-level), 2922 (protein-level) (inference performed using proteotypic peptides only)
[791:07] Quantification
[791:08] Precursors with scored PTMs at 1% FDR: 26 out of 55 considered
[791:08] Precursors with all scored PTM sites unoccupied at 1% FDR: 23452
[791:08] Precursors with PTMs localised (when required) with > 90% confidence: 24 out of 26
[791:09] Quantification information saved to D:\USERS\Soumyadeep\020725\E3\HA_E3_T2.wiff.quant

[791:11] File #30/36
[791:11] Loading run D:\USERS\Soumyadeep\020725\E3\HA_E3_T3.wiff
[801:59] Pre-processing...
[802:03] 1117 MS1 and 111700 MS2 scans in 1117 (inferred) and 1117 (encoded) cycles, 6606089 precursors in range
[802:04] Calibrating with mass accuracies 28 (MS1), 25 (MS2)
[805:27] RT window set to 2.03146
[805:28] Recommended MS1 mass accuracy setting: 31 ppm
[807:12] Searching decoys
[810:27] Main search
[816:54] Removing low confidence identifications
[817:09] Removing interfering precursors
[817:18] Training neural networks on 46574 target and 23600 decoy PSMs
[818:09] Training neural networks on 46574 target and 22534 decoy PSMs
[818:53] Number of IDs at 0.01 FDR: 25646
[818:53] Precursors at 1% peptidoform FDR: 23985
[818:55] Calculating protein q-values
[818:56] Number of genes identified at 1% FDR: 3432 (precursor-level), 2931 (protein-level) (inference performed using proteotypic peptides only)
[818:56] Quantification
[818:57] Precursors with scored PTMs at 1% FDR: 18 out of 46 considered
[818:57] Precursors with all scored PTM sites unoccupied at 1% FDR: 23967
[818:57] Precursors with PTMs localised (when required) with > 90% confidence: 18 out of 18
[818:59] Quantification information saved to D:\USERS\Soumyadeep\020725\E3\HA_E3_T3.wiff.quant

[819:01] File #31/36
[819:01] Loading run D:\USERS\Soumyadeep\020725\E3\HI_E3_T1.wiff
[833:25] Pre-processing...
[833:28] 1117 MS1 and 111700 MS2 scans in 1117 (inferred) and 1117 (encoded) cycles, 6606089 precursors in range
[833:30] Calibrating with mass accuracies 28 (MS1), 25 (MS2)
[836:47] RT window set to 2.11822
[836:48] Recommended MS1 mass accuracy setting: 30 ppm
[838:23] Searching decoys
[841:36] Main search
[847:56] Removing low confidence identifications
[848:10] Removing interfering precursors
[848:19] Training neural networks on 44651 target and 22310 decoy PSMs
[849:07] Training neural networks on 44651 target and 21249 decoy PSMs
[849:46] Number of IDs at 0.01 FDR: 25384
[849:47] Precursors at 1% peptidoform FDR: 24347
[849:49] Calculating protein q-values
[849:50] Number of genes identified at 1% FDR: 3452 (precursor-level), 3050 (protein-level) (inference performed using proteotypic peptides only)
[849:50] Quantification
[849:51] Precursors with scored PTMs at 1% FDR: 45 out of 66 considered
[849:51] Precursors with all scored PTM sites unoccupied at 1% FDR: 24302
[849:51] Precursors with PTMs localised (when required) with > 90% confidence: 45 out of 45
[849:52] Quantification information saved to D:\USERS\Soumyadeep\020725\E3\HI_E3_T1.wiff.quant

[849:54] File #32/36
[849:54] Loading run D:\USERS\Soumyadeep\020725\E3\HI_E3_T2.wiff
[860:46] Pre-processing...
[860:49] 1117 MS1 and 111700 MS2 scans in 1117 (inferred) and 1117 (encoded) cycles, 6606089 precursors in range
[860:51] Calibrating with mass accuracies 28 (MS1), 25 (MS2)
[864:40] RT window set to 2.2544
[864:40] Recommended MS1 mass accuracy setting: 32 ppm
[866:19] Searching decoys
[869:51] Main search
[876:40] Removing low confidence identifications
[876:55] Removing interfering precursors
[877:04] Training neural networks on 43862 target and 21630 decoy PSMs
[877:52] Training neural networks on 43862 target and 20278 decoy PSMs
[878:32] Number of IDs at 0.01 FDR: 25590
[878:32] Precursors at 1% peptidoform FDR: 24243
[878:34] Calculating protein q-values
[878:35] Number of genes identified at 1% FDR: 3430 (precursor-level), 2859 (protein-level) (inference performed using proteotypic peptides only)
[878:35] Quantification
[878:36] Precursors with scored PTMs at 1% FDR: 37 out of 72 considered
[878:36] Precursors with all scored PTM sites unoccupied at 1% FDR: 24206
[878:36] Precursors with PTMs localised (when required) with > 90% confidence: 36 out of 37
[878:38] Quantification information saved to D:\USERS\Soumyadeep\020725\E3\HI_E3_T2.wiff.quant

[878:40] File #33/36
[878:40] Loading run D:\USERS\Soumyadeep\020725\E3\HI_E3_T3.wiff
[889:40] Pre-processing...
[889:43] 1117 MS1 and 111700 MS2 scans in 1117 (inferred) and 1117 (encoded) cycles, 6606089 precursors in range
[889:45] Calibrating with mass accuracies 28 (MS1), 25 (MS2)
[893:51] RT window set to 2.02602
[893:52] Recommended MS1 mass accuracy setting: 30 ppm
[895:29] Searching decoys
[898:38] Main search
[904:50] Removing low confidence identifications
[905:05] Removing interfering precursors
[905:14] Training neural networks on 49325 target and 26120 decoy PSMs
[906:09] Training neural networks on 49325 target and 24772 decoy PSMs
[906:53] Number of IDs at 0.01 FDR: 26021
[906:54] Precursors at 1% peptidoform FDR: 24835
[906:56] Calculating protein q-values
[906:56] Number of genes identified at 1% FDR: 3480 (precursor-level), 3003 (protein-level) (inference performed using proteotypic peptides only)
[906:56] Quantification
[906:57] Precursors with scored PTMs at 1% FDR: 40 out of 56 considered
[906:57] Precursors with all scored PTM sites unoccupied at 1% FDR: 24795
[906:57] Precursors with PTMs localised (when required) with > 90% confidence: 38 out of 40
[906:59] Quantification information saved to D:\USERS\Soumyadeep\020725\E3\HI_E3_T3.wiff.quant

[907:01] File #34/36
[907:01] Loading run D:\USERS\Soumyadeep\020725\E3\HIA_E3_T1.wiff
[919:14] Pre-processing...
[919:17] 1117 MS1 and 111700 MS2 scans in 1117 (inferred) and 1117 (encoded) cycles, 6606089 precursors in range
[919:19] Calibrating with mass accuracies 28 (MS1), 25 (MS2)
[922:55] RT window set to 2.1864
[922:55] Recommended MS1 mass accuracy setting: 31 ppm
[925:08] Searching decoys
[929:29] Main search
[937:59] Removing low confidence identifications
[938:15] Removing interfering precursors
[938:23] Training neural networks on 46295 target and 23417 decoy PSMs
[939:17] Training neural networks on 46295 target and 21999 decoy PSMs
[940:01] Number of IDs at 0.01 FDR: 26865
[940:01] Precursors at 1% peptidoform FDR: 25653
[940:03] Calculating protein q-values
[940:04] Number of genes identified at 1% FDR: 3539 (precursor-level), 2995 (protein-level) (inference performed using proteotypic peptides only)
[940:04] Quantification
[940:05] Precursors with scored PTMs at 1% FDR: 29 out of 62 considered
[940:05] Precursors with all scored PTM sites unoccupied at 1% FDR: 25624
[940:05] Precursors with PTMs localised (when required) with > 90% confidence: 29 out of 29
[940:07] Quantification information saved to D:\USERS\Soumyadeep\020725\E3\HIA_E3_T1.wiff.quant

[940:11] File #35/36
[940:11] Loading run D:\USERS\Soumyadeep\020725\E3\HIA_E3_T2.wiff
[952:49] Pre-processing...
[952:53] 1117 MS1 and 111700 MS2 scans in 1117 (inferred) and 1117 (encoded) cycles, 6606089 precursors in range
[952:54] Calibrating with mass accuracies 28 (MS1), 25 (MS2)
[956:31] RT window set to 2.30134
[956:32] Recommended MS1 mass accuracy setting: 32 ppm
[958:35] Searching decoys
[963:14] Main search
[972:25] Removing low confidence identifications
[972:41] Removing interfering precursors
[972:50] Training neural networks on 48804 target and 24990 decoy PSMs
[973:46] Training neural networks on 48804 target and 23757 decoy PSMs
[974:34] Number of IDs at 0.01 FDR: 27613
[974:34] Precursors at 1% peptidoform FDR: 26194
[974:36] Calculating protein q-values
[974:37] Number of genes identified at 1% FDR: 3614 (precursor-level), 3092 (protein-level) (inference performed using proteotypic peptides only)
[974:37] Quantification
[974:39] Precursors with scored PTMs at 1% FDR: 44 out of 76 considered
[974:39] Precursors with all scored PTM sites unoccupied at 1% FDR: 26150
[974:39] Precursors with PTMs localised (when required) with > 90% confidence: 42 out of 44
[974:41] Quantification information saved to D:\USERS\Soumyadeep\020725\E3\HIA_E3_T2.wiff.quant

[974:44] File #36/36
[974:44] Loading run D:\USERS\Soumyadeep\020725\E3\HIA_E3_T3.wiff
[987:46] Pre-processing...
[987:49] 1117 MS1 and 111700 MS2 scans in 1117 (inferred) and 1117 (encoded) cycles, 6606089 precursors in range
[987:51] Calibrating with mass accuracies 28 (MS1), 25 (MS2)
[992:08] RT window set to 2.32892
[992:08] Recommended MS1 mass accuracy setting: 31 ppm
[994:02] Searching decoys
[997:51] Main search
[1005:11] Removing low confidence identifications
[1005:26] Removing interfering precursors
[1005:35] Training neural networks on 43430 target and 21350 decoy PSMs
[1006:22] Training neural networks on 43430 target and 20093 decoy PSMs
[1007:01] Number of IDs at 0.01 FDR: 25570
[1007:01] Precursors at 1% peptidoform FDR: 24189
[1007:03] Calculating protein q-values
[1007:04] Number of genes identified at 1% FDR: 3402 (precursor-level), 2839 (protein-level) (inference performed using proteotypic peptides only)
[1007:04] Quantification
[1007:05] Precursors with scored PTMs at 1% FDR: 56 out of 85 considered
[1007:05] Precursors with all scored PTM sites unoccupied at 1% FDR: 24133
[1007:05] Precursors with PTMs localised (when required) with > 90% confidence: 55 out of 56
[1007:07] Quantification information saved to D:\USERS\Soumyadeep\020725\E3\HIA_E3_T3.wiff.quant

[1007:10] Cross-run analysis
[1007:10] Reading quantification information: 36 files
[1007:54] Quantifying peptides
[1010:38] Assembling protein groups
[1010:41] Quantifying proteins
[1010:42] Calculating q-values for protein and gene groups
[1011:10] Calculating global q-values for protein and gene groups
[1011:10] Protein groups with global q-value <= 0.01: 3820
[1011:17] Compressed report saved to D:\USERS\Soumyadeep\CellLysate_DIANN2\HvHIvHA_out\all_celllysates_comparison-first-pass.parquet. Use R 'arrow' or Python 'PyArrow' package to process
[1011:17] Site report saved to D:\USERS\Soumyadeep\CellLysate_DIANN2\HvHIvHA_out\all_celllysates_comparison-first-pass.site_report.parquet
[1011:17] Saving precursor levels matrix
[1011:18] Precursor levels matrix (1% precursor and protein group FDR) saved to D:\USERS\Soumyadeep\CellLysate_DIANN2\HvHIvHA_out\all_celllysates_comparison-first-pass.pr_matrix.tsv.
[1011:18] Manifest saved to D:\USERS\Soumyadeep\CellLysate_DIANN2\HvHIvHA_out\all_celllysates_comparison-first-pass.manifest.txt
[1011:18] Stats report saved to D:\USERS\Soumyadeep\CellLysate_DIANN2\HvHIvHA_out\all_celllysates_comparison-first-pass.stats.tsv
[1011:19] Generating spectral library:
[1011:21] 46809 target and 2365 decoy precursors saved
[1011:21] Spectral library saved to D:\USERS\Soumyadeep\CellLysate_DIANN2\HvHIvHA_out\all_celllysates_comparison-lib.parquet

[1011:25] Loading spectral library D:\USERS\Soumyadeep\CellLysate_DIANN2\HvHIvHA_out\all_celllysates_comparison-lib.parquet
[1011:27] Spectral library loaded: 7488 protein isoforms, 7278 protein groups and 49174 precursors in 43315 elution groups.
[1011:27] Loading protein annotations from FASTA camprotR_240512_cRAP_20190401_full_tags.fasta
[1011:27] Loading protein annotations from FASTA D:\USERS\Soumyadeep\Fasta 070725\clean_humwholeprot_reviewed_can.fasta
[1011:27] Annotating library proteins with information from the FASTA database
[1011:27] Gene names missing for some isoforms
[1011:27] Library contains 7488 proteins, and 7470 genes
[1011:28] Initialising library
[1011:33] Saving the library to D:\USERS\Soumyadeep\CellLysate_DIANN2\HvHIvHA_out\all_celllysates_comparison-lib.parquet.skyline.speclib


Second pass: using the newly created spectral library to reanalyse the data

[1011:33] File #1/36
[1011:33] Loading run D:\USERS\Soumyadeep\Cell_Lysates\270625_cell\H_T1.wiff
[1020:49] Pre-processing...
[1020:49] 1117 MS1 and 111700 MS2 scans in 1117 (inferred) and 1117 (encoded) cycles, 46809 precursors in range
[1020:49] Calibrating with mass accuracies 28 (MS1), 25 (MS2)
[1020:51] RT window set to 0.676807
[1020:51] Recommended MS1 mass accuracy setting: 34 ppm
[1020:52] Searching decoys
[1020:53] Main search
[1020:54] Removing low confidence identifications
[1020:56] Removing interfering precursors
[1020:57] Training neural networks on 26556 target and 12914 decoy PSMs
[1021:22] Training neural networks on 26507 target and 12894 decoy PSMs
[1021:46] Number of IDs at 0.01 FDR: 20100
[1021:46] Precursors at 1% peptidoform FDR: 16160
[1021:46] Calculating protein q-values
[1021:46] Number of genes identified at 1% FDR: 3266 (precursor-level), 2878 (protein-level) (inference performed using proteotypic peptides only)
[1021:46] Quantification
[1021:47] Precursors with scored PTMs at 1% FDR: 42 out of 55 considered
[1021:47] Precursors with all scored PTM sites unoccupied at 1% FDR: 16118
[1021:47] Precursors with PTMs localised (when required) with > 90% confidence: 40 out of 42
[1021:49] XICs saved to D:\USERS\Soumyadeep\CellLysate_DIANN2\HvHIvHA_out/all_celllysates_comparison_xic/H_T1.xic.parquet

[1021:50] File #2/36
[1021:50] Loading run D:\USERS\Soumyadeep\Cell_Lysates\270625_cell\H_T2.wiff
[1031:00] Pre-processing...
[1031:01] 1117 MS1 and 111700 MS2 scans in 1117 (inferred) and 1117 (encoded) cycles, 46809 precursors in range
[1031:01] Calibrating with mass accuracies 28 (MS1), 25 (MS2)
[1031:02] RT window set to 0.674846
[1031:02] Recommended MS1 mass accuracy setting: 31 ppm
[1031:03] Searching decoys
[1031:04] Main search
[1031:06] Removing low confidence identifications
[1031:08] Removing interfering precursors
[1031:08] Training neural networks on 26734 target and 13054 decoy PSMs
[1031:33] Training neural networks on 26687 target and 12946 decoy PSMs
[1031:58] Number of IDs at 0.01 FDR: 20060
[1031:58] Precursors at 1% peptidoform FDR: 15967
[1031:58] Calculating protein q-values
[1031:58] Number of genes identified at 1% FDR: 3237 (precursor-level), 2849 (protein-level) (inference performed using proteotypic peptides only)
[1031:58] Quantification
[1031:59] Precursors with scored PTMs at 1% FDR: 41 out of 49 considered
[1031:59] Precursors with all scored PTM sites unoccupied at 1% FDR: 15926
[1031:59] Precursors with PTMs localised (when required) with > 90% confidence: 40 out of 41
[1032:00] XICs saved to D:\USERS\Soumyadeep\CellLysate_DIANN2\HvHIvHA_out/all_celllysates_comparison_xic/H_T2.xic.parquet

[1032:02] File #3/36
[1032:02] Loading run D:\USERS\Soumyadeep\Cell_Lysates\270625_cell\H_T3.wiff
[1041:55] Pre-processing...
[1041:56] 1117 MS1 and 111700 MS2 scans in 1117 (inferred) and 1117 (encoded) cycles, 46809 precursors in range
[1041:56] Calibrating with mass accuracies 28 (MS1), 25 (MS2)
[1041:57] RT window set to 0.662399
[1041:58] Recommended MS1 mass accuracy setting: 33 ppm
[1041:59] Searching decoys
[1041:59] Main search
[1042:01] Removing low confidence identifications
[1042:03] Removing interfering precursors
[1042:04] Training neural networks on 26716 target and 13128 decoy PSMs
[1042:28] Training neural networks on 26676 target and 12956 decoy PSMs
[1042:53] Number of IDs at 0.01 FDR: 20263
[1042:53] Precursors at 1% peptidoform FDR: 15973
[1042:53] Calculating protein q-values
[1042:53] Number of genes identified at 1% FDR: 3262 (precursor-level), 2824 (protein-level) (inference performed using proteotypic peptides only)
[1042:53] Quantification
[1042:54] Precursors with scored PTMs at 1% FDR: 40 out of 55 considered
[1042:54] Precursors with all scored PTM sites unoccupied at 1% FDR: 15933
[1042:54] Precursors with PTMs localised (when required) with > 90% confidence: 37 out of 40
[1042:56] XICs saved to D:\USERS\Soumyadeep\CellLysate_DIANN2\HvHIvHA_out/all_celllysates_comparison_xic/H_T3.xic.parquet

[1042:57] File #4/36
[1042:57] Loading run D:\USERS\Soumyadeep\Cell_Lysates\270625_cell\HA_T1.wiff
[1051:31] Pre-processing...
[1051:31] 1117 MS1 and 111700 MS2 scans in 1117 (inferred) and 1117 (encoded) cycles, 46809 precursors in range
[1051:31] Calibrating with mass accuracies 28 (MS1), 25 (MS2)
[1051:32] RT window set to 0.679749
[1051:32] Recommended MS1 mass accuracy setting: 32 ppm
[1051:33] Searching decoys
[1051:34] Main search
[1051:36] Removing low confidence identifications
[1051:38] Removing interfering precursors
[1051:38] Training neural networks on 30934 target and 15507 decoy PSMs
[1052:08] Training neural networks on 30889 target and 15771 decoy PSMs
[1052:36] Number of IDs at 0.01 FDR: 20273
[1052:36] Precursors at 1% peptidoform FDR: 14534
[1052:36] Calculating protein q-values
[1052:36] Number of genes identified at 1% FDR: 3112 (precursor-level), 2733 (protein-level) (inference performed using proteotypic peptides only)
[1052:36] Quantification
[1052:37] Precursors with scored PTMs at 1% FDR: 34 out of 49 considered
[1052:37] Precursors with all scored PTM sites unoccupied at 1% FDR: 14500
[1052:37] Precursors with PTMs localised (when required) with > 90% confidence: 31 out of 34
[1052:38] XICs saved to D:\USERS\Soumyadeep\CellLysate_DIANN2\HvHIvHA_out/all_celllysates_comparison_xic/HA_T1.xic.parquet

[1052:39] File #5/36
[1052:39] Loading run D:\USERS\Soumyadeep\Cell_Lysates\270625_cell\HA_T2.wiff
[1061:06] Pre-processing...
[1061:06] 1117 MS1 and 111700 MS2 scans in 1117 (inferred) and 1117 (encoded) cycles, 46809 precursors in range
[1061:06] Calibrating with mass accuracies 28 (MS1), 25 (MS2)
[1061:08] RT window set to 0.686405
[1061:08] Recommended MS1 mass accuracy setting: 34 ppm
[1061:09] Searching decoys
[1061:09] Main search
[1061:11] Removing low confidence identifications
[1061:13] Removing interfering precursors
[1061:14] Training neural networks on 30881 target and 15216 decoy PSMs
[1061:46] Training neural networks on 30840 target and 15188 decoy PSMs
[1062:15] Number of IDs at 0.01 FDR: 19884
[1062:15] Precursors at 1% peptidoform FDR: 14751
[1062:15] Calculating protein q-values
[1062:15] Number of genes identified at 1% FDR: 3086 (precursor-level), 2732 (protein-level) (inference performed using proteotypic peptides only)
[1062:15] Quantification
[1062:16] Precursors with scored PTMs at 1% FDR: 38 out of 50 considered
[1062:16] Precursors with all scored PTM sites unoccupied at 1% FDR: 14713
[1062:16] Precursors with PTMs localised (when required) with > 90% confidence: 35 out of 38
[1062:17] XICs saved to D:\USERS\Soumyadeep\CellLysate_DIANN2\HvHIvHA_out/all_celllysates_comparison_xic/HA_T2.xic.parquet

[1062:18] File #6/36
[1062:18] Loading run D:\USERS\Soumyadeep\Cell_Lysates\270625_cell\HA_T3.wiff
[1071:08] Pre-processing...
[1071:09] 1117 MS1 and 111700 MS2 scans in 1117 (inferred) and 1117 (encoded) cycles, 46809 precursors in range
[1071:09] Calibrating with mass accuracies 28 (MS1), 25 (MS2)
[1071:11] RT window set to 0.676161
[1071:11] Recommended MS1 mass accuracy setting: 33 ppm
[1071:12] Searching decoys
[1071:13] Main search
[1071:15] Removing low confidence identifications
[1071:17] Removing interfering precursors
[1071:18] Training neural networks on 30963 target and 15516 decoy PSMs
[1071:48] Training neural networks on 30911 target and 15435 decoy PSMs
[1072:15] Number of IDs at 0.01 FDR: 20768
[1072:16] Precursors at 1% peptidoform FDR: 15557
[1072:16] Calculating protein q-values
[1072:16] Number of genes identified at 1% FDR: 3180 (precursor-level), 2720 (protein-level) (inference performed using proteotypic peptides only)
[1072:16] Quantification
[1072:16] Precursors with scored PTMs at 1% FDR: 37 out of 52 considered
[1072:16] Precursors with all scored PTM sites unoccupied at 1% FDR: 15520
[1072:16] Precursors with PTMs localised (when required) with > 90% confidence: 36 out of 37
[1072:18] XICs saved to D:\USERS\Soumyadeep\CellLysate_DIANN2\HvHIvHA_out/all_celllysates_comparison_xic/HA_T3.xic.parquet

[1072:19] File #7/36
[1072:19] Loading run D:\USERS\Soumyadeep\Cell_Lysates\270625_cell\HI_T1.wiff
[1085:21] Pre-processing...
[1085:22] 1117 MS1 and 111700 MS2 scans in 1117 (inferred) and 1117 (encoded) cycles, 46809 precursors in range
[1085:22] Calibrating with mass accuracies 28 (MS1), 25 (MS2)
[1085:23] RT window set to 0.65929
[1085:23] Recommended MS1 mass accuracy setting: 34 ppm
[1085:24] Searching decoys
[1085:25] Main search
[1085:26] Removing low confidence identifications
[1085:29] Removing interfering precursors
[1085:29] Training neural networks on 28291 target and 14014 decoy PSMs
[1085:58] Training neural networks on 28250 target and 14041 decoy PSMs
[1086:24] Number of IDs at 0.01 FDR: 19613
[1086:24] Precursors at 1% peptidoform FDR: 15308
[1086:24] Calculating protein q-values
[1086:24] Number of genes identified at 1% FDR: 3184 (precursor-level), 2787 (protein-level) (inference performed using proteotypic peptides only)
[1086:24] Quantification
[1086:25] Precursors with scored PTMs at 1% FDR: 41 out of 52 considered
[1086:25] Precursors with all scored PTM sites unoccupied at 1% FDR: 15267
[1086:25] Precursors with PTMs localised (when required) with > 90% confidence: 39 out of 41
[1086:26] XICs saved to D:\USERS\Soumyadeep\CellLysate_DIANN2\HvHIvHA_out/all_celllysates_comparison_xic/HI_T1.xic.parquet

[1086:27] File #8/36
[1086:27] Loading run D:\USERS\Soumyadeep\Cell_Lysates\270625_cell\HI_T2.wiff
[1096:07] Pre-processing...
[1096:07] 1117 MS1 and 111700 MS2 scans in 1117 (inferred) and 1117 (encoded) cycles, 46809 precursors in range
[1096:07] Calibrating with mass accuracies 28 (MS1), 25 (MS2)
[1096:09] RT window set to 0.649061
[1096:09] Recommended MS1 mass accuracy setting: 33 ppm
[1096:10] Searching decoys
[1096:10] Main search
[1096:12] Removing low confidence identifications
[1096:14] Removing interfering precursors
[1096:15] Training neural networks on 27788 target and 13784 decoy PSMs
[1096:43] Training neural networks on 27746 target and 13704 decoy PSMs
[1097:10] Number of IDs at 0.01 FDR: 19200
[1097:10] Precursors at 1% peptidoform FDR: 14849
[1097:10] Calculating protein q-values
[1097:10] Number of genes identified at 1% FDR: 3159 (precursor-level), 2797 (protein-level) (inference performed using proteotypic peptides only)
[1097:10] Quantification
[1097:11] Precursors with scored PTMs at 1% FDR: 42 out of 54 considered
[1097:11] Precursors with all scored PTM sites unoccupied at 1% FDR: 14807
[1097:11] Precursors with PTMs localised (when required) with > 90% confidence: 40 out of 42
[1097:12] XICs saved to D:\USERS\Soumyadeep\CellLysate_DIANN2\HvHIvHA_out/all_celllysates_comparison_xic/HI_T2.xic.parquet

[1097:13] File #9/36
[1097:13] Loading run D:\USERS\Soumyadeep\Cell_Lysates\270625_cell\HI_T3.wiff
[1106:41] Pre-processing...
[1106:42] 1117 MS1 and 111700 MS2 scans in 1117 (inferred) and 1117 (encoded) cycles, 46809 precursors in range
[1106:42] Calibrating with mass accuracies 28 (MS1), 25 (MS2)
[1106:43] RT window set to 0.653008
[1106:44] Recommended MS1 mass accuracy setting: 33 ppm
[1106:45] Searching decoys
[1106:45] Main search
[1106:47] Removing low confidence identifications
[1106:50] Removing interfering precursors
[1106:50] Training neural networks on 27945 target and 13928 decoy PSMs
[1107:19] Training neural networks on 27895 target and 13923 decoy PSMs
[1107:46] Number of IDs at 0.01 FDR: 19654
[1107:46] Precursors at 1% peptidoform FDR: 15252
[1107:46] Calculating protein q-values
[1107:46] Number of genes identified at 1% FDR: 3174 (precursor-level), 2761 (protein-level) (inference performed using proteotypic peptides only)
[1107:46] Quantification
[1107:47] Precursors with scored PTMs at 1% FDR: 39 out of 48 considered
[1107:47] Precursors with all scored PTM sites unoccupied at 1% FDR: 15213
[1107:47] Precursors with PTMs localised (when required) with > 90% confidence: 37 out of 39
[1107:48] XICs saved to D:\USERS\Soumyadeep\CellLysate_DIANN2\HvHIvHA_out/all_celllysates_comparison_xic/HI_T3.xic.parquet

[1107:49] File #10/36
[1107:49] Loading run D:\USERS\Soumyadeep\Cell_Lysates\270625_cell\HIA_T1.wiff
[1117:35] Pre-processing...
[1117:36] 1117 MS1 and 111700 MS2 scans in 1117 (inferred) and 1117 (encoded) cycles, 46809 precursors in range
[1117:36] Calibrating with mass accuracies 28 (MS1), 25 (MS2)
[1117:37] RT window set to 0.664953
[1117:37] Recommended MS1 mass accuracy setting: 34 ppm
[1117:38] Searching decoys
[1117:39] Main search
[1117:41] Removing low confidence identifications
[1117:43] Removing interfering precursors
[1117:44] Training neural networks on 28390 target and 13968 decoy PSMs
[1118:12] Training neural networks on 28357 target and 13944 decoy PSMs
[1118:38] Number of IDs at 0.01 FDR: 20372
[1118:38] Precursors at 1% peptidoform FDR: 15634
[1118:38] Calculating protein q-values
[1118:38] Number of genes identified at 1% FDR: 3232 (precursor-level), 2781 (protein-level) (inference performed using proteotypic peptides only)
[1118:38] Quantification
[1118:39] Precursors with scored PTMs at 1% FDR: 43 out of 57 considered
[1118:39] Precursors with all scored PTM sites unoccupied at 1% FDR: 15591
[1118:39] Precursors with PTMs localised (when required) with > 90% confidence: 42 out of 43
[1118:40] XICs saved to D:\USERS\Soumyadeep\CellLysate_DIANN2\HvHIvHA_out/all_celllysates_comparison_xic/HIA_T1.xic.parquet

[1118:41] File #11/36
[1118:41] Loading run D:\USERS\Soumyadeep\Cell_Lysates\270625_cell\HIA_T2.wiff
[1128:49] Pre-processing...
[1128:49] 1117 MS1 and 111700 MS2 scans in 1117 (inferred) and 1117 (encoded) cycles, 46809 precursors in range
[1128:49] Calibrating with mass accuracies 28 (MS1), 25 (MS2)
[1128:51] RT window set to 0.668215
[1128:51] Recommended MS1 mass accuracy setting: 34 ppm
[1128:52] Searching decoys
[1128:53] Main search
[1128:55] Removing low confidence identifications
[1128:57] Removing interfering precursors
[1128:58] Training neural networks on 28270 target and 14029 decoy PSMs
[1129:26] Training neural networks on 28220 target and 13859 decoy PSMs
[1129:53] Number of IDs at 0.01 FDR: 19906
[1129:53] Precursors at 1% peptidoform FDR: 15462
[1129:53] Calculating protein q-values
[1129:53] Number of genes identified at 1% FDR: 3214 (precursor-level), 2792 (protein-level) (inference performed using proteotypic peptides only)
[1129:53] Quantification
[1129:54] Precursors with scored PTMs at 1% FDR: 43 out of 60 considered
[1129:54] Precursors with all scored PTM sites unoccupied at 1% FDR: 15419
[1129:54] Precursors with PTMs localised (when required) with > 90% confidence: 41 out of 43
[1129:55] XICs saved to D:\USERS\Soumyadeep\CellLysate_DIANN2\HvHIvHA_out/all_celllysates_comparison_xic/HIA_T2.xic.parquet

[1129:56] File #12/36
[1129:56] Loading run D:\USERS\Soumyadeep\Cell_Lysates\270625_cell\HIA_T3.wiff
[1140:11] Pre-processing...
[1140:11] 1117 MS1 and 111700 MS2 scans in 1117 (inferred) and 1117 (encoded) cycles, 46809 precursors in range
[1140:11] Calibrating with mass accuracies 28 (MS1), 25 (MS2)
[1140:13] RT window set to 0.669737
[1140:13] Recommended MS1 mass accuracy setting: 34 ppm
[1140:14] Searching decoys
[1140:15] Main search
[1140:17] Removing low confidence identifications
[1140:19] Removing interfering precursors
[1140:20] Training neural networks on 28253 target and 14008 decoy PSMs
[1140:47] Training neural networks on 28210 target and 13830 decoy PSMs
[1141:13] Number of IDs at 0.01 FDR: 20161
[1141:13] Precursors at 1% peptidoform FDR: 14764
[1141:13] Calculating protein q-values
[1141:13] Number of genes identified at 1% FDR: 3205 (precursor-level), 2814 (protein-level) (inference performed using proteotypic peptides only)
[1141:13] Quantification
[1141:14] Precursors with scored PTMs at 1% FDR: 40 out of 55 considered
[1141:14] Precursors with all scored PTM sites unoccupied at 1% FDR: 14724
[1141:14] Precursors with PTMs localised (when required) with > 90% confidence: 39 out of 40
[1141:16] XICs saved to D:\USERS\Soumyadeep\CellLysate_DIANN2\HvHIvHA_out/all_celllysates_comparison_xic/HIA_T3.xic.parquet

[1141:17] File #13/36
[1141:17] Loading run D:\USERS\Soumyadeep\020725\E2\H_E2_T1.wiff
[1151:08] Pre-processing...
[1151:09] 1117 MS1 and 111700 MS2 scans in 1117 (inferred) and 1117 (encoded) cycles, 46809 precursors in range
[1151:09] Calibrating with mass accuracies 28 (MS1), 25 (MS2)
[1151:11] RT window set to 0.684934
[1151:11] Recommended MS1 mass accuracy setting: 36 ppm
[1151:11] Searching decoys
[1151:12] Main search
[1151:15] Removing low confidence identifications
[1151:18] Removing interfering precursors
[1151:19] Training neural networks on 35256 target and 17388 decoy PSMs
[1151:53] Training neural networks on 35207 target and 17608 decoy PSMs
[1152:25] Number of IDs at 0.01 FDR: 27006
[1152:25] Precursors at 1% peptidoform FDR: 21830
[1152:25] Calculating protein q-values
[1152:25] Number of genes identified at 1% FDR: 3647 (precursor-level), 3141 (protein-level) (inference performed using proteotypic peptides only)
[1152:25] Quantification
[1152:26] Precursors with scored PTMs at 1% FDR: 46 out of 63 considered
[1152:26] Precursors with all scored PTM sites unoccupied at 1% FDR: 21784
[1152:26] Precursors with PTMs localised (when required) with > 90% confidence: 43 out of 46
[1152:28] XICs saved to D:\USERS\Soumyadeep\CellLysate_DIANN2\HvHIvHA_out/all_celllysates_comparison_xic/H_E2_T1.xic.parquet

[1152:29] File #14/36
[1152:29] Loading run D:\USERS\Soumyadeep\020725\E2\H_E2_T2.wiff
[1162:42] Pre-processing...
[1162:43] 1117 MS1 and 111700 MS2 scans in 1117 (inferred) and 1117 (encoded) cycles, 46809 precursors in range
[1162:43] Calibrating with mass accuracies 28 (MS1), 25 (MS2)
[1162:44] RT window set to 0.682908
[1162:44] Recommended MS1 mass accuracy setting: 34 ppm
[1162:45] Searching decoys
[1162:46] Main search
[1162:49] Removing low confidence identifications
[1162:52] Removing interfering precursors
[1162:53] Training neural networks on 35367 target and 17311 decoy PSMs
[1163:27] Training neural networks on 35318 target and 17427 decoy PSMs
[1163:59] Number of IDs at 0.01 FDR: 26952
[1164:00] Precursors at 1% peptidoform FDR: 21915
[1164:00] Calculating protein q-values
[1164:00] Number of genes identified at 1% FDR: 3597 (precursor-level), 3225 (protein-level) (inference performed using proteotypic peptides only)
[1164:00] Quantification
[1164:00] Precursors with scored PTMs at 1% FDR: 49 out of 60 considered
[1164:00] Precursors with all scored PTM sites unoccupied at 1% FDR: 21866
[1164:00] Precursors with PTMs localised (when required) with > 90% confidence: 48 out of 49
[1164:02] XICs saved to D:\USERS\Soumyadeep\CellLysate_DIANN2\HvHIvHA_out/all_celllysates_comparison_xic/H_E2_T2.xic.parquet

[1164:04] File #15/36
[1164:04] Loading run D:\USERS\Soumyadeep\020725\E2\H_E2_T3.wiff
[1174:07] Pre-processing...
[1174:07] 1117 MS1 and 111700 MS2 scans in 1117 (inferred) and 1117 (encoded) cycles, 46809 precursors in range
[1174:07] Calibrating with mass accuracies 28 (MS1), 25 (MS2)
[1174:09] RT window set to 0.68776
[1174:09] Recommended MS1 mass accuracy setting: 35 ppm
[1174:10] Searching decoys
[1174:11] Main search
[1174:14] Removing low confidence identifications
[1174:17] Removing interfering precursors
[1174:18] Training neural networks on 35328 target and 17377 decoy PSMs
[1174:51] Training neural networks on 35280 target and 17534 decoy PSMs
[1175:23] Number of IDs at 0.01 FDR: 27246
[1175:23] Precursors at 1% peptidoform FDR: 21467
[1175:23] Calculating protein q-values
[1175:23] Number of genes identified at 1% FDR: 3650 (precursor-level), 3137 (protein-level) (inference performed using proteotypic peptides only)
[1175:23] Quantification
[1175:24] Precursors with scored PTMs at 1% FDR: 49 out of 63 considered
[1175:24] Precursors with all scored PTM sites unoccupied at 1% FDR: 21418
[1175:24] Precursors with PTMs localised (when required) with > 90% confidence: 48 out of 49
[1175:26] XICs saved to D:\USERS\Soumyadeep\CellLysate_DIANN2\HvHIvHA_out/all_celllysates_comparison_xic/H_E2_T3.xic.parquet

[1175:28] File #16/36
[1175:28] Loading run D:\USERS\Soumyadeep\020725\E2\HA_E2_T1.wiff
[1183:26] Pre-processing...
[1183:26] 1117 MS1 and 111700 MS2 scans in 1117 (inferred) and 1117 (encoded) cycles, 46809 precursors in range
[1183:26] Calibrating with mass accuracies 28 (MS1), 25 (MS2)
[1183:28] RT window set to 0.691293
[1183:28] Recommended MS1 mass accuracy setting: 35 ppm
[1183:29] Searching decoys
[1183:29] Main search
[1183:31] Removing low confidence identifications
[1183:33] Removing interfering precursors
[1183:34] Training neural networks on 33805 target and 17403 decoy PSMs
[1184:07] Training neural networks on 33760 target and 17494 decoy PSMs
[1184:40] Number of IDs at 0.01 FDR: 22126
[1184:40] Precursors at 1% peptidoform FDR: 16861
[1184:40] Calculating protein q-values
[1184:40] Number of genes identified at 1% FDR: 3241 (precursor-level), 2843 (protein-level) (inference performed using proteotypic peptides only)
[1184:40] Quantification
[1184:41] Precursors with scored PTMs at 1% FDR: 46 out of 57 considered
[1184:41] Precursors with all scored PTM sites unoccupied at 1% FDR: 16815
[1184:41] Precursors with PTMs localised (when required) with > 90% confidence: 45 out of 46
[1184:42] XICs saved to D:\USERS\Soumyadeep\CellLysate_DIANN2\HvHIvHA_out/all_celllysates_comparison_xic/HA_E2_T1.xic.parquet

[1184:43] File #17/36
[1184:43] Loading run D:\USERS\Soumyadeep\020725\E2\HA_E2_T2.wiff
[1193:21] Pre-processing...
[1193:22] 1117 MS1 and 111700 MS2 scans in 1117 (inferred) and 1117 (encoded) cycles, 46809 precursors in range
[1193:22] Calibrating with mass accuracies 28 (MS1), 25 (MS2)
[1193:23] RT window set to 0.694299
[1193:23] Recommended MS1 mass accuracy setting: 35 ppm
[1193:24] Searching decoys
[1193:25] Main search
[1193:27] Removing low confidence identifications
[1193:30] Removing interfering precursors
[1193:31] Training neural networks on 34074 target and 17255 decoy PSMs
[1194:03] Training neural networks on 34028 target and 17457 decoy PSMs
[1194:35] Number of IDs at 0.01 FDR: 22329
[1194:35] Precursors at 1% peptidoform FDR: 16695
[1194:35] Calculating protein q-values
[1194:35] Number of genes identified at 1% FDR: 3224 (precursor-level), 2831 (protein-level) (inference performed using proteotypic peptides only)
[1194:35] Quantification
[1194:36] Precursors with scored PTMs at 1% FDR: 40 out of 56 considered
[1194:36] Precursors with all scored PTM sites unoccupied at 1% FDR: 16655
[1194:36] Precursors with PTMs localised (when required) with > 90% confidence: 38 out of 40
[1194:38] XICs saved to D:\USERS\Soumyadeep\CellLysate_DIANN2\HvHIvHA_out/all_celllysates_comparison_xic/HA_E2_T2.xic.parquet

[1194:39] File #18/36
[1194:39] Loading run D:\USERS\Soumyadeep\020725\E2\HA_E2_T3.wiff
[1203:28] Pre-processing...
[1203:29] 1117 MS1 and 111700 MS2 scans in 1117 (inferred) and 1117 (encoded) cycles, 46809 precursors in range
[1203:29] Calibrating with mass accuracies 28 (MS1), 25 (MS2)
[1203:30] RT window set to 0.676833
[1203:31] Recommended MS1 mass accuracy setting: 34 ppm
[1203:31] Searching decoys
[1203:32] Main search
[1203:34] Removing low confidence identifications
[1203:37] Removing interfering precursors
[1203:38] Training neural networks on 33790 target and 16990 decoy PSMs
[1204:12] Training neural networks on 33738 target and 17224 decoy PSMs
[1204:44] Number of IDs at 0.01 FDR: 22374
[1204:45] Precursors at 1% peptidoform FDR: 16080
[1204:45] Calculating protein q-values
[1204:45] Number of genes identified at 1% FDR: 3246 (precursor-level), 2860 (protein-level) (inference performed using proteotypic peptides only)
[1204:45] Quantification
[1204:45] Precursors with scored PTMs at 1% FDR: 36 out of 49 considered
[1204:45] Precursors with all scored PTM sites unoccupied at 1% FDR: 16044
[1204:45] Precursors with PTMs localised (when required) with > 90% confidence: 35 out of 36
[1204:47] XICs saved to D:\USERS\Soumyadeep\CellLysate_DIANN2\HvHIvHA_out/all_celllysates_comparison_xic/HA_E2_T3.xic.parquet

[1204:48] File #19/36
[1204:48] Loading run D:\USERS\Soumyadeep\020725\E2\HI_E2_T1.wiff
[1212:37] Pre-processing...
[1212:38] 1117 MS1 and 111700 MS2 scans in 1117 (inferred) and 1117 (encoded) cycles, 46809 precursors in range
[1212:38] Calibrating with mass accuracies 28 (MS1), 25 (MS2)
[1212:39] RT window set to 0.697163
[1212:39] Recommended MS1 mass accuracy setting: 35 ppm
[1212:40] Searching decoys
[1212:41] Main search
[1212:43] Removing low confidence identifications
[1212:45] Removing interfering precursors
[1212:46] Training neural networks on 33286 target and 16964 decoy PSMs
[1213:19] Training neural networks on 33252 target and 17161 decoy PSMs
[1213:50] Number of IDs at 0.01 FDR: 21129
[1213:50] Precursors at 1% peptidoform FDR: 15581
[1213:50] Calculating protein q-values
[1213:50] Number of genes identified at 1% FDR: 3222 (precursor-level), 2845 (protein-level) (inference performed using proteotypic peptides only)
[1213:50] Quantification
[1213:51] Precursors with scored PTMs at 1% FDR: 28 out of 52 considered
[1213:51] Precursors with all scored PTM sites unoccupied at 1% FDR: 15553
[1213:51] Precursors with PTMs localised (when required) with > 90% confidence: 27 out of 28
[1213:52] XICs saved to D:\USERS\Soumyadeep\CellLysate_DIANN2\HvHIvHA_out/all_celllysates_comparison_xic/HI_E2_T1.xic.parquet

[1213:53] File #20/36
[1213:53] Loading run D:\USERS\Soumyadeep\020725\E2\HI_E2_T2.wiff
[1222:31] Pre-processing...
[1222:31] 1117 MS1 and 111700 MS2 scans in 1117 (inferred) and 1117 (encoded) cycles, 46809 precursors in range
[1222:31] Calibrating with mass accuracies 28 (MS1), 25 (MS2)
[1222:33] RT window set to 0.690456
[1222:33] Recommended MS1 mass accuracy setting: 33 ppm
[1222:34] Searching decoys
[1222:35] Main search
[1222:37] Removing low confidence identifications
[1222:39] Removing interfering precursors
[1222:40] Training neural networks on 33095 target and 16712 decoy PSMs
[1223:13] Training neural networks on 33030 target and 17001 decoy PSMs
[1223:44] Number of IDs at 0.01 FDR: 21072
[1223:44] Precursors at 1% peptidoform FDR: 15108
[1223:44] Calculating protein q-values
[1223:44] Number of genes identified at 1% FDR: 3187 (precursor-level), 2755 (protein-level) (inference performed using proteotypic peptides only)
[1223:44] Quantification
[1223:45] Precursors with scored PTMs at 1% FDR: 33 out of 51 considered
[1223:45] Precursors with all scored PTM sites unoccupied at 1% FDR: 15075
[1223:45] Precursors with PTMs localised (when required) with > 90% confidence: 33 out of 33
[1223:47] XICs saved to D:\USERS\Soumyadeep\CellLysate_DIANN2\HvHIvHA_out/all_celllysates_comparison_xic/HI_E2_T2.xic.parquet

[1223:47] File #21/36
[1223:47] Loading run D:\USERS\Soumyadeep\020725\E2\HI_E2_T3.wiff
[1232:28] Pre-processing...
[1232:29] 1117 MS1 and 111700 MS2 scans in 1117 (inferred) and 1117 (encoded) cycles, 46809 precursors in range
[1232:29] Calibrating with mass accuracies 28 (MS1), 25 (MS2)
[1232:30] RT window set to 0.691913
[1232:30] Recommended MS1 mass accuracy setting: 33 ppm
[1232:31] Searching decoys
[1232:32] Main search
[1232:34] Removing low confidence identifications
[1232:37] Removing interfering precursors
[1232:38] Training neural networks on 33212 target and 17002 decoy PSMs
[1233:11] Training neural networks on 33170 target and 17046 decoy PSMs
[1233:41] Number of IDs at 0.01 FDR: 20906
[1233:41] Precursors at 1% peptidoform FDR: 15814
[1233:41] Calculating protein q-values
[1233:41] Number of genes identified at 1% FDR: 3169 (precursor-level), 2812 (protein-level) (inference performed using proteotypic peptides only)
[1233:41] Quantification
[1233:42] Precursors with scored PTMs at 1% FDR: 35 out of 49 considered
[1233:42] Precursors with all scored PTM sites unoccupied at 1% FDR: 15779
[1233:42] Precursors with PTMs localised (when required) with > 90% confidence: 35 out of 35
[1233:43] XICs saved to D:\USERS\Soumyadeep\CellLysate_DIANN2\HvHIvHA_out/all_celllysates_comparison_xic/HI_E2_T3.xic.parquet

[1233:44] File #22/36
[1233:44] Loading run D:\USERS\Soumyadeep\020725\E2\HIA_E2_T1.wiff
[1242:48] Pre-processing...
[1242:48] 1117 MS1 and 111700 MS2 scans in 1117 (inferred) and 1117 (encoded) cycles, 46809 precursors in range
[1242:48] Calibrating with mass accuracies 28 (MS1), 25 (MS2)
[1242:50] RT window set to 0.716822
[1242:50] Recommended MS1 mass accuracy setting: 34 ppm
[1242:51] Searching decoys
[1242:52] Main search
[1242:54] Removing low confidence identifications
[1242:57] Removing interfering precursors
[1242:58] Training neural networks on 34455 target and 17423 decoy PSMs
[1243:34] Training neural networks on 34414 target and 17673 decoy PSMs
[1244:08] Number of IDs at 0.01 FDR: 21907
[1244:08] Precursors at 1% peptidoform FDR: 15690
[1244:08] Calculating protein q-values
[1244:08] Number of genes identified at 1% FDR: 3234 (precursor-level), 2889 (protein-level) (inference performed using proteotypic peptides only)
[1244:08] Quantification
[1244:09] Precursors with scored PTMs at 1% FDR: 36 out of 59 considered
[1244:09] Precursors with all scored PTM sites unoccupied at 1% FDR: 15654
[1244:09] Precursors with PTMs localised (when required) with > 90% confidence: 34 out of 36
[1244:11] XICs saved to D:\USERS\Soumyadeep\CellLysate_DIANN2\HvHIvHA_out/all_celllysates_comparison_xic/HIA_E2_T1.xic.parquet

[1244:12] File #23/36
[1244:12] Loading run D:\USERS\Soumyadeep\020725\E2\HIA_E2_T2.wiff
[1253:32] Pre-processing...
[1253:33] 1117 MS1 and 111700 MS2 scans in 1117 (inferred) and 1117 (encoded) cycles, 46809 precursors in range
[1253:33] Calibrating with mass accuracies 28 (MS1), 25 (MS2)
[1253:34] RT window set to 0.729658
[1253:34] Recommended MS1 mass accuracy setting: 33 ppm
[1253:36] Searching decoys
[1253:36] Main search
[1253:39] Removing low confidence identifications
[1253:41] Removing interfering precursors
[1253:42] Training neural networks on 34326 target and 17174 decoy PSMs
[1254:16] Training neural networks on 34282 target and 17414 decoy PSMs
[1254:49] Number of IDs at 0.01 FDR: 21902
[1254:49] Precursors at 1% peptidoform FDR: 14936
[1254:49] Calculating protein q-values
[1254:49] Number of genes identified at 1% FDR: 3214 (precursor-level), 2857 (protein-level) (inference performed using proteotypic peptides only)
[1254:49] Quantification
[1254:50] Precursors with scored PTMs at 1% FDR: 35 out of 56 considered
[1254:50] Precursors with all scored PTM sites unoccupied at 1% FDR: 14901
[1254:50] Precursors with PTMs localised (when required) with > 90% confidence: 33 out of 35
[1254:51] XICs saved to D:\USERS\Soumyadeep\CellLysate_DIANN2\HvHIvHA_out/all_celllysates_comparison_xic/HIA_E2_T2.xic.parquet

[1254:52] File #24/36
[1254:52] Loading run D:\USERS\Soumyadeep\020725\E2\HIA_E2_T3.wiff
[1264:23] Pre-processing...
[1264:24] 1117 MS1 and 111700 MS2 scans in 1117 (inferred) and 1117 (encoded) cycles, 46809 precursors in range
[1264:24] Calibrating with mass accuracies 28 (MS1), 25 (MS2)
[1264:26] RT window set to 0.722653
[1264:26] Recommended MS1 mass accuracy setting: 34 ppm
[1264:27] Searching decoys
[1264:28] Main search
[1264:30] Removing low confidence identifications
[1264:33] Removing interfering precursors
[1264:33] Training neural networks on 34384 target and 17323 decoy PSMs
[1265:08] Training neural networks on 34336 target and 17548 decoy PSMs
[1265:41] Number of IDs at 0.01 FDR: 21988
[1265:41] Precursors at 1% peptidoform FDR: 15843
[1265:41] Calculating protein q-values
[1265:41] Number of genes identified at 1% FDR: 3178 (precursor-level), 2791 (protein-level) (inference performed using proteotypic peptides only)
[1265:41] Quantification
[1265:42] Precursors with scored PTMs at 1% FDR: 39 out of 57 considered
[1265:42] Precursors with all scored PTM sites unoccupied at 1% FDR: 15804
[1265:42] Precursors with PTMs localised (when required) with > 90% confidence: 37 out of 39
[1265:44] XICs saved to D:\USERS\Soumyadeep\CellLysate_DIANN2\HvHIvHA_out/all_celllysates_comparison_xic/HIA_E2_T3.xic.parquet

[1265:45] File #25/36
[1265:45] Loading run D:\USERS\Soumyadeep\020725\E3\H_E3_T1.wiff
[1277:00] Pre-processing...
[1277:00] 1117 MS1 and 111700 MS2 scans in 1117 (inferred) and 1117 (encoded) cycles, 46809 precursors in range
[1277:00] Calibrating with mass accuracies 28 (MS1), 25 (MS2)
[1277:01] RT window set to 0.767323
[1277:01] Recommended MS1 mass accuracy setting: 36 ppm
[1277:02] Searching decoys
[1277:03] Main search
[1277:07] Removing low confidence identifications
[1277:10] Removing interfering precursors
[1277:12] Training neural networks on 41367 target and 21342 decoy PSMs
[1277:50] Training neural networks on 41314 target and 22002 decoy PSMs
[1278:31] Number of IDs at 0.01 FDR: 33555
[1278:31] Precursors at 1% peptidoform FDR: 25424
[1278:31] Calculating protein q-values
[1278:31] Number of genes identified at 1% FDR: 3874 (precursor-level), 3381 (protein-level) (inference performed using proteotypic peptides only)
[1278:31] Quantification
[1278:32] Precursors with scored PTMs at 1% FDR: 58 out of 78 considered
[1278:32] Precursors with all scored PTM sites unoccupied at 1% FDR: 25366
[1278:32] Precursors with PTMs localised (when required) with > 90% confidence: 56 out of 58
[1278:34] XICs saved to D:\USERS\Soumyadeep\CellLysate_DIANN2\HvHIvHA_out/all_celllysates_comparison_xic/H_E3_T1.xic.parquet

[1278:36] File #26/36
[1278:36] Loading run D:\USERS\Soumyadeep\020725\E3\H_E3_T2.wiff
[1290:16] Pre-processing...
[1290:17] 1117 MS1 and 111700 MS2 scans in 1117 (inferred) and 1117 (encoded) cycles, 46809 precursors in range
[1290:17] Calibrating with mass accuracies 28 (MS1), 25 (MS2)
[1290:18] RT window set to 0.766498
[1290:18] Recommended MS1 mass accuracy setting: 36 ppm
[1290:18] Searching decoys
[1290:20] Main search
[1290:23] Removing low confidence identifications
[1290:27] Removing interfering precursors
[1290:28] Training neural networks on 41430 target and 21371 decoy PSMs
[1291:09] Training neural networks on 41383 target and 21779 decoy PSMs
[1291:48] Number of IDs at 0.01 FDR: 33271
[1291:48] Precursors at 1% peptidoform FDR: 26387
[1291:48] Calculating protein q-values
[1291:48] Number of genes identified at 1% FDR: 3866 (precursor-level), 3366 (protein-level) (inference performed using proteotypic peptides only)
[1291:48] Quantification
[1291:49] Precursors with scored PTMs at 1% FDR: 60 out of 74 considered
[1291:49] Precursors with all scored PTM sites unoccupied at 1% FDR: 26327
[1291:49] Precursors with PTMs localised (when required) with > 90% confidence: 57 out of 60
[1291:51] XICs saved to D:\USERS\Soumyadeep\CellLysate_DIANN2\HvHIvHA_out/all_celllysates_comparison_xic/H_E3_T2.xic.parquet

[1291:53] File #27/36
[1291:53] Loading run D:\USERS\Soumyadeep\020725\E3\H_E3_T3.wiff
[1303:21] Pre-processing...
[1303:22] 1117 MS1 and 111700 MS2 scans in 1117 (inferred) and 1117 (encoded) cycles, 46809 precursors in range
[1303:22] Calibrating with mass accuracies 28 (MS1), 25 (MS2)
[1303:23] RT window set to 0.765428
[1303:23] Recommended MS1 mass accuracy setting: 37 ppm
[1303:23] Searching decoys
[1303:25] Main search
[1303:28] Removing low confidence identifications
[1303:32] Removing interfering precursors
[1303:33] Training neural networks on 41408 target and 21200 decoy PSMs
[1304:14] Training neural networks on 41363 target and 21738 decoy PSMs
[1304:52] Number of IDs at 0.01 FDR: 33469
[1304:52] Precursors at 1% peptidoform FDR: 24786
[1304:52] Calculating protein q-values
[1304:52] Number of genes identified at 1% FDR: 3875 (precursor-level), 3405 (protein-level) (inference performed using proteotypic peptides only)
[1304:52] Quantification
[1304:53] Precursors with scored PTMs at 1% FDR: 52 out of 72 considered
[1304:53] Precursors with all scored PTM sites unoccupied at 1% FDR: 24734
[1304:53] Precursors with PTMs localised (when required) with > 90% confidence: 50 out of 52
[1304:55] XICs saved to D:\USERS\Soumyadeep\CellLysate_DIANN2\HvHIvHA_out/all_celllysates_comparison_xic/H_E3_T3.xic.parquet

[1304:57] File #28/36
[1304:57] Loading run D:\USERS\Soumyadeep\020725\E3\HA_E3_T1.wiff
[1315:57] Pre-processing...
[1315:58] 1117 MS1 and 111700 MS2 scans in 1117 (inferred) and 1117 (encoded) cycles, 46809 precursors in range
[1315:58] Calibrating with mass accuracies 28 (MS1), 25 (MS2)
[1315:58] RT window set to 0.763081
[1315:58] Recommended MS1 mass accuracy setting: 38 ppm
[1315:59] Searching decoys
[1316:01] Main search
[1316:04] Removing low confidence identifications
[1316:07] Removing interfering precursors
[1316:09] Training neural networks on 41217 target and 20801 decoy PSMs
[1316:50] Training neural networks on 41155 target and 21423 decoy PSMs
[1317:30] Number of IDs at 0.01 FDR: 32020
[1317:30] Precursors at 1% peptidoform FDR: 23679
[1317:30] Calculating protein q-values
[1317:30] Number of genes identified at 1% FDR: 3771 (precursor-level), 3321 (protein-level) (inference performed using proteotypic peptides only)
[1317:30] Quantification
[1317:31] Precursors with scored PTMs at 1% FDR: 54 out of 74 considered
[1317:31] Precursors with all scored PTM sites unoccupied at 1% FDR: 23625
[1317:31] Precursors with PTMs localised (when required) with > 90% confidence: 52 out of 54
[1317:33] XICs saved to D:\USERS\Soumyadeep\CellLysate_DIANN2\HvHIvHA_out/all_celllysates_comparison_xic/HA_E3_T1.xic.parquet

[1317:35] File #29/36
[1317:35] Loading run D:\USERS\Soumyadeep\020725\E3\HA_E3_T2.wiff
[1332:44] Pre-processing...
[1332:45] 1117 MS1 and 111700 MS2 scans in 1117 (inferred) and 1117 (encoded) cycles, 46809 precursors in range
[1332:45] Calibrating with mass accuracies 28 (MS1), 25 (MS2)
[1332:45] RT window set to 0.763776
[1332:46] Recommended MS1 mass accuracy setting: 37 ppm
[1332:46] Searching decoys
[1332:48] Main search
[1332:51] Removing low confidence identifications
[1332:54] Removing interfering precursors
[1332:55] Training neural networks on 41322 target and 21243 decoy PSMs
[1333:36] Training neural networks on 41267 target and 21677 decoy PSMs
[1334:12] Number of IDs at 0.01 FDR: 31935
[1334:12] Precursors at 1% peptidoform FDR: 23606
[1334:12] Calculating protein q-values
[1334:12] Number of genes identified at 1% FDR: 3776 (precursor-level), 3371 (protein-level) (inference performed using proteotypic peptides only)
[1334:12] Quantification
[1334:13] Precursors with scored PTMs at 1% FDR: 53 out of 68 considered
[1334:13] Precursors with all scored PTM sites unoccupied at 1% FDR: 23553
[1334:13] Precursors with PTMs localised (when required) with > 90% confidence: 50 out of 53
[1334:15] XICs saved to D:\USERS\Soumyadeep\CellLysate_DIANN2\HvHIvHA_out/all_celllysates_comparison_xic/HA_E3_T2.xic.parquet

[1334:17] File #30/36
[1334:17] Loading run D:\USERS\Soumyadeep\020725\E3\HA_E3_T3.wiff
[1345:26] Pre-processing...
[1345:27] 1117 MS1 and 111700 MS2 scans in 1117 (inferred) and 1117 (encoded) cycles, 46809 precursors in range
[1345:27] Calibrating with mass accuracies 28 (MS1), 25 (MS2)
[1345:27] RT window set to 0.76579
[1345:28] Recommended MS1 mass accuracy setting: 37 ppm
[1345:28] Searching decoys
[1345:30] Main search
[1345:33] Removing low confidence identifications
[1345:36] Removing interfering precursors
[1345:37] Training neural networks on 41282 target and 20956 decoy PSMs
[1346:18] Training neural networks on 41224 target and 21554 decoy PSMs
[1346:56] Number of IDs at 0.01 FDR: 31433
[1346:56] Precursors at 1% peptidoform FDR: 24043
[1346:56] Calculating protein q-values
[1346:56] Number of genes identified at 1% FDR: 3710 (precursor-level), 3262 (protein-level) (inference performed using proteotypic peptides only)
[1346:56] Quantification
[1346:57] Precursors with scored PTMs at 1% FDR: 53 out of 68 considered
[1346:57] Precursors with all scored PTM sites unoccupied at 1% FDR: 23990
[1346:57] Precursors with PTMs localised (when required) with > 90% confidence: 51 out of 53
[1346:59] XICs saved to D:\USERS\Soumyadeep\CellLysate_DIANN2\HvHIvHA_out/all_celllysates_comparison_xic/HA_E3_T3.xic.parquet

[1347:01] File #31/36
[1347:01] Loading run D:\USERS\Soumyadeep\020725\E3\HI_E3_T1.wiff
[1358:28] Pre-processing...
[1358:29] 1117 MS1 and 111700 MS2 scans in 1117 (inferred) and 1117 (encoded) cycles, 46809 precursors in range
[1358:29] Calibrating with mass accuracies 28 (MS1), 25 (MS2)
[1358:30] RT window set to 0.757275
[1358:30] Recommended MS1 mass accuracy setting: 36 ppm
[1358:31] Searching decoys
[1358:32] Main search
[1358:36] Removing low confidence identifications
[1358:39] Removing interfering precursors
[1358:41] Training neural networks on 40839 target and 20646 decoy PSMs
[1359:20] Training neural networks on 40792 target and 21277 decoy PSMs
[1359:57] Number of IDs at 0.01 FDR: 31336
[1359:57] Precursors at 1% peptidoform FDR: 23651
[1359:57] Calculating protein q-values
[1359:57] Number of genes identified at 1% FDR: 3732 (precursor-level), 3267 (protein-level) (inference performed using proteotypic peptides only)
[1359:57] Quantification
[1359:58] Precursors with scored PTMs at 1% FDR: 55 out of 71 considered
[1359:58] Precursors with all scored PTM sites unoccupied at 1% FDR: 23596
[1359:58] Precursors with PTMs localised (when required) with > 90% confidence: 54 out of 55
[1360:01] XICs saved to D:\USERS\Soumyadeep\CellLysate_DIANN2\HvHIvHA_out/all_celllysates_comparison_xic/HI_E3_T1.xic.parquet

[1360:02] File #32/36
[1360:02] Loading run D:\USERS\Soumyadeep\020725\E3\HI_E3_T2.wiff
[1371:26] Pre-processing...
[1371:27] 1117 MS1 and 111700 MS2 scans in 1117 (inferred) and 1117 (encoded) cycles, 46809 precursors in range
[1371:27] Calibrating with mass accuracies 28 (MS1), 25 (MS2)
[1371:28] RT window set to 0.773671
[1371:28] Recommended MS1 mass accuracy setting: 36 ppm
[1371:29] Searching decoys
[1371:30] Main search
[1371:34] Removing low confidence identifications
[1371:37] Removing interfering precursors
[1371:38] Training neural networks on 40970 target and 20626 decoy PSMs
[1372:16] Training neural networks on 40924 target and 21247 decoy PSMs
[1372:53] Number of IDs at 0.01 FDR: 31708
[1372:54] Precursors at 1% peptidoform FDR: 23371
[1372:54] Calculating protein q-values
[1372:54] Number of genes identified at 1% FDR: 3775 (precursor-level), 3304 (protein-level) (inference performed using proteotypic peptides only)
[1372:54] Quantification
[1372:54] Precursors with scored PTMs at 1% FDR: 48 out of 67 considered
[1372:54] Precursors with all scored PTM sites unoccupied at 1% FDR: 23323
[1372:54] Precursors with PTMs localised (when required) with > 90% confidence: 46 out of 48
[1372:57] XICs saved to D:\USERS\Soumyadeep\CellLysate_DIANN2\HvHIvHA_out/all_celllysates_comparison_xic/HI_E3_T2.xic.parquet

[1372:58] File #33/36
[1372:58] Loading run D:\USERS\Soumyadeep\020725\E3\HI_E3_T3.wiff
[1384:24] Pre-processing...
[1384:25] 1117 MS1 and 111700 MS2 scans in 1117 (inferred) and 1117 (encoded) cycles, 46809 precursors in range
[1384:25] Calibrating with mass accuracies 28 (MS1), 25 (MS2)
[1384:26] RT window set to 0.763146
[1384:26] Recommended MS1 mass accuracy setting: 35 ppm
[1384:27] Searching decoys
[1384:28] Main search
[1384:31] Removing low confidence identifications
[1384:35] Removing interfering precursors
[1384:36] Training neural networks on 40946 target and 21026 decoy PSMs
[1385:17] Training neural networks on 40903 target and 21637 decoy PSMs
[1385:57] Number of IDs at 0.01 FDR: 31655
[1385:57] Precursors at 1% peptidoform FDR: 23870
[1385:57] Calculating protein q-values
[1385:57] Number of genes identified at 1% FDR: 3761 (precursor-level), 3257 (protein-level) (inference performed using proteotypic peptides only)
[1385:57] Quantification
[1385:58] Precursors with scored PTMs at 1% FDR: 49 out of 70 considered
[1385:58] Precursors with all scored PTM sites unoccupied at 1% FDR: 23821
[1385:58] Precursors with PTMs localised (when required) with > 90% confidence: 47 out of 49
[1386:00] XICs saved to D:\USERS\Soumyadeep\CellLysate_DIANN2\HvHIvHA_out/all_celllysates_comparison_xic/HI_E3_T3.xic.parquet

[1386:02] File #34/36
[1386:02] Loading run D:\USERS\Soumyadeep\020725\E3\HIA_E3_T1.wiff
[1398:51] Pre-processing...
[1398:52] 1117 MS1 and 111700 MS2 scans in 1117 (inferred) and 1117 (encoded) cycles, 46809 precursors in range
[1398:52] Calibrating with mass accuracies 28 (MS1), 25 (MS2)
[1398:53] RT window set to 0.76766
[1398:53] Recommended MS1 mass accuracy setting: 38 ppm
[1398:54] Searching decoys
[1398:55] Main search
[1398:59] Removing low confidence identifications
[1399:03] Removing interfering precursors
[1399:04] Training neural networks on 41514 target and 21173 decoy PSMs
[1399:44] Training neural networks on 41455 target and 21692 decoy PSMs
[1400:20] Number of IDs at 0.01 FDR: 33827
[1400:20] Precursors at 1% peptidoform FDR: 26210
[1400:20] Calculating protein q-values
[1400:20] Number of genes identified at 1% FDR: 3947 (precursor-level), 3442 (protein-level) (inference performed using proteotypic peptides only)
[1400:20] Quantification
[1400:21] Precursors with scored PTMs at 1% FDR: 61 out of 75 considered
[1400:21] Precursors with all scored PTM sites unoccupied at 1% FDR: 26149
[1400:21] Precursors with PTMs localised (when required) with > 90% confidence: 60 out of 61
[1400:23] XICs saved to D:\USERS\Soumyadeep\CellLysate_DIANN2\HvHIvHA_out/all_celllysates_comparison_xic/HIA_E3_T1.xic.parquet

[1400:26] File #35/36
[1400:26] Loading run D:\USERS\Soumyadeep\020725\E3\HIA_E3_T2.wiff
[1413:53] Pre-processing...
[1413:54] 1117 MS1 and 111700 MS2 scans in 1117 (inferred) and 1117 (encoded) cycles, 46809 precursors in range
[1413:54] Calibrating with mass accuracies 28 (MS1), 25 (MS2)
[1413:55] RT window set to 0.762642
[1413:55] Recommended MS1 mass accuracy setting: 39 ppm
[1413:56] Searching decoys
[1413:57] Main search
[1414:01] Removing low confidence identifications
[1414:05] Removing interfering precursors
[1414:06] Training neural networks on 41623 target and 21310 decoy PSMs
[1414:45] Training neural networks on 41567 target and 21821 decoy PSMs
[1415:24] Number of IDs at 0.01 FDR: 33900
[1415:24] Precursors at 1% peptidoform FDR: 26092
[1415:24] Calculating protein q-values
[1415:24] Number of genes identified at 1% FDR: 3930 (precursor-level), 3473 (protein-level) (inference performed using proteotypic peptides only)
[1415:24] Quantification
[1415:25] Precursors with scored PTMs at 1% FDR: 64 out of 83 considered
[1415:25] Precursors with all scored PTM sites unoccupied at 1% FDR: 26028
[1415:25] Precursors with PTMs localised (when required) with > 90% confidence: 61 out of 64
[1415:28] XICs saved to D:\USERS\Soumyadeep\CellLysate_DIANN2\HvHIvHA_out/all_celllysates_comparison_xic/HIA_E3_T2.xic.parquet

[1415:31] File #36/36
[1415:31] Loading run D:\USERS\Soumyadeep\020725\E3\HIA_E3_T3.wiff
[1428:13] Pre-processing...
[1428:14] 1117 MS1 and 111700 MS2 scans in 1117 (inferred) and 1117 (encoded) cycles, 46809 precursors in range
[1428:14] Calibrating with mass accuracies 28 (MS1), 25 (MS2)
[1428:14] RT window set to 0.77242
[1428:14] Recommended MS1 mass accuracy setting: 37 ppm
[1428:15] Searching decoys
[1428:17] Main search
[1428:20] Removing low confidence identifications
[1428:24] Removing interfering precursors
[1428:25] Training neural networks on 41480 target and 21287 decoy PSMs
[1429:05] Training neural networks on 41424 target and 21684 decoy PSMs
[1429:42] Number of IDs at 0.01 FDR: 33336
[1429:43] Precursors at 1% peptidoform FDR: 25369
[1429:43] Calculating protein q-values
[1429:43] Number of genes identified at 1% FDR: 3882 (precursor-level), 3458 (protein-level) (inference performed using proteotypic peptides only)
[1429:43] Quantification
[1429:44] Precursors with scored PTMs at 1% FDR: 60 out of 77 considered
[1429:44] Precursors with all scored PTM sites unoccupied at 1% FDR: 25309
[1429:44] Precursors with PTMs localised (when required) with > 90% confidence: 58 out of 60
[1429:46] XICs saved to D:\USERS\Soumyadeep\CellLysate_DIANN2\HvHIvHA_out/all_celllysates_comparison_xic/HIA_E3_T3.xic.parquet

[1429:48] Cross-run analysis
[1429:48] Reading quantification information: 36 files
[1429:53] Quantifying peptides
[1435:21] Quantification parameters: 0.359406, 0.0555465, 1.30147, 0.0136352, 0.166776, 0.0329797, 0.237273, 0.220561, 0.224366, 0.0436949, 0.252609, 0.15286, 0.283934, 0.145661, 0.182778, 0.0827297
[1437:09] Quantifying proteins
[1437:09] Calculating q-values for protein and gene groups
[1437:10] Calculating global q-values for protein and gene groups
[1437:10] Protein groups with global q-value <= 0.01: 3614
[1437:20] Compressed report saved to D:\USERS\Soumyadeep\CellLysate_DIANN2\HvHIvHA_out\all_celllysates_comparison.parquet. Use R 'arrow' or Python 'PyArrow' package to process
[1437:20] Site report saved to D:\USERS\Soumyadeep\CellLysate_DIANN2\HvHIvHA_out\all_celllysates_comparison.site_report.parquet
[1437:20] Saving precursor levels matrix
[1437:21] Precursor levels matrix (1% precursor and protein group FDR) saved to D:\USERS\Soumyadeep\CellLysate_DIANN2\HvHIvHA_out\all_celllysates_comparison.pr_matrix.tsv.
[1437:21] Saving protein group levels matrix
[1437:22] Protein groups matrix saved to D:\USERS\Soumyadeep\CellLysate_DIANN2\HvHIvHA_out\all_celllysates_comparison.pg_matrix.tsv.
[1437:22] Saving gene group levels matrix
[1437:22] Gene groups matrix saved to D:\USERS\Soumyadeep\CellLysate_DIANN2\HvHIvHA_out\all_celllysates_comparison.gg_matrix.tsv.
[1437:22] Saving unique genes levels matrix
[1437:22] Unique genes matrix saved to D:\USERS\Soumyadeep\CellLysate_DIANN2\HvHIvHA_out\all_celllysates_comparison.unique_genes_matrix.tsv.
[1437:22] Manifest saved to D:\USERS\Soumyadeep\CellLysate_DIANN2\HvHIvHA_out\all_celllysates_comparison.manifest.txt
[1437:22] Stats report saved to D:\USERS\Soumyadeep\CellLysate_DIANN2\HvHIvHA_out\all_celllysates_comparison.stats.tsv
