## Supplementary File 4 for "Dynamic s-acylation of schizophrenia-linked antioxidants explicate the redox flexibility of human microglia during inflammation"

DIA-NN 2.2.0 Academia  (Data-Independent Acquisition by Neural Networks)
Compiled on May 29 2025 21:29:29
Current date and time: Fri Jul 11 11:07:59 2025
CPU: GenuineIntel 12th Gen Intel(R) Core(TM) i7-12700
SIMD instructions: AVX AVX2 FMA SSE4.1 SSE4.2 
Logical CPU cores: 20
diann.exe --f D:\Soumyadeep\Tissue\Tissue Run 2 MS Data\RAW FILES\B184F_T1.wiff  --f D:\Soumyadeep\Tissue\Tissue Run 2 MS Data\RAW FILES\B184H_T1.wiff  --f D:\Soumyadeep\Tissue\Tissue Run 2 MS Data\RAW FILES\B185F_T1.wiff  --f D:\Soumyadeep\Tissue\Tissue Run 2 MS Data\RAW FILES\B212F_T1.wiff  --f D:\Soumyadeep\Tissue\Tissue Run 2 MS Data\RAW FILES\B214F_T1.wiff  --f D:\Soumyadeep\Tissue\Tissue Run 2 MS Data\RAW FILES\B229F_T1.wiff  --f D:\Soumyadeep\Tissue\Tissue Run 3 MS Data\280425_SD_CJ\B184F_T2.wiff  --f D:\Soumyadeep\Tissue\Tissue Run 3 MS Data\280425_SD_CJ\B184H_T2.wiff  --f D:\Soumyadeep\Tissue\Tissue Run 3 MS Data\280425_SD_CJ\B185F_T2.wiff  --f D:\Soumyadeep\Tissue\Tissue Run 3 MS Data\280425_SD_CJ\B212F_T2.wiff  --f D:\Soumyadeep\Tissue\Tissue Run 3 MS Data\280425_SD_CJ\B214F_T2.wiff  --f D:\Soumyadeep\Tissue\Tissue Run 3 MS Data\280425_SD_CJ\B229F_T2.wiff  --lib D:\Soumyadeep\Cell Click\lib\humwp_uniprot_canfinal_lib.predicted.speclib --threads 18 --verbose 1 --out D:\Soumyadeep\Tissue\out\tissue_diannreport.parquet --qvalue 0.05 --matrices --temp D:\Soumyadeep\Tissue\temp --xic --fasta camprotR_240512_cRAP_20190401_full_tags.fasta --cont-quant-exclude cRAP- --fasta D:\Soumyadeep\Cell Click\clean_humwholeprot_reviewed_can.fasta --met-excision --cut K*,R*,!*P --missed-cleavages 2 --unimod4 --var-mods 3 --var-mod UniMod:35,15.994915,M --reanalyse --rt-profiling 

Thread number set to 18
Output will be filtered at 0.05 FDR
Precursor/protein x samples expression level matrices will be saved along with the main report
XICs within 10 seconds from the apex will be extracted for each precursor and saved in .parquet format, a folder will be created next to the main report for the XICs storage
Peptides corresponding to protein sequence IDs tagged with cRAP- will be excluded from normalisation as well as quantification of protein groups that do not include proteins bearing the tag
N-terminal methionine excision enabled
In silico digest will involve cuts at K*,R*
But excluding cuts at *P
Maximum number of missed cleavages set to 2
Cysteine carbamidomethylation enabled as a fixed modification
Maximum number of variable modifications set to 3
Modification UniMod:35 with mass delta 15.9949 at M will be considered as variable
MBR enabled; .quant files will only be saved to disk during the first pass
The spectral library (if generated) will retain the original spectra but will include empirically-aligned RTs
DIA-NN will automatically optimise the mass accuracy for the first run of the experiment, use this mode for preliminary analyses only
WARNING: peptidoform scoring enabled because variable modifications have been declared; to disable, use --no-peptidoforms
The following variable modifications will be localised: UniMod:35 
WARNING: protein inference will only be performed for precursors identified with global q-value <= 0.05 or q-value <= 0.05.

12 files will be processed
[0:00] Loading spectral library D:\Soumyadeep\Cell Click\lib\humwp_uniprot_canfinal_lib.predicted.speclib
[0:05] Library annotated with sequence database(s): D:\Soumyadeep\Cell Click\clean_humwholeprot_reviewed_can.fasta
[0:06] Spectral library loaded: 20368 protein isoforms, 29405 protein groups and 7700783 precursors in 2804689 elution groups.
[0:06] Loading protein annotations from FASTA camprotR_240512_cRAP_20190401_full_tags.fasta
[0:06] Loading protein annotations from FASTA D:\Soumyadeep\Cell Click\clean_humwholeprot_reviewed_can.fasta
[0:06] Annotating library proteins with information from the FASTA database
[0:07] Gene names missing for some isoforms
[0:07] Library contains 20368 proteins, and 20174 genes
[0:11] Initialising library

First pass: generating a spectral library from DIA data

[0:26] File #1/12
[0:26] Loading run D:\Soumyadeep\Tissue\Tissue Run 2 MS Data\RAW FILES\B184F_T1.wiff
[1:44] Pre-processing...
[1:44] 1600 MS1 and 44800 MS2 scans in 1600 (inferred) and 1600 (encoded) cycles, 7700783 precursors in range
[1:45] Calibrating with mass accuracies 25 (MS1), 25 (MS2)
[18:16] RT window set to 2.8193
[18:16] Peak width: 3.872
[18:16] Scan window radius set to 8
[18:17] Recommended MS1 mass accuracy setting: 10 ppm
[63:27] Optimised mass accuracy: 16 ppm
[76:25] Searching decoys
[80:45] Main search
[89:17] Removing low confidence identifications
[89:21] Removing interfering precursors
[89:24] Training neural networks on 3894 target and 2328 decoy PSMs
[89:32] Training neural networks on 3894 target and 2073 decoy PSMs
[89:37] Number of IDs at 0.01 FDR: 2056
[89:37] Precursors at 1% peptidoform FDR: 1798
[89:37] Calculating protein q-values
[89:38] Number of genes identified at 1% FDR: 619 (precursor-level), 563 (protein-level) (inference performed using proteotypic peptides only)
[89:38] Quantification
[89:38] Precursors with scored PTMs at 1% FDR: 9 out of 18 considered
[89:38] Precursors with all scored PTM sites unoccupied at 1% FDR: 1789
[89:38] Precursors with PTMs localised (when required) with > 90% confidence: 9 out of 9
[89:38] Quantification information saved to D:\Soumyadeep\Tissue\temp/D__Soumyadeep_Tissue_Tissue Run 2 MS Data_RAW FILES_B184F_T1_wiff.quant

[89:38] File #2/12
[89:38] Loading run D:\Soumyadeep\Tissue\Tissue Run 2 MS Data\RAW FILES\B184H_T1.wiff
[91:10] Pre-processing...
[91:11] 1600 MS1 and 44800 MS2 scans in 1600 (inferred) and 1600 (encoded) cycles, 7700783 precursors in range
[91:11] Calibrating with mass accuracies 25 (MS1), 25 (MS2)
[103:03] RT window set to 2.66904
[103:04] Recommended MS1 mass accuracy setting: 11 ppm
[110:16] Searching decoys
[114:47] Main search
[123:39] Removing low confidence identifications
[123:44] Removing interfering precursors
[123:46] Training neural networks on 4870 target and 3063 decoy PSMs
[123:54] Training neural networks on 4870 target and 2671 decoy PSMs
[124:00] Number of IDs at 0.01 FDR: 2524
[124:00] Precursors at 1% peptidoform FDR: 2308
[124:01] Calculating protein q-values
[124:01] Number of genes identified at 1% FDR: 727 (precursor-level), 656 (protein-level) (inference performed using proteotypic peptides only)
[124:01] Quantification
[124:01] Precursors with scored PTMs at 1% FDR: 8 out of 10 considered
[124:01] Precursors with all scored PTM sites unoccupied at 1% FDR: 2300
[124:01] Precursors with PTMs localised (when required) with > 90% confidence: 7 out of 8
[124:02] Quantification information saved to D:\Soumyadeep\Tissue\temp/D__Soumyadeep_Tissue_Tissue Run 2 MS Data_RAW FILES_B184H_T1_wiff.quant

[124:02] File #3/12
[124:02] Loading run D:\Soumyadeep\Tissue\Tissue Run 2 MS Data\RAW FILES\B185F_T1.wiff
[125:34] Pre-processing...
[125:34] 1600 MS1 and 44800 MS2 scans in 1600 (inferred) and 1600 (encoded) cycles, 7700783 precursors in range
[125:35] Calibrating with mass accuracies 25 (MS1), 25 (MS2)
[142:57] RT window set to 2.53038
[142:58] Recommended MS1 mass accuracy setting: 12 ppm
[148:40] Searching decoys
[152:25] Main search
[159:47] Removing low confidence identifications
[159:52] Removing interfering precursors
[159:54] Training neural networks on 4353 target and 2695 decoy PSMs
[160:02] Training neural networks on 4353 target and 2437 decoy PSMs
[160:07] Number of IDs at 0.01 FDR: 2260
[160:08] Precursors at 1% peptidoform FDR: 2114
[160:08] Calculating protein q-values
[160:08] Number of genes identified at 1% FDR: 676 (precursor-level), 633 (protein-level) (inference performed using proteotypic peptides only)
[160:09] Quantification
[160:09] Precursors with scored PTMs at 1% FDR: 6 out of 9 considered
[160:09] Precursors with all scored PTM sites unoccupied at 1% FDR: 2108
[160:09] Precursors with PTMs localised (when required) with > 90% confidence: 5 out of 6
[160:09] Quantification information saved to D:\Soumyadeep\Tissue\temp/D__Soumyadeep_Tissue_Tissue Run 2 MS Data_RAW FILES_B185F_T1_wiff.quant

[160:09] File #4/12
[160:09] Loading run D:\Soumyadeep\Tissue\Tissue Run 2 MS Data\RAW FILES\B212F_T1.wiff
[162:03] Pre-processing...
[162:03] 1600 MS1 and 44800 MS2 scans in 1600 (inferred) and 1600 (encoded) cycles, 7700783 precursors in range
[162:04] Calibrating with mass accuracies 25 (MS1), 25 (MS2)
[174:49] RT window set to 2.45348
[174:49] Recommended MS1 mass accuracy setting: 10 ppm
[180:14] Searching decoys
[185:00] Main search
[194:24] Removing low confidence identifications
[194:29] Removing interfering precursors
[194:32] Training neural networks on 7307 target and 4968 decoy PSMs
[194:41] Training neural networks on 7307 target and 4348 decoy PSMs
[194:47] Number of IDs at 0.01 FDR: 3152
[194:47] Precursors at 1% peptidoform FDR: 2766
[194:48] Calculating protein q-values
[194:48] Number of genes identified at 1% FDR: 847 (precursor-level), 777 (protein-level) (inference performed using proteotypic peptides only)
[194:48] Quantification
[194:48] Precursors with scored PTMs at 1% FDR: 22 out of 35 considered
[194:48] Precursors with all scored PTM sites unoccupied at 1% FDR: 2744
[194:48] Precursors with PTMs localised (when required) with > 90% confidence: 18 out of 22
[194:49] Quantification information saved to D:\Soumyadeep\Tissue\temp/D__Soumyadeep_Tissue_Tissue Run 2 MS Data_RAW FILES_B212F_T1_wiff.quant

[194:49] File #5/12
[194:49] Loading run D:\Soumyadeep\Tissue\Tissue Run 2 MS Data\RAW FILES\B214F_T1.wiff
[196:42] Pre-processing...
[196:43] 1600 MS1 and 44800 MS2 scans in 1600 (inferred) and 1600 (encoded) cycles, 7700783 precursors in range
[196:44] Calibrating with mass accuracies 25 (MS1), 25 (MS2)
[207:09] RT window set to 3.05716
[207:09] Recommended MS1 mass accuracy setting: 12 ppm
[214:17] Searching decoys
[221:43] Main search
[236:27] Removing low confidence identifications
[236:32] Removing interfering precursors
[236:35] Training neural networks on 5595 target and 3966 decoy PSMs
[236:43] Training neural networks on 5595 target and 3462 decoy PSMs
[236:49] Number of IDs at 0.01 FDR: 2606
[236:49] Precursors at 1% peptidoform FDR: 2216
[236:50] Calculating protein q-values
[236:50] Number of genes identified at 1% FDR: 821 (precursor-level), 749 (protein-level) (inference performed using proteotypic peptides only)
[236:50] Quantification
[236:50] Precursors with scored PTMs at 1% FDR: 6 out of 22 considered
[236:50] Precursors with all scored PTM sites unoccupied at 1% FDR: 2210
[236:50] Precursors with PTMs localised (when required) with > 90% confidence: 5 out of 6
[236:51] Quantification information saved to D:\Soumyadeep\Tissue\temp/D__Soumyadeep_Tissue_Tissue Run 2 MS Data_RAW FILES_B214F_T1_wiff.quant

[236:51] File #6/12
[236:51] Loading run D:\Soumyadeep\Tissue\Tissue Run 2 MS Data\RAW FILES\B229F_T1.wiff
[238:41] Pre-processing...
[238:42] 1600 MS1 and 44800 MS2 scans in 1600 (inferred) and 1600 (encoded) cycles, 7700783 precursors in range
[238:43] Calibrating with mass accuracies 25 (MS1), 25 (MS2)
[256:55] RT window set to 2.04728
[256:56] Recommended MS1 mass accuracy setting: 12 ppm
[262:22] Searching decoys
[265:43] Main search
[272:17] Removing low confidence identifications
[272:21] Removing interfering precursors
[272:24] Training neural networks on 6023 target and 4350 decoy PSMs
[272:32] Training neural networks on 6023 target and 3933 decoy PSMs
[272:38] Number of IDs at 0.01 FDR: 2444
[272:39] Precursors at 1% peptidoform FDR: 2221
[272:39] Calculating protein q-values
[272:39] Number of genes identified at 1% FDR: 751 (precursor-level), 672 (protein-level) (inference performed using proteotypic peptides only)
[272:39] Quantification
[272:39] Precursors with scored PTMs at 1% FDR: 6 out of 13 considered
[272:39] Precursors with all scored PTM sites unoccupied at 1% FDR: 2215
[272:39] Precursors with PTMs localised (when required) with > 90% confidence: 4 out of 6
[272:40] Quantification information saved to D:\Soumyadeep\Tissue\temp/D__Soumyadeep_Tissue_Tissue Run 2 MS Data_RAW FILES_B229F_T1_wiff.quant

[272:40] File #7/12
[272:40] Loading run D:\Soumyadeep\Tissue\Tissue Run 3 MS Data\280425_SD_CJ\B184F_T2.wiff
[274:01] Pre-processing...
[274:01] 1600 MS1 and 44800 MS2 scans in 1600 (inferred) and 1600 (encoded) cycles, 7700783 precursors in range
[274:02] Calibrating with mass accuracies 25 (MS1), 25 (MS2)
[285:25] RT window set to 2.48792
[285:26] Recommended MS1 mass accuracy setting: 11 ppm
[292:03] Searching decoys
[297:31] Main search
[308:18] Removing low confidence identifications
[308:22] Removing interfering precursors
[308:25] Training neural networks on 4647 target and 2776 decoy PSMs
[308:33] Training neural networks on 4647 target and 2402 decoy PSMs
[308:38] Number of IDs at 0.01 FDR: 2729
[308:38] Precursors at 1% peptidoform FDR: 2255
[308:38] Calculating protein q-values
[308:39] Number of genes identified at 1% FDR: 806 (precursor-level), 750 (protein-level) (inference performed using proteotypic peptides only)
[308:39] Quantification
[308:39] Precursors with scored PTMs at 1% FDR: 3 out of 13 considered
[308:39] Precursors with all scored PTM sites unoccupied at 1% FDR: 2252
[308:39] Precursors with PTMs localised (when required) with > 90% confidence: 3 out of 3
[308:39] Quantification information saved to D:\Soumyadeep\Tissue\temp/D__Soumyadeep_Tissue_Tissue Run 3 MS Data_280425_SD_CJ_B184F_T2_wiff.quant

[308:40] File #8/12
[308:40] Loading run D:\Soumyadeep\Tissue\Tissue Run 3 MS Data\280425_SD_CJ\B184H_T2.wiff
[310:02] Pre-processing...
[310:02] 1600 MS1 and 44800 MS2 scans in 1600 (inferred) and 1600 (encoded) cycles, 7700783 precursors in range
[310:03] Calibrating with mass accuracies 25 (MS1), 25 (MS2)
[321:39] RT window set to 2.74807
[321:40] Recommended MS1 mass accuracy setting: 11 ppm
[329:13] Searching decoys
[334:05] Main search
[343:39] Removing low confidence identifications
[343:43] Removing interfering precursors
[343:46] Training neural networks on 4174 target and 2579 decoy PSMs
[343:54] Training neural networks on 4174 target and 2177 decoy PSMs
[343:59] Number of IDs at 0.01 FDR: 2431
[344:00] Precursors at 1% peptidoform FDR: 2161
[344:00] Calculating protein q-values
[344:00] Number of genes identified at 1% FDR: 723 (precursor-level), 650 (protein-level) (inference performed using proteotypic peptides only)
[344:00] Quantification
[344:00] Precursors with scored PTMs at 1% FDR: 23 out of 28 considered
[344:00] Precursors with all scored PTM sites unoccupied at 1% FDR: 2138
[344:00] Precursors with PTMs localised (when required) with > 90% confidence: 21 out of 23
[344:01] Quantification information saved to D:\Soumyadeep\Tissue\temp/D__Soumyadeep_Tissue_Tissue Run 3 MS Data_280425_SD_CJ_B184H_T2_wiff.quant

[344:01] File #9/12
[344:01] Loading run D:\Soumyadeep\Tissue\Tissue Run 3 MS Data\280425_SD_CJ\B185F_T2.wiff
[345:26] Pre-processing...
[345:27] 1600 MS1 and 44800 MS2 scans in 1600 (inferred) and 1600 (encoded) cycles, 7700783 precursors in range
[345:28] Calibrating with mass accuracies 25 (MS1), 25 (MS2)
[356:55] RT window set to 3.47766
[356:56] Recommended MS1 mass accuracy setting: 11 ppm
[368:16] Searching decoys
[373:35] Main search
[384:03] Removing low confidence identifications
[384:07] Removing interfering precursors
[384:10] Training neural networks on 5294 target and 3562 decoy PSMs
[384:18] Training neural networks on 5294 target and 3148 decoy PSMs
[384:24] Number of IDs at 0.01 FDR: 2622
[384:24] Precursors at 1% peptidoform FDR: 2301
[384:24] Calculating protein q-values
[384:24] Number of genes identified at 1% FDR: 801 (precursor-level), 738 (protein-level) (inference performed using proteotypic peptides only)
[384:25] Quantification
[384:25] Precursors with scored PTMs at 1% FDR: 7 out of 15 considered
[384:25] Precursors with all scored PTM sites unoccupied at 1% FDR: 2294
[384:25] Precursors with PTMs localised (when required) with > 90% confidence: 6 out of 7
[384:25] Quantification information saved to D:\Soumyadeep\Tissue\temp/D__Soumyadeep_Tissue_Tissue Run 3 MS Data_280425_SD_CJ_B185F_T2_wiff.quant

[384:25] File #10/12
[384:25] Loading run D:\Soumyadeep\Tissue\Tissue Run 3 MS Data\280425_SD_CJ\B212F_T2.wiff
[386:02] Pre-processing...
[386:03] 1600 MS1 and 44800 MS2 scans in 1600 (inferred) and 1600 (encoded) cycles, 7700783 precursors in range
[386:03] Calibrating with mass accuracies 25 (MS1), 25 (MS2)
[394:28] RT window set to 2.9
[394:28] Recommended MS1 mass accuracy setting: 13 ppm
[400:01] Searching decoys
[405:02] Main search
[414:56] Removing low confidence identifications
[415:01] Removing interfering precursors
[415:04] Training neural networks on 6313 target and 4085 decoy PSMs
[415:12] Training neural networks on 6313 target and 3417 decoy PSMs
[415:18] Number of IDs at 0.01 FDR: 3204
[415:19] Precursors at 1% peptidoform FDR: 2933
[415:19] Calculating protein q-values
[415:19] Number of genes identified at 1% FDR: 854 (precursor-level), 773 (protein-level) (inference performed using proteotypic peptides only)
[415:19] Quantification
[415:20] Precursors with scored PTMs at 1% FDR: 32 out of 42 considered
[415:20] Precursors with all scored PTM sites unoccupied at 1% FDR: 2901
[415:20] Precursors with PTMs localised (when required) with > 90% confidence: 28 out of 32
[415:20] Quantification information saved to D:\Soumyadeep\Tissue\temp/D__Soumyadeep_Tissue_Tissue Run 3 MS Data_280425_SD_CJ_B212F_T2_wiff.quant

[415:20] File #11/12
[415:20] Loading run D:\Soumyadeep\Tissue\Tissue Run 3 MS Data\280425_SD_CJ\B214F_T2.wiff
[417:00] Pre-processing...
[417:00] 1600 MS1 and 44800 MS2 scans in 1600 (inferred) and 1600 (encoded) cycles, 7700783 precursors in range
[417:01] Calibrating with mass accuracies 25 (MS1), 25 (MS2)
[426:55] RT window set to 2.82366
[426:56] Recommended MS1 mass accuracy setting: 13 ppm
[433:10] Searching decoys
[437:26] Main search
[445:49] Removing low confidence identifications
[445:54] Removing interfering precursors
[445:57] Training neural networks on 6457 target and 4671 decoy PSMs
[446:05] Training neural networks on 6457 target and 4188 decoy PSMs
[446:11] Number of IDs at 0.01 FDR: 2763
[446:12] Precursors at 1% peptidoform FDR: 2405
[446:12] Calculating protein q-values
[446:12] Number of genes identified at 1% FDR: 866 (precursor-level), 796 (protein-level) (inference performed using proteotypic peptides only)
[446:12] Quantification
[446:13] Precursors with scored PTMs at 1% FDR: 9 out of 23 considered
[446:13] Precursors with all scored PTM sites unoccupied at 1% FDR: 2396
[446:13] Precursors with PTMs localised (when required) with > 90% confidence: 8 out of 9
[446:13] Quantification information saved to D:\Soumyadeep\Tissue\temp/D__Soumyadeep_Tissue_Tissue Run 3 MS Data_280425_SD_CJ_B214F_T2_wiff.quant

[446:13] File #12/12
[446:13] Loading run D:\Soumyadeep\Tissue\Tissue Run 3 MS Data\280425_SD_CJ\B229F_T2.wiff
[447:47] Pre-processing...
[447:47] 1600 MS1 and 44800 MS2 scans in 1600 (inferred) and 1600 (encoded) cycles, 7700783 precursors in range
[447:48] Calibrating with mass accuracies 25 (MS1), 25 (MS2)
[459:39] RT window set to 2.46563
[459:39] Recommended MS1 mass accuracy setting: 12 ppm
[464:01] Searching decoys
[467:26] Main search
[474:08] Removing low confidence identifications
[474:13] Removing interfering precursors
[474:15] Training neural networks on 6262 target and 4428 decoy PSMs
[474:24] Training neural networks on 6262 target and 4017 decoy PSMs
[474:30] Number of IDs at 0.01 FDR: 2507
[474:31] Precursors at 1% peptidoform FDR: 2303
[474:31] Calculating protein q-values
[474:31] Number of genes identified at 1% FDR: 759 (precursor-level), 724 (protein-level) (inference performed using proteotypic peptides only)
[474:31] Quantification
[474:31] Precursors with scored PTMs at 1% FDR: 11 out of 17 considered
[474:31] Precursors with all scored PTM sites unoccupied at 1% FDR: 2292
[474:31] Precursors with PTMs localised (when required) with > 90% confidence: 11 out of 11
[474:32] Quantification information saved to D:\Soumyadeep\Tissue\temp/D__Soumyadeep_Tissue_Tissue Run 3 MS Data_280425_SD_CJ_B229F_T2_wiff.quant

[474:32] Cross-run analysis
[474:32] Reading quantification information: 12 files
[474:46] Quantifying peptides
[474:50] Assembling protein groups
[474:51] Quantifying proteins
[474:51] Calculating q-values for protein and gene groups
[474:54] Calculating global q-values for protein and gene groups
[474:54] Protein groups with global q-value <= 0.01: 1234
[474:54] Compressed report saved to D:\Soumyadeep\Tissue\out\tissue_diannreport-first-pass.parquet. Use R 'arrow' or Python 'PyArrow' package to process
[474:54] Site report saved to D:\Soumyadeep\Tissue\out\tissue_diannreport-first-pass.site_report.parquet
[474:54] Saving precursor levels matrix
[474:54] Precursor levels matrix (1% precursor and protein group FDR) saved to D:\Soumyadeep\Tissue\out\tissue_diannreport-first-pass.pr_matrix.tsv.
[474:54] Manifest saved to D:\Soumyadeep\Tissue\out\tissue_diannreport-first-pass.manifest.txt
[474:54] Stats report saved to D:\Soumyadeep\Tissue\out\tissue_diannreport-first-pass.stats.tsv
[474:54] Generating spectral library:
[474:55] 5963 target and 304 decoy precursors saved
[474:55] Spectral library saved to D:\Soumyadeep\Tissue\out\tissue_diannreport-lib.parquet

[474:56] Loading spectral library D:\Soumyadeep\Tissue\out\tissue_diannreport-lib.parquet
[474:56] Spectral library loaded: 2013 protein isoforms, 1928 protein groups and 6267 precursors in 4913 elution groups.
[474:56] Loading protein annotations from FASTA camprotR_240512_cRAP_20190401_full_tags.fasta
[474:56] Loading protein annotations from FASTA D:\Soumyadeep\Cell Click\clean_humwholeprot_reviewed_can.fasta
[474:56] Annotating library proteins with information from the FASTA database
[474:56] Gene names missing for some isoforms
[474:56] Library contains 2013 proteins, and 2011 genes
[474:56] Initialising library
[474:57] Saving the library to D:\Soumyadeep\Tissue\out\tissue_diannreport-lib.parquet.skyline.speclib


Second pass: using the newly created spectral library to reanalyse the data

[474:57] File #1/12
[474:57] Loading run D:\Soumyadeep\Tissue\Tissue Run 2 MS Data\RAW FILES\B184F_T1.wiff
[476:13] Pre-processing...
[476:13] 1600 MS1 and 44800 MS2 scans in 1600 (inferred) and 1600 (encoded) cycles, 5963 precursors in range
[476:13] Calibrating with mass accuracies 25 (MS1), 25 (MS2)
[476:14] RT window set to 0.516552
[476:14] Recommended MS1 mass accuracy setting: 13 ppm
[476:14] Searching decoys
[476:15] Main search
[476:15] Removing low confidence identifications
[476:15] Removing interfering precursors
[476:15] Training neural networks on 4629 target and 3981 decoy PSMs
[476:20] Training neural networks on 4621 target and 3762 decoy PSMs
[476:25] Number of IDs at 0.01 FDR: 2690
[476:25] Precursors at 1% peptidoform FDR: 1312
[476:25] Calculating protein q-values
[476:25] Number of genes identified at 1% FDR: 796 (precursor-level), 703 (protein-level) (inference performed using proteotypic peptides only)
[476:25] Quantification
[476:25] Precursors with scored PTMs at 1% FDR: 16 out of 27 considered
[476:25] Precursors with all scored PTM sites unoccupied at 1% FDR: 1296
[476:25] Precursors with PTMs localised (when required) with > 90% confidence: 15 out of 16
[476:25] XICs saved to D:\Soumyadeep\Tissue\out/tissue_diannreport_xic/B184F_T1.xic.parquet

[476:26] File #2/12
[476:26] Loading run D:\Soumyadeep\Tissue\Tissue Run 2 MS Data\RAW FILES\B184H_T1.wiff
[477:58] Pre-processing...
[477:58] 1600 MS1 and 44800 MS2 scans in 1600 (inferred) and 1600 (encoded) cycles, 5963 precursors in range
[477:58] Calibrating with mass accuracies 25 (MS1), 25 (MS2)
[477:59] RT window set to 0.502692
[477:59] Recommended MS1 mass accuracy setting: 14 ppm
[477:59] Searching decoys
[478:00] Main search
[478:00] Removing low confidence identifications
[478:00] Removing interfering precursors
[478:01] Training neural networks on 4672 target and 4029 decoy PSMs
[478:06] Training neural networks on 4668 target and 3756 decoy PSMs
[478:12] Number of IDs at 0.01 FDR: 3445
[478:12] Precursors at 1% peptidoform FDR: 2296
[478:12] Calculating protein q-values
[478:12] Number of genes identified at 1% FDR: 948 (precursor-level), 864 (protein-level) (inference performed using proteotypic peptides only)
[478:12] Quantification
[478:12] Precursors with scored PTMs at 1% FDR: 13 out of 19 considered
[478:12] Precursors with all scored PTM sites unoccupied at 1% FDR: 2283
[478:12] Precursors with PTMs localised (when required) with > 90% confidence: 12 out of 13
[478:12] XICs saved to D:\Soumyadeep\Tissue\out/tissue_diannreport_xic/B184H_T1.xic.parquet

[478:12] File #3/12
[478:12] Loading run D:\Soumyadeep\Tissue\Tissue Run 2 MS Data\RAW FILES\B185F_T1.wiff
[479:45] Pre-processing...
[479:45] 1600 MS1 and 44800 MS2 scans in 1600 (inferred) and 1600 (encoded) cycles, 5963 precursors in range
[479:45] Calibrating with mass accuracies 25 (MS1), 25 (MS2)
[479:46] RT window set to 0.539136
[479:46] Recommended MS1 mass accuracy setting: 13 ppm
[479:47] Searching decoys
[479:47] Main search
[479:47] Removing low confidence identifications
[479:48] Removing interfering precursors
[479:48] Training neural networks on 4608 target and 3997 decoy PSMs
[479:53] Training neural networks on 4601 target and 3772 decoy PSMs
[479:59] Number of IDs at 0.01 FDR: 3068
[479:59] Precursors at 1% peptidoform FDR: 1022
[479:59] Calculating protein q-values
[479:59] Number of genes identified at 1% FDR: 902 (precursor-level), 749 (protein-level) (inference performed using proteotypic peptides only)
[479:59] Quantification
[479:59] Precursors with scored PTMs at 1% FDR: 7 out of 27 considered
[479:59] Precursors with all scored PTM sites unoccupied at 1% FDR: 1015
[479:59] Precursors with PTMs localised (when required) with > 90% confidence: 7 out of 7
[479:59] XICs saved to D:\Soumyadeep\Tissue\out/tissue_diannreport_xic/B185F_T1.xic.parquet

[479:59] File #4/12
[479:59] Loading run D:\Soumyadeep\Tissue\Tissue Run 2 MS Data\RAW FILES\B212F_T1.wiff
[481:54] Pre-processing...
[481:55] 1600 MS1 and 44800 MS2 scans in 1600 (inferred) and 1600 (encoded) cycles, 5963 precursors in range
[481:55] Calibrating with mass accuracies 25 (MS1), 25 (MS2)
[481:56] RT window set to 0.5157
[481:56] Recommended MS1 mass accuracy setting: 15 ppm
[481:57] Searching decoys
[481:57] Main search
[481:57] Removing low confidence identifications
[481:58] Removing interfering precursors
[481:58] Training neural networks on 4734 target and 4087 decoy PSMs
[482:04] Training neural networks on 4725 target and 3806 decoy PSMs
[482:09] Number of IDs at 0.01 FDR: 3586
[482:09] Precursors at 1% peptidoform FDR: 2626
[482:09] Calculating protein q-values
[482:09] Number of genes identified at 1% FDR: 962 (precursor-level), 827 (protein-level) (inference performed using proteotypic peptides only)
[482:09] Quantification
[482:10] Precursors with scored PTMs at 1% FDR: 29 out of 41 considered
[482:10] Precursors with all scored PTM sites unoccupied at 1% FDR: 2597
[482:10] Precursors with PTMs localised (when required) with > 90% confidence: 26 out of 29
[482:10] XICs saved to D:\Soumyadeep\Tissue\out/tissue_diannreport_xic/B212F_T1.xic.parquet

[482:10] File #5/12
[482:10] Loading run D:\Soumyadeep\Tissue\Tissue Run 2 MS Data\RAW FILES\B214F_T1.wiff
[484:03] Pre-processing...
[484:04] 1600 MS1 and 44800 MS2 scans in 1600 (inferred) and 1600 (encoded) cycles, 5963 precursors in range
[484:04] Calibrating with mass accuracies 25 (MS1), 25 (MS2)
[484:05] RT window set to 0.514763
[484:05] Recommended MS1 mass accuracy setting: 13 ppm
[484:05] Searching decoys
[484:06] Main search
[484:06] Removing low confidence identifications
[484:06] Removing interfering precursors
[484:06] Training neural networks on 4630 target and 3962 decoy PSMs
[484:12] Training neural networks on 4624 target and 3752 decoy PSMs
[484:18] Number of IDs at 0.01 FDR: 3261
[484:18] Precursors at 1% peptidoform FDR: 2172
[484:18] Calculating protein q-values
[484:18] Number of genes identified at 1% FDR: 978 (precursor-level), 827 (protein-level) (inference performed using proteotypic peptides only)
[484:18] Quantification
[484:18] Precursors with scored PTMs at 1% FDR: 13 out of 21 considered
[484:18] Precursors with all scored PTM sites unoccupied at 1% FDR: 2159
[484:18] Precursors with PTMs localised (when required) with > 90% confidence: 12 out of 13
[484:18] XICs saved to D:\Soumyadeep\Tissue\out/tissue_diannreport_xic/B214F_T1.xic.parquet

[484:19] File #6/12
[484:19] Loading run D:\Soumyadeep\Tissue\Tissue Run 2 MS Data\RAW FILES\B229F_T1.wiff
[486:09] Pre-processing...
[486:09] 1600 MS1 and 44800 MS2 scans in 1600 (inferred) and 1600 (encoded) cycles, 5963 precursors in range
[486:09] Calibrating with mass accuracies 25 (MS1), 25 (MS2)
[486:10] RT window set to 0.520158
[486:10] Recommended MS1 mass accuracy setting: 14 ppm
[486:11] Searching decoys
[486:11] Main search
[486:11] Removing low confidence identifications
[486:12] Removing interfering precursors
[486:12] Training neural networks on 4599 target and 3982 decoy PSMs
[486:18] Training neural networks on 4590 target and 3763 decoy PSMs
[486:23] Number of IDs at 0.01 FDR: 3204
[486:23] Precursors at 1% peptidoform FDR: 1611
[486:23] Calculating protein q-values
[486:23] Number of genes identified at 1% FDR: 952 (precursor-level), 804 (protein-level) (inference performed using proteotypic peptides only)
[486:23] Quantification
[486:23] Precursors with scored PTMs at 1% FDR: 10 out of 22 considered
[486:23] Precursors with all scored PTM sites unoccupied at 1% FDR: 1601
[486:23] Precursors with PTMs localised (when required) with > 90% confidence: 9 out of 10
[486:24] XICs saved to D:\Soumyadeep\Tissue\out/tissue_diannreport_xic/B229F_T1.xic.parquet

[486:24] File #7/12
[486:24] Loading run D:\Soumyadeep\Tissue\Tissue Run 3 MS Data\280425_SD_CJ\B184F_T2.wiff
[487:45] Pre-processing...
[487:45] 1600 MS1 and 44800 MS2 scans in 1600 (inferred) and 1600 (encoded) cycles, 5963 precursors in range
[487:45] Calibrating with mass accuracies 25 (MS1), 25 (MS2)
[487:46] RT window set to 0.505332
[487:46] Recommended MS1 mass accuracy setting: 14 ppm
[487:46] Searching decoys
[487:46] Main search
[487:47] Removing low confidence identifications
[487:47] Removing interfering precursors
[487:47] Training neural networks on 4683 target and 4047 decoy PSMs
[487:53] Training neural networks on 4676 target and 3753 decoy PSMs
[487:59] Number of IDs at 0.01 FDR: 3644
[487:59] Precursors at 1% peptidoform FDR: 1781
[487:59] Calculating protein q-values
[487:59] Number of genes identified at 1% FDR: 1004 (precursor-level), 809 (protein-level) (inference performed using proteotypic peptides only)
[487:59] Quantification
[487:59] Precursors with scored PTMs at 1% FDR: 12 out of 26 considered
[487:59] Precursors with all scored PTM sites unoccupied at 1% FDR: 1769
[487:59] Precursors with PTMs localised (when required) with > 90% confidence: 12 out of 12
[487:59] XICs saved to D:\Soumyadeep\Tissue\out/tissue_diannreport_xic/B184F_T2.xic.parquet

[487:59] File #8/12
[487:59] Loading run D:\Soumyadeep\Tissue\Tissue Run 3 MS Data\280425_SD_CJ\B184H_T2.wiff
[489:22] Pre-processing...
[489:22] 1600 MS1 and 44800 MS2 scans in 1600 (inferred) and 1600 (encoded) cycles, 5963 precursors in range
[489:22] Calibrating with mass accuracies 25 (MS1), 25 (MS2)
[489:23] RT window set to 0.505389
[489:23] Recommended MS1 mass accuracy setting: 13 ppm
[489:23] Searching decoys
[489:24] Main search
[489:24] Removing low confidence identifications
[489:24] Removing interfering precursors
[489:24] Training neural networks on 4696 target and 4026 decoy PSMs
[489:30] Training neural networks on 4693 target and 3754 decoy PSMs
[489:36] Number of IDs at 0.01 FDR: 3271
[489:36] Precursors at 1% peptidoform FDR: 1381
[489:36] Calculating protein q-values
[489:36] Number of genes identified at 1% FDR: 927 (precursor-level), 819 (protein-level) (inference performed using proteotypic peptides only)
[489:36] Quantification
[489:36] Precursors with scored PTMs at 1% FDR: 18 out of 36 considered
[489:36] Precursors with all scored PTM sites unoccupied at 1% FDR: 1363
[489:36] Precursors with PTMs localised (when required) with > 90% confidence: 17 out of 18
[489:36] XICs saved to D:\Soumyadeep\Tissue\out/tissue_diannreport_xic/B184H_T2.xic.parquet

[489:36] File #9/12
[489:36] Loading run D:\Soumyadeep\Tissue\Tissue Run 3 MS Data\280425_SD_CJ\B185F_T2.wiff
[491:02] Pre-processing...
[491:02] 1600 MS1 and 44800 MS2 scans in 1600 (inferred) and 1600 (encoded) cycles, 5963 precursors in range
[491:02] Calibrating with mass accuracies 25 (MS1), 25 (MS2)
[491:03] RT window set to 0.540045
[491:03] Recommended MS1 mass accuracy setting: 15 ppm
[491:04] Searching decoys
[491:04] Main search
[491:05] Removing low confidence identifications
[491:05] Removing interfering precursors
[491:05] Training neural networks on 4637 target and 4000 decoy PSMs
[491:11] Training neural networks on 4630 target and 3748 decoy PSMs
[491:17] Number of IDs at 0.01 FDR: 3235
[491:17] Precursors at 1% peptidoform FDR: 1476
[491:17] Calculating protein q-values
[491:17] Number of genes identified at 1% FDR: 947 (precursor-level), 855 (protein-level) (inference performed using proteotypic peptides only)
[491:17] Quantification
[491:17] Precursors with scored PTMs at 1% FDR: 8 out of 27 considered
[491:17] Precursors with all scored PTM sites unoccupied at 1% FDR: 1468
[491:17] Precursors with PTMs localised (when required) with > 90% confidence: 8 out of 8
[491:17] XICs saved to D:\Soumyadeep\Tissue\out/tissue_diannreport_xic/B185F_T2.xic.parquet

[491:17] File #10/12
[491:17] Loading run D:\Soumyadeep\Tissue\Tissue Run 3 MS Data\280425_SD_CJ\B212F_T2.wiff
[492:55] Pre-processing...
[492:55] 1600 MS1 and 44800 MS2 scans in 1600 (inferred) and 1600 (encoded) cycles, 5963 precursors in range
[492:55] Calibrating with mass accuracies 25 (MS1), 25 (MS2)
[492:57] RT window set to 0.520244
[492:57] Recommended MS1 mass accuracy setting: 14 ppm
[492:57] Searching decoys
[492:57] Main search
[492:58] Removing low confidence identifications
[492:58] Removing interfering precursors
[492:59] Training neural networks on 4740 target and 4129 decoy PSMs
[493:05] Training neural networks on 4733 target and 3789 decoy PSMs
[493:10] Number of IDs at 0.01 FDR: 3817
[493:10] Precursors at 1% peptidoform FDR: 2817
[493:10] Calculating protein q-values
[493:10] Number of genes identified at 1% FDR: 1007 (precursor-level), 841 (protein-level) (inference performed using proteotypic peptides only)
[493:10] Quantification
[493:10] Precursors with scored PTMs at 1% FDR: 31 out of 41 considered
[493:10] Precursors with all scored PTM sites unoccupied at 1% FDR: 2786
[493:10] Precursors with PTMs localised (when required) with > 90% confidence: 30 out of 31
[493:11] XICs saved to D:\Soumyadeep\Tissue\out/tissue_diannreport_xic/B212F_T2.xic.parquet

[493:11] File #11/12
[493:11] Loading run D:\Soumyadeep\Tissue\Tissue Run 3 MS Data\280425_SD_CJ\B214F_T2.wiff
[494:51] Pre-processing...
[494:51] 1600 MS1 and 44800 MS2 scans in 1600 (inferred) and 1600 (encoded) cycles, 5963 precursors in range
[494:51] Calibrating with mass accuracies 25 (MS1), 25 (MS2)
[494:52] RT window set to 0.51681
[494:52] Recommended MS1 mass accuracy setting: 15 ppm
[494:52] Searching decoys
[494:53] Main search
[494:53] Removing low confidence identifications
[494:54] Removing interfering precursors
[494:54] Training neural networks on 4628 target and 3965 decoy PSMs
[494:59] Training neural networks on 4623 target and 3739 decoy PSMs
[495:05] Number of IDs at 0.01 FDR: 3385
[495:05] Precursors at 1% peptidoform FDR: 1566
[495:05] Calculating protein q-values
[495:05] Number of genes identified at 1% FDR: 1010 (precursor-level), 879 (protein-level) (inference performed using proteotypic peptides only)
[495:05] Quantification
[495:05] Precursors with scored PTMs at 1% FDR: 13 out of 30 considered
[495:05] Precursors with all scored PTM sites unoccupied at 1% FDR: 1553
[495:05] Precursors with PTMs localised (when required) with > 90% confidence: 13 out of 13
[495:05] XICs saved to D:\Soumyadeep\Tissue\out/tissue_diannreport_xic/B214F_T2.xic.parquet

[495:06] File #12/12
[495:06] Loading run D:\Soumyadeep\Tissue\Tissue Run 3 MS Data\280425_SD_CJ\B229F_T2.wiff
[496:39] Pre-processing...
[496:39] 1600 MS1 and 44800 MS2 scans in 1600 (inferred) and 1600 (encoded) cycles, 5963 precursors in range
[496:39] Calibrating with mass accuracies 25 (MS1), 25 (MS2)
[496:40] RT window set to 0.519307
[496:40] Recommended MS1 mass accuracy setting: 15 ppm
[496:41] Searching decoys
[496:41] Main search
[496:41] Removing low confidence identifications
[496:42] Removing interfering precursors
[496:42] Training neural networks on 4572 target and 3950 decoy PSMs
[496:48] Training neural networks on 4569 target and 3742 decoy PSMs
[496:53] Number of IDs at 0.01 FDR: 3110
[496:53] Precursors at 1% peptidoform FDR: 1792
[496:53] Calculating protein q-values
[496:53] Number of genes identified at 1% FDR: 922 (precursor-level), 823 (protein-level) (inference performed using proteotypic peptides only)
[496:53] Quantification
[496:53] Precursors with scored PTMs at 1% FDR: 10 out of 18 considered
[496:53] Precursors with all scored PTM sites unoccupied at 1% FDR: 1782
[496:53] Precursors with PTMs localised (when required) with > 90% confidence: 10 out of 10
[496:54] XICs saved to D:\Soumyadeep\Tissue\out/tissue_diannreport_xic/B229F_T2.xic.parquet

[496:54] Cross-run analysis
[496:54] Reading quantification information: 12 files
[496:54] Quantifying peptides
[497:00] Quantification parameters: 0.383601, 0.0275203, 1.66336, 0.128056, 0.0622094, 0.0904893, 0.323597, 0.136739, 0.178614, 0.014778, 0.0158689, 0.0151559, 0.198748, 0.0131428, 0.137389, 0.012973
[497:03] Quantifying proteins
[497:03] Calculating q-values for protein and gene groups
[497:03] Calculating global q-values for protein and gene groups
[497:03] Protein groups with global q-value <= 0.01: 1132
[497:03] Compressed report saved to D:\Soumyadeep\Tissue\out\tissue_diannreport.parquet. Use R 'arrow' or Python 'PyArrow' package to process
[497:03] Site report saved to D:\Soumyadeep\Tissue\out\tissue_diannreport.site_report.parquet
[497:03] Saving precursor levels matrix
[497:03] Precursor levels matrix (1% precursor and protein group FDR) saved to D:\Soumyadeep\Tissue\out\tissue_diannreport.pr_matrix.tsv.
[497:03] Saving protein group levels matrix
[497:03] Protein groups matrix saved to D:\Soumyadeep\Tissue\out\tissue_diannreport.pg_matrix.tsv.
[497:03] Saving gene group levels matrix
[497:03] Gene groups matrix saved to D:\Soumyadeep\Tissue\out\tissue_diannreport.gg_matrix.tsv.
[497:03] Saving unique genes levels matrix
[497:03] Unique genes matrix saved to D:\Soumyadeep\Tissue\out\tissue_diannreport.unique_genes_matrix.tsv.
[497:03] Manifest saved to D:\Soumyadeep\Tissue\out\tissue_diannreport.manifest.txt
[497:03] Stats report saved to D:\Soumyadeep\Tissue\out\tissue_diannreport.stats.tsv
